# Sex-Specific Pathophysiological Signatures in Allometric Dosing-Controlled Bleomycin Acute Lung Injury Model

**DOI:** 10.64898/2026.03.10.710851

**Authors:** Samuel Gillman, Alice Ngu, Michael Lush, Nikolay Karpuk, Kevin M. Hu, Steven J. Lisco, Han-Jun Wang

**Author notes:** Both authors contributed equally to this work (Samuel Gillman, Alice Ngu). **Corresponding Author:** Han-Jun Wang, M.D., Department of Anesthesiology, University of Nebraska Medical Center, Omaha, NE 68198, USA.

## Abstract

**Introduction:** In acute lung injury (ALI), clinical data show that while mortality rates are similar between sexes, women require shorter ventilation times and intensive care unit stays than men, yet preclinical studies show conflicting sex-specific vulnerabilities. We reasoned that a hidden dosing bias may explain the inconsistency, as intratracheal bleomycin is scaled to body weight, even though lung mass grows more slowly than total body mass, so age-matched males, whose body mass outpaces lung growth, inevitably receive more drug per gram of lung than females.

**Methods:** We compared age-matched (12-week) and body-weight-matched (∼300g) Sprague-Dawley rats receiving intratracheal bleomycin (2.5mg/kg) or saline. Both cohorts underwent functional assessments (plethysmography, lung mechanics, arterial gases, histology) at day 7; weight-matched animals exclusively underwent mechanistic profiling (BALF analysis, cytokine multiplex, paired mRNA/miRNA-sequencing, immunoblotting).

**Results:** Males developed worse hypoxemia (PaO₂: age-matched p = 0.045; weight-matched p = 0.027) with higher respiratory rates (both cohorts p < 0.05). Weight-matched males showed greater compliance loss (p = 0.029), increased BALF protein (p = 0.008), and elevated IL-1β (p =0.005) and TNF-α (p = 0.017). RNA-sequencing identified 2,393 male versus 1,533 female differentially-expressed genes, with males activating complement-coagulation cascades while females enriched ECM-remodeling/BMP-signaling pathways. Males exhibited significant miR-672-3p suppression (p < 0.0001), inversely correlating with inflammatory targets. SERPINA3 and its upstream regulator STAT3 showed significantly higher induction in males (both p < 0.0001), whereas females exhibited higher BMPR2 protein levels (p = 0.009) and preserved IL-10 (p = 0.023).

**Conclusions:** Body-weight matching corrects unrecognized allometric bias affecting preclinical ALI sex-difference studies. Both cohorts demonstrated male vulnerability with worse hypoxemia and increased respiratory rates. Weight-matched molecular analyses revealed distinct responses: males showed significant miR-672-3p suppression with concurrent inflammatory mediator upregulation, including higher SERPINA3, IL-1β, and TNF-α. In contrast, females maintained higher miR-672-3p levels alongside elevated BMPR2/IL-10, suggesting that divergent post-transcriptional regulation contributes to functional differences and may inform sex-specific therapeutic strategies.

## Introduction

Acute lung injury (ALI) and its more severe manifestation, acute respiratory distress syndrome (ARDS), are characterized by the disruption of the alveolar-capillary barrier. This disruption leads to a triad of protein-rich pulmonary edema, surfactant dysfunction, and neutrophilic infiltration, culminating in hypoxemia and respiratory failure (1). Despite the adoption of lung protective ventilation and adjunctive rescue therapies, global outcomes remain poor. Mechanistic understanding gaps and limited targeted therapeutics persist as significant barriers to progress. In the 50-nation LUNG-SAFE cohort, hospital mortality ranged between 35-46% (2) with no significant difference between males and females, however, when matched for disease severity, females had shorter ventilation times and number of days in the ICU (3). However, shorter females are less likely to receive guideline-concordant low-tidal-volume ventilation (3), indicating that superior female outcomes cannot be ascribed to better care and suggest that sex-dependent pulmonary physiology contributes to recovery.

ALI models are deliberately reductionist, each capturing a distinct anatomic or mechanistic facet. Endothelial-dominant injuries (e.g., oleic-acid), epithelial-focused injuries (e.g., intratracheal bleomycin), biophysical trauma (e.g., high-strain ventilation), and innate immune triggers (e.g., lipopolysaccharide) collectively inform pathogenesis (4, 5).

Sex-specific findings across these models are discordant. In mice, males sustain greater neutrophilic alveolitis, impaired mechanics, robust inflammatory transcriptomes, and amplified matrix deposition following bleomycin-induced ALI (6–8). Hyperoxia triggers broader male transcriptomic activation (9), and a meta-analysis of sex differences in preclinical LPS studies reveals males developing a more severe phenotype (10). Conversely, female rats exhibit worse fibrosis and higher pro-collagen transcription following bleomycin-induced ALI (11). Because bleomycin causes direct epithelial apoptosis and pulmonary edema (12), it offers an ideal context to elucidate the basis of these conflicting findings. Brief hormone-manipulation experiments suggest that androgens amplify bleomycin injury (13).

A critical methodological thread and potential confound is the age matching of males and females. Between 10 and 14 weeks of age, Sprague-Dawley rats nearly double in body weight, while lung weight increases by only 20% (14–16). This disproportionate growth means that when bleomycin, which is dosed based on body weight, is administered intratracheally to male rats, they receive higher concentrations per gram of lung tissue than females of the same age. This size-related disparity can make male injury appear worse in age-matched approaches, emphasizing why body weight matching (or lung volume matching) is a critical consideration when studying sex differences in a bleomycin model of ALI. Indeed, investigations in select murine models of lung injury (LPS, *Streptococcus pneumoniae*, and influenza A) have demonstrated that the choice of dosing strategy (e.g., identical inoculum/dose versus body weight normalization versus lung-weight normalization) can significantly alter observed sex-specific outcomes and interpretation of sex differences in susceptibility and response (17).

To disentangle scaling artifacts from intrinsic sex biology, we implemented a dual-cohort approach. Alongside a conventional age-matched group of 12-week-old males and females, we included a novel bodyweight-matched (BW-matched) cohort where we used slightly older females (∼14 weeks) and younger males (∼10 weeks) so that both sexes averaged ∼300 g at dosing. Seven days following bleomycin-induced ALI, we comprehensively phenotyped both cohorts using non-invasive plethysmography, invasive resistance/compliance measurements, arterial blood gases, and immunohistochemistry. More detailed mechanistic studies (bronchoalveolar lavage, lung tissue cytokines, paired lung RNA/miRNA sequencing, and protein analyses) were performed only in the BW-matched cohort to eliminate size-related confounding during molecular comparison. This approach addresses two critical questions: Do sex-specific responses to bleomycin ALI persist when animals are matched for body weight, and what molecular mechanisms might explain these differences?

## Materials and Methods

All animal experimentation was approved by the Institutional Animal Care and Use Committee (IACUC) of the University of Nebraska Medical Center (UNMC) (Omaha, NE). Experiments were performed in accordance with the ARRIVE guidelines and the National Institutes of Health’s Guide for the Care and Use of Laboratory Animals (18, 19). Experiments were performed on male (Age-match ∼ 380g, BW-match ∼ 300g) and female (Age-match ∼ 280g, BW-match ∼ 300g) Sprague-Dawley rats purchased from Charles River Laboratories (Wilmington, MA, USA). Animals were housed on-site and acclimated for one week before experiments. Food and water were supplied *ad libitum*, and rats were kept on 12-h light/dark cycles.

### Induction of Acute Lung Injury (ALI)

ALI was induced via a single intra-tracheal (IT) instillation of bleomycin sulfate (Sigma-Aldrich, St. Louis, MO, USA) (2.5 mg/kg, ∼0.15 ml) under 3% isoflurane anesthesia. Immediately post-instillation, rats were ventilated with 1.5% isoflurane for 3 minutes to ensure uniform pulmonary distribution. Animals were monitored daily; those losing ≥20% of initial body weight were euthanized according to approved protocols. Sham animals (non-lung injury) received equal volume sterile saline (∼0.15mL)

### Acclimation to Whole Body Plethysmography (WBP) Chambers

Following the facility acclimation period, animals were weighed and placed into the Rat WBP standard chamber (DSI Buxco respiratory solutions, DSI Inc., St. Paul, MN, USA) for one hour on two consecutive days, totaling two hours of chamber acclimation per animal before data collection.

### Non-Invasive Functional Respiratory Parameters

After chamber acclimation, baseline recordings were obtained before bleomycin exposure. Animals were weighed and placed in the Rat WBP standard chamber for a 20-minute acclimation period, followed by a one-hour recording. Data was recorded using DSI FinePointe WBP software (Data Sciences International, St. Paul, MN, USA). Parameters measured included respiratory rate (RR), tidal volume (TV), minute volume (MV), inspiratory time (Ti), expiratory time (Te), relaxation Time (Tr), expiratory flow at 50% TV (EF50), peak inspiratory flow (PIF), peak expiratory flow (PEF), relative time to peak expiratory flow (Rpef), enhanced pause (Penh), and O_2_ Consumption (VO_2_).

### Invasive Airway Mechanics

Seven days following induction of lung injury we performed direct measurement of airway resistance (RI) and dynamic compliance (Cdyn) with the DSI Buxco Resistance/Compliance (RC) apparatus. Animals were anesthetized with an intraperitoneal injection of urethane (0.15g/0.1kg). After confirming the anesthetic depth, the trachea was cannulated with a 14-gauge cannula, secured with suture, and the animal was moved to the RC system. End-expiratory pressure was set at 3.0 cmH2O, and the ventilator was set at 90 breaths/minute. Resistance and compliance were recorded for 5 minutes with averages reported.

### Arterial Blood Gas

Seven days post-bleomycin ALI, blood (∼0.1 mL) was collected from the ventral tail artery to assess arterial blood gases. The animal was placed in a restrainer, and the tail was aseptically prepared with alternating alcohol and iodine prep pads. The artery was punctured with a 24G needle, and a small volume (∼0.1 mL) of blood was aspirated. The blood was analyzed via iSTAT (Abbott, Chicago, IL, USA).

### Immunohistochemistry (IHC)

At the seven-day timepoint after bleomycin, animals were euthanized and perfused with 4% paraformaldehyde (PFA). The lungs were then excised, stored in 4% PFA, and transferred to 70% ethanol after 48 hours. Lung tissues were processed at the UNMC Tissue Sciences Facility. Stains included hematoxylin & eosin (H&E), Masson’s trichrome (MTC), CD3, CD68, and myeloperoxidase (MPO). Imaging was performed with Aperio ScanScope XT (Aperio Technologies, Vista, CA). Analysis of each respective stain was performed with QuPath (20).

### Bronchoalveolar Lavage Fluid (BALF) Collection and Analysis

Animals were anesthetized with urethane (0.15g/0.1kg) and exsanguinated. The trachea was cannulated with a 14-gauge blunt-end catheter, and three 2 mL washes of lavage buffer (1× PBS + 0.5 mM EDTA) were performed. BALF was centrifuged for 6 minutes at 1,200 rpm. The supernatant was collected and stored at -80°C. Total BALF cell count was determined using a Countess 3 Automated Cell Counter (Thermo Fisher Scientific, Waltham, MA, USA). BALF protein concentration was measured via BCA Protein Assay Kit (Thermo Fisher Scientific).

### Lung Tissue Cytokine Profiling via Multiplex ELISA

Lungs from BW-matched male and female rats (sham and bleomycin) were harvested one week after treatment, homogenized, and analyzed using Quansys Q-Plex sample testing service (Logan, UT, United States). Samples were tested for IL-1α, IL-1β, IL-2, IL-4, IL-6, IL-10, IL-12, IFN-γ, and TNF-α. Samples were diluted with Quansys buffer at 1:2, 1:20, and 1:200 dilutions, with each measured in triplicate. Polypropylene low-binding 96-well plates were used for sample preparation. Total protein concentration was determined via absorbance at 280 nm using a NanoDrop ND-1000 spectrophotometer (Thermo Fisher Scientific, Wilmington, DE). Images were captured with a 270-second exposure on Q-View Imager LS, and light emission was measured as pixel density units by Q-View Software. The optimal dilution for samples was selected by the Q-view software, which finds the dilution where the pixel intensity values fall on the most linear portion of the standard curve. Results were normalized to total protein concentration and presented as pg/mg.

### mRNA- and miRNA-sequencing: Paired Library Preparation, Sequencing, and Bioinformatics Workflow

Paired mRNA and miRNA sequencing utilized services offered by LC Sciences (Houston, TX, USA). Total lung RNA was extracted from the BW-matched cohort only (five sham-male, five sham-female, five bleomycin-male, five bleomycin-female) with TRIzol (Invitrogen) and randomized for downstream processing. Integrity was verified on a Bioanalyzer 2100 (Agilent); every sample showed RIN ≥ 7.0. Two complementary libraries were prepared from the same RNA aliquot for each animal.

**mRNA (poly-A) libraries** were generated with the TruSeq-Stranded mRNA kit (Illumina). Poly-A RNA was captured with two rounds of oligo-dT magnetic-bead purification, chemically fragmented, converted to cDNA, end-repaired, A-tailed, ligated to indexed adapters, and amplified (15 cycles). Paired-end 150-bp reads were produced on an Illumina NovaSeq 6000 at LC Sciences (Houston, TX, USA), yielding a median of 7.9 G bases per sample with ≥97 % valid reads and Q30 > 98 %. Adapters and low-quality bases were removed with Cutadapt 1.10; quality was confirmed with FastQC 0.10.1. Clean reads were aligned to the *Rattus norvegicus* genome (Ensembl v112, Rnor_6.0) using HISAT2 2.0. Transcript assembly and gene-level quantification (FPKM) were performed with StringTie 1.3.4; counts were imported into DESeq2 (21) and edgeR (22) for differential expression. Genes with |log₂ fold-change| ≥ 1 and FDR < 0.05 were considered significant. Downstream GO and KEGG enrichment used clusterProfiler with Benjamini–Hochberg correction.

**miRNA libraries** were built from the remaining 1 µg RNA using the TruSeq Small-RNA kit (Illumina). Size-selected libraries (18–26 nt inserts) were single-end-sequenced (50 bp) on an Illumina HiSeq 2500, producing a median 9.2 million raw reads per sample. Filtering (adapter, low-quality, rRNA/tRNA/snoRNA) and mapping to miRBase v22 rat precursors were performed with ACGT101-miR v2023.1, allowing one mismatch in the mature region. Unmapped reads were BLASTed to Rnor_6.0; hairpins that met LC Sciences structural criteria (≥16 stem pairs, ΔG ≤ −15 kcal mol⁻¹, loop ≤ 20 nt) were annotated as novel miRNAs. Normalization used the median-ratio (DESeq) method; edgeR identified differentially expressed miRNAs (p < 0.05, |log₂ FC| ≥ 1). Putative targets were predicted by the intersection of miRanda and TargetScan-Rat (23, 24).

### cDNA synthesis and RT-qPCR

Tissue-extracted total RNA (1µg) was polyadenylated and reverse transcribed using microScript microRNA cDNA synthesis kit (Norgen Biotek Corporation 54410, ON, Canada). The miRNA expression was assessed by RT-qPCR using QuantiNova SYBR green PCR kit (Qiagen 208056, MD, USA), a universal reverse primer (Norgen Biotek Corporation 54410, ON, Canada), and miRNA-specific forward primers designed using miRprimer (25). RT-qPCR analysis was performed on StepOnePlus real-time PCR system (Applied Biosystems, MA, USA). The cycling conditions were 95°C for 2 minutes, followed by 35 cycles of 95°C for 5 seconds and 60°C for 10 seconds. U6 small nuclear RNA (snRNAs, NCBI Accession XR_005499917.1) was used as an endogenous reference gene for normalization of miRNA expression. The miRNA primers are shown in the supplementary materials (**Table S9**).

Tissue-extracted total RNA (500ng) was polyadenylated and reverse transcribed using miRCURY LNA RT kit (Qiagen 339340, MD, USA). The miRNA expression was assessed by RT-qPCR using miRCURY LNA SYBR green PCR kit (Qiagen 339345, MD, USA) and miRCURY LNA miRNA PCR assays (Qiagen 339306, MD, USA). RT-qPCR analysis was performed on StepOnePlus real-time PCR system (Applied Biosystems, MA, USA). The cycling conditions were 95°C for 2 minutes, followed by 40 cycles of 95°C for 10 seconds and 56°C for 1 minute. U6 small nuclear RNA was used as an endogenous reference gene for normalization of miRNA expression. The Geneglobe ID of miRNA PCR primer sets is shown in Supplementary Table 9.

### Immunoblotting

The following antibodies were used for Western blotting: SERPINA3 (PA5-86755, Thermo Fisher), Vimentin (MA511883, Thermo Fisher Scientific) STAT3 (ab76315, ABCAM), pSTAT3 (ab68153, ABCAM), MFGE8 (PA5109955, Life Technologies), and BMPR2 (MA515827, Life Technologies). Protein was extracted from lung tissue samples, quantified, and equal amounts were loaded onto SDS-PAGE gels. Following electrophoresis, proteins were transferred to PVDF membranes. Membranes were blocked and incubated with primary antibodies overnight at 4°C, followed by species-appropriate HRP-conjugated secondary antibodies. Immunoreactive bands were visualized using enhanced chemiluminescence and imaged with the iBright FL750 Imaging System (Thermo Fisher Scientific). Band intensities were quantified by densitometry using Image Studio Software (LI-COR Biosciences). Results were normalized to appropriate loading controls.

### Cell Culture and Transfection with Synthetic Nucleic Acid

RAW264.7 cells were cultured in complete media, which comprised RPMI 1640 medium (Thermo Fisher Scientific 11875085, MA, USA) with the final concentration of 10% FBS (ATCC 30-2020, VA, USA) and 1% Penicillin-Streptomycin (Thermo Fisher Scientific 15140148, MA, USA) at 37°C and 5% CO2. Around 4 × 106 cells were seeded in a T75 flask (Corning 430641U, NY, USA) overnight. 30µl of transfection reagent (Thermo Fisher Scientific 13778150, MA, USA) was diluted with 470µl of OptiMEM (Thermo Fisher Scientific 31985070, MA, USA) while synthetic nucleic acids (Table 1), which had been dissolved in molecular free water, were diluted with OptiMEM to a working concentration of 2uM. Diluted transfected reagent (500µl) and 2uM synthetic nucleic acids (500µl) were mixed together and incubated at room temperature for around 5 minutes. The total volume of the mixture (1000 µl) was added dropwise to the seeded cells, and at this point, the final concentration of synthetic nucleic acids in the complete medium was 100nM. The cells were later incubated at 37°C and 5% CO2 overnight. Without changing the medium, a final concentration of 50 ng/ml LPS (InvivoGen tlrl-eblps LEB-39-03, CA, USA) was added to the cells for approximately 4 hours. The media supernatant was collected after centrifugation at 300xg for 10 minutes. The protein concentration of the media supernatant was measured using the BCA protein assay (Thermo Fisher Scientific 23225, MA, USA) according to the manufacturer’s instructions.

**Table 1.**
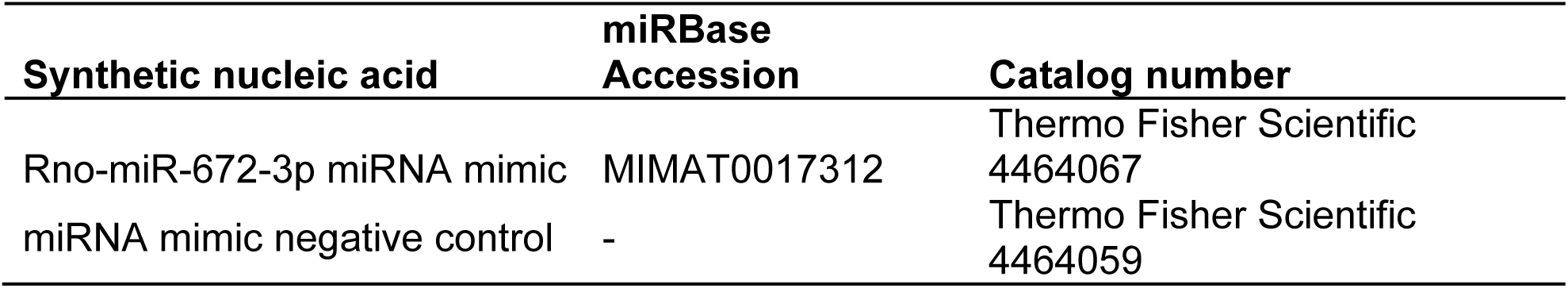
List of synthetic nucleic acids transfected to RAW264.7 cells.

Cytokine profiling of the media supernatant was performed using a Multiplex ELISA assay through the Quansys Q-Plex sample testing service (Logan, UT, USA). Results were normalized to total protein concentration and presented as pg/mg.

### Statistical Analysis

Comparisons between treatment and sex groups were analyzed via two-way ANOVA with Tukey’s multiple comparisons test using GraphPad Prism (version 10.5.0) (San Diego, CA, USA). Data are presented as mean ± SEM unless otherwise specified. Statistical significance was established at p < 0.05.

## Results

### Sex Differences in Non-Invasively and Invasively Measured Lung Mechanics

#### Non-invasive Lung Mechanics

Whole Body Plethysmography (WBP) was employed to capture respiratory metrics at baseline and post-bleomycin-induced ALI. Injury resulted in significant alterations in breathing patterns measured at one week (**Figure 1A**), with respiratory rate (RR) representing the most perturbed non-invasively derived metric. In age and weight-matched cohorts, males (both p < 0.0001) and females (both p < 0.0001) demonstrated a significant increase in RR following bleomycin compared to controls, with male animals having significantly higher RR than female animals post-injury across both cohorts (both p < 0.05) (**Figure 1B**). Tidal volume per kilogram (TV/kg) decreased significantly in both sexes post-bleomycin across both cohorts, with females demonstrating higher TV/kg compared to males under baseline and injury conditions in both cohorts (**Figure 1C**). Minute ventilation-per-kilogram (MV/kg) was significantly elevated following bleomycin in both sexes in the age-match (both p < 0.001) and BW-match cohorts (both p < 0.001) (**Figure 1D**). No sex differences in MV/kg were observed except under baseline conditions in the age-match cohort, where females had a higher MV/kg than males (p = 0.03). Mid-tidal expiratory flow (EF50) increased post-injury in both cohorts. In the age-matched cohort, EF50 in males increased compared to baseline in both males and females (both p < 0.0001), with males having a higher EF50 post-injury compared to females (p < 0.0001) (**Figure 1E**). BW-matched males and females also showed increased EF50 compared to respective shams (both p < 0.001), though no sex difference was observed post-injury (**Figure 1E**). Minimal changes in enhanced pause (Penh) were noted, with only baseline sex difference in the age-match cohort observed, where males had higher Penh than females (p = 0.01) (**Figure 1F**). Relative time to peak expiratory flow (Rpef) was decreased significantly post-injury in females in age and BW-match cohorts (both p < 0.05) (**Figure 1G**). Other WBP-derived metrics, including peak expiratory flow, peak inspiratory flow, and normalized oxygen consumption, also showed variable alteration depending on sex and treatment across cohorts (**Figure S1A-C**). Collectively, these results demonstrate that both sexes respond to bleomycin with altered respiratory mechanics and breathing patterns across both cohorts, with males exhibiting higher breathing frequencies compared to females in age- and body-weight-matched cohorts, as well as higher EF50 in age-matched cohorts.

**Figure 1.**
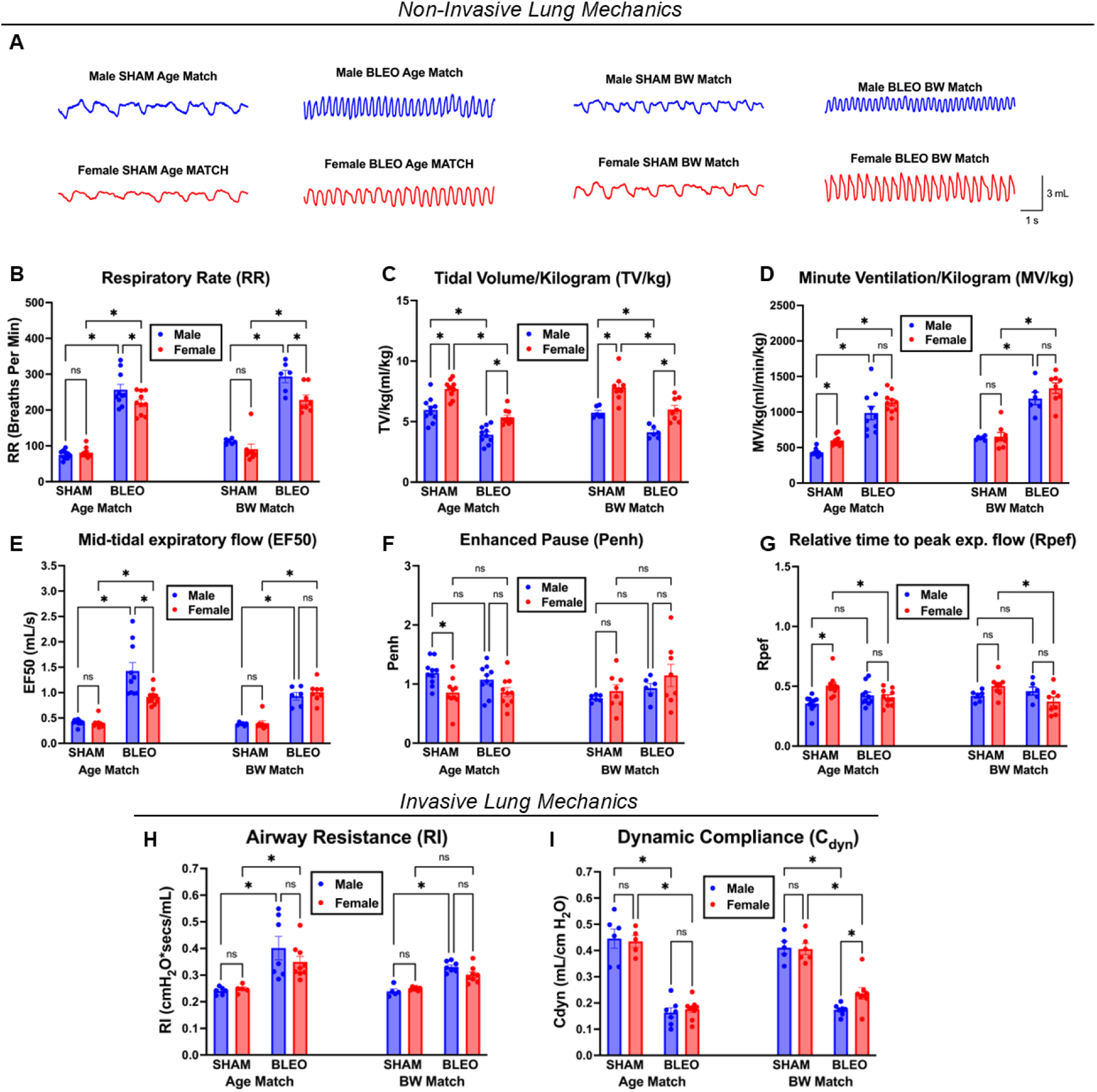
Sex differences in lung function following bleomycin-induced acute lung injury (ALI). (**A**) Representative waveform traces demonstrating altered breathing patterns in SHAM and bleomycin-treated (BLEO) male and female rats in both matching approaches. (**B-G**). Non-invasive whole-body plethysmography (WBP) measurements showing (**B**) respiratory rate (RR, breaths/min), (**C**) tidal normalized to body weight (TV/kg, mL/kg), (**D**) minute ventilation normalized to body weight (MV/kg, ml/min/kg), (**E**) mid-tidal expiratory flow (EF50, mL/s), (**F**) enhanced pause (Penh), and (**G**) relative time to peak expiratory flow (Rpef). Invasive lung mechanics measurements of (**H**) airway resistance (RI, cmH_2_O·s/mL) and (**I**) dynamic compliance (Cdyn, mL/cmH_2_O). Data are presented as mean ± SEM. For B-G, age matched cohort: n = 10 per group; BW matched group: n = 6 males, n = 8 females. For H-I, age matched group: n = 5-7 for SHAM, n = 7-9 for BLEO; BW-matched group: n = 5-6 for SHAM, n = 7-8 for BLEO. *p < 0.05, ns = not significant. Statistical analysis performed using two-way ANOVA with Tukey’s multiple comparisons test.

#### Invasive Lung Mechanics

Direct airway mechanics demonstrated robust bleomycin-induced lung injury across both cohorts. Airway resistance increased significantly in males (p < 0.0001) and females (p = 0.007) of the age-matched cohort following bleomycin, compared to their respective shams (**Figure 1H**). In the body weight matched cohort, only males showed a significant increase in airway resistance post-bleomycin (p = 0.02). No sex differences were detected under both baseline and injury conditions.

Dynamic compliance decreased significantly in age-matched males and females (both p < 0.0001) post-bleomycin compared to sham animals (**Figure 1I**). These findings were replicated in the body-weight-matched cohort, with males and females (both p < 0.001) exhibiting decreased compliance following lung injury. However, only the body weight-matched cohort revealed significant sex differences, with males exhibiting greater compliance loss than females (p = 0.029), whereas age-matched animals showed no sex difference (p = 0.633) (**Figure 1I**).

In summary, direct measurements of airway resistance and dynamic compliance revealed that significant sex differences in mechanical dysfunction only became evident when animals were matched for body weight, underscoring the importance of dosing normalization in detecting intrinsic biological disparities.

### Arterial blood gases demonstrate greater hypoxemia in males

Bleomycin-induced ALI resulted in a broad disturbance of oxygenation and acid-base balance across both cohorts, yielding sex-specific findings.

In age-matched and BW-matched cohorts, partial pressure arterial oxygen (PaO₂) significantly decreased in males and females post-bleomycin (all p< 0.001). In both cohorts, injured males exhibited significantly lower PaO₂ than injured females (age-match p = 0.045; BW-match p = 0.027) (**Figure 2A**). Blood oxygen saturation (sO₂) was significantly decreased in males in both cohorts compared to females post-bleomycin (both p < 0.01 vs. respective shams) (**Figure 2B**).

**Figure 2.**
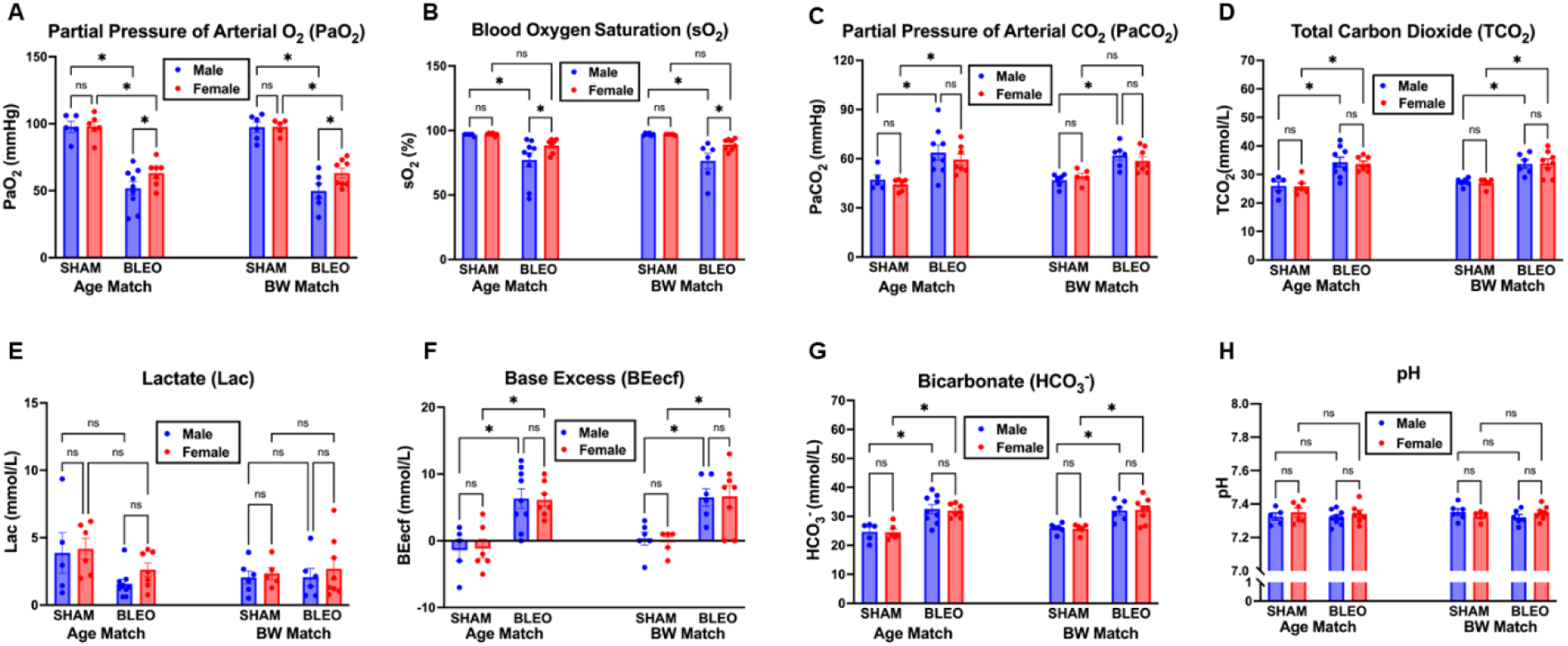
Blood gas analysis reveals sex-dependent differences in hypoxemia severity following bleomycin-induced ALI. Arterial blood gas parameters were measured in male and female rats 7 days after intratracheal instillation of saline (SHAM) or bleomycin (BLEO). (**A**) Partial pressure of oxygen (PaO_2_, mmHg) demonstrates significant hypoxemia in BLEO-treated rats, with female rats maintaining higher PaO_2_ levels compared to males. (**B**) Oxygen saturation (SO_2_, %) confirms impaired oxygenation following bleomycin injury. (**C**) Partial pressure of carbon dioxide (PaCO_2_, mmHg) and (**D**) Total carbon dioxide (TCO_2_, mmol/L) levels. (**E**) Lactate levels (mmol/L) are paradoxically decreased in BLEO groups. (**F**) Base excess (mmol/L) demonstrates acid-base disturbances. (**G**) Bicarbonate (HCO_3 -_, mmol/L) concentration indicating metabolic compensation. (**H**) Blood pH remains relatively stable despite respiratory changes. Data are shown for both age-matched and body weight (BW)-matched cohorts. Male rats are represented by blue bars and female rats by red bars. Female BLEO-treated rats exhibit less severe hypoxemia compared to male BLEO-treated rats in both matching strategies. Data are presented as mean ± SEM (n = 5-9 per group). *p < 0.05, ns = not significant. Statistical analysis performed using two-way ANOVA with Tukey’s multiple comparisons test.

Arterial CO_2_ (PaCO_2_) rose significantly after injury. In the age-matched cohort, PaCO₂ was heightened in both males and females following bleomycin ALI (p < 0.02) (**Figure 2C**). Similar elevations were seen in male BW-matched animals (p = 0.015), though females did not show a significant increase (p = 0.21) (**Figure 2C**). Total CO₂ (TCO₂) increased in both sexes and cohorts (all p < 0.01) (**Figure 2D**). No sex difference was detected for either parameter within the sham or the bleomycin groups.

Consistent with respiratory acidosis, serum bicarbonate (HCO_3_ ^-^) and base excess (BEecf) rose robustly in both male and female animals following bleomycin (all p < 0.05). (**Figure 2G & F**). The magnitude of metabolic compensation did not differ between males and females.

Systemic lactate remained within the low-millimolar range and was unaffected by sex or injury. Arterial pH consequently showed only minor, non-significant fluctuations in neutrality. (**Figure 2E & H**).

Taken together, bleomycin-induced ALI caused significant hypoxemia that was most severe in males across both age- and body weight-matched cohorts.

### Immunohistochemistry

To elucidate the cellular and structural basis underlying these sex-specific physiological impairments, we performed a comprehensive immunohistochemical analysis of lung tissue harvested at the seven-day timepoint in both experimental cohorts.

Histopathological Analysis Reveals Matching Strategy-Dependent Patterns of Inflammatory and Fibrotic Responses

#### Neutrophil Infiltration

Myeloperoxidase (MPO) staining revealed variable neutrophilic infiltration patterns following bleomycin-induced ALI across the two experimental cohorts (**Figure 3A**). In the age-matched cohort, neutrophil density showed non-significant increases in both sexes (males p = 0.376, females p = 0.058). No significant sex differences were observed within sham (p = 0.569) or bleomycin-treated groups (p = 0.654).

**Figure 3.**
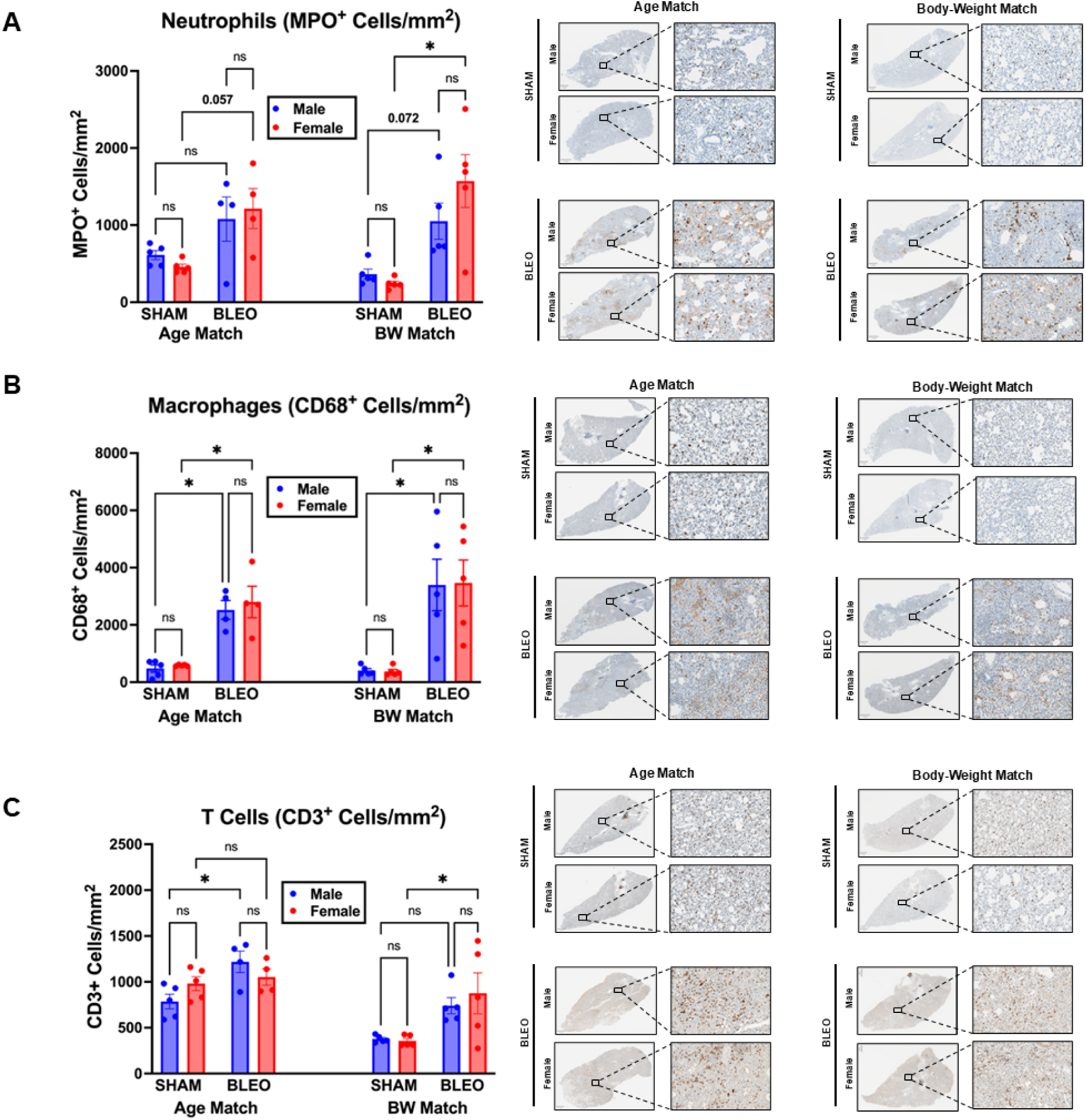
Histological assessment of lung immune cell infiltration post-bleomycin induced ALI. (**A**) Neutrophil infiltration, measured by myeloperoxidase (MPO)+ cells density (cells/mm^2^), demonstrates an acute inflammatory response, notable accumulation in the injured lung, especially in BW-matched female lungs. (**B**) Macrophage density, evaluated by CD68+ density (cells/mm^2^) increased in both sexes following lung injury in both age and BW matched cohorts. (**C**) CD3+ T-lymphocyte density (cells/mm^2^) was significantly increased in age-matched males and BW-matched females. Data is presented as mean ± SEM. Age match cohort: n = 4-5 per group; BW match cohort: n =5. *p < 0.05, ns = not significant. Statistical analysis performed using two-way ANOVA with Tukey’s multiple comparisons test.

The body weight-matched cohort revealed different patterns of neutrophilic response. While males showed a trend toward increased neutrophil infiltration (p = 0.073), females demonstrated a significant 6.5-fold increase (p < 0.001). Injured females exhibited a trend toward greater neutrophil density compared to injured males (p = 0.063). No sex differences were observed in the sham groups (p = 0.651).

#### Macrophage Accumulation

CD68 immunostaining demonstrated marked macrophage accumulation in response to bleomycin injury across both cohorts (**Figure 3B**). In the age-matched cohort, macrophage density increased significantly in both males (p = 0.037) and females (p = 0.021). In the BW-matched cohort, both males and females increased lung macrophage infiltration post-bleomycin (both p < 0.001) (**Figure 3B**). No significant sex differences were detected within sham or bleomycin-treated groups in either cohort.

#### T Cell Infiltration

CD3 staining revealed distinct patterns of T cell infiltration that varied by matching strategy (**Figure 3C**). In the age-matched cohort, males showed significant T cell accumulation following bleomycin injury (p = 0.048), while females showed no significant change (p = 0.969). The body weight-matched cohort displayed a different pattern. Females exhibited significant T cell infiltration (p = 0.008), while males showed only a trend toward increased T cell density (p = 0.094) (**Figure 3C**). No sex differences were observed within either the sham or bleomycin-treated groups.

#### Matrix Deposition

Masson’s Trichrome staining quantified collagen deposition seven days post-injury (**Figure S2**). In the age-matched cohort, bleomycin induced significant collagen deposition in both sexes (both p < 0.0001). Males exhibited significantly greater fibrotic burden compared to females (p = 0.005). The body weight-matched cohort also showed significant fibrosis in both sexes (both p < 0.001). However, in this cohort, females demonstrated significantly greater collagen deposition than males (p = 0.04).

Overall, patterns of inflammatory cell infiltration and early matrix deposition were influenced by both sex and matching strategy, with the direction and magnitude of sex differences in neutrophil, T cell, and collagen responses sometimes reversing depending on whether animals were age- or body weight matched.

Collectively, our functional, mechanical, and histopathological investigations suggest that intrinsic sex-specific differences exist in the severity and nature of bleomycin-induced lung injury, with these distinctions being clarified substantially by adopting a body-weight-matched approach. To further elucidate the molecular underpinnings of these observed sex-specific physiological and cellular responses, and to distinguish intrinsic biological vulnerability from confounding size-related effects, we next undertook an unbiased multi-omics analysis restricted to the body-weight-matched cohort.

### Bulk-RNA Sequencing

To elucidate the transcriptional mechanisms driving these sex differences and to uncover potential upstream regulatory pathways, we performed comprehensive parallel bulk RNA and miRNA sequencing of lung tissue from animals in the body-weight-matched cohort. This integrative transcriptomic analysis allowed for unbiased identification of sex-specific gene expression signatures and their potential miRNA regulators.

#### Sex-Specific Transcriptomic Responses to Bleomycin

Comparative analysis of bleomycin-induced ALI transcriptional responses demonstrated marked sex differences in both magnitude and directionality (**Figure 4B, Fig S4A-C**). Males exhibited significantly greater transcriptional perturbation with 2,393 differentially expressed genes (DEGs) (1,602 upregulated, 791 downregulated) versus 1,533 DEGs in females (1,129 upregulated, 404 downregulated) (|log_2_FC| ≥ 1, FDR < 0.05) (**Figure S4A-C**). Direct comparison of bleomycin-treated animals identified 826 DEGs (307 female-enriched, 519 male-enriched) (**Figure 4B-C**). Baseline sex differences were minimal, with only 262 DEGs between sham-treated animals (51 female-enriched, 211 male-enriched) (**Figure S4C**), indicating that observed transcriptional divergence primarily reflects sex-specific injury responses rather than constitutive sexual dimorphism.

**Figure 4.**
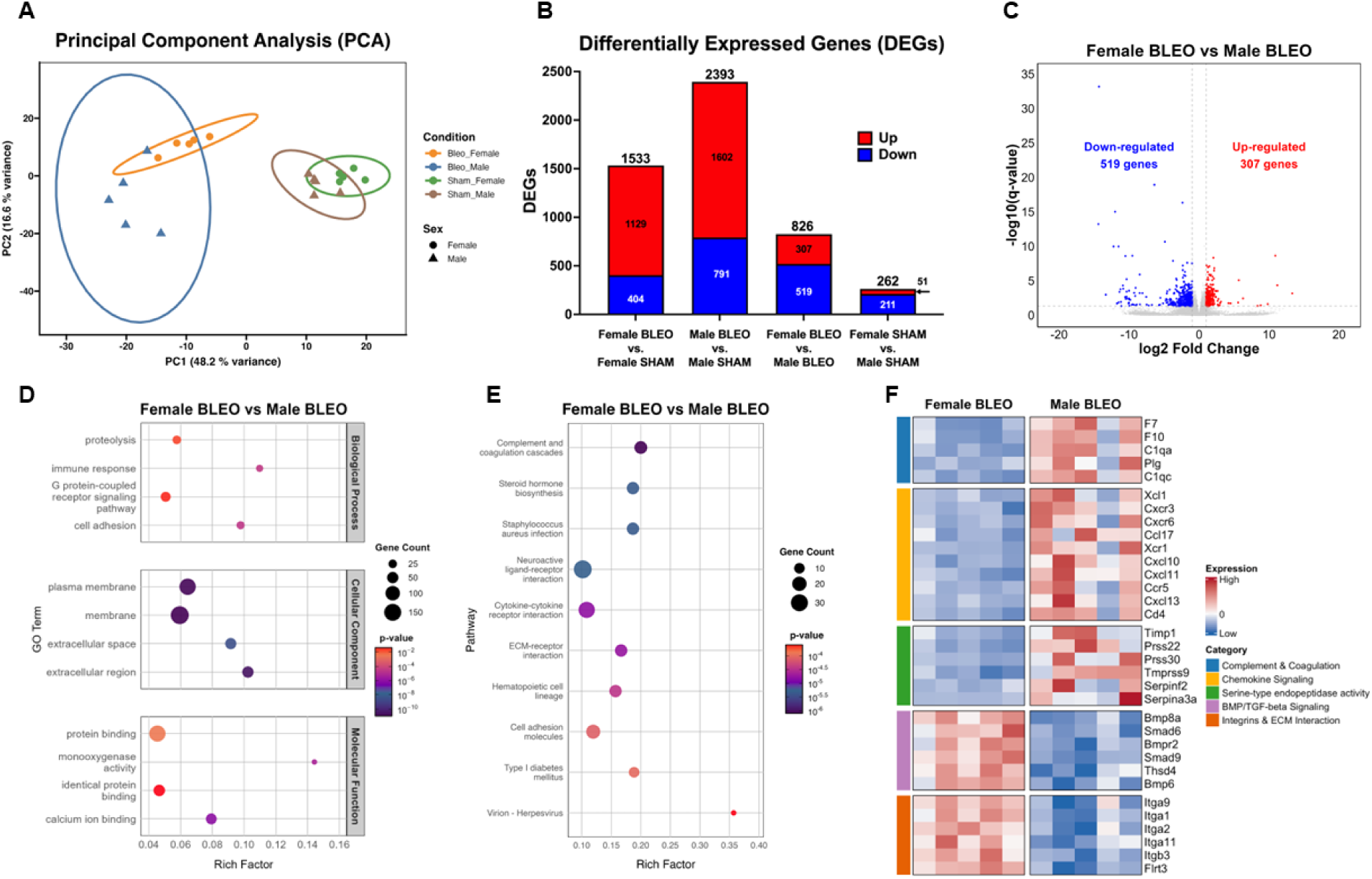
Sex-biased lung transcriptomic responses to bleomycin-induced ALI. (**A**) Principal component analysis of 20 lung RNA-seq libraries (5 sham male, 5 sham female, 5 bleomycin treated male, 5 bleomycin treated females) separates samples chiefly by injury along PC1 (48.2% of variance) and by sex along PC2 (16.6%). (**B**) Plot summarizing differentially expressed genes (DEGs; log_2_FC > |1|, q-value < 0.05). Bleomycin-induced ALI produced 2,393 DEGs in males and 1,533 females compared to their respective shams, whereas direct comparison under injury conditions identified 826 DEGs. (**C**) Volcano plot for Female BLEO vs Male BLEO highlights the 519 male-enriched (blue) and 307 female-enriched (red) genes that meet significance criteria. (**D**) Gene ontology (GO) enrichment of the 826 DEGs between Female BLEO vs Male BLEO mapped to GO terms across function classes, including biological processes terms linked to proteolysis, immune response, chemokine signaling, cell component terms associated with membrane and extracellular space, and molecular functions associated with protein and calcium ion binding. (**E**) KEGG analysis identified enrichment in pathways including complement coagulation cascades, cytokine-cytokine receptor interaction, and ECM-receptor interaction. (**F**) Heatmap of selected genes, grouped by functional theme, showing males have higher expression of coagulation factors (*F7, F10*), complement components (*C1qa, C1qc*), chemokines and chemokine receptors (*Cxcl10, Cxcl11, Cxcl13, Cxcr3*), endopeptidase activity components (*Serpina3a, Timp1*), whereas females displayed higher levels of BMP signaling mediators (*Bmpr2, Bmp8a, Smad9, Smad6*) and integrin/ECM genes (*Itga1/2/9/11, Flrt3*). Expression levels reflect gene-wise Z-scores (red = high, blue = low). All data derive from body-weight matched cohort (n = 5 biological replicates per sex and treatment).

Gene Ontology (GO) analysis of these 826 DEGs (**Figure 4D**) began to define this polarity. The biological processes with the most divergent expression patterns included proteolysis (GO:0006508) and immune response (GO:0006955). Cellular components included both membrane and extracellular space (GO:0016020, GO:0005615), and molecular functions included protein binding (GO:0005515).

KEGG pathway analysis further highlighted the contrast. Direct comparison between females and males revealed divergent pathways, including: cytokine-cytokine receptor interaction (ko04060), complement and coagulation cascades (ko04610), and ECM-receptor interaction (ko04512) (**Figure 4E**). A more granular analysis of the DEGs comprising these divergent functional categories and pathways identified male lungs selectively upregulating chemokines (*Cxcl10*, *Cxcl11*, *Ccl17*), complement/coagulation factors, (*F7*, *Plg*, *C1qa*), and components of the acute-phase protease inhibitor system (*Timp1*, *Serpina3a*, *Tmprss9*), whereas females induced BMP signaling genes (*Bmpr2*, *Smad9*, *Bmp6*) and an integrin scaffolding proteins (*Itga1/2/9/11*, *Itgb3*, *Flrt3*) (**Figure 4F**).

Notable findings from within-sex analyses showed females with enriched GO and KEGG themes related to extracellular matrix organization and cell-cycle progression, while males were more focused on immune pathways, specifically cytokine induction and viral response modules. (**Figures S4D-L**). A complete list of all DEGs across comparisons is provided in the supplementary materials (**Table S1-S4**).

### Lung microRNA Sequencing and Molecular Integration

We performed parallel RNA-miRNA sequencing to comprehensively and unbiasedly examine the transcriptional and post-transcriptional landscape of lung tissue in female and male rats following bleomycin ALI. Male animals displayed the greatest miRNA perturbation following lung injury with 112 miRNAs differentially expressed (75 upregulated, 37 downregulated) (**Figure 5A**; |log_2_FC| ≥ 1 p < 0.05). Female animals demonstrated markedly fewer differentially expressed miRNAs, with 66 total miRNAs (45 upregulated, 21 downregulated). Direct comparison between female and male animals’ post-injury revealed 54 differentially expressed miRNAs (8 upregulated, 46 downregulated). Baseline comparison between female and male miRNA profiles turned up 23 miRNAs (6 upregulated, 17 downregulated). A full list of significantly differentially expressed miRNAs is available in the supplementary materials (**Table S5-S8**).

**Figure 5.**
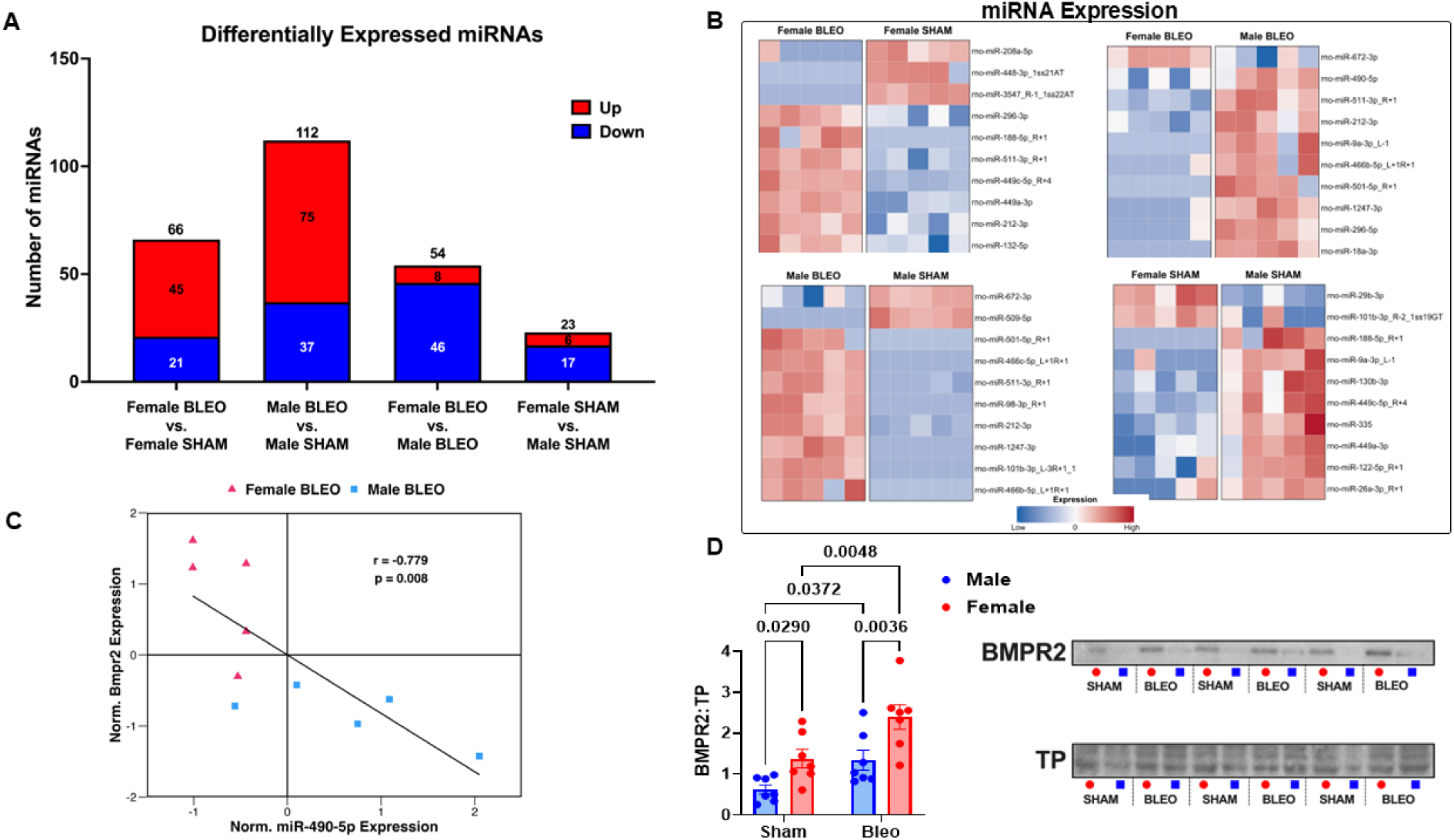
Sex-specific microRNA (miRNA) program and post-transcriptional regulation of BMPR2 after bleomycin-induced ALI. (**A**) miRNA-seq identified markedly different miRNA response magnitudes among the four comparisons (|log_2_FC|≥1, p < 0.05). Bleomycin lung injury produced 112 differentially expressed miRNAs in males (75 up, 37 down) and 66 differentially expressed miRNAs in females (45 up, 21 down) relative to their shams, whereas direct comparison of injured lungs yielded 54 sex-specific miRNAs (8 up, 46 down) and baseline dimorphism accounted for 23 miRNAs (6 up, 17 down). (**B**) Heatmaps show a subset of differentially expressed miRNAs within each comparison. Direct comparison between injured females and males highlights miR-511-3p_R+1, miR-490-5p, and miR-212-3p being higher in post-bleomycin males, while miR-672-3p was higher in females compared to males post-bleomycin. (**C**) *in silico* prediction identified Bmpr2 as a target of miR-490-5p. Leveraging the paired bulk and small RNA sequencing we performed, we discovered that within bleomycin-injured lungs, miR-490-5p and Bmpr2 expression is inversely correlated (Pearson r = -0.779, p = 0.008; red triangles = female, blue squares = male), consistent with the miRNA target relationship. (**D**) Immunoblotting confirms sex-divergent BMPR2 protein expression, baseline BMPR2 expression were significantly higher levels in females (p = 0.03), and levels of BMPR2 post-injury were significantly higher in females than males (p = 0.004). BMPR2 band intensity was first normalized to correspond to total protein (TP) intensity, then expressed as a ratio to the mean of the pooled sham samples. Bars represent mean ± SEM (n = 6 per group). Statistical analysis performed using two-way ANOVA with Tukey’s multiple comparisons test.

Among the sex-biased miRNAs, miR-490-5p showed significant downregulation in females versus males following bleomycin-induced lung injury (log_2_FC = -2.40) (**Figure 5B**). This was particularly intriguing as miR-490-5p is predicted to target *Bmpr2* (TargetScan score = 88), a critical component for vascular endothelial integrity and the BMP/TGF-β signaling pathway. Correlation analysis of *Bmpr2* expression and miR-490-5p expression revealed a negative correlation with females having a lower expression of miR-490-5p and a higher expression of *Bmpr2* compared to males (**Figure 5C).** Protein validation of BMPR2 revealed a baseline trend towards females having higher BMPR2 levels compared to males (p = 0.09), with females having significantly higher BMPR2 induction following lung injury compared to males (p = 0.009) (**Figure 5D**).

rno-miR-672-3p emerged as a noteworthy differentially expressed miRNA, a sex-specific regulator with opposing expression patterns between sexes (**Figure 6A**). In males, miR-672-3p was significantly downregulated following bleomycin injury (log_2_FC = -2.46, p = 0.0003), while females showed no significant change (log_2_FC = 0.375, p = 0.202). Direct comparison revealed that miR-672-3p was significantly higher in bleomycin-treated females compared to bleomycin-treated males (log_2_FC = 2.21, p = 0.006).

**Figure 6.**
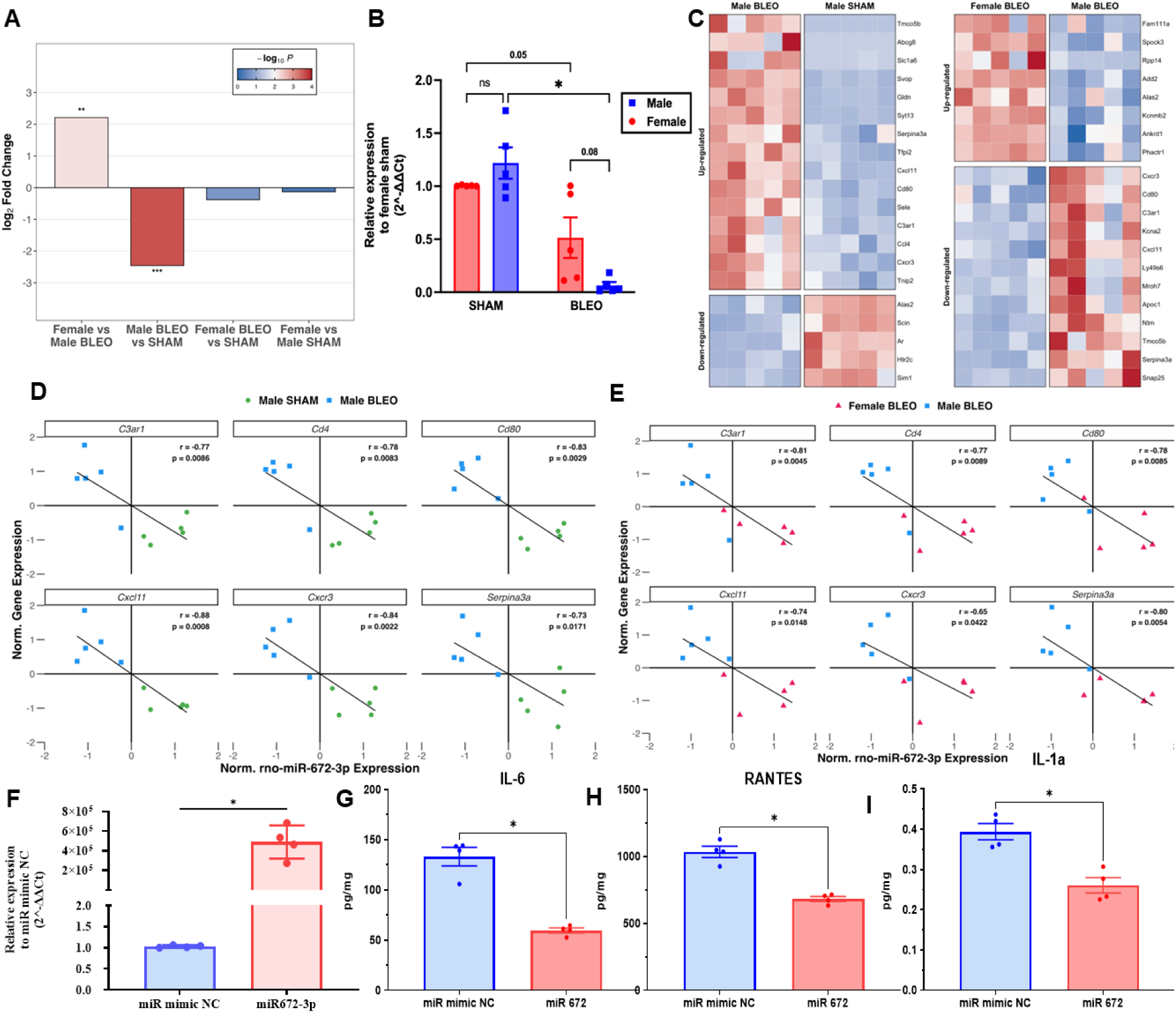
Sex-specific miR-672-3p expression association with a pro-inflammatory module after bleomycin-induced ALI. (**A**) miRNA-seq log_2_ fold-changes for miR-672-3p across the four group comparisons. miR-672-3p expression is significantly higher in Female BLEO than Male BLEO (+2.21 log_2_FC, p = 0.006), but significantly suppressed in Male BLEO versus SHAM (-2.46 log_2_FC, p = 0.0003). (**B**) RT-qPCR validation (2^ΔΔCt, normalized to U6 and ratioed to the pooled Female-Sham) reveals similar findings as sequencing results. While females show a trend towards reduced miR-672-3p expression post-bleomycin (p = 0.05), males display a robust decrease in miR-672-3p compared to their shams (p < 0.0001), and though non-significant, females retained higher levels of miR-672-3p post-injury (p = 0.08). (**C**) Heatmap of predicted miR-672-3p targets (e.g., *Cxcl11*, *Serpina3a*) are upregulated in males (**D**). Within-sex correlations (Male BLEO vs Male SHAM). Normalized miR-672-3p expression is plotted against six target genes; Pearson r values range from -0.73 to -0.88 (all p ≤ 0.02). (**E**) Inverse relationships persist among those six inflammatory mediators when the higher miR-672-3p expression in females correlates with lower expression of inflammatory mediators compared to males (r = -0.65 to -0.81; all p < 0.05). **(F)** Successful transfection of miR-672-3p in RAW264.7 cells. **(G–I)** Transfection of miR-672-3p reduces the production of pro-inflammatory cytokines, including IL-6, IL-1α, and RANTES, in LPS-treated RAW264.7 cells.

qPCR validation revealed a more nuanced pattern than the sequencing data suggested (**Figure 6B**). Both sexes showed suppression of miR-672-3p expression following ALI compared to their respective sham controls. Male animals demonstrated the most dramatic reduction (p < 0.0001), with approximately 95% suppression relative to sham. Females also showed downregulation (approximately a 50% reduction, p = 0.0516), although the magnitude was substantially less pronounced. The direct comparison between injured males and females showed a trend toward significance (p = 0.0801), indicating high retention of miR-672-3p expression in injured females compared to males.

Target prediction identified 242 potential miR-672-3p targets among male bleomycin versus sham and female versus male bleomycin DEGs, including high-confidence targeting of inflammatory mediators: *Cxcr3, Cxcl11*, *C3ar1*, *Cd80*, *Cd4*, and *Serpina3a*, all of which were upregulated in males in the parallel RNA-seq performed (**Figure 6C**). The biological coherence of these targets, which encompass chemokine signaling, complement activation, and acute-phase responses, suggests a functional significance.

To examine whether predicted miR-672-3p targets showed expression patterns consistent with mRNA expression, we analyzed correlations between miR-672-3p and target gene expression. In males, comparing injury to baseline, miR-672-3p showed negative correlations with inflammatory mediators including *Cxcl11* (r = -0.88, p = 0.0008), *Cxcr3* (r = -0.84, p =0.0022), *Cd80* (r = -0.83, p = 0.0029), *Cd4* (r = -0.78, p = 0.0083), *Serpina3*a (r = -0.73, p = 0.017), and *C3ar1* (r = -0.77, p = 0.0086) (**Figure 6D**). These inverse relationships were also observed in the sex comparison with similar correlation strengths: *C3ar1* (r = -0.81, p = 0.0045), *Serpina3a* (r = -0.80, p = 0.0054), *Cd80* (r = -0.78, p = 0.0085), *Cxcl11* (r = -0.78, p = 0.0148), *Cxcr3* (r = -0.65, p = 0.042), and *Cd4* (r = -0.77, p = 0.0089) (**Figure 6E**). To examine a potential cause–effect relationship between **miR-672-3p** and immune cell function, RAW264.7 cells were transfected with either miR-672-3p or a negative control (miR mimic NC) and subsequently treated with LPS. Cytokine levels in the culture supernatants were then measured and compared between the miR-672-3p and miR mimic NC groups. **Figure 6F** demonstrates successful transfection of miR-672-3p in RAW264.7 cells. **Figures 6G–I** show that transfection of miR-672-3p significantly suppressed the production of pro-inflammatory cytokines, including IL-6, RANTES, and IL-1α. Interestingly, transfection of miR-672-3p also significantly reduced the anti-inflammatory cytokine IL-10 (**Figure S6**).

Because *Serpina3a* (rodent ortholog of human SERPINA3) emerged as a key negatively correlated transcript, we probed its upstream signaling axis (**Figure 7A**). STAT3, an established transcription factor that upregulates Serpina3 gene expression, showed higher total, phosphorylated (pSTAT3) protein levels and higher pSTAT3:STAT3 ratio in both males and females following bleomycin-induced ALI (**Figure 7B**). Protein validation of SERPINA3, a key acute-phase protein and miR-672-3p target, further provided evidence for a more robust inflammatory response in males, with males displaying a significant increase in SERPINA3 protein induction post-injury (p = 0.0002), whereas females did not show a significant increase, yielding significantly higher post-injury levels in males (p = 0.008) (**Figure 7C**).

**Figure 7.**
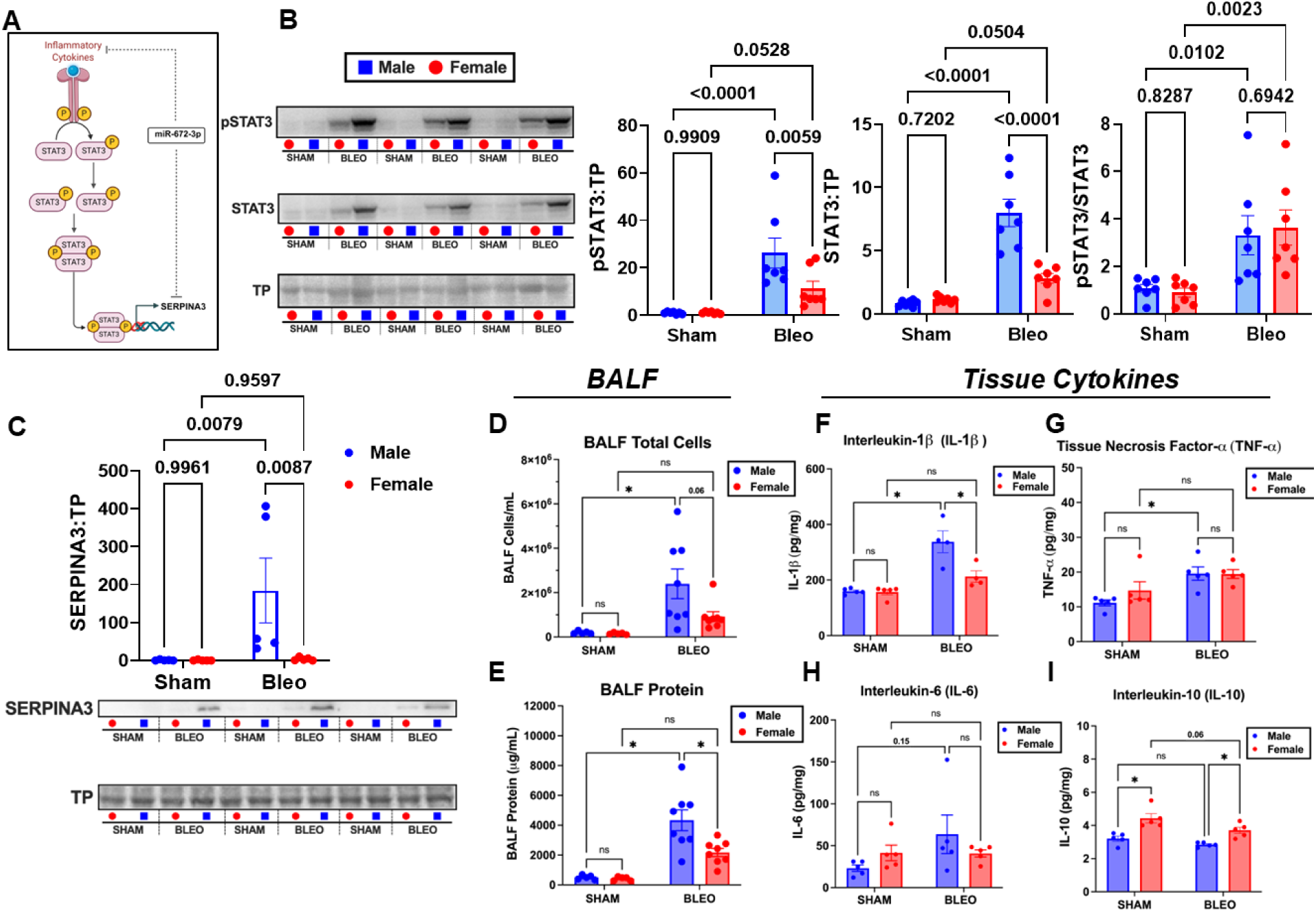
Sex-dependent activation of STAT3 signaling and SERPINA3 expression in bleomycin-induced ALI. (**A**) Inflammatory cytokines bind respective receptors, activating STAT3 signaling. pSTAT3 will translocate to the nucleus and induce *Serpina3*. Dashed lines indicate miR-672-3p is predicted to target multiple effectors of this pathway and dampen overall signaling. (**B**) STAT3 and pSTAT3 sharply increase in males following bleomycin ALI relative to their shams and bleomycin females (all p < 0.0001), whereas females mount a smaller increase in total STAT3 and pSTAT3 but a similar pSTAT3:STAT3 ratio. (**C**) SERPINA3 is significantly elevated in post-bleomycin males (p < 0.0001). (**D**) BALF cells significantly increase in males post-bleomycin (p = 0.01). (**E**) BALF protein is higher in males compared to females post-bleomycin (p = 0.008). (**G-I**) Lung tissue IL-1β levels are higher in males than females (p = 0.005), and IL-10 levels are higher at baseline and post-injury in females (both p < 0.05). Sample sizes: miRNA – RNAseq n = 5 per group RT-qPCR n = 5 per group, immunoblots n = 6, BALF n = 5-8 per group, tissue cytokines n = 4-5 per group. Data are mean ± SEM. *p < 0.05, ns = not significant. Statistical analysis performed using two-way ANOVA with Tukey’s multiple comparisons test.

Vimentin, a marker of mesenchymal transition and tissue remodeling, demonstrated male-specific induction following bleomycin lung injury (p = 0.02), while females showed no significant change compared to sham animals (p = 0.33) (**Figure S7**). Similarly, MFGE8 protein levels were significantly elevated in both males and females following injury, with bleomycin-treated males exhibiting significantly higher MFGE8 levels than females (p = 0.011520) (**Figure S7**).

To examine whether the observed miR-672-3p suppression patterns correspond to inflammatory outcomes, we analyzed cellular recruitment to the alveolar compartment through bronchoalveolar lavage fluid (BALF) analysis. The cellular infiltration patterns showed similar sex-specific trends as the miR-672-3p expression differences (**Figure 7D**). Males, who demonstrated pronounced miR-672-3p suppression, exhibited substantial BALF cell accumulation with an 11.7-fold increase in total cell counts (p = 0.012). Females, who maintained higher residual miR-672-3p levels, did not display a significant increase in BALF cells post-bleomycin (p = 0.63). There was a non-significant difference post-injury in BALF cellularity between males and females, with males presenting higher BALF total cell counts than females (p = 0.068).

BALF protein concentration analysis revealed sex-specific patterns of epithelial barrier dysfunction that paralleled the miR-672-3p and cellular recruitment findings (**Figure 7E**). Males demonstrated substantial barrier compromise following bleomycin injury (p < 0.0001), representing an 8.6-fold increase from baseline. Females exhibited more moderate barrier disruption with a 5.2-fold increase that approached but did not reach statistical significance (p = 0.074). Notably, the comparison between injured males and females revealed significantly higher protein concentrations in males (p = 0.008). No baseline differences were observed between male and female sham animals.

Multiplex cytokine analysis of lung tissue revealed sex-specific inflammatory mediator production that showed similar patterns to the miR-672-3p findings (**Figure 7F-I).** Males demonstrated robust pro-inflammatory cytokine elevation post-bleomycin, including significant IL-1β induction (p < 0.001), representing a 2.1-fold increase compared to sham males (**Figure 7F**). In contrast, females showed minimal IL-1β elevation that did not reach statistical significance (p = 0.252). Direct comparison between bleomycin-treated males and females revealed significantly higher IL-1β levels in males versus females (p = 0.005).

TNF-α levels were significantly elevated in males following injury (p = 0.017), while females exhibited no significant change compared to sham animals (p = 0.26) (**Figure 7G**). Direct comparison between injured males and females showed no significant difference.

Analysis of anti-inflammatory mediators highlighted constitutive sex differences in IL-10 levels. Males maintained significantly lower baseline IL-10 levels compared to females (p = 0.001) (**Figure 7I**). This baseline sex difference persisted post-injury with females maintaining significantly higher IL-10 levels than males (p = 0.023).

Several other cytokines displayed no significant changes in either sex. IL-6 levels remained unchanged in both males (p = 0.172) and females (p > 0.999), with no sex differences observed (p = 0.628) (**Figure 7H**). Similarly, IL-4 levels showed no injury-induced changes in males (p = 0.312) or females (p = 0.912) and no sex differences (p = 0.168) (**Supplementary Figure 3A**). IL-12 levels also remained stable following injury in males (p = 0.265) and females (p = 0.165), with no post-injury sex differences (p = 0.518) (**Supplementary Figure 3B**).

The consistency of these negative correlations across both analytical approaches, despite modest sample sizes, strengthens evidence for functional regulation. The biological coherence of correlated targets, encompassing chemokine signaling, complement activation, and acute-phase responses, suggests coordinated post-transcriptional control. Furthermore, transfection of miR-672-3p significantly suppressed the production of pro-inflammatory cytokines, including IL-6, RANTES, and IL-1α in RAW264.7 cells (**Figure 6G-I**). Combined with the dramatic sex difference in miR-672-3p suppression observed in the qPCR validation (95% in males versus 50% in females), these data support a model where loss of miR-672-3p-mediated repression enables the exaggerated male inflammatory phenotype.

#### Broader miRNA Expression Patterns

A more granular analysis of top differentially expressed miRNAs uncovered multiple miRNAs that were upregulated in both bleomycin-treated females and males relative to their respective sham, notably miR-511-3p_R+1, though with males having higher miR-511-3p_R+1 expression relative to females following bleomycin-induced lung injury (**Figure 5B. Figure S5A-D**). Another miRNA mirroring a similar pattern of upregulation in both sexes post-ALI, but higher in males than females was miR-212-3p (**Figure 5B).** Together, these support the previous evidence of a more perturbed miRNA response in males than females. This coordinated dysregulation suggests that multiple miRNA networks beyond miR-672-3p alone contribute to sex-specific-post-transcriptional programs.

In the body-weight-matched cohort, where we attempted to limit dosing artifacts, a distinct and convergent sex-specific pathophysiology emerged. The functionally more severe injury in males, defined by elevated respiratory rate, reduced lung compliance, and marked hypoxemia, was underpinned by evidence of alveolar-capillary dysfunction, indicated by elevated BALF protein in male versus female animals. This barrier compromise in males corresponded to a robust pro-inflammatory milieu, characterized by increased lung tissue levels of IL-1β and TNF-α and a transcriptomic signature dominated by the complement, coagulation, and cytokine signaling pathways. In stark contrast, females displayed a molecular program that skewed toward cellular proliferation and tissue remodeling, along with constitutively higher levels of IL-10. A potential regulatory node driving these divergent fates was identified in our parallel mRNA-miRNA sequencing and validated with qPCR: the profound suppression of miR-672-3p specifically in males. The validated upregulation of one of its inflammatory protein targets, SERPINA3, establishes a strong molecular association between this post-transcriptional event and the observed hyper-inflammatory state. Therefore, our multi-modal data, spanning from lung mechanics and arterial blood gas down to specific miRNA-target interactions, demonstrate that males and females engage distinct and often opposing molecular programs in response to bleomycin-induced acute lung injury, leading to divergent physiological outcomes.

## Discussion

Clinical outcomes in acute lung injury (ALI) have shown males and females to have similar mortality rates, but females have been reported to have shorter ventilation times and reduced length of hospital stays compared to males (3, 26). Rodent studies using the bleomycin ALI model probing the mechanistic underpinnings of this clinical observation have yielded conflicting conclusions, with mouse studies consistently showing a more severe male phenotype(6–8, 13), and limited rat studies showing a more severe response in females (11). Conventional rodent ALI studies administer a fixed mg/kg intratracheal dose, even though lung mass increases far more slowly than body mass, meaning that heavier males receive substantially more drug per gram of lung than lighter females. Growth-curve data from Sprague Dawley rats illustrate this issue: between 10- and 14-weeks, body weight nearly doubles while lung weight only rises ∼20%, yielding a clear sub-linear allometry (14, 16). Consequently, an age-matched male receives markedly more bleomycin per gram of lung than a female. Toxicology and pharmacology fields routinely correct for such non-linear organ-to-body allometry in sex difference studies (27, 28). In this study, we employed a dual-cohort design to correct for an inherent confound present in preclinical studies, finding a male-biased propensity to a more severe response to bleomycin-induced acute lung injury as measured across functional, molecular, and transcriptional dimensions. The functionally worse outcome in males, defined by severe hypoxemia and impaired lung mechanics, was underpinned by a dysregulated inflammatory program. In contrast, females mounted a more restrained, pro-reparative response, transcriptionally biased toward tissue remodeling. Our unique mRNA-miRNA analysis identified distinct, sex-specific post-transcriptional regulatory nodes, implicating the loss of miR-672-3p in the male inflammatory cascade and enhanced BMPR2 in the female protective phenotype. Our central finding is that after normalizing the effective drug dose via body weight matching, a more severe male-biased injury profile was clarified and confirmed. The greater hypoxemia present in males was observed in both cohorts, while analyses exclusive to the BW-matched cohort revealed greater barrier permeability. This fundamentally divergent molecular response underpinned the functionally worse outcome in males. Whereas males launched a dysregulated inflammatory program, females mounted a more restrained response with a transcriptional signature biased toward cell cycle activation and tissue remodeling. By optimizing our experimental design to limit dosing confounds, we have provided a multi-modal body of evidence that shows the male response to bleomycin exposure results in severe acute lung dysfunction in this model.

Physiological endpoints across both cohorts revealed that males are less resilient to bleomycin-induced ALI, displaying higher respiratory rates, lower oxygenation, and, in the BW-matched cohort, a more pronounced decrease in lung compliance. Male mice exposed to bleomycin displayed lower compliance and higher airway resistance compared to females at seven days (8) and 21 days post-bleomycin (13). Other models of lung injury using hydrochloric acid (HCl) and nitrogen mustard (NM) have also reported females to have less perturbed lung resistance/compliance compared to males (29). Arterial oxygenation (PaO_2_ and sO_2_%), a clinically relevant metric not commonly measured in rodent models (30), was lower in males compared to females post-injury. In an LPS ALI model, no sex differences were reported between young and aged males and females in measuring PaO_2_ and sO_2_% post-LPS (31). Overall, our study confirms that the functional differences are not an artifact of larger male bodies because they persisted under the weight-matched bleomycin-dosed cohort, a finding that has been observed in LPS models where size-normalized LPS inoculation still yielded worse outcomes (17).

BALF cellularity and protein concentration, proxies for alveolar-capillary barrier integrity and overall immune response(32), were significantly elevated in males compared to females post-bleomycin. A mouse model of bleomycin-induced ALI reported BALF cells and protein levels to be higher in males at 7 and 14 days post-bleomycin (7). In line with this are LPS ALI mouse models, which found higher BALF protein in males (10, 33).

In a model of bleomycin fibrosis, miR-672-3p was identified as a potential regulatory node in a miRNA-circRNA network (34). Outside of respiratory research, overexpression of miR-672-3p has been shown to be protective in spinal cord contusion by curtailing ferroptosis through FSP1 repression (35, 36). In cardiovascular studies, miR-672-5p attenuates pressure overload hypertrophy by restraining JUN-dependent cytokine signaling (37). In ovariectomized mice, miR-672-5p showed therapeutic rescue against estrogen deficiency-induced osteopenia and sarcopenia (38). Expression of miR-672-3p is X-linked and constrained to rodents, with no human ortholog identified (39, 40).

A novel aspect of our work is the parallel mRNA-miRNA analysis in the BW-matched cohort, which identified post-transcriptional regulators driving divergent responses in males versus females. A key finding was the identification of miR-672-3p being profoundly suppressed in males after bleomycin, whereas females retained significantly higher levels after bleomycin. Using *in silico* prediction, miR-672-3p expression was inversely related to several high-confidence targets, including multiple inflammatory mediators significantly differentially expressed in the mRNA-seq data *(Cxcr3*, *Cxcl11*, *Serpina3a*, *Cd80*). Loss of *Cxcr3* reduces acute lung inflammation, and its ligand *Cxcl11* is a known marker of pulmonary inflammation; together, *Cxcr3-Cxcl11* signaling is crucial for immune cell recruitment to the injured lung (41–43). *Serpina3a*, a rodent equivalent of human *Serpina3*, was significantly more expressed in males compared to females at the gene level. Additionally, probing of SERPINA3 showed higher levels in males than in females after bleomycin exposure ALI. Although the predicted rodent miR-672-3-*Serpina3a* interaction only carries a moderate TargetScan score, human *Serpina3* scores much higher (23). SERPINA3 functions as an acute-phase protein and an inflammatory protease inhibitor, with its main target being cathepsin G released from neutrophil granules (44, 45). Proteomic studies have identified Serpina3 to be elevated in lung and blood tissue of humans with ARDS (46). Inhibition of Serpina3 *in vitro* reduces LPS-induced cytokine production (47). Linked with the male-biased SERPINA3 induction, total STAT3 and pSTAT3 levels were also higher in male lungs, consistent with the role of STAT3 and pro-inflammatory cytokines coordinating *Serpina3* expression (48).

Another sex-specific miRNA expression pattern in our data was miR-490-5p. We found miR-490-5p to be significantly lower in females than males after bleomycin, and females showed higher gene and protein expression of *Bmpr2* (bone morphogenetic protein receptor II), a predicted target of miR-490-5p. This is supported by studies in other systems demonstrating that miR-490-5p can directly bind to and suppress *Bmpr2* mRNA. (49, 50). BMPR2 is a key receptor in the BMP signaling pathway, which generally counter-regulates TGF-β signaling and also has roles in vascular integrity and tissue homeostasis (51–53). The female-biased increase in BMPR2 could tilt the balance towards BMP pathway activity in females. Indeed, our RNA-seq showed that females had higher expression of several BMP signaling components (*Bmp6, Bmp8a, Smad9, Bmpr2, Smad6*), consistent with an enhanced BMP signaling environment. In the lung, mutations in *Bmpr2* are the most common genetic cause of pulmonary hypertension (PAH) (54, 55). Stable *Bmpr2* is necessary for endothelial integrity and reducing vascular leak; activation of BMPR2 has been shown to reverse PAH (56, 57). Enhanced *Bmpr2* induction from the gene to protein level in females could explain lower BALF protein and thus preserved PaO_2_. In the context of lung injury exosomal BMPR2 delivery has been shown to accelerate tissue repair, and reduce IL-1β and IL-6 levels (58). We observed that females had lower IL-1β lung tissue levels compared to males. Our data highlight a previously unrecognized microRNA-mediated association (miR-490-5p – *Bmpr2*) that differs by sex and could influence ALI outcomes.

Beyond specific miRNAs, the global transcriptomic pathway analysis reinforces that male lungs were in a pro-inflammatory state, whereas female lungs had more pathways involved in tissue repair and resolution. Male upregulated genes included multiple neutrophil and macrophage chemoattractants (*Cxcl10 Ccl17*), pattern recognition receptors, and fibrinolysis factors. *Cxcl10* functions as neutrophil chemoattractants and has been implicated in the rat bleomycin ALI model (59), while *Ccl17* is responsible for macrophage migration to the lungs following bleomycin exposure (60). One particularly interesting finding was the sex difference in the complement and coagulation cascade pathway, where males had higher expression of coagulation factors (*F7*, *F10*) along with *Timp1*, implying more intense tissue remodeling and inflammation. *Timp1*, expressed on the X chromosome, functions to modulate inflammation and matrix turnover in bleomycin acute lung injury and is markedly increased under injury conditions (61–65). Clinically, decreased plasma levels of TIMP1 are correlated with lower oxygenation, fewer ventilator-free days, more extended hospital stays, and higher 30- and 90-day mortality exclusively among female ARDS patients (66, 67). Although we observed higher levels in males compared to females, the elevated levels of *Timp1* contributing to a more severe phenotype in males are backed up by clinical evidence. Females showed greater upregulation of multiple integrin subunits and focal adhesion proteins (*Itga2*, *Itgb3*, *Flrt3*). *Itga2* has been shown to suppress TNF-α while upregulating IL-10, demonstrating broad anti-inflammatory effects in an LPS model of ALI (68). *Itgb3* knockout mice show greater endothelial permeability and pro-inflammatory cytokine induction (69). *Flrt3*, when overexpressed, improves vascular integrity, reduces pulmonary edema, and ultimately improves PaO_2_/FiO_2_ ratios following lung injury (70). The divergent transcriptional programming was reflected in lung tissue cytokines, with IL-10 levels being higher in females under injury and sham conditions, and TNF-α and IL-1β being exclusively elevated in males.

In this study, we employed a dual cohort design to disentangle intrinsic sex-specific biology from dosing artifacts in the rat bleomycin model of acute lung injury. Our central finding is that after normalizing the effective drug dose via body weight matching, a more severe male-biased injury profile was clarified and confirmed. The greater hypoxemia present in males was observed in both cohorts, while analyses exclusive to the body weight match cohort revealed greater barrier permeability. This fundamentally divergent molecular response underpinned this functionally worse outcome in males: whereas males launched a dysregulated inflammatory program, females mounted a more restrained response with a transcriptional signature biased towards cell cycle activation and tissue remodeling. By optimizing our experimental design to limit dosing confounds we have provided a multi-modal body of evidence that shows the male response to bleomycin-induced ALI results in severe acute lung dysfunction in this model.

## Limitations

Our findings derive from the rat bleomycin model, which recapitulates direct epithelial injury, but not the full spectrum of clinical ALI/ARDS etiologies. We did not confirm the circulating sex-steroid hormone levels, and estrous cycle staging was not measured, leaving hormonal drivers unresolved. Further studies probing the role of the estrous cycle using the 4-vinylcyclohexene diepoxide (VCD) model of ovarian failure (71), which leads to a gradual, chemically induced menopause without the acute stress of surgery, would be a potential next step to isolate the contribution of female sex hormones to the protective phenotype conferred in females.

## Conclusions

By employing a dual-cohort strategy, specifically the body-weight-matched cohort, which mitigates dosing artifacts, we demonstrate that male rats exhibit more severe responses to bleomycin-induced ALI than females. Males developed worse hypoxemia, increased alveolar-capillary barrier permeability, and launched a dysregulated inflammatory program characterized by miR-672-3p suppression and excessive inflammatory mediator expression. In contrast, females better maintained miR-672-3p expression, had higher BMPR2 expression, which was correlated with reduced miR-490-5p expression, and overall maintained a controlled tissue repair response. These findings establish that male vulnerability to bleomycin ALI persists in age- and body-weight-matched conditions, highlighting the need for sex-specific considerations in preclinical ALI research and future therapeutic development.

## References

1. MA M, GA Z. Acute lung injury and the acute respiratory distress syndrome: four decades of inquiry into pathogenesis and rational management - PubMed. American journal of respiratory cell and molecular biology. 2005 Oct;33(4).

2. Bellani G, Laffey JG, Pham T, Fan E, Brochard L, Esteban A, et al. Trends in Acute Respiratory Distress Syndrome in 50 Countries. JAMA. 2016/02/23;315(8).

3. McNicholas BA, Madotto F, Pham T, Rezoagli E, Masterson CH, Horie S, et al. Demographics, management and outcome of females and males with acute respiratory distress syndrome in the LUNG SAFE prospective cohort study. European Respiratory Journal. 2019-10-17;54(4).

4. Matute-Bello G, Frevert CW, Martin TR. Animal models of acute lung injury. American Journal of Physiology - Lung Cellular and Molecular Physiology. 2008 Sep;295(3).

5. Matute-Bello G, Downey G, Moore BB, Groshong SD, Matthay MA, Slutsky AS, et al. An Official American Thoracic Society Workshop Report: Features and Measurements of Experimental Acute Lung Injury in Animals. American Journal of Respiratory Cell and Molecular Biology. 2011 May;44(5).

6. Redente EF, Jacobsen KM, Solomon JJ, Lara AR, Faubel S, Keith RC, et al. Age and sex dimorphisms contribute to the severity of bleomycin-induced lung injury and fibrosis. American Journal of Physiology - Lung Cellular and Molecular Physiology. 2011 Jul 8;301(4).

7. Lamichhane R, Patial S, Saini Y. Higher susceptibility of males to bleomycin-induced pulmonary inflammation is associated with sex-specific transcriptomic differences in myeloid cells. Toxicology and applied pharmacology. 2022 Sep 8;454.

8. Prasad C, Duraisamy SK, Sundar IK. Lung mechanics showing sex-based differences and circadian time-of-day response to bleomycin-induced lung injury in mice. Physiological Reports. 2023 Oct 5;11(19).

9. K L, C S, W J, L W, XI C, B M. Analysis of the transcriptome in hyperoxic lung injury and sex-specific alterations in gene expression - PubMed. PloS one. 07/08/2014;9(7).

10. Kuhar E, Chander N, Stewart DJ, Jahandideh F, Zhang H, Kristof AS, et al. A preclinical systematic review and meta-analysis assessing the effect of biological sex in lipopolysaccharide-induced acute lung injury. American Journal of Physiology-Lung Cellular and Molecular Physiology. 2024 May 16;326(6).

11. Gharaee-Kermani M, Hatano K, Nozaki Y, Phan SH. Gender-Based Differences in Bleomycin-Induced Pulmonary Fibrosis. The American Journal of Pathology. 2005 Jun;166(6).

12. J H, S S, G L. Mechanisms of bleomycin-induced lung damage - PubMed. Archives of toxicology. 1991;65(2).

13. JW V, JW C, MA C, LM D, CD F, GP F, et al. Male sex hormones exacerbate lung function impairment after bleomycin-induced pulmonary fibrosis - PubMed. American journal of respiratory cell and molecular biology. 2008 Jul;39(1).

14. Brower M, Grace M, Kotz CM, Koya V. Comparative analysis of growth characteristics of Sprague Dawley rats obtained from different sources. Laboratory Animal Research. 2015 Dec 22;31(4).

15. Toxicity studies of abrasive blasting agents administered by inhalation to F344/NTac rats and Sprague Dawley (Hsd:Sprague Dawley SD) Rats - PubMed. Toxicity report series. 2020 Jul(91).

16. Piao Y, Liu Y, Xie X. Change Trends of Organ Weight Background Data in Sprague Dawley Rats at Different Ages. Journal of Toxicologic Pathology. 2013 Apr 22;26(1).

17. Mock JR, Tune MK, Bose PG, McCullough MJ, Doerschuk CM. Comparison of different methods of initiating lung inflammation and the sex-specific effects on inflammatory parameters. American Journal of Physiology - Lung Cellular and Molecular Physiology. 2023 Jan 3;324(2).

18. Kilkenny C, Browne WJ, Cuthill IC, Emerson M, Altman DG. Improving Bioscience Research Reporting: The ARRIVE Guidelines for Reporting Animal Research. PLoS Biology. 2010 Jun 29;8(6).

19. the NRCUCftUotGf, Animals CaUoL. Guide for the Care and Use of Laboratory Animals. 2011.

20. Bankhead P, Loughrey MB, Fernández JA, Dombrowski Y, McArt DG, Dunne PD, et al. QuPath: Open source software for digital pathology image analysis. Scientific Reports 2017 7:1. 2017-12-04;7(1).

21. MI L, W H, S A. Moderated estimation of fold change and dispersion for RNA-seq data with DESeq2 - PubMed. Genome biology. 2014;15(12).

22. MD R, DJ M, GK S. edgeR: a Bioconductor package for differential expression analysis of digital gene expression data - PubMed. Bioinformatics (Oxford, England). 01/01/2010;26(1).

23. Agarwal V, Bell GW, Nam J-W, Bartel DP. Predicting effective microRNA target sites in mammalian mRNAs. eLife. 2015 Aug 12;4.

24. John B, Enright AJ, Aravin A, Tuschl T, Sander C, Marks DS. Human MicroRNA Targets. PLoS Biology. 2004 Oct 5;2(11).

25. PK B. A tool for design of primers for microRNA-specific quantitative RT-qPCR - PubMed. BMC bioinformatics. 01/28/2014;15(1).

26. Zhao H, Yang B, Dai H, Li C, Ruan H, Li Y. SEX DIFFERENCES IN SEPSIS-RELATED ACUTE RESPIRATORY DISTRESS SYNDROME AND OTHER SHORT-TERM OUTCOMES AMONG CRITICALLY ILL PATIENTS WITH SEPSIS: A RETROSPECTIVE STUDY IN CHINA. Shock. May 2025;63(5).

27. Wilson LAB, Zajitschek SRK, Lagisz M, Mason J, Haselimashhadi H, Nakagawa S. Sex differences in allometry for phenotypic traits in mice indicate that females are not scaled males. Nature Communications. 2022 Dec 12;13(1).

28. Sharma V, McNeill JH. To scale or not to scale: the principles of dose extrapolation. British Journal of Pharmacology. 2009 Jun 5;157(6).

29. Solopov P, Biancatelli RMLC, Dimitropoulou C, Catravas JD. Sex-Related Differences in Murine Models of Chemically Induced Pulmonary Fibrosis. International Journal of Molecular Sciences. 2021 May 31;22(11).

30. E F, V B, DJ K, O G. SpO2/FiO2 ratio on hospital admission is an indicator of early acute respiratory distress syndrome development among patients at risk - PubMed. Journal of intensive care medicine. 2015 May;30(4).

31. Crispens C, Fleckenstein E, Wilken-Schmitz A, Weber S, Gröger M, Hoffmann A, et al. Sex- and age-related differences in LPS-induced lung injury: establishing a mouse intensive care unit. Intensive Care Medicine Experimental 2025 13:1. 2025-05-06;13(1).

32. ME D, D Y, YP D. Using Bronchoalveolar Lavage to Evaluate Changes in Pulmonary Diseases - PubMed. Methods in molecular biology (Clifton, NJ). 2020;2102.

33. Card JW, Carey MA, Bradbury JA, DeGraff LM, Morgan DL, Moorman MP, et al. Gender Differences in Murine Airway Responsiveness and Lipopolysaccharide-Induced Inflammation. Journal of immunology (Baltimore, Md : 1950). 2006 Jul 1;177(1).

34. Liu X, Liu H, Jia X, He R, Zhang X, Zhang W. Frontiers | Changing Expression Profiles of Messenger RNA, MicroRNA, Long Non-coding RNA, and Circular RNA Reveal the Key Regulators and Interaction Networks of Competing Endogenous RNA in Pulmonary Fibrosis. Frontiers in Genetics. 2020/09/24;11.

35. Wang F, Li J, Zhao Y, Guo D, Liu D, Chang Se, et al. miR-672-3p Promotes Functional Recovery in Rats with Contusive Spinal Cord Injury by Inhibiting Ferroptosis Suppressor Protein 1. Oxidative Medicine and Cellular Longevity. 2022 Feb 21;2022(1).

36. Sun C, Liu D, Gao S, Xiu M, Zhang Z, Sun C, et al. Propofol Ameliorates Spinal Cord Injury Process by Mediating miR-672-3p/TNIP2 Axis. Biochemical Genetics 2024 62:6. 2024-02-20;62(6).

37. Y L, F W. A new miRNA regulator, miR-672, reduces cardiac hypertrophy by inhibiting JUN expression - PubMed. Gene. 03/30/2018;648.

38. Ahmad N, Kushwaha P, Karvande A, Tripathi AK, Kothari P, Adhikary S, et al. MicroRNA-672-5p Identified during Weaning Reverses Osteopenia and Sarcopenia in Ovariectomized Mice. Molecular Therapy Nucleic Acids. 2019 Jan 10;14.

39. National Center for Biotechnology Information. Mir672 microRNA 672 [Mus musculus (house mouse)] 2024 [Available from: https://www.ncbi.nlm.nih.gov/gene/751535.

40. Smith A, Calley J, Mathur S, Qian H-R, Wu H, Farmen M, et al. The Rat microRNA body atlas; Evaluation of the microRNA content of rat organs through deep sequencing and characterization of pancreas enriched miRNAs as biomarkers of pancreatic toxicity in the rat and dog. BMC Genomics 2016 17:1. 2016-08-30;17(1).

41. Nie L, Xiang R-l, Liu Y, Zhou W-x, Jiang L, Lu B, et al. Acute pulmonary inflammation is inhibited in CXCR3 knockout mice after short-term cigarette smoke exposure. Acta Pharmacologica Sinica 2008 29:12. 2008/12;29(12).

42. Kameda M, Otsuka M, Chiba H, Kuronuma K, Hasegawa T, Takahashi H, et al. CXCL9, CXCL10, and CXCL11; biomarkers of pulmonary inflammation associated with autoimmunity in patients with collagen vascular diseases–associated interstitial lung disease and interstitial pneumonia with autoimmune features. PLoS ONE. 2020 Nov 2;15(11).

43. Torraca V, Cui C, Boland R, Bebelman J-P, van der Sar AM, Smit MJ, et al. The CXCR3-CXCL11 signaling axis mediates macrophage recruitment and dissemination of mycobacterial infection. Disease Models & Mechanisms. 2015/03/01;8(3).

44. Jin Y, Wang W, Wang Q, Zhang Y, Zahid KR, Raza U, et al. Alpha-1-antichymotrypsin as a novel biomarker for diagnosis, prognosis, and therapy prediction in human diseases. Cancer Cell International. 2022 Apr 19;22(1).

45. Sánchez-Navarro A, González-Soria I, Caldiño-Bohn R, Bobadilla NA. An integrative view of serpins in health and disease: the contribution of SerpinA3. American Journal of Physiology-Cell Physiology. 2021 Jan 27.

46. R G, H L, G L, J X, C H, X Z, et al. Integrative proteomic profiling of lung tissues and blood in acute respiratory distress syndrome - PubMed. Frontiers in immunology. 05/01/2023;14.

47. X W, B C, C C. Identification of biomarkers and candidate small-molecule drugs in lipopolysaccharide (LPS)-induced acute lung injury by bioinformatics analysis - PubMed. Allergologia et immunopathologia. 01/01/2023;51(1).

48. Li Y, Guo L. Frontiers | The versatile role of Serpina3c in physiological and pathological processes: a review of recent studies. Frontiers in Endocrinology. 2023/05/23;14.

49. Yang Q, Guo J, Ren Z, Li B, Huang H, Yang Z. LncRNA NONHSAT030515 promotes the chondrogenic differentiation of human adipose-derived stem cells via regulating the miR-490-5p/BMPR2 axis. Journal of Orthopaedic Surgery and Research. 2021 Nov 6;16(1).

50. Yang Z, Hao J, Hu ZM. MicroRNA expression profiles in human adipose-derived stem cells during chondrogenic differentiation. International Journal of Molecular Medicine. 2015-03-01;35(3).

51. Wu M, Wu S, Chen W, Li Y-P, Wu M, Wu S, et al. The roles and regulatory mechanisms of TGF-β and BMP signaling in bone and cartilage development, homeostasis and disease. Cell Research 2024 34:2. 2024-01-24;34(2).

52. Hiepen C, Jatzlau J, Hildebrandt S, Kampfrath B, Goktas M, Murgai A, et al. BMPR2 acts as a gatekeeper to protect endothelial cells from increased TGFβ responses and altered cell mechanics. PLOS Biology. Dec 11, 2019;17(12).

53. Teichert-Kuliszewska K, Kutryk MJB, Kuliszewski MA, Karoubi G, Courtman DW, Zucco L, et al. Bone Morphogenetic Protein Receptor-2 Signaling Promotes Pulmonary Arterial Endothelial Cell Survival. Circulation Research. 2006-02-03;98(2).

54. Li W, Quigley K. Bone morphogenetic protein signalling in pulmonary arterial hypertension: revisiting the BMPRII connection. Biochemical Society Transactions. 2024 May 8;52(3).

55. Evans JD, Girerd B, Montani D, Wang X-J, Galiè N, Austin ED, et al. BMPR2 mutations and survival in pulmonary arterial hypertension: an individual participant data meta-analysis. The Lancet Respiratory Medicine. 2016;4(2).

56. Long L, Ormiston ML, Yang X, Southwood M, Gräf S, Machado RD, et al. Selective enhancement of endothelial BMPR-II with BMP9 reverses pulmonary arterial hypertension. Nature Medicine. 2015;21(7).

57. Spiekerkoetter E, Tian X, Cai J, Hopper RK, Sudheendra D, Li CG, et al. FK506 activates BMPR2, rescues endothelial dysfunction, and reverses pulmonary hypertension. Journal of Clinical Investigation. 2013;123(8).

58. Exosomal BMPR2 Macromolecule Facilitates Alveolar Epithelial Cell Repair Through Functional Complex Formation with BMPR1B in Acute Lung Injury. International Journal of Nanomedicine. 2025;Volume 20.

59. Hirani DV, Thielen F, Mansouri S, Danopoulos S, Vohlen C, Haznedar-Karakaya P, et al. CXCL10 deficiency limits macrophage infiltration, preserves lung matrix, and enables lung growth in bronchopulmonary dysplasia. Inflammation and Regeneration 2023 43:1. 2023-10-24;43(1).

60. Ishida Y, Kuninaka Y, Mukaida N, Kondo T. Immune Mechanisms of Pulmonary Fibrosis with Bleomycin. International Journal of Molecular Sciences. 2023 Feb 5;24(4).

61. B M, S C-M, I G, V L, E B. TIMP-1 is a key factor of fibrogenic response to bleomycin in mouse lung - PubMed. International journal of immunopathology and pharmacology. 2006 Jul-Sep;19(3).

62. Kim K-H, Burkhart K, Chen P, Frevert CW, Randolph-Habecker J, Hackman RC, et al. Tissue Inhibitor of Metalloproteinase-1 Deficiency Amplifies Acute Lung Injury in Bleomycin-Exposed Mice. American Journal of Respiratory Cell and Molecular Biology. 2005 Jun 9;33(3).

63. Brew K, Nagase H. The tissue inhibitors of metalloproteinases (TIMPs): An ancient family with structural and functional diversity. Biochimica et biophysica acta. 2010 Jan 15;1803(1).

64. CL F, F G, G H, BR P, LA O. Epithelial expression of TIMP-1 does not alter sensitivity to bleomycin-induced lung injury in C57BL/6 mice - PubMed. American journal of physiology Lung cellular and molecular physiology. 2008 Mar;294(3).

65. Sen’kova AV, Savin IA, Brenner EV, Zenkova MA, Markov AV. Core genes involved in the regulation of acute lung injury and their association with COVID-19 and tumor progression: A bioinformatics and experimental study. PLOS ONE. Nov 22, 2021;16(11).

66. Almuntashiri S, Dutta S, Zhu Y, Gamare S, Ramírez G, Irineo-Moreno V, et al. Estrogen-dependent gene regulation: Molecular basis of TIMP-1 as a sex-specific biomarker for acute lung injury. Physiological Reports. 2024 Sep 12;12(17).

67. Almuntashiri S, Jones TW, Wang X, Sikora A, Zhang D, Almuntashiri S, et al. Plasma TIMP-1 as a sex-specific biomarker for acute lung injury. Biology of Sex Differences 2022 13:1. 2022-12-08;13(1).

68. Kang H, Kim O-H, Chang ES, Kim J, Kim J-Y, Shin G-S, et al. Enhanced engraftment and immunomodulatory effects of integrin alpha-2-overexpressing mesenchymal stromal cells in lipopolysaccharide-induced acute lung injury. Stem Cell Research & Therapy 2025 16:1. 2025-06-03;16(1).

69. Tong Y, Bao C, Xu Y-Q, Tao L, Zhou Y, Zhuang L, et al. The β3/5 Integrin-MMP9 Axis Regulates Pulmonary Inflammatory Response and Endothelial Leakage in Acute Lung Injury. Journal of Inflammation Research. 2021 Oct 5;14.

70. Cao Y, Sheng S, Zhong Y, Shang J, Jin C, Tan Q, et al. FLRT3 Overexpression Attenuates Ischemia–Reperfusion Induced Vascular Hyperpermeability and Lung Injury Through RND3. Lung. 2025 Mar 6;203(1).

71. HL B, DP P, PB H. The VCD Mouse Model of Menopause and Perimenopause for the Study of Sex Differences in Cardiovascular Disease and the Metabolic Syndrome - PubMed. Physiology (Bethesda, Md). 2016 Jul;31(4).

