## Supplemental Figure for "Sex-Specific Pathophysiological Signatures in Allometric Dosing-Controlled Bleomycin Acute Lung Injury Model"

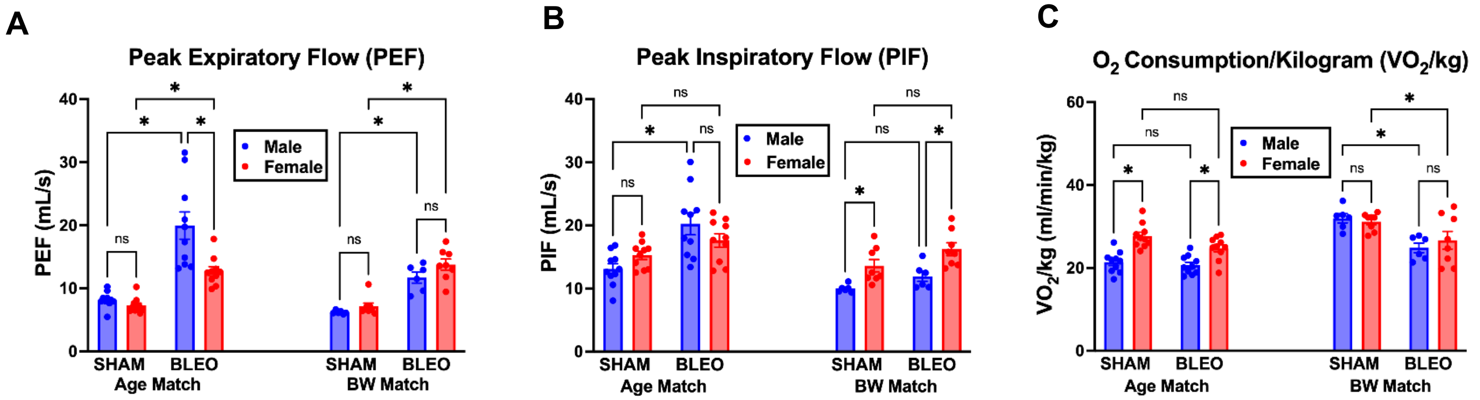


**Supplementary Figure 1**: Peak Expiratory Flow (PEF), Peak Inspiratory Flow (PIF), and weight normalized Oxygen Consumption ($VO_2$/kg). Data are presented as mean ± SEM. For B-G, age matched cohort: n = 10 per group; BW matched group: n = 6 males, n = 8 females. *$p < 0.05$, ns = not significant. Statistical analysis performed using two-way ANOVA with Tukey's multiple comparisons test.

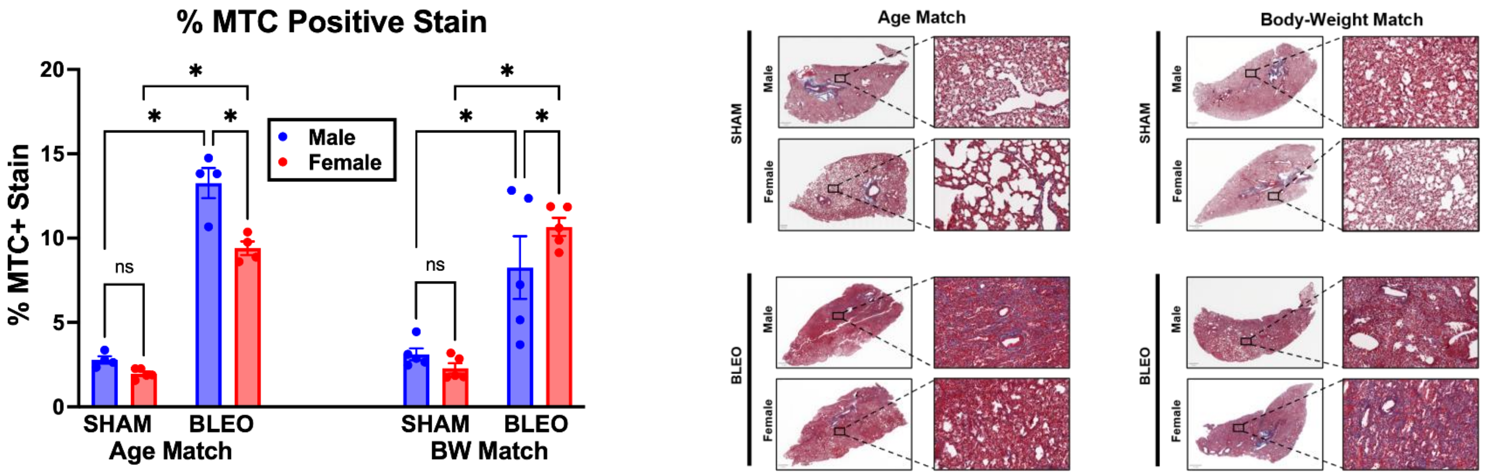


**Supplementary Figure 2:** Masson’s trichrome (MTC) staining quantification revealing sex differences in collagen deposition with males showing greater collagen staining compared to females in the age-match cohort following bleomycin-induced ALI (13.3% vs 9.4%, $p = 0.0046$). In the BW-matched cohort females displayed greater collagen staining post-injury compared to males (10.7% vs 8.3%, $p = 0.04$). Data are plotted as Mean ± SEM. Age-matched group: n = 4-5. BW matched group: n = 5. *$p < 0.05$, ns = not significant. Statistical analysis performed using two-way ANOVA with Tukey’s multiple comparisons test.

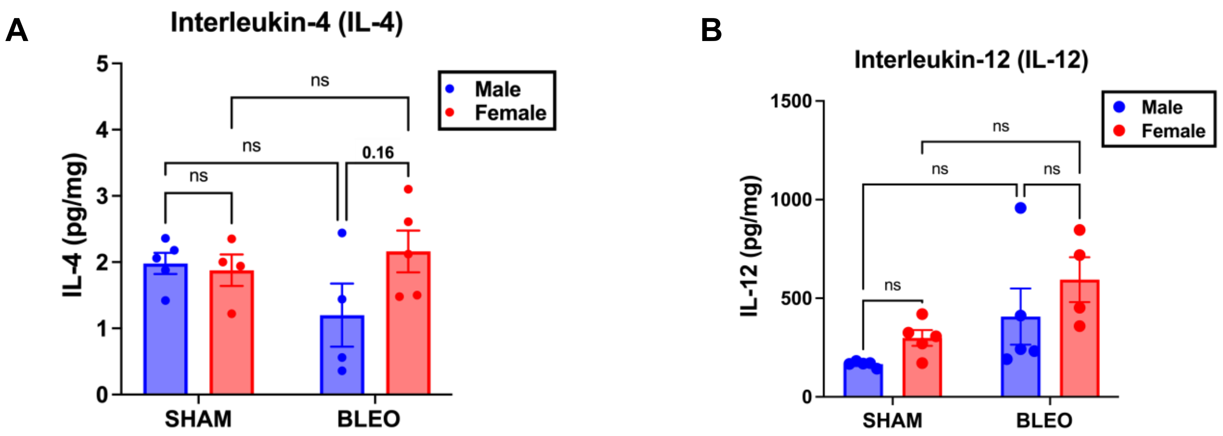


**Supplementary Figure 3: (A)** Interleukin 4 (IL-4) and (B) Interleukin-12 (IL-12) levels assessed at day 7 post-bleomycin or sham in weight matched males and females. Neither cytokine changed significantly post-bleomycin, and no sex differences were observed (IL-4 $p = 0.168$, IL-12 $p = 0.518$). Data are mean ± SEM, n = 4,5 per group. Two-way ANOVA with Tukey's multiple comparisons test; ns = not significant.

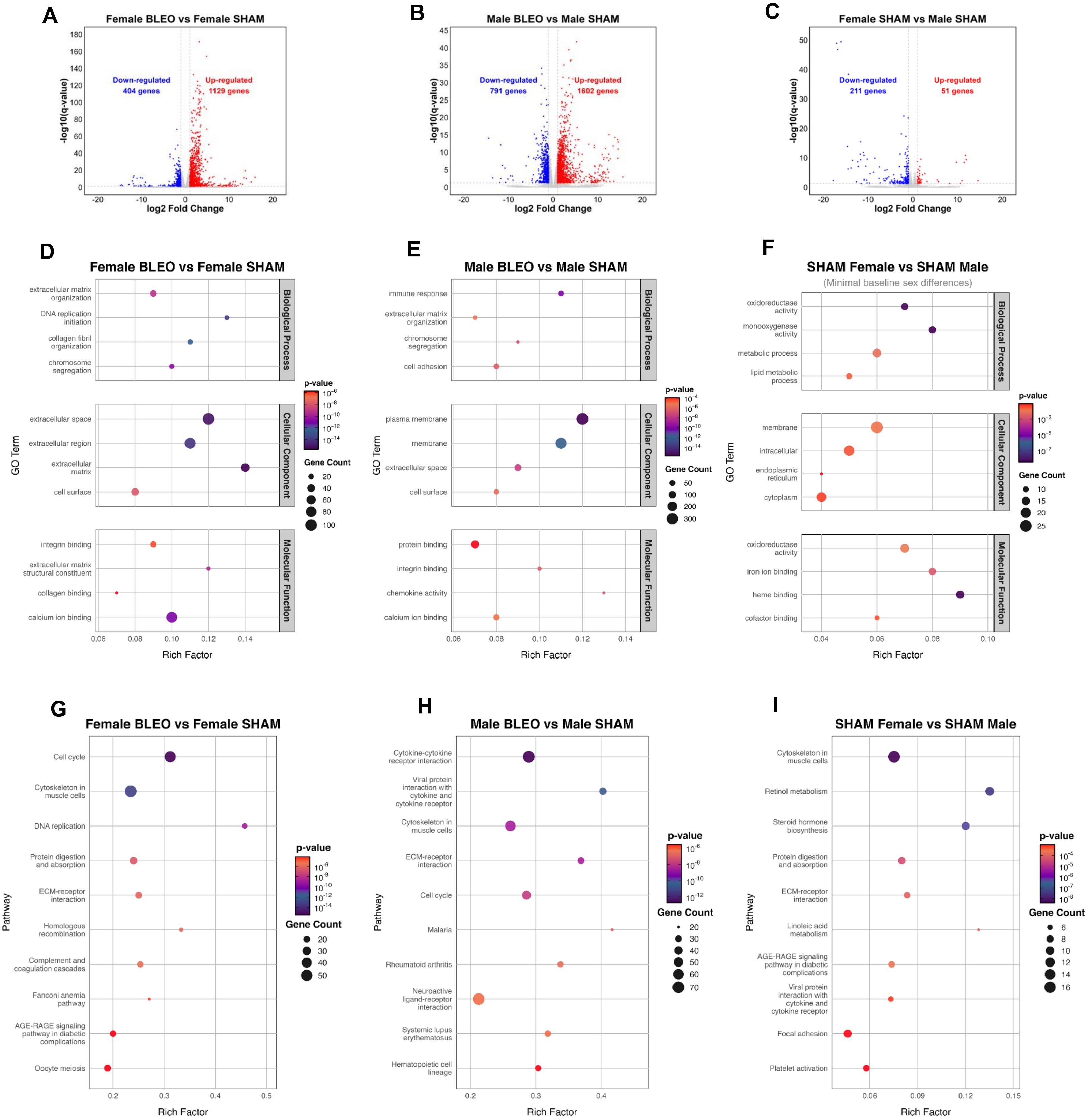
A
Female BLEO vs Female SHAM
Down-regulated
404 genes
Up-regulated
1129 genes
-log10(q-value)
log2 Fold Change
B
Male BLEO vs Male SHAM
Down-regulated
791 genes
Up-regulated
1602 genes
-log10(q-value)
log2 Fold Change
C
Female SHAM vs Male SHAM
Down-regulated
211 genes
Up-regulated
51 genes
-log10(q-value)
log2 Fold Change
D
Female BLEO vs Female SHAM
extracellular matrix organization
DNA replication initiation
collagen fibril organization
chromosome segregation
extracellular space
extracellular region
extracellular matrix
cell surface
integrin binding
extracellular matrix structural constituent
collagen binding
calcium ion binding
Biological Process
Cellular Component
Molecular Function
GO Term
Rich Factor
p-value
Gene Count
E
Male BLEO vs Male SHAM
immune response
extracellular matrix organization
chromosome segregation
cell adhesion
plasma membrane
membrane
extracellular space
cell surface
protein binding
integrin binding
chemokine activity
calcium ion binding
Biological Process
Cellular Component
Molecular Function
GO Term
Rich Factor
p-value
Gene Count
F
SHAM Female vs SHAM Male
(Minimal baseline sex differences)
oxidoreductase activity
monooxygenase activity
metabolic process
lipid metabolic process
membrane
intracellular
endoplasmic reticulum
cytoplasm
oxidoreductase activity
iron ion binding
heme binding
cofactor binding
Biological Process
Cellular Component
Molecular Function
GO Term
Rich Factor
p-value
Gene Count
G
Female BLEO vs Female SHAM
Cell cycle
Cytoskeleton in muscle cells
DNA replication
Protein digestion and absorption
ECM-receptor interaction
Homologous recombination
Complement and coagulation cascades
Fanconi anemia pathway
AGE-RAGE signaling pathway in diabetic complications
Oocyte meiosis
Pathway
Rich Factor
p-value
Gene Count
H
Male BLEO vs Male SHAM
Cytokine-cytokine receptor interaction
Viral protein interaction with cytokine and cytokine receptor
Cytoskeleton in muscle cells
ECM-receptor interaction
Cell cycle
Malaria
Rheumatoid arthritis
Neuroactive ligand-receptor interaction
Systemic lupus erythematosus
Hematopoietic cell lineage
Pathway
Rich Factor
p-value
Gene Count
I
SHAM Female vs SHAM Male
Cytoskeleton in muscle cells
Retinol metabolism
Steroid hormone biosynthesis
Protein digestion and absorption
ECM-receptor interaction
Linoleic acid metabolism
AGE-RAGE signaling pathway in diabetic complications
Viral protein interaction with cytokine and cytokine receptor
Focal adhesion
Platelet activation
Pathway
Rich Factor
p-value
Gene Count

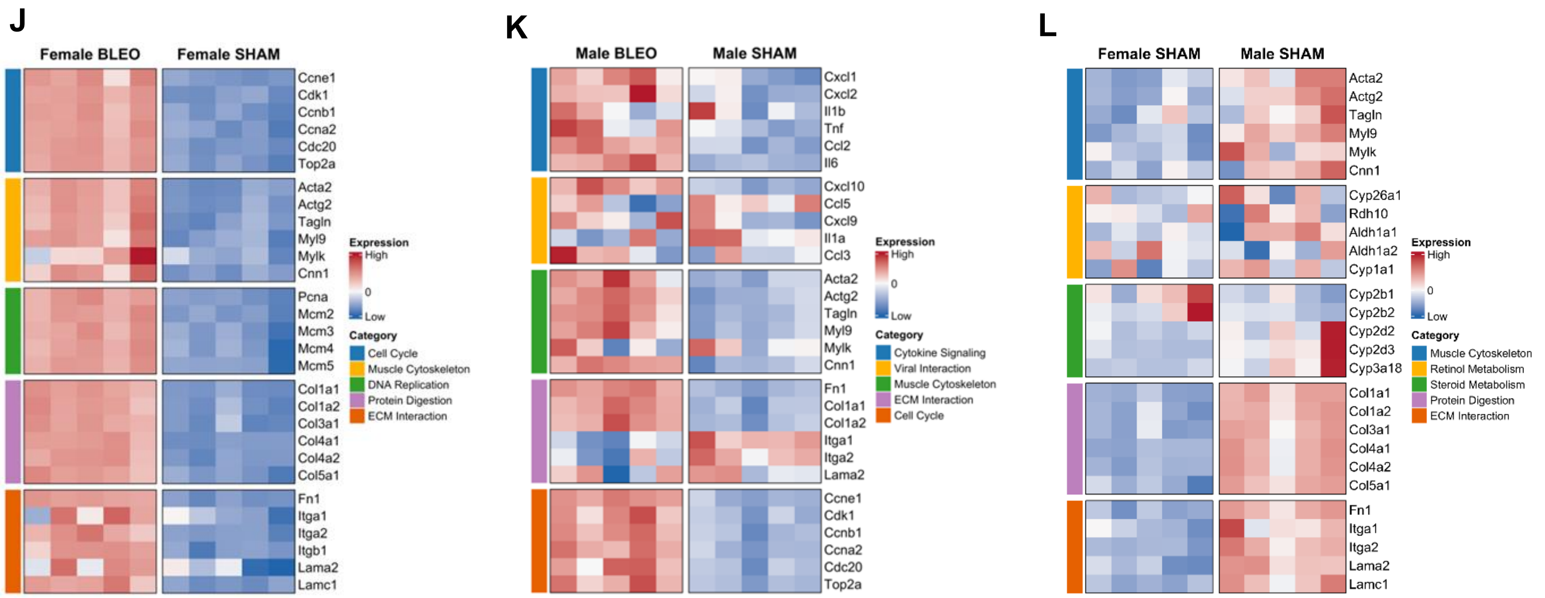


**Supplementary Figure 4:** (**A-C**) Volcano plots of differentially expressed genes (DEGs) for: (**A**) Female BLEO vs Female SHAM, (**B**), Male BLEO vs Male SHAM, and (**C**) Female SHAM vs Male SHAM. (**D-F**) GO enrichment analysis for DEGs in each comparison ranked by rich factor and p-value. (**G-I**) KEGG pathway enrichment for DEGs in each comparison. (**J-L**) Heatmaps of selected DEGs grouped by functional themes. DEGs determine by |$\log_2$FC| ≥, FDR < 0.05. n = 5 biological replicates per sex/treatment.

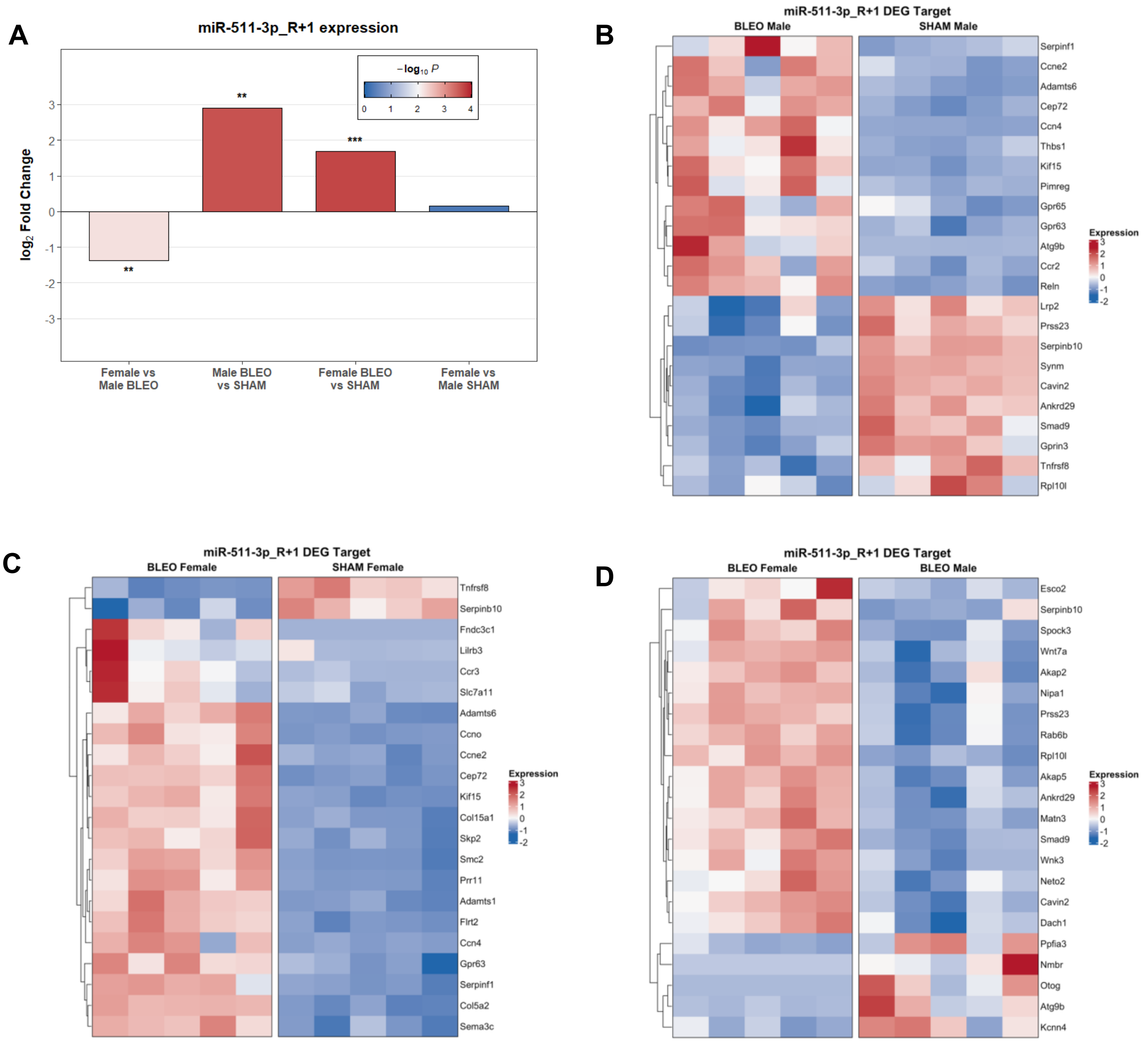


**Supplementary Figure 5:** Lung miRNA-Sequencing. (**A**) Relative expression of miR-511-3p_R+1 and (**B-D**) its predicted mRNA targets that were identified as differentially expressed genes (DEGs) in the parallel lung RNA-sequencing dataset.

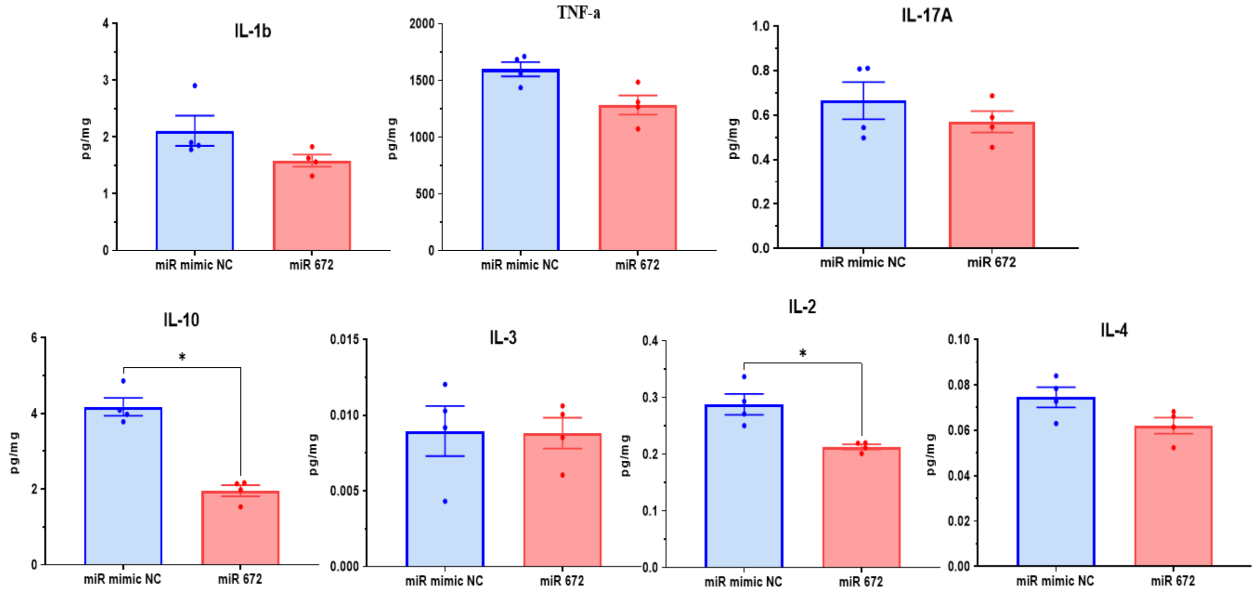


**Supplementary Figure 6:** Effects of miR-672-3p transfection on the production of multiple pro- and anti-inflammatory cytokines in LPS-treated RAW264.7 cells. Cytokines measured include IL-6, IL-1β, TNF-α, IL-17A, IL-10, IL-3, IL-2, and IL-4. Data are presented as mean ± SEM, $n$ = 4 per group. *$p$<0.05. Statistical analysis was performed using the Mann–Whitney test.

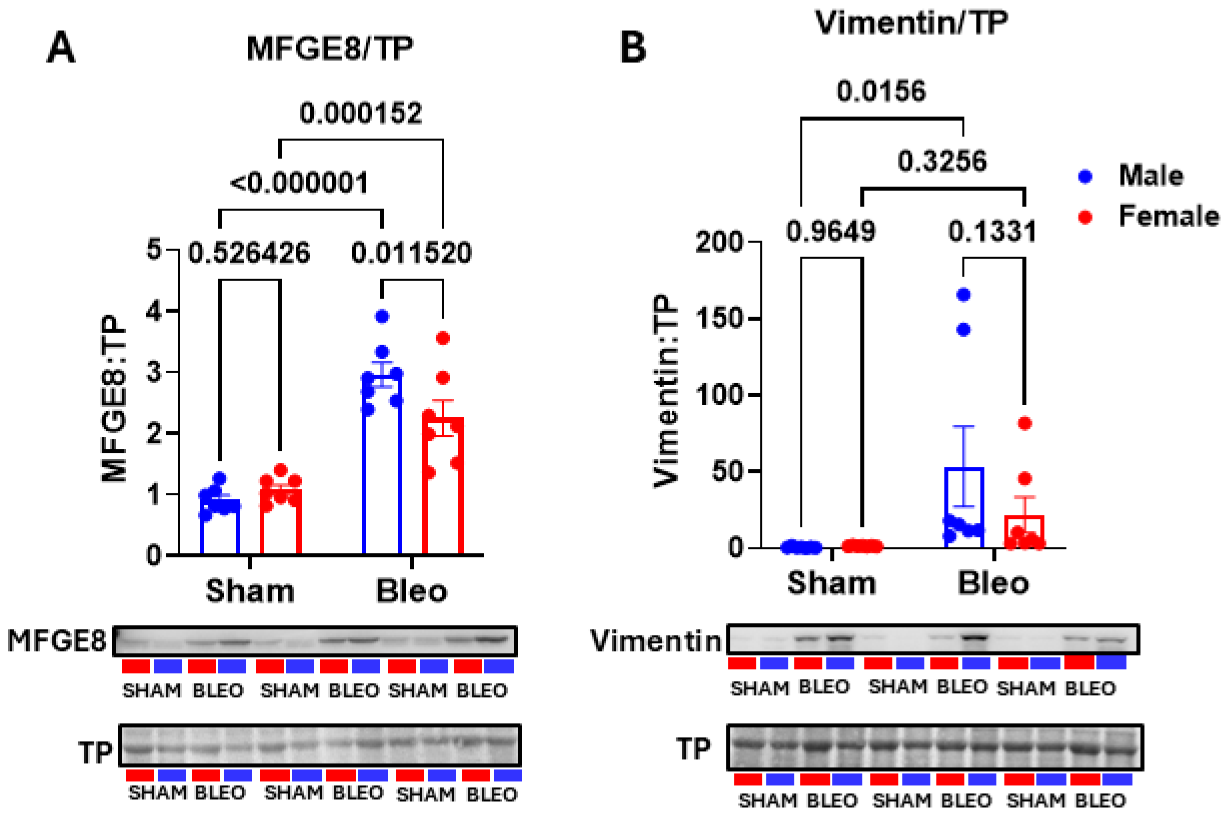


**Supplementary Figure 7:** (**A**) Milk Fat Globule EGF Factor 8 (MFGE8) protein levels were significantly increased in both male and female lungs post-bleomycin, with males having higher MFEG8 protein than females. (**B**) Vimentin was significantly increased in male lungs post-bleomycin but not females. Females had lower vimentin levels than males post-bleomycin, but this did not reach significance. Data are mean ± SEM, n = 6 per group. Two-way ANOVA with Tukey's multiple comparisons test.

**Table 1. Female BLEO vs Female SHAM mRNA-seq**
Significantly Differentially Expressed Genes ($Log_2FC > |1|$; qvalue < 0.05)

| Gene | fc | $log_2(fc)$ | pval | qval | regulation |
|---|---|---|---|---|---|
| Fn1 | 9.095423 | 3.1851408 | 1.4E-176 | 3.1E-172 | up |
| Cemip | 29.5196 | 4.8836014 | 4.4E-159 | 4.9E-155 | up |
| Loxl1 | 3.600238 | 1.8480922 | 2E-137 | 1.5E-133 | up |
| Racgap1 | 6.918776 | 2.7905169 | 2.7E-129 | 1.5E-125 | up |
| Cdc20 | 7.185219 | 2.845032 | 2.9E-124 | 1.3E-120 | up |
| Cdkn3 | 10.06326 | 3.3310254 | 8.6E-124 | 3.2E-120 | up |
| Hmmr | 6.485458 | 2.6972084 | 1E-120 | 3.2E-117 | up |
| Spag5 | 8.691038 | 3.1195286 | 7.5E-119 | 2.1E-115 | up |
| Stab1 | 5.160394 | 2.3674812 | 2.5E-117 | 6.3E-114 | up |
| Diaph3 | 10.43182 | 3.3829191 | 8E-117 | 1.8E-113 | up |
| Knstrn | 10.10082 | 3.3364008 | 4.5E-116 | 9.2E-113 | up |
| Eln | 11.45819 | 3.5183074 | 5.1E-116 | 9.5E-113 | up |
| Ube2c | 9.30507 | 3.218017 | 5.1E-114 | 8.4E-111 | up |
| Tk1 | 17.99321 | 4.1693805 | 5.3E-114 | 8.4E-111 | up |
| Nid1 | 3.542109 | 1.8246088 | 6.3E-109 | 9.4E-106 | up |
| Iqgap3 | 8.520276 | 3.0909002 | 4.7E-108 | 6.6E-105 | up |
| Ccnb2 | 12.60818 | 3.6562886 | 5.8E-108 | 7.7E-105 | up |
| Mastl | 9.068146 | 3.1808075 | 9.1E-107 | 1.1E-103 | up |
| Ckap2 | 10.55353 | 3.3996541 | 6.9E-105 | 8.2E-102 | up |
| Cdk1 | 9.187008 | 3.199595 | 1.5E-103 | 1.7E-100 | up |
| Plk1 | 7.887798 | 2.9796226 | 7.7E-102 | 8.2E-99 | up |
| Tnc | 19.95964 | 4.3190139 | 1.55E-98 | 1.58E-95 | up |
| Col1a1 | 10.77887 | 3.4301336 | 5.42E-98 | 5.28E-95 | up |
| Dtl | 7.729021 | 2.9502857 | 2.62E-97 | 2.44E-94 | up |
| Ltbp2 | 7.475957 | 2.9022583 | 7.55E-97 | 6.76E-94 | up |
| Anln | 11.9805 | 3.5826156 | 1.07E-96 | 9.25E-94 | up |
| Serpine1 | 28.54091 | 4.8349595 | 7.01E-96 | 5.82E-93 | up |
| Cdca3 | 10.16081 | 3.3449433 | 2.88E-95 | 2.3E-92 | up |
| Fkbp10 | 3.581374 | 1.8405131 | 2.19E-94 | 1.7E-91 | up |
| Bgn | 4.64262 | 2.2149392 | 1.06E-93 | 7.95E-91 | up |
| Tpx2 | 10.1295 | 3.340491 | 4.12E-93 | 2.98E-90 | up |
| Fstl1 | 2.665886 | 1.4146153 | 4.11E-92 | 2.88E-89 | up |
| Pttg1 | 9.744623 | 3.2846063 | 6.74E-92 | 4.58E-89 | up |
| Bub1 | 7.336362 | 2.8750648 | 1.92E-91 | 1.27E-88 | up |
| Efemp2 | 2.908399 | 1.540225 | 1.57E-90 | 1.01E-87 | up |
| Nek2l1 (ENSRNOG00000004487) | 6.336209 | 2.6636198 | 2.17E-90 | 1.35E-87 | up |
| Loxl2 | 6.048154 | 2.5964948 | 3.75E-89 | 2.27E-86 | up |
| Ndc80 | 9.375924 | 3.2289609 | 7.02E-89 | 4.14E-86 | up |

| | | | | | |
|---|---|---|---|---|---|
| Top2a | 8.678702 | 3.1174792 | 1.26E-88 | 7.22E-86 | up |
| Nuf2 | 10.27139 | 3.3605591 | 5.15E-88 | 2.88E-85 | up |
| Mcm6 | 4.093313 | 2.0332689 | 3.96E-85 | 2.16E-82 | up |
| Myh10 | 2.884511 | 1.5283268 | 4.73E-85 | 2.52E-82 | up |
| Fbn1 | 5.424774 | 2.4395632 | 1.85E-82 | 9.66E-80 | up |
| Tgfb1i1 | 2.703129 | 1.4346305 | 2.3E-81 | 1.17E-78 | up |
| Ppic | 3.046574 | 1.6071879 | 3.15E-80 | 1.57E-77 | up |
| Ankrd1 | 11.8013 | 3.5608738 | 1.55E-76 | 7.54E-74 | up |
| Mcm2 | 3.523379 | 1.8169596 | 2.01E-76 | 9.59E-74 | up |
| Melk | 10.70296 | 3.4199374 | 9.07E-75 | 4.23E-72 | up |
| Aldh1l2 | 30.63902 | 4.9372982 | 5.33E-72 | 2.44E-69 | up |
| Gsn | 0.265323 | -1.914178 | 1.02E-71 | 4.58E-69 | down |
| Pcna | 2.517839 | 1.3321859 | 3.03E-70 | 1.33E-67 | up |
| Kif11 | 10.17055 | 3.3463262 | 1.57E-68 | 6.77E-66 | up |
| Mki67 | 6.616517 | 2.7260719 | 1.73E-68 | 7.33E-66 | up |
| Cdca5 | 7.855813 | 2.9737606 | 3.47E-68 | 1.44E-65 | up |
| Fanci | 5.380458 | 2.4277291 | 3.92E-68 | 1.6E-65 | up |
| Plpp1 | 3.011513 | 1.5904886 | 4.78E-68 | 1.91E-65 | up |
| Mybl2 | 7.115253 | 2.830915 | 1.69E-67 | 6.64E-65 | up |
| ENSRNOG00000064136 | 3.874535 | 1.9540231 | 3.27E-67 | 1.26E-64 | up |
| Kif20a | 5.195418 | 2.3772398 | 1.22E-66 | 4.64E-64 | up |
| Kif18b | 11.07532 | 3.4692764 | 4.95E-64 | 1.85E-61 | up |
| Ect2 | 9.691779 | 3.2767615 | 2.96E-63 | 1.09E-60 | up |
| Ccnb1 | 7.267505 | 2.8614601 | 3.58E-63 | 1.29E-60 | up |
| Bub1b | 8.566664 | 3.0987335 | 5.37E-63 | 1.91E-60 | up |
| Cit | 9.820171 | 3.2957482 | 9.51E-63 | 3.33E-60 | up |
| Kif20b | 9.087321 | 3.183855 | 2.17E-62 | 7.47E-60 | up |
| Ncaph | 8.650343 | 3.1127573 | 2.88E-62 | 9.79E-60 | up |
| Kntc1 | 8.843348 | 3.1445926 | 4.04E-62 | 1.35E-59 | up |
| Zwilch | 3.191596 | 1.6742779 | 5.51E-62 | 1.81E-59 | up |
| Ncapd2 | 3.911728 | 1.9678059 | 8.2E-62 | 2.66E-59 | up |
| Calu | 2.648449 | 1.4051476 | 1.36E-61 | 4.36E-59 | up |
| Tnfrsf12a | 5.575213 | 2.4790268 | 8.21E-61 | 2.59E-58 | up |
| Cks1b | 3.962104 | 1.9862668 | 1.37E-60 | 4.26E-58 | up |
| Col18a1 | 2.624416 | 1.3919962 | 5.67E-60 | 1.74E-57 | up |
| Stil | 8.485334 | 3.0849714 | 5.42E-59 | 1.64E-56 | up |
| Col4a1 | 3.126422 | 1.6445125 | 3.28E-58 | 9.79E-56 | up |
| Kif2c | 7.760635 | 2.9561746 | 3.4E-58 | 1E-55 | up |
| Olfml2b | 4.989893 | 2.319009 | 4.08E-58 | 1.18E-55 | up |
| Nusap1 | 9.410093 | 3.234209 | 4.12E-58 | 1.18E-55 | up |
| Shcbp1 | 9.797965 | 3.2924821 | 4.71E-58 | 1.33E-55 | up |

| | | | | | |
|---|---|---|---|---|---|
| Col5a2 | 3.190991 | 1.6740047 | 1.03E-56 | 2.88E-54 | up |
| Hmgb2 | 4.659963 | 2.2203186 | 2.52E-56 | 6.96E-54 | up |
| Vcan | 7.927954 | 2.9869486 | 1.82E-55 | 4.98E-53 | up |
| Pole | 6.206892 | 2.6338709 | 4.76E-55 | 1.28E-52 | up |
| Fancd2 | 5.764773 | 2.5272639 | 6.34E-55 | 1.69E-52 | up |
| Cenpe | 10.19575 | 3.3498955 | 2E-54 | 5.26E-52 | up |
| Ckap2l | 10.98154 | 3.4570082 | 6.7E-54 | 1.75E-51 | up |
| Pimreg | 42.41853 | 5.4066226 | 2.73E-53 | 7.04E-51 | up |
| Brca1 | 4.794033 | 2.26124 | 4.21E-53 | 1.07E-50 | up |
| Slc5a1 | 0.311067 | -1.684702 | 1.03E-52 | 2.6E-50 | down |
| Ccna2 | 10.26617 | 3.3598258 | 1.16E-52 | 2.9E-50 | up |
| Col5a1 | 5.360285 | 2.4223097 | 5.42E-52 | 1.33E-49 | up |
| Thbs1 | 5.709682 | 2.5134104 | 1.39E-51 | 3.39E-49 | up |
| Cdca8 | 4.765723 | 2.2526951 | 2.23E-51 | 5.36E-49 | up |
| Rad51 | 7.900873 | 2.982012 | 3.75E-51 | 8.95E-49 | up |
| Rrm2 | 7.407522 | 2.8889911 | 6.84E-50 | 1.61E-47 | up |
| Fblim1 | 4.1066 | 2.0379443 | 1.26E-49 | 2.95E-47 | up |
| Dbf4 | 4.014328 | 2.0051586 | 2.47E-49 | 5.7E-47 | up |
| Pdzrn3 | 3.367447 | 1.7516551 | 3.57E-49 | 8.17E-47 | up |
| Acta2 | 4.210924 | 2.0741369 | 4.85E-49 | 1.1E-46 | up |
| Rmi2 | 8.229453 | 3.0407965 | 8.3E-49 | 1.86E-46 | up |
| Birc5 | 9.133873 | 3.1912268 | 2.26E-48 | 5.02E-46 | up |
| Dlgap5 | 6.01009 | 2.5873866 | 2.53E-48 | 5.55E-46 | up |
| Cyp4b1 | 0.209581 | -2.254418 | 4.71E-48 | 1.02E-45 | down |
| Col6a3 | 2.984708 | 1.5775897 | 2.22E-47 | 4.74E-45 | up |
| Dysf | 2.60559 | 1.3816099 | 3.48E-47 | 7.35E-45 | up |
| Ccn2 | 8.315412 | 3.0557877 | 8.98E-47 | 1.88E-44 | up |
| Kifc1 | 7.508318 | 2.9084898 | 9.48E-47 | 1.97E-44 | up |
| Tubb6 | 6.247612 | 2.643305 | 1.42E-46 | 2.92E-44 | up |
| Tcf19 | 2.842442 | 1.5071309 | 2E-46 | 4.08E-44 | up |
| Fbln5 | 3.190107 | 1.6736048 | 2.08E-46 | 4.21E-44 | up |
| Arhgap11a | 4.300534 | 2.1045158 | 5.87E-46 | 1.17E-43 | up |
| P3h4 | 2.152552 | 1.106048 | 6.89E-46 | 1.37E-43 | up |
| Actg2 | 8.454654 | 3.0797457 | 1.55E-45 | 3.04E-43 | up |
| Rfc5 | 2.049232 | 1.0350832 | 1.77E-45 | 3.43E-43 | up |
| Mcm3 | 4.087523 | 2.0312269 | 1.78E-45 | 3.43E-43 | up |
| Col1a2 | 6.148506 | 2.6202359 | 1.94E-45 | 3.71E-43 | up |
| Mcm5 | 6.306266 | 2.656786 | 1.99E-45 | 3.78E-43 | up |
| Cdc6 | 7.780649 | 2.9598905 | 3.92E-45 | 7.38E-43 | up |
| Cep55 | 8.502932 | 3.0879603 | 6.63E-45 | 1.23E-42 | up |
| H2az1 | 2.055947 | 1.039803 | 7.31E-45 | 1.34E-42 | up |

| | | | | | |
|---|---|---|---|---|---|
| Lox | 5.633177 | 2.4939487 | 2.2E-44 | 4E-42 | up |
| Fen1 | 3.365593 | 1.7508608 | 2.27E-44 | 4.1E-42 | up |
| Pbk | 15.22511 | 3.9283808 | 3.46E-44 | 6.2E-42 | up |
| Fndc1 | 8.60043 | 3.1044088 | 3.76E-44 | 6.69E-42 | up |
| Oip5 | 7.84214 | 2.9712475 | 6.15E-44 | 1.09E-41 | up |
| Bard1 | 7.038623 | 2.8152932 | 7.84E-44 | 1.37E-41 | up |
| Cpz | 8.790084 | 3.135877 | 1.29E-43 | 2.25E-41 | up |
| Uhrf1 | 7.371756 | 2.8820083 | 2.8E-43 | 4.82E-41 | up |
| Osmr | 4.76536 | 2.2525851 | 3.75E-43 | 6.42E-41 | up |
| Cdc7 | 6.322901 | 2.6605866 | 5.52E-43 | 9.37E-41 | up |
| Col4a2 | 2.724333 | 1.445903 | 1.04E-42 | 1.76E-40 | up |
| Asf1b | 6.636936 | 2.7305173 | 1.71E-42 | 2.86E-40 | up |
| Ccnf | 3.813831 | 1.9312408 | 1.77E-42 | 2.94E-40 | up |
| Wscd2 | 5.156345 | 2.3663489 | 2.09E-42 | 3.45E-40 | up |
| Amtn | 8.701363 | 3.1212414 | 3.27E-42 | 5.35E-40 | up |
| Hjurp | 6.520496 | 2.7049816 | 3.49E-42 | 5.67E-40 | up |
| Plk4 | 6.35161 | 2.6671223 | 6.74E-42 | 1.09E-39 | up |
| Cdt1 | 5.14032 | 2.3618581 | 1.02E-41 | 1.63E-39 | up |
| Galnt14 | 0.089028 | -3.489593 | 1.02E-41 | 1.63E-39 | down |
| Lum | 6.261244 | 2.6464494 | 4.78E-41 | 7.54E-39 | up |
| Slc7a5 | 3.296503 | 1.7209366 | 6.82E-41 | 1.07E-38 | up |
| Lmnb1 | 4.475743 | 2.1621273 | 1.45E-40 | 2.24E-38 | up |
| Smtn | 2.863607 | 1.5178336 | 2.65E-40 | 4.07E-38 | up |
| Tgfb3 | 3.230614 | 1.6918085 | 3.24E-40 | 4.94E-38 | up |
| Exo1 | 12.39285 | 3.6314362 | 4.66E-40 | 7.06E-38 | up |
| Pcsk5 | 3.139925 | 1.6507303 | 9.21E-40 | 1.38E-37 | up |
| ENSRNOG00000067254 | 6.817301 | 2.7692007 | 1.41E-39 | 2.11E-37 | up |
| Ccno | 5.922757 | 2.566269 | 4.87E-39 | 7.17E-37 | up |
| Cthrc1 | 7.727588 | 2.9500181 | 5.06E-39 | 7.41E-37 | up |
| Aldh3a1 | 0.088448 | -3.499023 | 7.78E-39 | 1.13E-36 | down |
| Col3a1 | 4.708336 | 2.2352174 | 7.9E-39 | 1.14E-36 | up |
| Tacc3 | 4.40878 | 2.1403794 | 9.69E-39 | 1.39E-36 | up |
| Angptl2 | 2.962579 | 1.5668539 | 1.83E-38 | 2.62E-36 | up |
| Smc2 | 2.903173 | 1.5376307 | 1.96E-38 | 2.78E-36 | up |
| Nid2 | 2.370011 | 1.2448938 | 2.3E-38 | 3.25E-36 | up |
| Dkk3 | 2.275314 | 1.1860655 | 3.02E-38 | 4.23E-36 | up |
| Adamts12 | 3.531436 | 1.820255 | 4.51E-38 | 6.28E-36 | up |
| Aspm | 4.298955 | 2.1039861 | 5.77E-38 | 7.98E-36 | up |
| Scn3b | 4.830059 | 2.2720408 | 8.44E-38 | 1.16E-35 | up |
| Mettl7a | 0.358638 | -1.4794 | 9.86E-38 | 1.35E-35 | down |
| Cd163 | 7.963665 | 2.9934325 | 1.02E-37 | 1.38E-35 | up |

| | | | | | |
|---|---|---|---|---|---|
| Gja3 | 59.9895 | 5.9066382 | 1.55E-37 | 2.08E-35 | up |
| Adamts7 | 2.917004 | 1.5444874 | 1.64E-37 | 2.19E-35 | up |
| Rrm1 | 2.258006 | 1.1750494 | 2.4E-37 | 3.18E-35 | up |
| Pmf1 | 2.83853 | 1.5051441 | 3.87E-37 | 5.1E-35 | up |
| Plod2 | 5.287651 | 2.4026269 | 5.53E-37 | 7.24E-35 | up |
| Ccl2 | 12.86062 | 3.6848888 | 6.66E-37 | 8.67E-35 | up |
| Col6a2 | 3.228163 | 1.6907136 | 7.99E-37 | 1.04E-34 | up |
| Incenp | 3.778101 | 1.9176612 | 8.59E-37 | 1.11E-34 | up |
| Ticrr | 7.986157 | 2.9975015 | 1.39E-36 | 1.78E-34 | up |
| Cenpn | 4.121913 | 2.0433142 | 1.46E-36 | 1.86E-34 | up |
| Pcdh17 | 3.182006 | 1.6699363 | 1.67E-36 | 2.11E-34 | up |
| Rarres2 | 2.590309 | 1.3731242 | 1.95E-36 | 2.45E-34 | up |
| Atp1a2 | 0.148431 | -2.752131 | 1.95E-36 | 2.45E-34 | down |
| Ccne1 | 8.356557 | 3.0629086 | 2.3E-36 | 2.87E-34 | up |
| Knl1 | 6.168849 | 2.6250014 | 5.15E-36 | 6.34E-34 | up |
| Spc24 | 8.813969 | 3.1397919 | 6.4E-36 | 7.83E-34 | up |
| Mmp23 | 4.223134 | 2.0783142 | 6.74E-36 | 8.17E-34 | up |
| Grem1 | 23.44605 | 4.5512732 | 6.83E-36 | 8.23E-34 | up |
| Emp1 | 2.194132 | 1.1336502 | 9.68E-36 | 1.16E-33 | up |
| Spc25 | 6.91973 | 2.7907158 | 1.27E-35 | 1.52E-33 | up |
| Aoc1 | 25.10602 | 4.6499616 | 2.24E-35 | 2.66E-33 | up |
| Troap | 10.22226 | 3.3536426 | 2.92E-35 | 3.45E-33 | up |
| Tmem176a | 3.806923 | 1.9286256 | 4.85E-35 | 5.68E-33 | up |
| Mxd3 | 12.66589 | 3.6628766 | 1.36E-34 | 1.59E-32 | up |
| Tubb5 | 2.748237 | 1.4585064 | 1.46E-34 | 1.69E-32 | up |
| H2ax | 2.867625 | 1.5198566 | 1.79E-34 | 2.05E-32 | up |
| Trabd2b | 2.791848 | 1.4812204 | 2E-34 | 2.28E-32 | up |
| Cenpa | 8.840433 | 3.1441171 | 2.34E-34 | 2.66E-32 | up |
| Wdhd1 | 3.068802 | 1.6176755 | 2.35E-34 | 2.66E-32 | up |
| Mcm4 | 3.623242 | 1.857281 | 2.49E-34 | 2.8E-32 | up |
| Pxdn | 2.048766 | 1.0347554 | 3.73E-34 | 4.16E-32 | up |
| Col6a1 | 3.158181 | 1.6590941 | 4.62E-34 | 5.12E-32 | up |
| Ccnb2-ps2 | 5.642796 | 2.4964103 | 4.89E-34 | 5.4E-32 | up |
| Selenbp1 | 0.340445 | -1.554507 | 5.84E-34 | 6.42E-32 | down |
| Stmn1 | 3.074036 | 1.6201343 | 6.04E-34 | 6.61E-32 | up |
| Cenpw | 8.419622 | 3.0737555 | 9.06E-34 | 9.86E-32 | up |
| Pclaf | 8.369271 | 3.0651019 | 1.04E-33 | 1.13E-31 | up |
| Ercc6l | 6.789114 | 2.7632233 | 1.12E-33 | 1.21E-31 | up |
| Cyp3a9 | 0.370457 | -1.432623 | 1.2E-33 | 1.28E-31 | down |
| Lig1 | 2.22531 | 1.1540064 | 1.93E-33 | 2.05E-31 | up |
| Rfc3 | 2.285697 | 1.1926341 | 3.1E-33 | 3.27E-31 | up |

| | | | | | |
|---|---|---|---|---|---|
| Gas7 | 2.781522 | 1.4758745 | 6.22E-33 | 6.55E-31 | up |
| Ube2t | 7.078913 | 2.8235278 | 6.44E-33 | 6.74E-31 | up |
| Amigo1 | 0.405274 | -1.303029 | 7.53E-33 | 7.84E-31 | down |
| Kif22 | 34.35631 | 5.1025031 | 8.37E-33 | 8.69E-31 | up |
| Cald1 | 2.690038 | 1.4276264 | 1.09E-32 | 1.12E-30 | up |
| Ckap4 | 2.396148 | 1.2607169 | 1.15E-32 | 1.18E-30 | up |
| E2f8 | 9.654372 | 3.2711824 | 2.47E-32 | 2.51E-30 | up |
| Fscn1 | 3.562484 | 1.8328836 | 2.9E-32 | 2.94E-30 | up |
| Cpxm2 | 3.578397 | 1.8393133 | 5.32E-32 | 5.37E-30 | up |
| Reep1 | 2.311126 | 1.208596 | 5.7E-32 | 5.73E-30 | up |
| Cdca2 | 2.93198 | 1.551875 | 8.61E-32 | 8.58E-30 | up |
| Mnda | 3.817445 | 1.9326076 | 1.1E-31 | 1.09E-29 | up |
| Kif14 | 10.22325 | 3.3537823 | 1.27E-31 | 1.25E-29 | up |
| Adamts2 | 3.245627 | 1.6984971 | 1.96E-31 | 1.92E-29 | up |
| Pdlim7 | 2.508354 | 1.3267408 | 3.37E-31 | 3.3E-29 | up |
| Ncapd3 | 2.953043 | 1.5622023 | 3.48E-31 | 3.39E-29 | up |
| Ttk | 12.1343 | 3.6010191 | 4.65E-31 | 4.51E-29 | up |
| Espl1 | 7.838894 | 2.97065 | 7.23E-31 | 6.95E-29 | up |
| Pole2 | 5.464107 | 2.4499858 | 8.44E-31 | 8.08E-29 | up |
| Gas2l3 | 5.418508 | 2.4378958 | 9.82E-31 | 9.37E-29 | up |
| Cenpt | 4.161958 | 2.0572624 | 1.14E-30 | 1.08E-28 | up |
| Cldn1 | 0.260644 | -1.939849 | 1.24E-30 | 1.17E-28 | down |
| Kif15 | 4.678403 | 2.2260161 | 1.32E-30 | 1.24E-28 | up |
| Adam12 | 5.781231 | 2.5313768 | 1.33E-30 | 1.25E-28 | up |
| Arhgap19 | 3.306591 | 1.7253446 | 1.39E-30 | 1.3E-28 | up |
| Gpsm2 | 2.617498 | 1.3881885 | 1.43E-30 | 1.33E-28 | up |
| Rbl1 | 2.671645 | 1.4177286 | 1.54E-30 | 1.43E-28 | up |
| Kif23 | 2.653489 | 1.4078907 | 1.85E-30 | 1.71E-28 | up |
| Dhfr | 3.362759 | 1.7496455 | 1.93E-30 | 1.78E-28 | up |
| Tmem176b | 2.91132 | 1.5416732 | 2.11E-30 | 1.93E-28 | up |
| Depdc1 | 14.22079 | 3.8299292 | 2.62E-30 | 2.38E-28 | up |
| Basp1 | 3.815683 | 1.9319412 | 3.47E-30 | 3.15E-28 | up |
| P3h3 | 2.278546 | 1.1881135 | 4.64E-30 | 4.2E-28 | up |
| Cibar1 | 2.088279 | 1.0623143 | 7.54E-30 | 6.78E-28 | up |
| Fam114a1 | 2.331351 | 1.2211664 | 1.05E-29 | 9.46E-28 | up |
| Trip13 | 5.075477 | 2.3435435 | 1.2E-29 | 1.07E-27 | up |
| Pola1 | 2.644465 | 1.4029757 | 1.25E-29 | 1.11E-27 | up |
| Ces1d | 0.316257 | -1.660829 | 1.31E-29 | 1.16E-27 | down |
| LOC291863 | 0.411567 | -1.280801 | 1.99E-29 | 1.75E-27 | down |
| Gprin3 | 0.287934 | -1.796192 | 2.79E-29 | 2.44E-27 | down |
| Aurka | 3.21505 | 1.684841 | 3.3E-29 | 2.88E-27 | up |

| | | | | | |
|---|---|---|---|---|---|
| Cep72 | 8.834456 | 3.1431412 | 3.46E-29 | 3.01E-27 | up |
| Nasp | 2.18147 | 1.1253008 | 4.71E-29 | 4.06E-27 | up |
| Recql4 | 5.359554 | 2.4221129 | 5.68E-29 | 4.86E-27 | up |
| Lratd1 | 0.320083 | -1.643483 | 5.99E-29 | 5.11E-27 | down |
| Anlnl1 | 10.01869 | 3.3246213 | 7.92E-29 | 6.72E-27 | up |
| Gyg1 | 2.455846 | 1.2962202 | 8.48E-29 | 7.17E-27 | up |
| Sgo1 | 11.52116 | 3.5262138 | 1.42E-28 | 1.2E-26 | up |
| Fbxo40 | 0.187928 | -2.411751 | 1.67E-28 | 1.4E-26 | down |
| Mmp2 | 2.634153 | 1.3973391 | 1.98E-28 | 1.65E-26 | up |
| Fmo2 | 0.179696 | -2.47637 | 2.13E-28 | 1.78E-26 | down |
| Cenpi | 7.911906 | 2.9840253 | 2.51E-28 | 2.08E-26 | up |
| Cenpq | 5.616339 | 2.4896299 | 2.6E-28 | 2.15E-26 | up |
| Sapcd2 | 8.313249 | 3.0554125 | 2.92E-28 | 2.4E-26 | up |
| Cdc45 | 7.805475 | 2.9644865 | 3.14E-28 | 2.58E-26 | up |
| Cenpu | 4.584835 | 2.1968698 | 3.48E-28 | 2.84E-26 | up |
| Fstl3 | 3.486535 | 1.8017939 | 4.9E-28 | 3.99E-26 | up |
| Nipsnap3a | 3.708517 | 1.8908423 | 5.03E-28 | 4.09E-26 | up |
| Col12a1 | 2.731236 | 1.4495538 | 5.31E-28 | 4.29E-26 | up |
| Rcn3 | 2.407903 | 1.2677775 | 5.43E-28 | 4.38E-26 | up |
| Gpr153 | 2.56164 | 1.3570678 | 6.42E-28 | 5.16E-26 | up |
| Trim13 | 0.346042 | -1.53098 | 7.12E-28 | 5.7E-26 | down |
| Itga2 | 4.403825 | 2.138757 | 1.02E-27 | 8.05E-26 | up |
| Lgals1 | 2.523071 | 1.3351806 | 1.06E-27 | 8.35E-26 | up |
| Rad51ap1 | 5.08111 | 2.3451436 | 1.14E-27 | 8.98E-26 | up |
| Chaf1b | 9.442616 | 3.2391867 | 1.8E-27 | 1.41E-25 | up |
| Rab32 | 2.347596 | 1.2311839 | 2.06E-27 | 1.61E-25 | up |
| C1qb | 4.804144 | 2.2642795 | 2.24E-27 | 1.75E-25 | up |
| LOC100359539 | 6.833284 | 2.7725791 | 2.63E-27 | 2.04E-25 | up |
| Nr3c2 | 0.467371 | -1.09736 | 3.63E-27 | 2.78E-25 | down |
| Chaf1a | 3.372461 | 1.7538018 | 6.62E-27 | 5.03E-25 | up |
| Thbs2 | 17.52232 | 4.1311221 | 9.51E-27 | 7.2E-25 | up |
| Serpinh1 | 2.193908 | 1.1335029 | 1.01E-26 | 7.65E-25 | up |
| Cp | 2.685281 | 1.4250731 | 1.31E-26 | 9.83E-25 | up |
| Fmo1 | 0.380147 | -1.395372 | 1.31E-26 | 9.83E-25 | down |
| LOC691995 | 4.222731 | 2.0781763 | 1.73E-26 | 1.29E-24 | up |
| C2h4orf46 | 3.979335 | 1.9925275 | 1.99E-26 | 1.48E-24 | up |
| Pycr1 | 5.454303 | 2.4473949 | 2.07E-26 | 1.54E-24 | up |
| Mcam | 2.466276 | 1.3023344 | 2.54E-26 | 1.87E-24 | up |
| Clec10a | 4.183569 | 2.0647343 | 2.76E-26 | 2.03E-24 | up |
| Wdr62 | 3.095292 | 1.6300757 | 3.51E-26 | 2.56E-24 | up |
| Gpx7 | 2.584147 | 1.369688 | 3.78E-26 | 2.75E-24 | up |

| | | | | | |
|---|---|---|---|---|---|
| Adamts1 | 2.952376 | 1.5618766 | 5.11E-26 | 3.68E-24 | up |
| Gpr88 | 4.460598 | 2.1572372 | 1.27E-25 | 9.09E-24 | up |
| E2f7 | 8.463323 | 3.0812242 | 1.38E-25 | 9.82E-24 | up |
| Clec11a | 6.170987 | 2.6255013 | 1.57E-25 | 1.12E-23 | up |
| Svep1 | 3.36722 | 1.751558 | 1.68E-25 | 1.19E-23 | up |
| Flrt2 | 2.651039 | 1.4065581 | 2.51E-25 | 1.77E-23 | up |
| Igfbp4 | 3.949744 | 1.9817591 | 2.96E-25 | 2.07E-23 | up |
| Cdkn2c | 3.32622 | 1.7338834 | 3.32E-25 | 2.32E-23 | up |
| Des | 3.670144 | 1.8758366 | 5.07E-25 | 3.53E-23 | up |
| Ssc5d | 2.512365 | 1.3290462 | 5.47E-25 | 3.78E-23 | up |
| Cenph | 7.45375 | 2.8979664 | 6.7E-25 | 4.61E-23 | up |
| Mybl1 | 3.510765 | 1.8117853 | 8.4E-25 | 5.75E-23 | up |
| Ybx2 | 4.662355 | 2.2210587 | 9.05E-25 | 6.18E-23 | up |
| Sema6b | 2.509731 | 1.327533 | 1.45E-24 | 9.85E-23 | up |
| Adamts9 | 2.510498 | 1.3279735 | 2.54E-24 | 1.71E-22 | up |
| Sgo2 | 9.404238 | 3.233311 | 2.56E-24 | 1.72E-22 | up |
| C10h17orf58 | 0.427412 | -1.2263 | 3.39E-24 | 2.26E-22 | down |
| Cyp2f4 | 0.225179 | -2.150856 | 3.73E-24 | 2.48E-22 | down |
| Mthfd2 | 2.968184 | 1.5695807 | 3.89E-24 | 2.57E-22 | up |
| Ncapg | 5.103447 | 2.3514719 | 4.77E-24 | 3.13E-22 | up |
| Haus4 | 3.55326 | 1.8291434 | 4.87E-24 | 3.19E-22 | up |
| ENSRNOG00000069437 | 13.66862 | 3.7727955 | 6.76E-24 | 4.4E-22 | up |
| Sertad4 | 5.460438 | 2.4490166 | 6.98E-24 | 4.53E-22 | up |
| Adamts6 | 2.771097 | 1.470457 | 7.48E-24 | 4.85E-22 | up |
| Aldh18a1 | 2.19443 | 1.1338462 | 8.2E-24 | 5.3E-22 | up |
| Fkbp11 | 2.129131 | 1.090265 | 9.46E-24 | 6.09E-22 | up |
| Tagln | 3.149907 | 1.6553094 | 9.79E-24 | 6.29E-22 | up |
| Apcdd1 | 2.193492 | 1.1332295 | 1.23E-23 | 7.87E-22 | up |
| Topbp1 | 2.014349 | 1.0103134 | 1.3E-23 | 8.27E-22 | up |
| Rad54l | 4.798664 | 2.2626328 | 1.39E-23 | 8.85E-22 | up |
| Lamc1 | 2.103389 | 1.0727154 | 1.83E-23 | 1.16E-21 | up |
| Irs3 | 0.240228 | -2.057521 | 1.96E-23 | 1.24E-21 | down |
| Nsl1 | 6.074001 | 2.6026472 | 1.97E-23 | 1.24E-21 | up |
| Rtkn2 | 0.448716 | -1.156127 | 2.09E-23 | 1.31E-21 | down |
| Smc4 | 2.634748 | 1.3976651 | 2.15E-23 | 1.35E-21 | up |
| Lmnb2 | 3.077917 | 1.6219542 | 2.64E-23 | 1.65E-21 | up |
| Ska1 | 13.65856 | 3.7717338 | 2.66E-23 | 1.66E-21 | up |
| Ncapg2 | 2.930846 | 1.5513171 | 2.88E-23 | 1.79E-21 | up |
| Emilin1 | 2.170258 | 1.1178667 | 3.69E-23 | 2.28E-21 | up |
| Gatm | 2.327449 | 1.2187498 | 4.06E-23 | 2.5E-21 | up |
| Rassf10 | 0.256072 | -1.965381 | 6.11E-23 | 3.72E-21 | down |

| | | | | | |
|---|---|---|---|---|---|
| Cercam | 2.893759 | 1.5329448 | 6.56E-23 | 3.97E-21 | up |
| Loxl3 | 2.790019 | 1.4802752 | 7.69E-23 | 4.65E-21 | up |
| Hacd1 | 2.082378 | 1.0582323 | 7.78E-23 | 4.68E-21 | up |
| Vcam1 | 2.891231 | 1.5316839 | 1.15E-22 | 6.87E-21 | up |
| E2f1 | 2.451586 | 1.2937157 | 1.43E-22 | 8.56E-21 | up |
| Fam13a | 0.427137 | -1.22723 | 1.75E-22 | 1.03E-20 | down |
| Chek2 | 2.62727 | 1.3935646 | 1.76E-22 | 1.04E-20 | up |
| Glod5 | 4.713964 | 2.2369408 | 1.94E-22 | 1.14E-20 | up |
| Sult1c3 | 26.18376 | 4.7106005 | 2.17E-22 | 1.28E-20 | up |
| Esco2 | 9.491151 | 3.2465831 | 2.29E-22 | 1.34E-20 | up |
| Sec14l3 | 0.37768 | -1.404762 | 2.89E-22 | 1.68E-20 | down |
| Cyp2b1 | 0.313293 | -1.674416 | 3.07E-22 | 1.79E-20 | down |
| Fanca | 4.388475 | 2.1337196 | 5.51E-22 | 3.18E-20 | up |
| Dctpp1 | 3.41743 | 1.7729118 | 6.3E-22 | 3.63E-20 | up |
| C7 | 3.071036 | 1.6187253 | 6.35E-22 | 3.65E-20 | up |
| C1qc | 4.11858 | 2.0421469 | 6.36E-22 | 3.65E-20 | up |
| Spred3 | 4.520666 | 2.1765354 | 7.18E-22 | 4.09E-20 | up |
| LOC120098765 | 4.010236 | 2.0036873 | 7.25E-22 | 4.12E-20 | up |
| Pla2g2d | 0.459401 | -1.122173 | 7.5E-22 | 4.25E-20 | down |
| Vwa1 | 3.086135 | 1.625801 | 7.88E-22 | 4.46E-20 | up |
| Fgf23 | 13554.41 | 13.726475 | 7.95E-22 | 4.49E-20 | up |
| Clec3b | 2.449349 | 1.2923981 | 9.2E-22 | 5.18E-20 | up |
| Prrx2 | 3.309093 | 1.7264359 | 9.78E-22 | 5.49E-20 | up |
| Arhgef39 | 11.53102 | 3.5274478 | 1.03E-21 | 5.78E-20 | up |
| C1qtnf6 | 4.954142 | 2.3086351 | 1.15E-21 | 6.41E-20 | up |
| Dbn1 | 3.332854 | 1.7367581 | 1.52E-21 | 8.4E-20 | up |
| Pcolce | 3.63567 | 1.8622213 | 1.56E-21 | 8.65E-20 | up |
| Arsi | 2.398347 | 1.2620403 | 1.74E-21 | 9.6E-20 | up |
| Insyn1 | 2.667407 | 1.4154381 | 2.02E-21 | 1.11E-19 | up |
| Igdcc4 | 3.166247 | 1.6627738 | 2.31E-21 | 1.27E-19 | up |
| Numbl | 2.436181 | 1.2846211 | 2.53E-21 | 1.38E-19 | up |
| Galnt15 | 0.31016 | -1.688917 | 3.01E-21 | 1.64E-19 | down |
| Dsn1 | 6.81719 | 2.7691772 | 4.06E-21 | 2.2E-19 | up |
| Scgb1a1 | 0.362766 | -1.462888 | 4.06E-21 | 2.2E-19 | down |
| Traip | 6.988194 | 2.8049197 | 4.34E-21 | 2.34E-19 | up |
| Mis18bp1 | 3.38881 | 1.7607786 | 4.34E-21 | 2.34E-19 | up |
| Ankle1 | 17.13051 | 4.098496 | 6.09E-21 | 3.27E-19 | up |
| Poc1a | 3.158027 | 1.6590236 | 8.21E-21 | 4.38E-19 | up |
| Chtf18 | 3.020291 | 1.5946876 | 1.02E-20 | 5.4E-19 | up |
| Tyms | 3.018355 | 1.5937623 | 1.35E-20 | 7.07E-19 | up |
| Psrc1 | 8.194354 | 3.0346302 | 1.46E-20 | 7.62E-19 | up |

| | | | | | |
|---|---|---|---|---|---|
| Kcnip2 | 0.243163 | -2.040002 | 1.61E-20 | 8.43E-19 | down |
| Cox7a1 | 0.315986 | -1.662068 | 2.63E-20 | 1.37E-18 | down |
| Aunip | 6.750973 | 2.7550955 | 2.8E-20 | 1.45E-18 | up |
| Sdk1 | 2.389809 | 1.2568953 | 2.92E-20 | 1.51E-18 | up |
| Itga9 | 2.222098 | 1.1519222 | 3.22E-20 | 1.66E-18 | up |
| Dhrs3 | 0.466987 | -1.098547 | 3.83E-20 | 1.97E-18 | down |
| Hunk | 4.278794 | 2.0972042 | 4.01E-20 | 2.05E-18 | up |
| Pld4 | 2.32186 | 1.2152812 | 4.73E-20 | 2.4E-18 | up |
| Col6a6 | 2.922327 | 1.5471178 | 4.77E-20 | 2.42E-18 | up |
| Macrod1 | 0.487182 | -1.037467 | 5.22E-20 | 2.64E-18 | down |
| Dnaja4 | 0.457509 | -1.128128 | 5.64E-20 | 2.84E-18 | down |
| ENSRNOG00000070694 | 5.421514 | 2.4386958 | 5.68E-20 | 2.85E-18 | up |
| Mcm10 | 5.563042 | 2.4758739 | 6.87E-20 | 3.45E-18 | up |
| Cnn1 | 4.29125 | 2.101398 | 7.44E-20 | 3.71E-18 | up |
| Fibin | 2.467451 | 1.3030215 | 7.92E-20 | 3.94E-18 | up |
| Col15a1 | 4.130823 | 2.0464294 | 8.81E-20 | 4.38E-18 | up |
| Prc1 | 2.5168 | 1.3315904 | 1.18E-19 | 5.8E-18 | up |
| Vegfc | 2.239642 | 1.1632679 | 1.18E-19 | 5.81E-18 | up |
| Gpr34 | 3.635275 | 1.8620645 | 1.34E-19 | 6.52E-18 | up |
| Slc39a2 | 0.415019 | -1.26875 | 1.74E-19 | 8.47E-18 | down |
| Lin9 | 2.287576 | 1.1938194 | 1.77E-19 | 8.59E-18 | up |
| Depdc7 | 3.345188 | 1.7420871 | 1.92E-19 | 9.28E-18 | up |
| Cdh11 | 2.884246 | 1.5281943 | 2.51E-19 | 1.2E-17 | up |
| Arid5a | 2.284336 | 1.1917751 | 2.61E-19 | 1.25E-17 | up |
| Pmepa1 | 2.457756 | 1.2973416 | 2.83E-19 | 1.35E-17 | up |
| Gen1 | 4.306996 | 2.1066821 | 3.36E-19 | 1.6E-17 | up |
| Myl9 | 2.225256 | 1.1539712 | 3.76E-19 | 1.78E-17 | up |
| C1qa | 4.449427 | 2.1536195 | 3.84E-19 | 1.81E-17 | up |
| Nt5dc2 | 2.178554 | 1.1233709 | 3.89E-19 | 1.83E-17 | up |
| Nrip3 | 3.445072 | 1.784534 | 4.03E-19 | 1.89E-17 | up |
| Timp1 | 4.056017 | 2.0200636 | 4.57E-19 | 2.14E-17 | up |
| ENSRNOG00000063706 | 4.214147 | 2.0752408 | 4.75E-19 | 2.22E-17 | up |
| Zfp367 | 4.10457 | 2.0372311 | 7.14E-19 | 3.31E-17 | up |
| Shisa8 | 5.452896 | 2.4470225 | 7.24E-19 | 3.34E-17 | up |
| Cgref1 | 6.891698 | 2.7848595 | 7.95E-19 | 3.67E-17 | up |
| P4ha3 | 2.564299 | 1.3585645 | 8.65E-19 | 3.97E-17 | up |
| Orc1 | 6.925222 | 2.7918603 | 1.07E-18 | 4.89E-17 | up |
| Ccdc198 | 0.209276 | -2.256523 | 1.12E-18 | 5.12E-17 | down |
| Slc15a2 | 0.260211 | -1.942247 | 1.71E-18 | 7.72E-17 | down |
| Gpt | 0.311485 | -1.682767 | 2.08E-18 | 9.38E-17 | down |
| Cyp2s1 | 0.303044 | -1.7224 | 2.1E-18 | 9.49E-17 | down |

| | | | | | |
|---|---|---|---|---|---|
| Osr1 | 2.434802 | 1.2838045 | 2.15E-18 | 9.66E-17 | up |
| Vgll3 | 2.904739 | 1.5384083 | 2.16E-18 | 9.69E-17 | up |
| Mad2l1 | 3.244204 | 1.6978645 | 2.3E-18 | 1.03E-16 | up |
| LOC100359539 | 7.671964 | 2.9395959 | 2.36E-18 | 1.05E-16 | up |
| Flrt3 | 0.465273 | -1.103851 | 2.38E-18 | 1.06E-16 | down |
| Col7a1 | 4.007458 | 2.0026873 | 2.5E-18 | 1.11E-16 | up |
| Fap | 9.199761 | 3.2015963 | 3.51E-18 | 1.55E-16 | up |
| Ppp1r1b | 0.329052 | -1.603611 | 4.36E-18 | 1.92E-16 | down |
| Folr2 | 7.325034 | 2.8728355 | 4.47E-18 | 1.96E-16 | up |
| Xkr5 | 9.786296 | 3.2907629 | 4.49E-18 | 1.97E-16 | up |
| Tppp3 | 0.481073 | -1.055673 | 4.73E-18 | 2.07E-16 | down |
| Tnfrsf8 | 0.270143 | -1.888207 | 5.64E-18 | 2.46E-16 | down |
| Frem1 | 3.245518 | 1.6984488 | 5.84E-18 | 2.55E-16 | up |
| Ptn | 4.524149 | 2.1776466 | 7.06E-18 | 3.05E-16 | up |
| Dscc1 | 6.583004 | 2.7187461 | 7.47E-18 | 3.22E-16 | up |
| Olfm2 | 8.98537 | 3.1675779 | 8.1E-18 | 3.49E-16 | up |
| Zrsr2-ps1 | 0.498911 | -1.003146 | 1.06E-17 | 4.53E-16 | down |
| Acta1 | 11.05263 | 3.4663173 | 1.35E-17 | 5.74E-16 | up |
| Serpinf1 | 3.206232 | 1.6808789 | 1.36E-17 | 5.81E-16 | up |
| C1qtnf5 | 2.690089 | 1.4276541 | 1.41E-17 | 6.01E-16 | up |
| Klf15 | 0.337777 | -1.565855 | 1.48E-17 | 6.29E-16 | down |
| Haspin | 2.036127 | 1.0258273 | 1.69E-17 | 7.18E-16 | up |
| Foxn4 | 7.147339 | 2.8374062 | 2.2E-17 | 9.24E-16 | up |
| C9h6orf132 | 0.4976 | -1.00694 | 2.42E-17 | 1.01E-15 | down |
| Ltbp3 | 2.14036 | 1.0978535 | 2.93E-17 | 1.23E-15 | up |
| Vegfd | 3.98254 | 1.9936888 | 3.33E-17 | 1.38E-15 | up |
| Tedc2 | 3.320896 | 1.7315724 | 4.73E-17 | 1.96E-15 | up |
| Timeless | 2.316982 | 1.2122467 | 4.74E-17 | 1.96E-15 | up |
| Scx | 2.708852 | 1.4376816 | 5.14E-17 | 2.12E-15 | up |
| Prr11 | 7.255059 | 2.8589874 | 6.12E-17 | 2.52E-15 | up |
| Jph2 | 2.197205 | 1.1356693 | 6.45E-17 | 2.65E-15 | up |
| Pparg | 0.464668 | -1.105726 | 7.48E-17 | 3.07E-15 | down |
| Mmp17 | 2.281538 | 1.1900068 | 7.54E-17 | 3.08E-15 | up |
| Adamts8 | 2.283371 | 1.1911652 | 7.83E-17 | 3.19E-15 | up |
| Gins1 | 4.566623 | 2.1911276 | 8.41E-17 | 3.42E-15 | up |
| Rasl11b | 4.005167 | 2.0018623 | 8.48E-17 | 3.44E-15 | up |
| Kcnrg | 0.485207 | -1.043328 | 8.97E-17 | 3.63E-15 | down |
| Deup1 | 2.467914 | 1.303292 | 9.54E-17 | 3.85E-15 | up |
| Krt76 | 0.29606 | -1.75604 | 1.14E-16 | 4.57E-15 | down |
| Sox9 | 0.401583 | -1.31623 | 1.18E-16 | 4.75E-15 | down |
| Col16a1 | 2.422433 | 1.2764568 | 1.27E-16 | 5.08E-15 | up |

| | | | | | |
|---|---|---|---|---|---|
| Mgp | 2.19468 | 1.1340107 | 1.39E-16 | 5.55E-15 | up |
| LOC108348201 | 3.274068 | 1.7110844 | 1.5E-16 | 6E-15 | up |
| Mindy2 | 0.438559 | -1.189158 | 1.72E-16 | 6.85E-15 | down |
| Ntrk2 | 4.081006 | 2.028925 | 2.34E-16 | 9.21E-15 | up |
| Dhcr7 | 2.327288 | 1.2186495 | 2.41E-16 | 9.49E-15 | up |
| Ttc34 | 0.370225 | -1.433525 | 2.57E-16 | 1.01E-14 | down |
| Pllp | 0.395253 | -1.339151 | 2.61E-16 | 1.02E-14 | down |
| Prob1 | 0.431149 | -1.213742 | 2.83E-16 | 1.1E-14 | down |
| Atad2 | 2.258563 | 1.1754051 | 3.17E-16 | 1.23E-14 | up |
| Snrpa | 2.195995 | 1.1348745 | 3.29E-16 | 1.28E-14 | up |
| Mdk | 2.118605 | 1.0831145 | 3.92E-16 | 1.51E-14 | up |
| Rexo5 | 4.303829 | 2.1056208 | 4.09E-16 | 1.57E-14 | up |
| Macroh2a2 | 2.944845 | 1.5581918 | 5.01E-16 | 1.92E-14 | up |
| Tpm2 | 2.53556 | 1.3423044 | 6.54E-16 | 2.5E-14 | up |
| Ddias | 5.062897 | 2.3399633 | 6.66E-16 | 2.54E-14 | up |
| Cntrob | 2.313503 | 1.2100792 | 1.08E-15 | 4.07E-14 | up |
| Ezh2 | 2.952288 | 1.5618333 | 1.16E-15 | 4.36E-14 | up |
| ENSRNOG00000064849 | 0.00029 | -11.74989 | 1.17E-15 | 4.39E-14 | down |
| Negr1 | 0.352445 | -1.504532 | 1.22E-15 | 4.55E-14 | down |
| Adgrv1 | 0.404104 | -1.307202 | 1.22E-15 | 4.55E-14 | down |
| Dse | 2.167998 | 1.1163632 | 1.31E-15 | 4.87E-14 | up |
| Ptch2 | 2.819082 | 1.4952254 | 1.55E-15 | 5.74E-14 | up |
| Slbp | 2.02697 | 1.0193249 | 1.56E-15 | 5.78E-14 | up |
| Chek1 | 3.548754 | 1.8273125 | 1.6E-15 | 5.91E-14 | up |
| Exph5 | 0.435991 | -1.197631 | 1.62E-15 | 5.94E-14 | down |
| Gstm7 | 0.399826 | -1.322557 | 1.67E-15 | 6.1E-14 | down |
| Rad51c | 2.83581 | 1.5037609 | 1.7E-15 | 6.21E-14 | up |
| Rfc4 | 2.224907 | 1.1537449 | 1.88E-15 | 6.88E-14 | up |
| Tubb3 | 9.379055 | 3.2294426 | 2.05E-15 | 7.47E-14 | up |
| Slc25a29 | 0.478454 | -1.063547 | 2.66E-15 | 9.65E-14 | down |
| Adhfe1 | 0.290255 | -1.784606 | 2.72E-15 | 9.82E-14 | down |
| Depdc1b | 3.956258 | 1.9841365 | 3.11E-15 | 1.12E-13 | up |
| Rbm3 | 2.399134 | 1.2625139 | 3.41E-15 | 1.22E-13 | up |
| Aaas | 2.045789 | 1.0326575 | 3.5E-15 | 1.26E-13 | up |
| Atad5 | 3.276269 | 1.7120539 | 3.62E-15 | 1.29E-13 | up |
| Colec12 | 2.042716 | 1.030489 | 3.7E-15 | 1.32E-13 | up |
| Fjx1 | 2.65543 | 1.4089458 | 3.8E-15 | 1.35E-13 | up |
| Fsbp | 5.118197 | 2.3556358 | 3.98E-15 | 1.41E-13 | up |
| Acsm5 | 0.407947 | -1.293548 | 4.29E-15 | 1.52E-13 | down |
| Pnma8b | 0.420559 | -1.249619 | 4.42E-15 | 1.56E-13 | down |
| Olah | 3.284263 | 1.7155697 | 5.62E-15 | 1.98E-13 | up |

| | | | | | |
|---|---|---|---|---|---|
| Gja4 | 2.131205 | 1.0916692 | 6.32E-15 | 2.22E-13 | up |
| Slc41a2 | 2.553921 | 1.352714 | 6.47E-15 | 2.26E-13 | up |
| Gadd45g | 2.836084 | 1.5039001 | 6.6E-15 | 2.31E-13 | up |
| Ecm2 | 0.443051 | -1.174454 | 7.17E-15 | 2.49E-13 | down |
| LOC361346 | 2.56122 | 1.3568311 | 7.39E-15 | 2.57E-13 | up |
| Ada | 2.374592 | 1.2476799 | 7.55E-15 | 2.62E-13 | up |
| Ror2 | 2.848816 | 1.5103624 | 7.6E-15 | 2.63E-13 | up |
| Azgp1 | 0.282876 | -1.821757 | 8.35E-15 | 2.88E-13 | down |
| Fmo3 | 0.329401 | -1.602082 | 8.56E-15 | 2.95E-13 | down |
| Ncam1 | 4.809076 | 2.2657598 | 9.09E-15 | 3.11E-13 | up |
| Dlg2 | 4.310235 | 2.1077664 | 1.08E-14 | 3.66E-13 | up |
| Emcn | 2.060463 | 1.0429686 | 1.2E-14 | 4.09E-13 | up |
| Hrob | 2.903396 | 1.5377412 | 1.26E-14 | 4.26E-13 | up |
| Gins3 | 3.309642 | 1.7266753 | 1.27E-14 | 4.31E-13 | up |
| Cbx7 | 0.400753 | -1.319215 | 1.41E-14 | 4.75E-13 | down |
| Slit2 | 2.030973 | 1.0221713 | 1.71E-14 | 5.72E-13 | up |
| Il6 | 95.39025 | 6.5757699 | 2.03E-14 | 6.76E-13 | up |
| Hells | 3.536396 | 1.8222797 | 2.03E-14 | 6.77E-13 | up |
| Cep128 | 2.065065 | 1.0461874 | 2.29E-14 | 7.6E-13 | up |
| Sema3c | 2.025186 | 1.0180547 | 2.39E-14 | 7.91E-13 | up |
| Prim1 | 2.306998 | 1.206017 | 2.71E-14 | 8.96E-13 | up |
| Cip2a | 6.258248 | 2.6457588 | 2.87E-14 | 9.44E-13 | up |
| Bhlhe40 | 3.096506 | 1.6306414 | 3.21E-14 | 1.05E-12 | up |
| Nos3 | 2.021281 | 1.0152697 | 3.49E-14 | 1.14E-12 | up |
| Adam19 | 2.433773 | 1.2831944 | 3.57E-14 | 1.16E-12 | up |
| Upk1a | 0.192079 | -2.380228 | 3.7E-14 | 1.2E-12 | down |
| Mis18a | 7.533469 | 2.9133144 | 4.29E-14 | 1.39E-12 | up |
| PCOLCE2 | 0.344769 | -1.5363 | 4.35E-14 | 1.41E-12 | down |
| Armcx4 | 2.958588 | 1.5649089 | 4.47E-14 | 1.44E-12 | up |
| Pkmyt1 | 3.333326 | 1.7369624 | 5.09E-14 | 1.63E-12 | up |
| Fam83d | 2.401331 | 1.263834 | 5.29E-14 | 1.69E-12 | up |
| Klhl29 | 3.792233 | 1.9230476 | 6.56E-14 | 2.09E-12 | up |
| Fcrl2 | 74.33681 | 6.2160049 | 6.71E-14 | 2.14E-12 | up |
| Fbxo5 | 4.210811 | 2.074098 | 7.09E-14 | 2.25E-12 | up |
| ENSRNOG00000066693 | 0.000257 | -11.92373 | 7.42E-14 | 2.35E-12 | down |
| Brinp2 | 10.00732 | 3.3229832 | 7.49E-14 | 2.37E-12 | up |
| Adra1a | 0.397248 | -1.331888 | 7.73E-14 | 2.44E-12 | down |
| Cenpf | 2.547548 | 1.3491093 | 7.85E-14 | 2.48E-12 | up |
| ENSRNOG00000065596 | 63321.99 | 15.950419 | 8.27E-14 | 2.6E-12 | up |
| C9 | 2.638804 | 1.399884 | 8.35E-14 | 2.62E-12 | up |
| Kcne4 | 3.189548 | 1.673352 | 8.51E-14 | 2.67E-12 | up |

| | | | | | |
|---|---|---|---|---|---|
| Mcph1 | 2.818556 | 1.4949565 | 9.63E-14 | 3.01E-12 | up |
| H1f0 | 2.243454 | 1.1657214 | 1.01E-13 | 3.14E-12 | up |
| Orc6 | 2.949069 | 1.5602597 | 1.17E-13 | 3.61E-12 | up |
| Ccne2 | 3.301105 | 1.7229491 | 1.19E-13 | 3.67E-12 | up |
| Crlf1 | 2.375533 | 1.248251 | 1.28E-13 | 3.92E-12 | up |
| Lca5 | 0.483091 | -1.049632 | 1.69E-13 | 5.17E-12 | down |
| Osgin1 | 0.490432 | -1.027876 | 1.73E-13 | 5.27E-12 | down |
| Erfe | 8.81404 | 3.1398034 | 1.76E-13 | 5.35E-12 | up |
| Suv39h1 | 2.092442 | 1.0651879 | 1.82E-13 | 5.52E-12 | up |
| C1qtnf3 | 47.81298 | 5.5793305 | 1.98E-13 | 6E-12 | up |
| Slc23a1 | 0.47274 | -1.080882 | 2.03E-13 | 6.15E-12 | down |
| Brip1 | 5.608807 | 2.487694 | 2.04E-13 | 6.15E-12 | up |
| Gnaz | 0.380738 | -1.393128 | 2.12E-13 | 6.4E-12 | down |
| Mfge8 | 2.53176 | 1.3401405 | 2.72E-13 | 8.16E-12 | up |
| Skp2 | 3.115209 | 1.6393292 | 2.77E-13 | 8.28E-12 | up |
| ENSRNOG00000069319 | 0.000847 | -10.20561 | 3.05E-13 | 9.12E-12 | down |
| Tbpl1 | 10567.11 | 13.367294 | 3.64E-13 | 1.08E-11 | up |
| Neurl1b | 2.094694 | 1.0667393 | 3.66E-13 | 1.08E-11 | up |
| Mmp14 | 2.631454 | 1.3958603 | 4.8E-13 | 1.41E-11 | up |
| Tril | 0.415495 | -1.267096 | 5.24E-13 | 1.54E-11 | down |
| Pcdh7 | 2.49708 | 1.3202419 | 5.38E-13 | 1.58E-11 | up |
| Tmem132c | 2.600402 | 1.3787345 | 6.48E-13 | 1.88E-11 | up |
| LOC100362814 | 3.358846 | 1.7479658 | 6.58E-13 | 1.9E-11 | up |
| Slc2a12 | 0.494305 | -1.016527 | 6.84E-13 | 1.97E-11 | down |
| Tmem132e | 5.123295 | 2.3570721 | 7.19E-13 | 2.06E-11 | up |
| ENSRNOG00000069825 | 0.001157 | -9.755241 | 7.56E-13 | 2.16E-11 | down |
| Rrlt | 12.68495 | 3.665046 | 1.13E-12 | 3.19E-11 | up |
| Asgr2 | 6.034325 | 2.5931924 | 1.14E-12 | 3.22E-11 | up |
| Six4 | 0.473152 | -1.079624 | 1.18E-12 | 3.34E-11 | down |
| Ackr4 | 0.420284 | -1.250565 | 1.2E-12 | 3.37E-11 | down |
| Col8a1 | 3.814306 | 1.9314204 | 1.23E-12 | 3.46E-11 | up |
| ENSRNOG00000069255 | 0.000476 | -11.03794 | 1.33E-12 | 3.73E-11 | down |
| Tmem59l | 5.074821 | 2.3433571 | 1.45E-12 | 4.03E-11 | up |
| Tuba8 | 7.599994 | 2.9259982 | 1.45E-12 | 4.04E-11 | up |
| Mtbp | 3.250955 | 1.7008637 | 1.46E-12 | 4.05E-11 | up |
| Ccdc34 | 2.12366 | 1.0865529 | 1.47E-12 | 4.07E-11 | up |
| Itga8 | 2.164502 | 1.114035 | 1.56E-12 | 4.31E-11 | up |
| Adamts14 | 2.273732 | 1.1850621 | 1.59E-12 | 4.38E-11 | up |
| Omd | 0.330685 | -1.59647 | 1.62E-12 | 4.46E-11 | down |
| Cited4 | 0.370043 | -1.434234 | 1.66E-12 | 4.57E-11 | down |
| Nwd1 | 0.351722 | -1.50749 | 1.98E-12 | 5.42E-11 | down |

| | | | | | |
|---|---|---|---|---|---|
| Clec4a | 8.259697 | 3.0460888 | 2.15E-12 | 5.83E-11 | up |
| Abcd2 | 0.379156 | -1.399137 | 2.17E-12 | 5.89E-11 | down |
| Efcc1 | 0.39889 | -1.325937 | 2.18E-12 | 5.89E-11 | down |
| Hmgcs1 | 2.019498 | 1.0139966 | 2.26E-12 | 6.11E-11 | up |
| Adamts4 | 9.648045 | 3.2702367 | 2.33E-12 | 6.28E-11 | up |
| Tnfaip6 | 5.576321 | 2.4793136 | 2.37E-12 | 6.39E-11 | up |
| Pgf | 2.718638 | 1.4428842 | 2.41E-12 | 6.48E-11 | up |
| Rcc1 | 2.02508 | 1.0179792 | 2.43E-12 | 6.55E-11 | up |
| Gmnn | 2.693438 | 1.4294491 | 2.56E-12 | 6.85E-11 | up |
| Tslp | 5.335981 | 2.4157534 | 2.69E-12 | 7.19E-11 | up |
| Mthfd1l | 2.671435 | 1.4176151 | 3.04E-12 | 8.09E-11 | up |
| Tnfrsf11a | 2.572536 | 1.3631912 | 3.28E-12 | 8.7E-11 | up |
| ENSRNOG00000062326 | 15579.78 | 13.927388 | 3.35E-12 | 8.89E-11 | up |
| Rtbdn | 0.361252 | -1.468922 | 3.45E-12 | 9.11E-11 | down |
| Steap2 | 2.226564 | 1.154819 | 3.83E-12 | 1.01E-10 | up |
| Hs3st6 | 0.384318 | -1.379626 | 3.86E-12 | 1.02E-10 | down |
| Marchf3 | 2.233992 | 1.1596239 | 4E-12 | 1.05E-10 | up |
| Zswim2 | 4.319789 | 2.1109609 | 4.13E-12 | 1.08E-10 | up |
| Aox3 | 0.234902 | -2.089871 | 4.17E-12 | 1.09E-10 | down |
| Penk | 7.289864 | 2.8658919 | 4.69E-12 | 1.22E-10 | up |
| Gper1 | 2.395734 | 1.2604675 | 4.72E-12 | 1.22E-10 | up |
| ENSRNOG00000065037 | 43.56727 | 5.4451728 | 4.95E-12 | 1.28E-10 | up |
| Adm | 2.201754 | 1.1386536 | 5E-12 | 1.29E-10 | up |
| Mmrn1 | 2.237488 | 1.1618802 | 5.06E-12 | 1.3E-10 | up |
| Clec4a1 | 2.872666 | 1.5223902 | 5.13E-12 | 1.32E-10 | up |
| Myh3 | 533.9806 | 9.0606434 | 5.36E-12 | 1.38E-10 | up |
| Slc41a3 | 2.762604 | 1.4660287 | 5.7E-12 | 1.46E-10 | up |
| F13a1 | 2.766871 | 1.4682553 | 5.88E-12 | 1.5E-10 | up |
| Slfn13 | 3.366469 | 1.7512363 | 6.05E-12 | 1.54E-10 | up |
| Prrx1 | 5.13236 | 2.3596225 | 6.52E-12 | 1.66E-10 | up |
| Gem | 3.082037 | 1.6238842 | 7.14E-12 | 1.81E-10 | up |
| Fignl1 | 6.701838 | 2.7445568 | 7.77E-12 | 1.96E-10 | up |
| Dctd | 2.308609 | 1.2070238 | 8.06E-12 | 2.03E-10 | up |
| Plppr2 | 2.290879 | 1.1959015 | 8.1E-12 | 2.03E-10 | up |
| Mnd1 | 5.336165 | 2.4158032 | 8.6E-12 | 2.16E-10 | up |
| Pof1b | 0.446299 | -1.163918 | 8.65E-12 | 2.17E-10 | down |
| Tspan11 | 2.56473 | 1.3588067 | 9.47E-12 | 2.36E-10 | up |
| Cdh24 | 5.471656 | 2.4519776 | 9.6E-12 | 2.39E-10 | up |
| Nes | 2.273444 | 1.1848797 | 1.11E-11 | 2.76E-10 | up |
| Sec14l5 | 0.386828 | -1.370235 | 1.14E-11 | 2.82E-10 | down |
| Pask | 2.408752 | 1.2682857 | 1.19E-11 | 2.93E-10 | up |

| | | | | | |
|---|---|---|---|---|---|
| Mturn | 0.440464 | -1.182903 | 1.22E-11 | 3E-10 | down |
| Kcna2 | 0.27931 | -1.840062 | 1.25E-11 | 3.07E-10 | down |
| Trim54 | 0.422458 | -1.243119 | 1.35E-11 | 3.32E-10 | down |
| Nol3 | 2.164903 | 1.1143026 | 1.43E-11 | 3.51E-10 | up |
| Dnah9 | 0.470401 | -1.088037 | 1.47E-11 | 3.6E-10 | down |
| Kcnj15 | 0.472907 | -1.08037 | 1.54E-11 | 3.76E-10 | down |
| Adora1 | 2.927193 | 1.5495181 | 1.56E-11 | 3.8E-10 | up |
| Scube3 | 2.760192 | 1.4647687 | 1.6E-11 | 3.89E-10 | up |
| Aspn | 2.654667 | 1.4085311 | 1.67E-11 | 4.06E-10 | up |
| Six1 | 0.405733 | -1.301397 | 1.85E-11 | 4.47E-10 | down |
| Esm1 | 0.22795 | -2.13321 | 1.99E-11 | 4.8E-10 | down |
| Myef2 | 2.14096 | 1.0982581 | 2.43E-11 | 5.78E-10 | up |
| H19 | 14.67025 | 3.874822 | 2.5E-11 | 5.94E-10 | up |
| Slc5a3 | 0.33132 | -1.593704 | 2.53E-11 | 5.99E-10 | down |
| Foxo6 | 0.487196 | -1.037426 | 2.56E-11 | 6.06E-10 | down |
| Tnfrsf10b | 4.162757 | 2.0575392 | 2.72E-11 | 6.4E-10 | up |
| Kcnn3 | 0.445504 | -1.166491 | 2.91E-11 | 6.83E-10 | down |
| Cenpk | 9.504501 | 3.2486109 | 3.05E-11 | 7.15E-10 | up |
| Caln1 | 0.255953 | -1.966048 | 3.16E-11 | 7.38E-10 | down |
| Apold1 | 4.193351 | 2.0681034 | 3.17E-11 | 7.41E-10 | up |
| Mrgprx3 | 0.353069 | -1.501977 | 3.26E-11 | 7.6E-10 | down |
| Sele | 5.252858 | 2.3931027 | 3.29E-11 | 7.67E-10 | up |
| Sgsm1 | 0.301127 | -1.731555 | 3.61E-11 | 8.38E-10 | down |
| Shc2 | 0.372765 | -1.423663 | 3.64E-11 | 8.45E-10 | down |
| Ube2q2l | 2.096811 | 1.068197 | 4.04E-11 | 9.36E-10 | up |
| Peg12 | 3.142001 | 1.6516838 | 4.08E-11 | 9.42E-10 | up |
| Arhgef25 | 2.246821 | 1.167885 | 4.7E-11 | 1.08E-09 | up |
| C3ar1 | 2.789561 | 1.4800383 | 4.75E-11 | 1.09E-09 | up |
| Bag2 | 2.077593 | 1.0549127 | 4.78E-11 | 1.1E-09 | up |
| Ubxn10 | 0.441838 | -1.178412 | 4.89E-11 | 1.12E-09 | down |
| Cpxm1 | 4.195592 | 2.0688745 | 5.02E-11 | 1.15E-09 | up |
| Ptprn | 8.188909 | 3.0336712 | 5.24E-11 | 1.2E-09 | up |
| Gli1 | 3.505801 | 1.8097442 | 5.6E-11 | 1.28E-09 | up |
| Klhdc7a | 0.368016 | -1.44216 | 6.26E-11 | 1.42E-09 | down |
| Prdm5 | 2.250553 | 1.1702794 | 6.54E-11 | 1.48E-09 | up |
| Gnmt | 0.291539 | -1.77824 | 6.74E-11 | 1.52E-09 | down |
| Vtcn1 | 0.280985 | -1.831432 | 6.75E-11 | 1.52E-09 | down |
| Col11a1 | 12.46859 | 3.6402265 | 6.9E-11 | 1.55E-09 | up |
| LOC681355 | 2.177408 | 1.1226118 | 7.19E-11 | 1.61E-09 | up |
| Cdo1 | 0.296797 | -1.752451 | 7.52E-11 | 1.68E-09 | down |
| Abcg1 | 0.492802 | -1.020919 | 8.11E-11 | 1.81E-09 | down |

| | | | | | |
|---|---|---|---|---|---|
| Dpysl5 | 3.94727 | 1.9808554 | 8.56E-11 | 1.91E-09 | up |
| Hcn4 | 0.411297 | -1.281747 | 9.06E-11 | 2.02E-09 | down |
| Cped1 | 0.480362 | -1.057806 | 9.51E-11 | 2.11E-09 | down |
| Thbs4 | 5.896401 | 2.5598347 | 1.02E-10 | 2.26E-09 | up |
| Mt2A | 6.700936 | 2.7443625 | 1.08E-10 | 2.38E-09 | up |
| Slc34a3 | 0.457741 | -1.127395 | 1.09E-10 | 2.4E-09 | down |
| Sox2 | 0.47571 | -1.071844 | 1.12E-10 | 2.45E-09 | down |
| ENSRNOG00000064568 | 2.806321 | 1.4886801 | 1.21E-10 | 2.64E-09 | up |
| Syt13 | 5.677627 | 2.5052879 | 1.28E-10 | 2.78E-09 | up |
| Spire2 | 0.499177 | -1.002375 | 1.32E-10 | 2.84E-09 | down |
| Flnc | 2.32212 | 1.2154427 | 1.34E-10 | 2.88E-09 | up |
| Cep295 | 2.681115 | 1.4228333 | 1.42E-10 | 3.07E-09 | up |
| Bmp15 | 0.328757 | -1.604908 | 1.43E-10 | 3.07E-09 | down |
| Zc2hc1c | 0.456135 | -1.132468 | 1.72E-10 | 3.68E-09 | down |
| Kcnk13 | 3.036692 | 1.6025007 | 1.72E-10 | 3.68E-09 | up |
| Lif | 4.227706 | 2.079875 | 1.78E-10 | 3.79E-09 | up |
| Pkhd1 | 4.00503 | 2.0018132 | 2.16E-10 | 4.6E-09 | up |
| Cdc25c | 2.974926 | 1.5728538 | 2.23E-10 | 4.72E-09 | up |
| Agr3 | 0.483146 | -1.049468 | 2.35E-10 | 4.96E-09 | down |
| Casq1 | 4.268337 | 2.0936742 | 2.56E-10 | 5.35E-09 | up |
| Brca2 | 2.157619 | 1.1094398 | 2.59E-10 | 5.42E-09 | up |
| Cdca7 | 3.049812 | 1.6087204 | 2.59E-10 | 5.42E-09 | up |
| LOC100911413 | 9.403257 | 3.2331605 | 2.69E-10 | 5.61E-09 | up |
| Slitrk6 | 0.308315 | -1.697523 | 3.08E-10 | 6.37E-09 | down |
| Frzb | 3.62918 | 1.8596436 | 3.19E-10 | 6.59E-09 | up |
| Cyp21a1 | 10.60687 | 3.4069269 | 3.49E-10 | 7.21E-09 | up |
| Ret | 0.494938 | -1.01468 | 3.56E-10 | 7.33E-09 | down |
| Esco2-ps2 | 6.179277 | 2.627438 | 3.61E-10 | 7.44E-09 | up |
| RGD1562029 | 0.405675 | -1.301602 | 3.77E-10 | 7.75E-09 | down |
| Socs3 | 2.878779 | 1.5254571 | 4.15E-10 | 8.5E-09 | up |
| Vnn3 | 0.462838 | -1.111422 | 4.28E-10 | 8.74E-09 | down |
| ENSRNOG00000064142 | 2.580995 | 1.3679273 | 4.47E-10 | 9.1E-09 | up |
| Palb2 | 2.615317 | 1.3869857 | 4.6E-10 | 9.34E-09 | up |
| Edil3 | 0.486105 | -1.04066 | 5.26E-10 | 1.06E-08 | down |
| Pclo | 0.420137 | -1.251068 | 5.69E-10 | 1.14E-08 | down |
| Efr3b | 2.311361 | 1.2087428 | 5.82E-10 | 1.16E-08 | up |
| Eme1 | 4.627884 | 2.2103528 | 6.53E-10 | 1.29E-08 | up |
| Tnfrsf22 | 4.335257 | 2.1161174 | 6.89E-10 | 1.36E-08 | up |
| Lvrn | 3.908777 | 1.9667171 | 7.92E-10 | 1.55E-08 | up |
| AABR07049134.4 | 3.308408 | 1.726137 | 8.15E-10 | 1.59E-08 | up |
| Ovol1 | 0.475999 | -1.070969 | 8.34E-10 | 1.63E-08 | down |

| | | | | | |
|---|---|---|---|---|---|
| Nme3 | 0.48032 | -1.057931 | 8.41E-10 | 1.64E-08 | down |
| Myrfl | 8.480601 | 3.0841666 | 9.34E-10 | 1.81E-08 | up |
| Stox1 | 0.478273 | -1.064093 | 9.57E-10 | 1.86E-08 | down |
| Adamtsl2 | 5.205664 | 2.3800822 | 1.02E-09 | 1.97E-08 | up |
| Pcsk1 | 0.433609 | -1.205533 | 1.07E-09 | 2.06E-08 | down |
| Prim2 | 2.862763 | 1.5174081 | 1.1E-09 | 2.12E-08 | up |
| Gas2l2 | 0.465932 | -1.101808 | 1.26E-09 | 2.4E-08 | down |
| Mdfi | 2.646074 | 1.4038536 | 1.33E-09 | 2.54E-08 | up |
| Ciart | 6.175231 | 2.6264932 | 1.33E-09 | 2.54E-08 | up |
| Ccbe1 | 2.420944 | 1.2755699 | 1.39E-09 | 2.64E-08 | up |
| Igsf10 | 2.437665 | 1.2854998 | 1.4E-09 | 2.66E-08 | up |
| AABR07032097.1 | 4.055525 | 2.0198886 | 1.43E-09 | 2.72E-08 | up |
| Tymp | 2.503037 | 1.3236797 | 1.52E-09 | 2.88E-08 | up |
| ENSRNOG00000065112 | 7.399804 | 2.887487 | 1.59E-09 | 2.99E-08 | up |
| Cfap44 | 0.481062 | -1.055704 | 1.71E-09 | 3.2E-08 | down |
| Ccsap | 3.162766 | 1.6611866 | 1.81E-09 | 3.38E-08 | up |
| Cntfr | 0.412015 | -1.279232 | 1.83E-09 | 3.42E-08 | down |
| Sp5 | 0.380966 | -1.392264 | 1.87E-09 | 3.49E-08 | down |
| Kif18a | 2.45947 | 1.2983477 | 2.12E-09 | 3.92E-08 | up |
| Mcm8 | 2.646615 | 1.4041481 | 2.41E-09 | 4.42E-08 | up |
| Zfp82 | 0.490006 | -1.02913 | 2.52E-09 | 4.61E-08 | down |
| Fgfbp1 | 0.398099 | -1.3288 | 2.52E-09 | 4.62E-08 | down |
| AABR07054614.1 | 2.13407 | 1.0936072 | 2.53E-09 | 4.62E-08 | up |
| Clec4a3 | 2.862467 | 1.517259 | 2.53E-09 | 4.63E-08 | up |
| Cidec | 0.284357 | -1.814224 | 2.66E-09 | 4.84E-08 | down |
| Tnfsf9 | 5.340229 | 2.4169016 | 3E-09 | 5.44E-08 | up |
| Ly6k | 0.000223 | -12.13235 | 3.25E-09 | 5.89E-08 | down |
| Cenpp | 4.828015 | 2.2714303 | 3.46E-09 | 6.26E-08 | up |
| Mgst1 | 0.487904 | -1.035332 | 3.59E-09 | 6.45E-08 | down |
| Artn | 0.420247 | -1.250691 | 3.69E-09 | 6.63E-08 | down |
| Rnf227 | 2.532664 | 1.3406559 | 3.72E-09 | 6.67E-08 | up |
| Tfpi2 | 5.073563 | 2.3429992 | 3.81E-09 | 6.81E-08 | up |
| Slc28a2 | 3.117937 | 1.6405917 | 3.93E-09 | 7.01E-08 | up |
| Nek2l1 (ENSRNOG00000012119) | 3.932662 | 1.9755062 | 4.12E-09 | 7.33E-08 | up |
| Polq | 5.742929 | 2.5217868 | 4.36E-09 | 7.73E-08 | up |
| Msc | 3.03448 | 1.6014493 | 4.38E-09 | 7.77E-08 | up |
| Sostdc1 | 0.264852 | -1.916744 | 4.79E-09 | 8.47E-08 | down |
| C1r | 2.079525 | 1.0562543 | 5.02E-09 | 8.84E-08 | up |
| Angpt4 | 4.461695 | 2.1575918 | 5.31E-09 | 9.32E-08 | up |
| Adgre1 | 2.61198 | 1.3851438 | 5.65E-09 | 9.88E-08 | up |
| Ankrd39 | 2.344437 | 1.2292418 | 6.22E-09 | 1.08E-07 | up |

| | | | | | |
|---|---|---|---|---|---|
| Rtn4rl2 | 4.713439 | 2.2367802 | 6.32E-09 | 1.1E-07 | up |
| Zfpm2 | 2.127455 | 1.0891287 | 6.48E-09 | 1.12E-07 | up |
| Nmnat2 | 0.467404 | -1.097258 | 6.52E-09 | 1.13E-07 | down |
| Pak1ip1 | 2.308025 | 1.2066591 | 6.75E-09 | 1.16E-07 | up |
| Clic6 | 0.376312 | -1.409998 | 7.36E-09 | 1.26E-07 | down |
| ENSRNOG00000064824 | 0.308287 | -1.697654 | 7.41E-09 | 1.27E-07 | down |
| AABR07019403.1 | 3.060972 | 1.6139896 | 7.72E-09 | 1.32E-07 | up |
| Prss46 | 2844.374 | 11.473895 | 7.74E-09 | 1.32E-07 | up |
| Cenpo | 2.418828 | 1.2743082 | 8.05E-09 | 1.37E-07 | up |
| Samd14 | 2.685513 | 1.4251977 | 8.28E-09 | 1.4E-07 | up |
| Ms4a4a | 2.233146 | 1.1590778 | 8.6E-09 | 1.45E-07 | up |
| Cenpm | 2.232836 | 1.158877 | 9.26E-09 | 1.56E-07 | up |
| Cyp7b1 | 2.479062 | 1.3097945 | 9.3E-09 | 1.57E-07 | up |
| Grhl3 | 0.274381 | -1.865747 | 9.77E-09 | 1.64E-07 | down |
| Cyp2e1 | 0.077448 | -3.690635 | 9.81E-09 | 1.65E-07 | down |
| Fancb | 3.224226 | 1.6889529 | 9.89E-09 | 1.66E-07 | up |
| Dmrtc2 | 0.430308 | -1.216557 | 1E-08 | 1.68E-07 | down |
| Abtb2 | 2.237785 | 1.1620712 | 1.03E-08 | 1.73E-07 | up |
| Tube1 | 2.631062 | 1.3956451 | 1.09E-08 | 1.82E-07 | up |
| Gpr19 | 2.264431 | 1.1791486 | 1.12E-08 | 1.86E-07 | up |
| Bpifb1 | 3.851134 | 1.9452831 | 1.13E-08 | 1.88E-07 | up |
| Pdgfrl | 2.482667 | 1.3118909 | 1.24E-08 | 2.06E-07 | up |
| Tcp11 | 2.365224 | 1.2419768 | 1.28E-08 | 2.12E-07 | up |
| Hpse2 | 2.667436 | 1.4154539 | 1.29E-08 | 2.13E-07 | up |
| Taf1d | 2.106547 | 1.0748804 | 1.33E-08 | 2.19E-07 | up |
| Cgas | 2.028462 | 1.0203865 | 1.39E-08 | 2.28E-07 | up |
| Ptx3 | 5.91207 | 2.5636633 | 1.49E-08 | 2.44E-07 | up |
| Col5a3 | 4.789534 | 2.2598852 | 1.56E-08 | 2.54E-07 | up |
| Samd11 | 2.524094 | 1.3357659 | 1.59E-08 | 2.58E-07 | up |
| Mafb | 2.801354 | 1.4861245 | 1.62E-08 | 2.62E-07 | up |
| Rnf152 | 2.893458 | 1.5327946 | 1.62E-08 | 2.62E-07 | up |
| Plppr4 | 2.713213 | 1.4400021 | 1.63E-08 | 2.64E-07 | up |
| Gjc1 | 2.064032 | 1.0454656 | 1.65E-08 | 2.66E-07 | up |
| Chia | 5.113442 | 2.3542947 | 1.77E-08 | 2.84E-07 | up |
| AABR07066753.1 | 2044.482 | 10.99752 | 1.8E-08 | 2.9E-07 | up |
| Klhl38 | 0.430497 | -1.215926 | 1.91E-08 | 3.06E-07 | down |
| Donson | 2.293109 | 1.1973049 | 1.99E-08 | 3.16E-07 | up |
| C6 | 17.52095 | 4.1310088 | 1.99E-08 | 3.17E-07 | up |
| Sectm1a | 7.517236 | 2.9102024 | 2.03E-08 | 3.22E-07 | up |
| Ccl20 | 0.173074 | -2.530536 | 2.12E-08 | 3.37E-07 | down |
| AABR07057438.1 | 6.153589 | 2.6214281 | 2.15E-08 | 3.4E-07 | up |

| | | | | | |
|---|---|---|---|---|---|
| ENSRNOG00000069461 | 0.428289 | -1.223344 | 2.15E-08 | 3.4E-07 | down |
| Ccdc18 | 2.474072 | 1.3068875 | 2.28E-08 | 3.59E-07 | up |
| Cd109 | 3.032341 | 1.6004319 | 2.33E-08 | 3.66E-07 | up |
| Ms4a7 | 9.254188 | 3.2101065 | 2.39E-08 | 3.75E-07 | up |
| Xrcc2 | 2.298014 | 1.2003877 | 2.44E-08 | 3.82E-07 | up |
| Sphk1 | 2.366954 | 1.2430317 | 2.47E-08 | 3.87E-07 | up |
| Pla2r1 | 2.418082 | 1.2738631 | 2.48E-08 | 3.88E-07 | up |
| Atp7b | 0.32638 | -1.615375 | 2.62E-08 | 4.08E-07 | down |
| Kirrel1 | 2.082285 | 1.0581678 | 2.64E-08 | 4.11E-07 | up |
| Sox21 | 0.254017 | -1.977006 | 2.72E-08 | 4.22E-07 | down |
| Rad18 | 2.152029 | 1.1056978 | 2.86E-08 | 4.43E-07 | up |
| Klra2 | 2.555314 | 1.3535007 | 2.94E-08 | 4.54E-07 | up |
| Cntln | 2.305106 | 1.2048329 | 3.02E-08 | 4.67E-07 | up |
| Selp | 3.376138 | 1.7553739 | 3.19E-08 | 4.91E-07 | up |
| Gstm1 | 0.498549 | -1.004194 | 3.24E-08 | 4.98E-07 | down |
| Sult1a1 | 0.442346 | -1.176752 | 3.27E-08 | 5.03E-07 | down |
| Strip2 | 2.297666 | 1.2001693 | 3.33E-08 | 5.12E-07 | up |
| Ms4a6a | 2.527764 | 1.3378619 | 3.56E-08 | 5.44E-07 | up |
| Enc1 | 2.015793 | 1.0113476 | 3.56E-08 | 5.44E-07 | up |
| Mfap2 | 2.696638 | 1.431162 | 3.68E-08 | 5.6E-07 | up |
| Gpr183 | 2.098451 | 1.069325 | 3.76E-08 | 5.71E-07 | up |
| Ddx11 | 2.545384 | 1.3478831 | 3.81E-08 | 5.77E-07 | up |
| Serpinb10 | 0.314239 | -1.670067 | 3.88E-08 | 5.87E-07 | down |
| Reg3b | 34221.17 | 15.062602 | 3.9E-08 | 5.89E-07 | up |
| Abhd14a | 0.446879 | -1.162043 | 4.05E-08 | 6.11E-07 | down |
| Sh3pxd2b | 2.01976 | 1.0141841 | 4.26E-08 | 6.42E-07 | up |
| Cxcr1 | 0.476149 | -1.070515 | 4.27E-08 | 6.42E-07 | down |
| Erich3 | 0.452182 | -1.145026 | 4.43E-08 | 6.63E-07 | down |
| Fbln2 | 2.051796 | 1.0368872 | 4.94E-08 | 7.34E-07 | up |
| Bank1 | 0.438055 | -1.190816 | 5.23E-08 | 7.75E-07 | down |
| Cdh26 | 0.438292 | -1.190036 | 5.25E-08 | 7.77E-07 | down |
| Gpihbp1 | 0.365272 | -1.452959 | 5.87E-08 | 8.66E-07 | down |
| Gins2 | 3.148581 | 1.6547017 | 5.87E-08 | 8.66E-07 | up |
| Spdl1 | 2.282152 | 1.1903949 | 6.54E-08 | 9.57E-07 | up |
| Gdf6 | 58.7067 | 5.8754533 | 6.64E-08 | 9.7E-07 | up |
| Lama1 | 42.06838 | 5.3946644 | 7.35E-08 | 1.07E-06 | up |
| Rgs16 | 5.525181 | 2.4660218 | 7.66E-08 | 1.11E-06 | up |
| Mmp19 | 2.010259 | 1.0073812 | 7.88E-08 | 1.14E-06 | up |
| Plcb1 | 2.075308 | 1.0533252 | 8.22E-08 | 1.18E-06 | up |
| Adamts19 | 3.309708 | 1.7267037 | 9.08E-08 | 1.29E-06 | up |
| Gpr39 | 2.615738 | 1.3872179 | 9.14E-08 | 1.3E-06 | up |

| | | | | | |
|---|---|---|---|---|---|
| Magix | 0.452114 | -1.145243 | 9.33E-08 | 1.33E-06 | down |
| Mms22l | 3.926414 | 1.9732124 | 9.34E-08 | 1.33E-06 | up |
| Bmp2 | 2.264159 | 1.178975 | 9.56E-08 | 1.36E-06 | up |
| Trhr | 0.001815 | -9.106008 | 9.74E-08 | 1.38E-06 | down |
| AABR07027854.1 | 2.108158 | 1.0759828 | 9.78E-08 | 1.38E-06 | up |
| Kctd8 | 0.376863 | -1.407887 | 9.88E-08 | 1.4E-06 | down |
| Lrriq4 | 0.480921 | -1.056129 | 1.01E-07 | 1.43E-06 | down |
| Cfi | 2.477391 | 1.3088219 | 1.04E-07 | 1.46E-06 | up |
| Orm1 | 6.699815 | 2.7441213 | 1.08E-07 | 1.51E-06 | up |
| Csprs | 0.421019 | -1.248042 | 1.09E-07 | 1.54E-06 | down |
| Lgr6 | 2.265411 | 1.1797731 | 1.2E-07 | 1.67E-06 | up |
| Tmem119 | 2.018884 | 1.0135582 | 1.26E-07 | 1.74E-06 | up |
| Fabp3 | 0.305569 | -1.710429 | 1.28E-07 | 1.78E-06 | down |
| Ska3 | 2.019841 | 1.0142417 | 1.35E-07 | 1.86E-06 | up |
| Bcl3 | 2.180462 | 1.1246338 | 1.39E-07 | 1.91E-06 | up |
| Scn3a | 5.757287 | 2.5253892 | 1.41E-07 | 1.94E-06 | up |
| Akr1c19 | 0.432453 | -1.209384 | 1.46E-07 | 2E-06 | down |
| Nr4a3 | 4.276121 | 2.0963026 | 1.6E-07 | 2.17E-06 | up |
| Fn3k | 0.255016 | -1.97134 | 1.65E-07 | 2.23E-06 | down |
| Necab2 | 3.879319 | 1.9558034 | 1.82E-07 | 2.45E-06 | up |
| Pif1 | 9.476612 | 3.2443713 | 1.91E-07 | 2.56E-06 | up |
| Spink8 | 3.004469 | 1.5871098 | 1.97E-07 | 2.63E-06 | up |
| Hmgb2-ps6 | 3.66067 | 1.8721076 | 2.47E-07 | 3.24E-06 | up |
| P2ry6 | 2.24345 | 1.165719 | 2.67E-07 | 3.48E-06 | up |
| Lefty2 | 1216.946 | 10.249049 | 2.7E-07 | 3.51E-06 | up |
| Scara5 | 2.667576 | 1.4155293 | 2.73E-07 | 3.55E-06 | up |
| Itgb3bp | 2.048272 | 1.0344076 | 2.74E-07 | 3.56E-06 | up |
| Rad51b | 2.704682 | 1.4354589 | 2.75E-07 | 3.57E-06 | up |
| Uchl3 | 2.373428 | 1.2469722 | 2.93E-07 | 3.78E-06 | up |
| Mapk11 | 3.246572 | 1.6989174 | 3.16E-07 | 4.07E-06 | up |
| Ccr3 | 8.033283 | 3.0059897 | 3.17E-07 | 4.08E-06 | up |
| ENSRNOG00000068860 | 0.442009 | -1.177854 | 3.39E-07 | 4.33E-06 | down |
| Proc | 0.328943 | -1.604092 | 3.7E-07 | 4.71E-06 | down |
| Bmal1 | 0.476751 | -1.068691 | 3.87E-07 | 4.92E-06 | down |
| Sgcd | 2.674539 | 1.4192901 | 3.88E-07 | 4.93E-06 | up |
| Ago4 | 0.495003 | -1.014491 | 4.19E-07 | 5.29E-06 | down |
| Gpc3 | 2.061099 | 1.0434136 | 4.24E-07 | 5.35E-06 | up |
| Otx1 | 0.369096 | -1.437931 | 5.07E-07 | 6.33E-06 | down |
| Ggn | 2.207792 | 1.1426041 | 5.34E-07 | 6.62E-06 | up |
| Pnma8c | 15.18867 | 3.924924 | 5.53E-07 | 6.84E-06 | up |
| Msx3 | 0.450298 | -1.151047 | 6.57E-07 | 8.06E-06 | down |

| | | | | | |
|---|---|---|---|---|---|
| Trim59 | 2.451228 | 1.2935049 | 6.7E-07 | 8.2E-06 | up |
| Foxs1 | 2.094958 | 1.0669212 | 6.79E-07 | 8.3E-06 | up |
| Ephb2 | 2.135587 | 1.0946324 | 6.98E-07 | 8.52E-06 | up |
| P2rx1 | 0.482697 | -1.050809 | 7.18E-07 | 8.75E-06 | down |
| Npas2 | 0.492052 | -1.023119 | 8.02E-07 | 9.7E-06 | down |
| Chrm3 | 0.35358 | -1.499891 | 8.47E-07 | 1.02E-05 | down |
| Ndufa10l1 | 0.3997 | -1.323011 | 8.77E-07 | 1.06E-05 | down |
| Mfap5 | 2.096649 | 1.0680852 | 8.98E-07 | 1.08E-05 | up |
| Cdsn | 2.240182 | 1.1636162 | 9.73E-07 | 1.16E-05 | up |
| Steap1 | 3.314464 | 1.7287758 | 1.02E-06 | 1.22E-05 | up |
| ENSRNOG00000067913 | 0.456001 | -1.13289 | 1.09E-06 | 1.28E-05 | down |
| Ppef1 | 3.959501 | 1.9853187 | 1.11E-06 | 1.31E-05 | up |
| Plek2 | 2.188486 | 1.1299328 | 1.13E-06 | 1.33E-05 | up |
| Arsj | 2.112309 | 1.0788207 | 1.16E-06 | 1.36E-05 | up |
| Fam114a1l1 | 2.482334 | 1.3116974 | 1.19E-06 | 1.39E-05 | up |
| Rd3 | 8.852567 | 3.1460958 | 1.2E-06 | 1.41E-05 | up |
| Sptbn4 | 2.662851 | 1.412972 | 1.25E-06 | 1.46E-05 | up |
| Fbxo43 | 3.079224 | 1.622567 | 1.32E-06 | 1.53E-05 | up |
| Ccl7 | 11.8093 | 3.5618522 | 1.36E-06 | 1.56E-05 | up |
| Lhx6 | 2.731579 | 1.449735 | 1.37E-06 | 1.57E-05 | up |
| Reln | 2.830491 | 1.5010523 | 1.38E-06 | 1.59E-05 | up |
| ENSRNOG00000068603 | 3.308032 | 1.7259731 | 1.4E-06 | 1.6E-05 | up |
| Slc6a7 | 0.184096 | -2.441472 | 1.4E-06 | 1.6E-05 | down |
| Sfrp1 | 2.289205 | 1.1948467 | 1.49E-06 | 1.7E-05 | up |
| Zbtb8b | 0.449122 | -1.154822 | 1.49E-06 | 1.7E-05 | down |
| AABR07054096.1 | 0.445918 | -1.165149 | 1.51E-06 | 1.72E-05 | down |
| Cep152 | 2.244508 | 1.166399 | 1.51E-06 | 1.72E-05 | up |
| Nnmt | 2.031363 | 1.0224478 | 1.51E-06 | 1.72E-05 | up |
| Bdkrb1 | 12.87222 | 3.6861889 | 1.57E-06 | 1.77E-05 | up |
| C1s | 2.229875 | 1.1569628 | 1.67E-06 | 1.88E-05 | up |
| Adora2b | 2.762591 | 1.4660218 | 1.69E-06 | 1.91E-05 | up |
| Nacad | 2.296089 | 1.1991786 | 1.7E-06 | 1.91E-05 | up |
| Lrr1 | 8.276758 | 3.0490657 | 1.7E-06 | 1.91E-05 | up |
| Trh | 102.2459 | 6.6758992 | 1.78E-06 | 2E-05 | up |
| Klhl30 | 2.279436 | 1.1886772 | 1.83E-06 | 2.05E-05 | up |
| Trmt9b | 0.400347 | -1.320677 | 1.85E-06 | 2.07E-05 | down |
| Cyp2b2 | 0.171748 | -2.541631 | 1.86E-06 | 2.09E-05 | down |
| Ereg | 6.139322 | 2.6180794 | 1.92E-06 | 2.15E-05 | up |
| Tcp11x2 | 3.691704 | 1.8842868 | 1.95E-06 | 2.17E-05 | up |
| Adcy1 | 0.385172 | -1.376426 | 1.97E-06 | 2.19E-05 | down |
| Hspb7 | 2.638966 | 1.3999728 | 1.99E-06 | 2.22E-05 | up |

| | | | | | |
|---|---|---|---|---|---|
| Hmox1 | 2.407221 | 1.2673686 | 2E-06 | 2.23E-05 | up |
| Ttc24 | 35.12717 | 5.1345155 | 2.06E-06 | 2.29E-05 | up |
| Myo7a | 2.056468 | 1.0401684 | 2.08E-06 | 2.31E-05 | up |
| Lrfn3 | 2.145351 | 1.1012136 | 2.15E-06 | 2.37E-05 | up |
| LOC100362216 | 0.486779 | -1.038661 | 2.24E-06 | 2.47E-05 | down |
| Ccl12 | 15.12656 | 3.919012 | 2.31E-06 | 2.54E-05 | up |
| Sec16b | 2.212196 | 1.1454795 | 2.35E-06 | 2.58E-05 | up |
| Ank1 | 0.467602 | -1.096647 | 2.39E-06 | 2.62E-05 | down |
| Susd3 | 2.162448 | 1.1126653 | 2.48E-06 | 2.71E-05 | up |
| Disp2 | 2.402705 | 1.2646594 | 2.57E-06 | 2.79E-05 | up |
| C4bpb | 2.36737 | 1.2432853 | 2.76E-06 | 2.98E-05 | up |
| Ccr5 | 2.095751 | 1.0674674 | 2.84E-06 | 3.05E-05 | up |
| Ihh | 0.309734 | -1.690897 | 2.89E-06 | 3.1E-05 | down |
| Matk | 3.149331 | 1.6550454 | 2.92E-06 | 3.12E-05 | up |
| Cfc1 | 0.291336 | -1.779246 | 3.03E-06 | 3.23E-05 | down |
| Tex52 | 5.807311 | 2.5378703 | 3.05E-06 | 3.26E-05 | up |
| RGD1311946 | 2.099015 | 1.0697124 | 3.16E-06 | 3.36E-05 | up |
| Gbx2 | 1208.996 | 10.239594 | 3.18E-06 | 3.38E-05 | up |
| ENSRNOG00000070452 | 9.272738 | 3.2129954 | 3.26E-06 | 3.45E-05 | up |
| Lrp8 | 2.230144 | 1.1571366 | 3.34E-06 | 3.53E-05 | up |
| Fbn2 | 2.729931 | 1.4488646 | 3.7E-06 | 3.88E-05 | up |
| Ppara | 0.434259 | -1.203374 | 3.73E-06 | 3.9E-05 | down |
| Tmsb15b2 | 2.192922 | 1.1328546 | 3.76E-06 | 3.92E-05 | up |
| ENSRNOG00000069208 | 2.487833 | 1.3148896 | 3.85E-06 | 4.01E-05 | up |
| Ccn4 | 32.3241 | 5.0145384 | 3.95E-06 | 4.11E-05 | up |
| Tnfrsf19 | 0.37781 | -1.404268 | 3.96E-06 | 4.12E-05 | down |
| Fcgr2b | 2.336966 | 1.2246368 | 4.03E-06 | 4.18E-05 | up |
| Np4 | 6416.528 | 12.647577 | 4.1E-06 | 4.25E-05 | up |
| C4b | 2.898664 | 1.5353879 | 4.18E-06 | 4.32E-05 | up |
| Mcoln2 | 0.490556 | -1.02751 | 4.2E-06 | 4.34E-05 | down |
| Calml3 | 0.415737 | -1.266257 | 4.2E-06 | 4.34E-05 | down |
| Zfp853 | 2.148126 | 1.1030788 | 4.23E-06 | 4.37E-05 | up |
| Itln1 | 22.4632 | 4.4894914 | 4.41E-06 | 4.53E-05 | up |
| Txnrd3 | 2.155414 | 1.1079647 | 4.44E-06 | 4.55E-05 | up |
| Vsig1 | 0.305244 | -1.711967 | 4.44E-06 | 4.55E-05 | down |
| LOC120098113 | 2.640801 | 1.4009756 | 4.5E-06 | 4.6E-05 | up |
| Gldn | 24.14526 | 4.593668 | 4.61E-06 | 4.69E-05 | up |
| ENSRNOG00000066291 | 0.000259 | -11.91454 | 4.71E-06 | 4.79E-05 | down |
| Cdhr3 | 0.470315 | -1.088302 | 4.73E-06 | 4.81E-05 | down |
| Gxylt2 | 2.22853 | 1.1560921 | 4.9E-06 | 4.96E-05 | up |
| Trpc6 | 2.502072 | 1.3231234 | 5.34E-06 | 5.38E-05 | up |

| | | | | | |
|---|---|---|---|---|---|
| Snora64 | 3.160682 | 1.660236 | 5.39E-06 | 5.42E-05 | up |
| Pcdhb10 | 2.115068 | 1.0807042 | 5.9E-06 | 5.9E-05 | up |
| Il22ra2 | 0.370659 | -1.431835 | 6.1E-06 | 6.08E-05 | down |
| P2ry12 | 2.073466 | 1.0520446 | 6.29E-06 | 6.25E-05 | up |
| Gria3 | 2.174574 | 1.1207325 | 6.38E-06 | 6.33E-05 | up |
| Dgki | 2.051065 | 1.0363729 | 6.52E-06 | 6.46E-05 | up |
| RGD1564696 | 0.223916 | -2.15897 | 6.54E-06 | 6.48E-05 | down |
| Sfn | 2.692797 | 1.4291052 | 6.71E-06 | 6.64E-05 | up |
| Clec2d | 7.397834 | 2.887103 | 7.16E-06 | 7.03E-05 | up |
| Timp4 | 0.466687 | -1.099473 | 7.27E-06 | 7.13E-05 | down |
| ENSRNOG00000068035 | 2.27732 | 1.1873368 | 7.4E-06 | 7.24E-05 | up |
| Suv39h2 | 2.26155 | 1.1773119 | 8.51E-06 | 8.26E-05 | up |
| Cnr1 | 0.470671 | -1.087208 | 8.78E-06 | 8.5E-05 | down |
| Adamts3 | 3.350825 | 1.7445165 | 9.12E-06 | 8.79E-05 | up |
| Mylk2 | 2.74195 | 1.4552024 | 9.2E-06 | 8.87E-05 | up |
| Olr59 | 3.563593 | 1.8333327 | 9.21E-06 | 8.88E-05 | up |
| AC118772.2 | 2.798806 | 1.4848113 | 9.37E-06 | 9.02E-05 | up |
| Fgf9 | 0.466427 | -1.100277 | 9.51E-06 | 9.14E-05 | down |
| Cyp2a3 | 0.248772 | -2.007105 | 9.74E-06 | 9.34E-05 | down |
| Tesmin | 4.211349 | 2.0742823 | 1E-05 | 9.59E-05 | up |
| Inmt | 0.227227 | -2.137796 | 1.04E-05 | 9.89E-05 | down |
| Etv4 | 2.01386 | 1.0099632 | 1.07E-05 | 0.000102 | up |
| Serpina3a | 6.190756 | 2.6301156 | 1.09E-05 | 0.000103 | up |
| Rgs20 | 3.098993 | 1.6317994 | 1.11E-05 | 0.000105 | up |
| Nkain1 | 4.628517 | 2.2105501 | 1.11E-05 | 0.000105 | up |
| Gpr63 | 2.217985 | 1.1492494 | 1.12E-05 | 0.000106 | up |
| Ablim2 | 0.41926 | -1.254082 | 1.15E-05 | 0.000109 | down |
| Ptafr | 2.402289 | 1.2644096 | 1.24E-05 | 0.000117 | up |
| Slamf9 | 3.069608 | 1.6180544 | 1.28E-05 | 0.00012 | up |
| Retnlg | 11.73929 | 3.5532734 | 1.28E-05 | 0.00012 | up |
| Ghrhr | 25.17187 | 4.6537407 | 1.31E-05 | 0.000122 | up |
| Uchl1 | 2.473611 | 1.3066184 | 1.35E-05 | 0.000125 | up |
| Slc38a4 | 2.735882 | 1.4520061 | 1.36E-05 | 0.000126 | up |
| Nipal4 | 2.928245 | 1.5500364 | 1.36E-05 | 0.000127 | up |
| Rspo2 | 0.204127 | -2.292462 | 1.37E-05 | 0.000127 | down |
| Matn3 | 11.07261 | 3.468923 | 1.38E-05 | 0.000128 | up |
| Drp2 | 2.254366 | 1.1727217 | 1.38E-05 | 0.000128 | up |
| Steap3 | 2.252098 | 1.1712694 | 1.45E-05 | 0.000134 | up |
| Rnft2 | 2.42674 | 1.2790195 | 1.46E-05 | 0.000134 | up |
| Eml5 | 0.470504 | -1.087721 | 1.47E-05 | 0.000136 | down |
| ENSRNOG00000068478 | 0.494952 | -1.014638 | 1.48E-05 | 0.000136 | down |

| | | | | | |
|---|---|---|---|---|---|
| Snord45a | 2.579323 | 1.3669923 | 1.53E-05 | 0.00014 | up |
| Mstn | 1005.48 | 9.9736687 | 1.54E-05 | 0.000141 | up |
| Kcnmb1 | 2.799834 | 1.4853413 | 1.57E-05 | 0.000143 | up |
| Tnni3 | 0.36884 | -1.438934 | 1.58E-05 | 0.000144 | down |
| Scube2 | 0.49875 | -1.003612 | 1.61E-05 | 0.000147 | down |
| Nr4a1 | 4.022983 | 2.0082656 | 1.62E-05 | 0.000148 | up |
| Mmp12 | 8.309857 | 3.0548237 | 1.62E-05 | 0.000148 | up |
| Dio3 | 13.64546 | 3.7703489 | 1.69E-05 | 0.000154 | up |
| Slc6a17 | 2.680809 | 1.4226684 | 1.73E-05 | 0.000157 | up |
| Cnih2 | 2.044039 | 1.0314224 | 1.83E-05 | 0.000165 | up |
| Lctl | 266.1237 | 8.0559532 | 1.85E-05 | 0.000167 | up |
| ENSRNOG00000062977 | 0.127148 | -2.975423 | 1.87E-05 | 0.000168 | down |
| Cilp | 2.041248 | 1.0294513 | 1.88E-05 | 0.000169 | up |
| Mmp7 | 92.48002 | 6.5310699 | 1.89E-05 | 0.000169 | up |
| Kcnj13 | 0.492945 | -1.0205 | 1.94E-05 | 0.000174 | down |
| H2az1-ps1 | 2.077227 | 1.0546589 | 1.95E-05 | 0.000174 | up |
| Gpr176 | 2.278841 | 1.1883002 | 1.98E-05 | 0.000177 | up |
| Rpl8 | 0.421824 | -1.245288 | 2E-05 | 0.000178 | down |
| Foxc1 | 0.29754 | -1.748847 | 2.01E-05 | 0.000179 | down |
| Muc5ac | 14.47345 | 3.8553374 | 2.07E-05 | 0.000184 | up |
| Nlrp10 | 3.177157 | 1.6677365 | 2.1E-05 | 0.000186 | up |
| Chrm1 | 7.156276 | 2.839209 | 2.23E-05 | 0.000196 | up |
| Ephb1 | 3.239935 | 1.6959648 | 2.24E-05 | 0.000197 | up |
| Matcap1 | 2.149008 | 1.1036712 | 2.26E-05 | 0.000199 | up |
| Slc29a4 | 3.769814 | 1.9144935 | 2.27E-05 | 0.0002 | up |
| Kcnd1 | 2.186768 | 1.1287998 | 2.47E-05 | 0.000215 | up |
| Tmem235 | 1832.436 | 10.839547 | 2.54E-05 | 0.000221 | up |
| ENSRNOG00000065520 | 14.81598 | 3.8890825 | 2.68E-05 | 0.000232 | up |
| Slc9b2 | 5.167012 | 2.3693301 | 2.79E-05 | 0.000241 | up |
| Rtkn2 | 8729.053 | 13.091609 | 2.84E-05 | 0.000244 | up |
| Kcna6 | 0.392591 | -1.3489 | 2.86E-05 | 0.000245 | down |
| Htr2b | 2.534066 | 1.3414543 | 2.87E-05 | 0.000246 | up |
| Stmn4 | 3.38948 | 1.761064 | 2.9E-05 | 0.000248 | up |
| Wasf1 | 3.595233 | 1.8460853 | 3.04E-05 | 0.000259 | up |
| Dusp9 | 3.081747 | 1.6237485 | 3.05E-05 | 0.00026 | up |
| C14h5orf52 | 3.221444 | 1.6877074 | 3.16E-05 | 0.000268 | up |
| Tnn | 8.169215 | 3.0301975 | 3.24E-05 | 0.000275 | up |
| Plin5 | 0.48861 | -1.033245 | 3.28E-05 | 0.000277 | down |
| Cd200r1l | 0.460952 | -1.117311 | 3.39E-05 | 0.000285 | down |
| Pcdhb7 | 2.902977 | 1.5375332 | 3.42E-05 | 0.000288 | up |
| Areg | 5.225106 | 2.3854604 | 3.67E-05 | 0.000306 | up |

| | | | | | |
|---|---|---|---|---|---|
| Cxcl10 | 2.219489 | 1.1502274 | 3.71E-05 | 0.000309 | up |
| Rspo4 | 29.093 | 4.8626003 | 3.78E-05 | 0.000314 | up |
| Scamp5 | 2.112143 | 1.0787075 | 3.88E-05 | 0.000321 | up |
| Sirpb3 | 0.498039 | -1.00567 | 3.89E-05 | 0.000322 | down |
| ENSRNOG00000063899 | 0.437237 | -1.193514 | 3.9E-05 | 0.000323 | down |
| Kcnj3 | 0.37286 | -1.423296 | 3.92E-05 | 0.000324 | down |
| Pcdhb6l | 2.071742 | 1.0508444 | 4.14E-05 | 0.00034 | up |
| Saal1 | 2.073232 | 1.0518816 | 4.16E-05 | 0.000341 | up |
| Myoz1 | 0.466907 | -1.098792 | 4.17E-05 | 0.000341 | down |
| Shisal1 | 3.76979 | 1.914484 | 4.24E-05 | 0.000346 | up |
| Sct | 3.924495 | 1.9725071 | 4.44E-05 | 0.000361 | up |
| Ngb | 0.325277 | -1.620259 | 4.57E-05 | 0.00037 | down |
| ENSRNOG00000066172 | 23.09296 | 4.5293814 | 4.59E-05 | 0.000371 | up |
| Pdzrn4 | 3.119271 | 1.641209 | 4.59E-05 | 0.000371 | up |
| RGD1561694 | 3.914271 | 1.9687437 | 4.6E-05 | 0.000372 | up |
| LOC685203 | 0.412223 | -1.278503 | 4.89E-05 | 0.000393 | down |
| ENSRNOG00000063341 | 0.000137 | -12.83027 | 5.12E-05 | 0.000411 | down |
| Htr7 | 5.01243 | 2.3255101 | 5.24E-05 | 0.00042 | up |
| Tmem86a | 2.346125 | 1.2302798 | 5.35E-05 | 0.000428 | up |
| Mcidas | 3.26303 | 1.7062122 | 5.48E-05 | 0.000436 | up |
| Plk5 | 2.430095 | 1.2810126 | 5.52E-05 | 0.000439 | up |
| Scn2a | 3.409738 | 1.7696609 | 5.62E-05 | 0.000446 | up |
| Sel1l3 | 2.161171 | 1.1118132 | 5.71E-05 | 0.000453 | up |
| Tmem200c | 2.265858 | 1.1800577 | 5.72E-05 | 0.000453 | up |
| ENSRNOG00000071049 | 0.071263 | -3.810695 | 5.78E-05 | 0.000457 | down |
| Oas2 | 0.450294 | -1.151061 | 5.86E-05 | 0.000463 | down |
| Itgam | 3.522688 | 1.8166765 | 5.87E-05 | 0.000463 | up |
| Prox1 | 4.055064 | 2.0197247 | 5.97E-05 | 0.00047 | up |
| Ppef2 | 3.040675 | 1.6043915 | 5.98E-05 | 0.000471 | up |
| Chst11 | 2.039878 | 1.0284827 | 6.13E-05 | 0.000482 | up |
| ENSRNOG00000070476 | 0.407147 | -1.296378 | 6.16E-05 | 0.000484 | down |
| LOC120094021 | 2.17811 | 1.1230769 | 6.53E-05 | 0.00051 | up |
| Lmo3 | 0.462995 | -1.110932 | 6.75E-05 | 0.000525 | down |
| Atf3 | 3.593375 | 1.8453396 | 6.92E-05 | 0.000537 | up |
| Pianp | 2.949349 | 1.5603966 | 7.05E-05 | 0.000546 | up |
| Bmp7 | 0.492517 | -1.021754 | 7.22E-05 | 0.000558 | down |
| Dscaml1 | 0.498752 | -1.003607 | 7.32E-05 | 0.000565 | down |
| Pitx1 | 0.492666 | -1.021318 | 7.36E-05 | 0.000567 | down |
| Ptprv | 28.31473 | 4.823481 | 7.38E-05 | 0.000569 | up |
| Cyp2j10 | 2.218101 | 1.149325 | 7.4E-05 | 0.000569 | up |
| Ddc | 2.229319 | 1.1566028 | 7.42E-05 | 0.00057 | up |

| | | | | | |
|---|---|---|---|---|---|
| LOC102551114 | 2.121807 | 1.0852937 | 7.48E-05 | 0.000574 | up |
| ENSRNOG00000069366 | 2.200356 | 1.1377367 | 7.51E-05 | 0.000576 | up |
| ENSRNOG00000064296 | 3.536831 | 1.8224571 | 8.01E-05 | 0.000611 | up |
| Cyp2c13 | 1183.928 | 10.209365 | 8.62E-05 | 0.000655 | up |
| Aldh1a7 | 0.360711 | -1.471086 | 8.76E-05 | 0.000664 | down |
| Cd300a | 2.239497 | 1.1631748 | 8.79E-05 | 0.000665 | up |
| Cyp2c12 | 0.000123 | -12.98787 | 8.98E-05 | 0.000679 | down |
| LOC298116 | 75.83395 | 6.244772 | 9.16E-05 | 0.000691 | up |
| Fat3 | 3.132725 | 1.6474183 | 9.41E-05 | 0.000707 | up |
| Hs6st2 | 2.280417 | 1.1892979 | 9.5E-05 | 0.000713 | up |
| ENSRNOG00000070458 | 0.350555 | -1.512288 | 9.5E-05 | 0.000713 | down |
| Fbln7 | 2.643393 | 1.402391 | 9.68E-05 | 0.000724 | up |
| ENSRNOG00000070389 | 2.237563 | 1.1619281 | 9.74E-05 | 0.000728 | up |
| Mrps17 | 3.708883 | 1.8909849 | 9.77E-05 | 0.000729 | up |
| Scn1a | 13.09071 | 3.7104712 | 0.000104 | 0.00077 | up |
| Ccl17 | 3.726113 | 1.8976715 | 0.000107 | 0.000793 | up |
| Chrna10 | 2.61371 | 1.3860991 | 0.000108 | 0.000795 | up |
| Ssmem1 | 0.444691 | -1.169123 | 0.000108 | 0.000795 | down |
| Prnd | 15.74813 | 3.9771089 | 0.000108 | 0.000797 | up |
| Ism2 | 2.055608 | 1.0395649 | 0.000108 | 0.000799 | up |
| Adgrd1 | 2.029183 | 1.0208989 | 0.000112 | 0.000826 | up |
| Rrad | 2.048208 | 1.0343623 | 0.000113 | 0.000828 | up |
| Krt23 | 0.496432 | -1.010331 | 0.000115 | 0.000842 | down |
| C2cd4b | 2.104682 | 1.0736025 | 0.000117 | 0.000856 | up |
| Fosb | 6.423792 | 2.6834252 | 0.000121 | 0.00088 | up |
| Pgam2 | 0.415552 | -1.266898 | 0.000124 | 0.000897 | down |
| Kcnj10 | 0.392015 | -1.351019 | 0.000125 | 0.000901 | down |
| Arhgap33 | 2.140534 | 1.0979708 | 0.000125 | 0.000902 | up |
| Adgrg3 | 2.485114 | 1.3133122 | 0.000126 | 0.00091 | up |
| ENSRNOG00000066421 | 9.439979 | 3.2387836 | 0.000126 | 0.00091 | up |
| Syt15 | 0.497576 | -1.007012 | 0.000127 | 0.000913 | down |
| Ptprh | 0.387356 | -1.368269 | 0.00013 | 0.000935 | down |
| Sfrp5 | 0.156809 | -2.672923 | 0.000131 | 0.000939 | down |
| ENSRNOG00000068365 | 0.487343 | -1.03699 | 0.000131 | 0.000944 | down |
| LOC100360095 | 2693.472 | 11.395251 | 0.000133 | 0.000955 | up |
| ENSRNOG00000071081 | 4.03E-05 | -14.59919 | 0.000143 | 0.001017 | down |
| Vash2 | 3.341125 | 1.7403338 | 0.000151 | 0.001071 | up |
| A2m | 44.43753 | 5.4737067 | 0.000159 | 0.001119 | up |
| LOC102553350 | 19.30162 | 4.2706504 | 0.000163 | 0.001138 | up |
| Cyp2b12 | 0.00148 | -9.400508 | 0.000167 | 0.001163 | down |
| Rps7-ps23 | 0.491207 | -1.025597 | 0.000168 | 0.00117 | down |

| | | | | | |
|---|---|---|---|---|---|
| Kcnma1 | 0.486661 | -1.03901 | 0.000181 | 0.001252 | down |
| Notum | 0.46688 | -1.098876 | 0.000183 | 0.00126 | down |
| Ctsk | 4.010602 | 2.0038189 | 0.000183 | 0.001265 | up |
| LOC120096111 | 2.261578 | 1.1773297 | 0.000184 | 0.001271 | up |
| Ly49s6 | 0.368329 | -1.440932 | 0.000186 | 0.00128 | down |
| ENSRNOG00000063847 | 3.345176 | 1.7420819 | 0.000191 | 0.001309 | up |
| Thrsp | 0.036123 | -4.790919 | 0.000191 | 0.001309 | down |
| Shank2 | 0.443497 | -1.173005 | 0.000197 | 0.001346 | down |
| Slc6a11 | 0.127595 | -2.970358 | 0.000199 | 0.001354 | down |
| Pi15 | 2.542651 | 1.3463332 | 0.000203 | 0.001383 | up |
| Pirt | 2.040754 | 1.029102 | 0.000206 | 0.001397 | up |
| Fcamr | 0.277763 | -1.848076 | 0.000214 | 0.001446 | down |
| Myh4 | 64.98181 | 6.021964 | 0.000215 | 0.001455 | up |
| Lypd1 | 2.409206 | 1.2685575 | 0.000235 | 0.001569 | up |
| Tnfsf18 | 3.348338 | 1.743445 | 0.000237 | 0.001584 | up |
| Galnt17 | 2.080776 | 1.057122 | 0.000239 | 0.001593 | up |
| Ccdc80 | 2.488942 | 1.3155329 | 0.000239 | 0.001594 | up |
| Plekhs1 | 0.245116 | -2.028463 | 0.000242 | 0.00161 | down |
| Sgpp2 | 0.467076 | -1.098272 | 0.000242 | 0.001614 | down |
| Mei4 | 8.975231 | 3.1659491 | 0.000248 | 0.001649 | up |
| Entrep2 | 12.80277 | 3.6783842 | 0.000254 | 0.001677 | up |
| AABR07017236.2 | 0.44797 | -1.158526 | 0.000256 | 0.001689 | down |
| ENSRNOG00000070845 | 0.265889 | -1.911103 | 0.000256 | 0.00169 | down |
| Fam167b | 2.908588 | 1.540319 | 0.000266 | 0.001747 | up |
| Cd80 | 2.469545 | 1.3042454 | 0.000267 | 0.001757 | up |
| Sapcd1 | 7.432644 | 2.8938756 | 0.000269 | 0.001767 | up |
| Rem1 | 2.012544 | 1.0090202 | 0.00028 | 0.001828 | up |
| Izumo4 | 2.319376 | 1.2137365 | 0.000284 | 0.001851 | up |
| Syngr3 | 0.489781 | -1.029792 | 0.000285 | 0.001855 | down |
| Usp13 | 2.114472 | 1.0802973 | 0.000286 | 0.001864 | up |
| Ffar3 | 11.38154 | 3.5086239 | 0.000292 | 0.001899 | up |
| AABR07037520.1 | 4.36351 | 2.1254891 | 0.000297 | 0.001924 | up |
| Scml2 | 3.461771 | 1.7915104 | 0.000299 | 0.001936 | up |
| Cyp2d3 | 2.272091 | 1.1840206 | 0.000301 | 0.001945 | up |
| Mt1 | 2.510111 | 1.3277509 | 0.000301 | 0.001947 | up |
| Gabbr2 | 23.53365 | 4.556653 | 0.000307 | 0.001981 | up |
| St6galnac1 | 740.488 | 9.5323325 | 0.000308 | 0.001986 | up |
| Enho | 0.357089 | -1.485643 | 0.000311 | 0.001999 | down |
| Mest | 2.091711 | 1.0646837 | 0.000326 | 0.002083 | up |
| Lmo1 | 3.430285 | 1.7783285 | 0.000341 | 0.002171 | up |
| LOC120100233 | 2.463378 | 1.3006383 | 0.000349 | 0.002213 | up |

| | | | | | |
|---|---|---|---|---|---|
| Kcns2 | 0.349848 | -1.515201 | 0.000353 | 0.002233 | down |
| ENSRNOG00000068655 | 2.450469 | 1.2930579 | 0.000371 | 0.002332 | up |
| Dnajc6 | 0.489963 | -1.029254 | 0.000376 | 0.002362 | down |
| Ms4a12 | 11.37209 | 3.507425 | 0.000378 | 0.002375 | up |
| Sstr1 | 0.467816 | -1.095986 | 0.000381 | 0.002389 | down |
| Frmpd4 | 0.439577 | -1.185812 | 0.000382 | 0.002394 | down |
| Kcnh4 | 0.423257 | -1.240394 | 0.000391 | 0.002442 | down |
| LOC102555043 | 0.352943 | -1.502494 | 0.000397 | 0.002472 | down |
| LOC120095690 | 2.827588 | 1.4995718 | 0.000412 | 0.002558 | up |
| ENSRNOG00000068528 | 0.473661 | -1.078072 | 0.000419 | 0.002596 | down |
| Sstr2 | 0.267351 | -1.903193 | 0.000423 | 0.002619 | down |
| Hormad2 | 2.529382 | 1.3387851 | 0.000432 | 0.00266 | up |
| Fam107a | 0.473601 | -1.078255 | 0.000441 | 0.00271 | down |
| Ccl22 | 4.213137 | 2.0748948 | 0.000448 | 0.002743 | up |
| Sptssb | 0.074959 | -3.73776 | 0.000449 | 0.002749 | down |
| Kcnip1 | 3.193059 | 1.6749393 | 0.000453 | 0.002771 | up |
| Il17b | 3.325276 | 1.7334741 | 0.000458 | 0.002798 | up |
| Mdga2 | 2.185441 | 1.1279244 | 0.000459 | 0.002799 | up |
| Neu2 | 0.373929 | -1.419163 | 0.000462 | 0.002817 | down |
| Caly | 0.000793 | -10.30013 | 0.000473 | 0.00288 | down |
| Upk2 | 12.25552 | 3.6153596 | 0.000483 | 0.002927 | up |
| Angptl7 | 3.720407 | 1.8954603 | 0.000496 | 0.002999 | up |
| Ppp1r14b-ps2 | 2.072161 | 1.0511362 | 0.000502 | 0.003033 | up |
| Ppp2r3b | 2.635234 | 1.3979312 | 0.000511 | 0.003077 | up |
| Aqp7 | 0.019645 | -5.669721 | 0.000514 | 0.003095 | down |
| Oas1k | 0.462713 | -1.11181 | 0.000525 | 0.00315 | down |
| Cyp2g1 | 0.175095 | -2.513788 | 0.000535 | 0.003208 | down |
| Vxn | 3.199522 | 1.6778562 | 0.00054 | 0.003234 | up |
| Rnase2 | 16.25617 | 4.0229159 | 0.000547 | 0.00327 | up |
| Tigd3 | 2.162071 | 1.1124136 | 0.000571 | 0.003399 | up |
| LOC120095179 | 2.622936 | 1.3911827 | 0.000572 | 0.003405 | up |
| Ascl1 | 0.388475 | -1.364105 | 0.00058 | 0.003447 | down |
| Cma1 | 2.467436 | 1.3030127 | 0.000582 | 0.00346 | up |
| Thy1 | 2.129487 | 1.0905057 | 0.000583 | 0.003466 | up |
| ENSRNOG00000063439 | 3.012808 | 1.5911088 | 0.000589 | 0.003493 | up |
| Pcdhb5 | 2.163094 | 1.1130965 | 0.000589 | 0.003493 | up |
| Brinp1 | 2.349045 | 1.2320741 | 0.000589 | 0.003495 | up |
| RGD1308065 | 3.00927 | 1.5894135 | 0.000597 | 0.003534 | up |
| Acnat2 | 952.604 | 9.8957328 | 0.000606 | 0.003581 | up |
| Dbp | 3.241307 | 1.6965758 | 0.000615 | 0.003625 | up |
| Sertm1 | 0.387571 | -1.367468 | 0.000622 | 0.003659 | down |

| | | | | | |
|---|---|---|---|---|---|
| Ucp1 | 0.003513 | -8.152963 | 0.000623 | 0.003666 | down |
| Ppargc1a | 0.464565 | -1.106049 | 0.000625 | 0.003673 | down |
| Slc26a4 | 5.993963 | 2.5835102 | 0.000641 | 0.003758 | up |
| Muc20 | 0.499026 | -1.002813 | 0.000642 | 0.00376 | down |
| Tac4 | 3.603731 | 1.8494913 | 0.000653 | 0.00382 | up |
| Qprt | 2.738573 | 1.4534244 | 0.000677 | 0.003943 | up |
| Meox1 | 2.50606 | 1.3254212 | 0.000679 | 0.003951 | up |
| LOC120093043 | 2.548581 | 1.3496942 | 0.00068 | 0.003958 | up |
| Hrc | 0.368337 | -1.440903 | 0.000685 | 0.003983 | down |
| ENSRNOG00000066084 | 0.320959 | -1.63954 | 0.000688 | 0.003996 | down |
| Prss35 | 2.447499 | 1.2913085 | 0.000699 | 0.004051 | up |
| Hpdl | 2.260253 | 1.176484 | 0.00071 | 0.004106 | up |
| Dmrta1 | 0.167993 | -2.573527 | 0.000743 | 0.004266 | down |
| Sult4a1 | 3.874383 | 1.9539666 | 0.000751 | 0.004305 | up |
| Rlbp1 | 534.11 | 9.0609931 | 0.000757 | 0.004331 | up |
| LOC100360522 | 0.499915 | -1.000246 | 0.000758 | 0.004336 | down |
| Lpar4 | 2.117732 | 1.0825202 | 0.000764 | 0.004369 | up |
| Bex1 | 0.334749 | -1.578847 | 0.000769 | 0.004394 | down |
| Stkld1 | 578.6508 | 9.1765492 | 0.000779 | 0.004437 | up |
| Plekhg4 | 8.262357 | 3.0465534 | 0.000786 | 0.004474 | up |
| Tmprss7 | 441.1291 | 8.7850572 | 0.000787 | 0.004477 | up |
| Or2b11 | 599 | 9.2264122 | 0.000793 | 0.004511 | up |
| Alkal1 | 41.16688 | 5.363412 | 0.000799 | 0.00454 | up |
| S100b | 0.405905 | -1.300787 | 0.0008 | 0.004545 | down |
| ENSRNOG00000062487 | 671.6502 | 9.3915663 | 0.000839 | 0.004735 | up |
| Ly6h | 11.05006 | 3.4659817 | 0.000843 | 0.004755 | up |
| H2ac4 | 12.17832 | 3.6062435 | 0.000846 | 0.004768 | up |
| Nxf7 | 42.92462 | 5.4237335 | 0.000855 | 0.004817 | up |
| Tex15 | 0.30471 | -1.714489 | 0.000859 | 0.004834 | down |
| AC129365.1 | 3.389368 | 1.7610161 | 0.000871 | 0.004893 | up |
| ENSRNOG00000068419 | 2.137175 | 1.095705 | 0.000874 | 0.004907 | up |
| Snord28b | 4.297526 | 2.1035062 | 0.000876 | 0.004913 | up |
| Gkn3 | 2.712816 | 1.4397912 | 0.000924 | 0.005143 | up |
| Scgb3a1 | 0.239583 | -2.0614 | 0.000928 | 0.005164 | down |
| Pheta2 | 2.058701 | 1.0417346 | 0.000947 | 0.005258 | up |
| Impg1 | 737.8373 | 9.5271589 | 0.000963 | 0.005335 | up |
| Nlgn3 | 2.234658 | 1.1600543 | 0.001003 | 0.00552 | up |
| Lhx2 | 615.74 | 9.2661775 | 0.001008 | 0.005547 | up |
| Sema4g | 2.023231 | 1.0166612 | 0.001009 | 0.005547 | up |
| Fam169a | 0.366311 | -1.44886 | 0.001038 | 0.005694 | down |
| Hif3a | 0.03003 | -5.057434 | 0.001055 | 0.00577 | down |

| | | | | | |
|---|---|---|---|---|---|
| Frmpd1 | 2.456044 | 1.2963364 | 0.001064 | 0.005811 | up |
| Fxyd2 | 2.022501 | 1.0161402 | 0.00109 | 0.005943 | up |
| ENSRNOG00000066012 | 201.724 | 7.6562387 | 0.001122 | 0.006085 | up |
| Aif1l | 2.100148 | 1.0704907 | 0.001123 | 0.006088 | up |
| LOC102550644 | 0.495234 | -1.013817 | 0.001127 | 0.006105 | down |
| Rab7b | 2.069784 | 1.0494803 | 0.001155 | 0.006236 | up |
| ENSRNOG00000070692 | 2.339809 | 1.226391 | 0.001181 | 0.00635 | up |
| ENSRNOG00000064944 | 2.185456 | 1.1279341 | 0.001182 | 0.006353 | up |
| ENSRNOG00000064366 | 0.39024 | -1.357567 | 0.001183 | 0.006353 | down |
| Ly49i4 | 0.364908 | -1.454397 | 0.0012 | 0.006434 | down |
| Zfp483 | 0.464645 | -1.105798 | 0.001202 | 0.006443 | down |
| Ranbp3l | 0.484878 | -1.044307 | 0.001209 | 0.006477 | down |
| Pde10a | 7.786829 | 2.9610359 | 0.001215 | 0.006502 | up |
| Fmo6 | 0.229798 | -2.121565 | 0.001219 | 0.006525 | down |
| LOC100364427 | 0.446745 | -1.162476 | 0.00125 | 0.006665 | down |
| Ifitm5 | 2.802983 | 1.486963 | 0.001259 | 0.006706 | up |
| Slc6a15 | 0.302342 | -1.725748 | 0.001278 | 0.0068 | down |
| Rcor2 | 2.198087 | 1.1362487 | 0.001315 | 0.006969 | up |
| Grin3a | 31.47317 | 4.9760508 | 0.001322 | 0.006997 | up |
| Scin | 0.433239 | -1.206765 | 0.001335 | 0.007055 | down |
| Armh4 | 2.351344 | 1.2334857 | 0.001341 | 0.007084 | up |
| Lilrb4 | 2.04881 | 1.0347862 | 0.001384 | 0.007281 | up |
| Faxc | 2.430561 | 1.2812895 | 0.001413 | 0.007417 | up |
| Hmgcs2 | 0.438157 | -1.190479 | 0.001419 | 0.007439 | down |
| Serpind1 | 0.473734 | -1.07785 | 0.001422 | 0.007451 | down |
| Sox8 | 0.143765 | -2.798214 | 0.001428 | 0.007476 | down |
| Pnma2 | 2.005825 | 1.004196 | 0.001433 | 0.007491 | up |
| Sh3rf3 | 2.404915 | 1.2659861 | 0.001442 | 0.007531 | up |
| Krt82 | 611.5937 | 9.2564297 | 0.001445 | 0.007542 | up |
| ENSRNOG00000066276 | 3.433451 | 1.7796595 | 0.001469 | 0.007651 | up |
| Bpifb2 | 12.09568 | 3.5964203 | 0.001482 | 0.007705 | up |
| Bcas1 | 5.616364 | 2.4896363 | 0.001491 | 0.007751 | up |
| Sectm1b | 4.621411 | 2.2083333 | 0.001516 | 0.007858 | up |
| LOC120103148 | 2.378491 | 1.2500466 | 0.001528 | 0.007912 | up |
| Sdsl | 2.316595 | 1.212006 | 0.001532 | 0.007925 | up |
| Abca13 | 0.125439 | -2.994942 | 0.001532 | 0.007925 | down |
| Defa5 | 4320.686 | 12.077045 | 0.001557 | 0.008035 | up |
| Zbtb16 | 0.263685 | -1.923114 | 0.001567 | 0.008075 | down |
| Per3 | 2.56139 | 1.3569271 | 0.001593 | 0.008193 | up |
| G0s2 | 0.430834 | -1.214797 | 0.001624 | 0.008319 | down |
| Gpr3 | 4.927221 | 2.3007743 | 0.001631 | 0.00835 | up |

| | | | | | |
|---|---|---|---|---|---|
| LOC120102039 | 2.892977 | 1.5325546 | 0.001659 | 0.008486 | up |
| ENSRNOG00000067342 | 0.461372 | -1.115997 | 0.00166 | 0.00849 | down |
| ENSRNOG00000063720 | 4.076214 | 2.0272299 | 0.001686 | 0.0086 | up |
| Pax9 | 0.272863 | -1.873752 | 0.001687 | 0.008602 | down |
| RGD1565779 | 11.74036 | 3.5534045 | 0.00169 | 0.008614 | up |
| Kng1 | 2.94754 | 1.5595115 | 0.00169 | 0.008615 | up |
| Retnla | 4.470004 | 2.1602761 | 0.0017 | 0.008664 | up |
| Adra1d | 2.002454 | 1.0017692 | 0.001715 | 0.008732 | up |
| Popdc3 | 0.354516 | -1.496078 | 0.001721 | 0.008759 | down |
| Bcat1 | 2.063253 | 1.044921 | 0.001746 | 0.008863 | up |
| LOC120097726 | 7.746245 | 2.9534971 | 0.001756 | 0.008903 | up |
| Myh2 | 8.206184 | 3.0367115 | 0.001779 | 0.009007 | up |
| Cd84 | 2.422897 | 1.2767331 | 0.001789 | 0.009047 | up |
| Trim9 | 0.384082 | -1.380512 | 0.001801 | 0.009091 | down |
| Tcfl5 | 6.870065 | 2.7803237 | 0.001806 | 0.009115 | up |
| ENSRNOG00000065998 | 2.668517 | 1.4160382 | 0.001843 | 0.009266 | up |
| LOC100911032 | 3.37E-05 | -14.85692 | 0.001849 | 0.009288 | down |
| Fndc3c1 | 224.9903 | 7.8137187 | 0.00186 | 0.009331 | up |
| Foxc2 | 2.644134 | 1.4027952 | 0.001862 | 0.009335 | up |
| Slc7a11 | 4.689566 | 2.2294543 | 0.001894 | 0.00947 | up |
| Tlcd2 | 2.80506 | 1.4880317 | 0.001896 | 0.009481 | up |
| Lrrc7 | 0.102324 | -3.288779 | 0.00191 | 0.009533 | down |
| Sema7a | 2.167246 | 1.1158627 | 0.001913 | 0.009549 | up |
| Neil3 | 10.1964 | 3.3499881 | 0.001942 | 0.009673 | up |
| Car1 | 8.755136 | 3.1301296 | 0.001952 | 0.009715 | up |
| Tmprss11b | 0.393179 | -1.346742 | 0.002004 | 0.009937 | down |
| LOC120097031 | 0.344945 | -1.53556 | 0.002066 | 0.010183 | down |
| Ankrd34a | 2.070937 | 1.0502836 | 0.002069 | 0.010194 | up |
| ENSRNOG00000068893 | 4.233484 | 2.0818453 | 0.002073 | 0.010207 | up |
| Ankdd1b | 3.214392 | 1.6845457 | 0.002105 | 0.010327 | up |
| ENSRNOG00000069434 | 17.61681 | 4.1388807 | 0.00219 | 0.010692 | up |
| ENSRNOG00000066286 | 659.396 | 9.3650013 | 0.002214 | 0.010798 | up |
| LOC100912568 | 0.365347 | -1.45266 | 0.002224 | 0.010839 | down |
| Gfpt2 | 2.266839 | 1.1806822 | 0.002279 | 0.011073 | up |
| ENSRNOG00000071123 | 2.524608 | 1.3360596 | 0.002289 | 0.011121 | up |
| Ms4a14 | 20.22104 | 4.3377854 | 0.0023 | 0.011159 | up |
| Nhs | 2.414977 | 1.2720094 | 0.00231 | 0.011204 | up |
| Inhba | 5.598914 | 2.4851471 | 0.00232 | 0.011233 | up |
| Soga3 | 0.486069 | -1.040766 | 0.002367 | 0.011428 | down |
| Oas1e | 2.283289 | 1.1911132 | 0.002383 | 0.011493 | up |
| Rbp7 | 2.338309 | 1.2254658 | 0.002425 | 0.01167 | up |

| | | | | | |
|---|---|---|---|---|---|
| LOC689757 | 5.471404 | 2.451911 | 0.002449 | 0.011766 | up |
| ENSRNOG00000068486 | 2.810782 | 1.4909715 | 0.00249 | 0.011931 | up |
| Ly6al | 11.30645 | 3.4990746 | 0.002518 | 0.012042 | up |
| Ctsg | 1402.792 | 10.454085 | 0.002523 | 0.012062 | up |
| Tex36 | 0.153595 | -2.702801 | 0.002588 | 0.012323 | down |
| Mttp | 0.365867 | -1.450611 | 0.002595 | 0.01235 | down |
| Dppa3 | 12.89199 | 3.688403 | 0.002616 | 0.012426 | up |
| AABR07048770.1 | 1500.764 | 10.551481 | 0.002669 | 0.012656 | up |
| Or5an6 | 519.8516 | 9.0219561 | 0.0027 | 0.012778 | up |
| Ces1f | 2.536018 | 1.342565 | 0.002717 | 0.012851 | up |
| Stac3 | 22.7718 | 4.5091763 | 0.00276 | 0.013024 | up |
| Krt87 | 13.25437 | 3.7283957 | 0.002775 | 0.013082 | up |
| Arpp21 | 2.289744 | 1.195186 | 0.00278 | 0.013097 | up |
| ENSRNOG00000065110 | 2.815373 | 1.4933258 | 0.002782 | 0.013103 | up |
| Sctr | 4.951486 | 2.3078614 | 0.002799 | 0.013176 | up |
| Lynx1 | 0.407658 | -1.294569 | 0.002817 | 0.01325 | down |
| Hsd17b14 | 0.31079 | -1.68599 | 0.002845 | 0.013355 | down |
| Asb11 | 0.240421 | -2.056363 | 0.002867 | 0.013445 | down |
| Khdrbs2 | 250.4449 | 7.9683493 | 0.002884 | 0.013513 | up |
| Plin4 | 0.455826 | -1.133445 | 0.002922 | 0.013676 | down |
| Atp6v1b1 | 0.410724 | -1.283757 | 0.002925 | 0.013687 | down |
| Dnajc5g | 620.6807 | 9.2777074 | 0.00294 | 0.013737 | up |
| Pcdhb12 | 2.109766 | 1.0770828 | 0.002949 | 0.013772 | up |
| LOC103690878 | 9.112972 | 3.1879216 | 0.00296 | 0.013813 | up |
| LOC102552404 | 7.695225 | 2.9439635 | 0.003061 | 0.014221 | up |
| Grifin | 0.014555 | -6.102321 | 0.003073 | 0.014263 | down |
| C4a | 2.534386 | 1.3416362 | 0.00308 | 0.014291 | up |
| Capn3 | 3.102223 | 1.6333023 | 0.003112 | 0.014422 | up |
| AABR07049223.1 | 2.347768 | 1.2312896 | 0.003162 | 0.014634 | up |
| Sprr2g | 0.000397 | -11.29742 | 0.00321 | 0.014814 | down |
| Unc5d | 0.08192 | -3.609643 | 0.003213 | 0.014829 | down |
| Il20ra | 0.189579 | -2.399125 | 0.003238 | 0.014928 | down |
| RGD1561722 | 4.83E-05 | -14.33658 | 0.003293 | 0.015158 | down |
| ENSRNOG00000066189 | 169.684 | 7.4067067 | 0.003324 | 0.015282 | up |
| Pla2g4c | 60.59013 | 5.9210109 | 0.003345 | 0.01537 | up |
| Hist1h4m | 10.15597 | 3.344256 | 0.003386 | 0.015527 | up |
| Htr2a | 2.747397 | 1.4580654 | 0.003418 | 0.015635 | up |
| Tceal5 | 3.253883 | 1.7021624 | 0.003458 | 0.015773 | up |
| Adgrb1 | 2.084855 | 1.0599469 | 0.003487 | 0.015868 | up |
| Npas1 | 0.497505 | -1.007217 | 0.003528 | 0.016024 | down |
| Lypd8 | 4.328512 | 2.1138712 | 0.003548 | 0.016092 | up |

| | | | | | |
|---|---|---|---|---|---|
| Acsbg1 | 2.19533 | 1.1344379 | 0.003549 | 0.016094 | up |
| AABR07030494.1 | 2.231141 | 1.1577818 | 0.003648 | 0.016487 | up |
| ENSRNOG00000065343 | 0.378134 | -1.403029 | 0.003658 | 0.016529 | down |
| LOC680920 | 6.594981 | 2.7213684 | 0.003729 | 0.016797 | up |
| Aard | 7.57943 | 2.9220894 | 0.003748 | 0.016863 | up |
| Cdh9 | 0.100717 | -3.311619 | 0.003752 | 0.016874 | down |
| L1td1 | 2.140745 | 1.0981127 | 0.003869 | 0.017331 | up |
| Adgrf3 | 3.156201 | 1.6581891 | 0.003889 | 0.017404 | up |
| Pax5 | 0.492308 | -1.022366 | 0.003914 | 0.017494 | down |
| Ces2c | 0.442097 | -1.177567 | 0.003938 | 0.01759 | down |
| ENSRNOG00000065629 | 0.23506 | -2.088897 | 0.00394 | 0.017596 | down |
| Trpa1 | 2.701712 | 1.4338741 | 0.003995 | 0.017815 | up |
| Hmgb2l1 | 2.902109 | 1.5371015 | 0.004072 | 0.018136 | up |
| Ar | 0.467265 | -1.097687 | 0.004174 | 0.018516 | down |
| Cdkn2a | 2.693489 | 1.4294762 | 0.004216 | 0.018685 | up |
| Sycp3 | 0.499083 | -1.002648 | 0.004264 | 0.018879 | down |
| LOC498465 | 0.000957 | -10.02844 | 0.004311 | 0.019076 | down |
| Hes2 | 3.741418 | 1.9035852 | 0.004315 | 0.019088 | up |
| Clec4b2 | 28.47377 | 4.8315616 | 0.004348 | 0.019198 | up |
| Mfsd4b2 | 2.803378 | 1.4871662 | 0.004389 | 0.019361 | up |
| Fam25a | 6.222064 | 2.6373933 | 0.00441 | 0.01943 | up |
| Igf2bp1 | 652.0492 | 9.348837 | 0.004416 | 0.019454 | up |
| Hist1h2ail1 | 5.79804 | 2.5355652 | 0.004565 | 0.019986 | up |
| Prkag3 | 0.00351 | -8.154312 | 0.004565 | 0.019986 | down |
| Pnoc | 24.00586 | 4.5853149 | 0.004619 | 0.020186 | up |
| Epm2a | 2.768258 | 1.4689786 | 0.004674 | 0.020374 | up |
| Htr2c | 0.421129 | -1.247664 | 0.004737 | 0.0206 | down |
| Spats1 | 2.210064 | 1.144088 | 0.004833 | 0.020962 | up |
| LOC100125362 | 2.444278 | 1.2894086 | 0.004857 | 0.021053 | up |
| Prss22 | 2.047384 | 1.0337815 | 0.004919 | 0.021281 | up |
| Chrnb2 | 23.58974 | 4.5600876 | 0.004928 | 0.021313 | up |
| Tlr11 | 2.335242 | 1.2235723 | 0.004979 | 0.021492 | up |
| Pla2g10 | 3.542068 | 1.8245918 | 0.004983 | 0.021508 | up |
| Mab21l1 | 6.054627 | 2.5980381 | 0.005152 | 0.022153 | up |
| ENSRNOG00000070816 | 0.171822 | -2.541011 | 0.005203 | 0.022339 | down |
| Fam115e | 0.003994 | -7.968079 | 0.005222 | 0.022419 | down |
| ENSRNOG00000069250 | 0.026975 | -5.212241 | 0.005281 | 0.022592 | down |
| Hao2 | 1056.795 | 10.04548 | 0.005293 | 0.022629 | up |
| Septin3 | 0.481684 | -1.053842 | 0.005464 | 0.023218 | down |
| Gabrp | 2.117 | 1.0820215 | 0.005466 | 0.02322 | up |
| Zscan18 | 0.465162 | -1.104196 | 0.005467 | 0.02322 | down |

| | | | | | |
|---|---|---|---|---|---|
| Tac1 | 0.156044 | -2.679973 | 0.005495 | 0.023307 | down |
| Esrrg | 0.489312 | -1.031173 | 0.005496 | 0.023308 | down |
| Grk1 | 400.1 | 8.6442168 | 0.005513 | 0.02337 | up |
| Pak5 | 201.784 | 7.656668 | 0.005522 | 0.023403 | up |
| AC131806.3 | 0.000308 | -11.6647 | 0.00553 | 0.02343 | down |
| RGD1565143 | 3.891728 | 1.960411 | 0.005535 | 0.023444 | up |
| LOC120102924 | 0.383757 | -1.381733 | 0.00576 | 0.024251 | down |
| Barx2 | 0.432018 | -1.210838 | 0.00578 | 0.024315 | down |
| Greb1 | 2.642372 | 1.4018337 | 0.005788 | 0.024344 | up |
| Vsx1 | 0.286904 | -1.801359 | 0.005829 | 0.024501 | down |
| Necab1 | 6.160844 | 2.623128 | 0.005848 | 0.024561 | up |
| Slc22a14 | 387.092 | 8.5965327 | 0.005858 | 0.024599 | up |
| Gjb6 | 4.152303 | 2.0539117 | 0.005877 | 0.024672 | up |
| Aox1 | 0.428191 | -1.223673 | 0.005879 | 0.024676 | down |
| Stfa2l3 | 0.486378 | -1.039849 | 0.005908 | 0.024776 | down |
| Or51e1 | 3.923845 | 1.9722682 | 0.005953 | 0.024957 | up |
| Dpy30 | 9.133911 | 3.1912327 | 0.00599 | 0.025088 | up |
| Wnt2b | 0.419072 | -1.254731 | 0.006014 | 0.02517 | down |
| LOC102550466 | 2.690282 | 1.4277572 | 0.00606 | 0.025332 | up |
| ENSRNOG00000063514 | 2.101625 | 1.071505 | 0.006077 | 0.025401 | up |
| Ccn1 | 2.872178 | 1.5221453 | 0.006168 | 0.025733 | up |
| Kash5 | 2.122248 | 1.0855933 | 0.006173 | 0.025745 | up |
| Dmgdh | 6.844966 | 2.7750435 | 0.006211 | 0.025879 | up |
| ENSRNOG00000069297 | 0.427003 | -1.227682 | 0.006277 | 0.026112 | down |
| Trem2 | 2.677685 | 1.4209864 | 0.006409 | 0.026569 | up |
| Atg9b | 5.740335 | 2.5211349 | 0.006425 | 0.02661 | up |
| Vpreb2 | 21.01989 | 4.3936829 | 0.006476 | 0.026785 | up |
| ENSRNOG00000071196 | 0.492521 | -1.021744 | 0.006507 | 0.026873 | down |
| ENSRNOG00000063859 | 0.142249 | -2.81351 | 0.006519 | 0.026903 | down |
| Gpr162 | 2.32592 | 1.2178016 | 0.006606 | 0.02722 | up |
| Lexm | 11.28327 | 3.4961134 | 0.006621 | 0.027263 | up |
| AABR07021591.1 | 8.128199 | 3.0229357 | 0.006634 | 0.027307 | up |
| Sbk2 | 0.350734 | -1.511553 | 0.006657 | 0.027386 | down |
| Serpina6 | 0.039655 | -4.656348 | 0.006659 | 0.027389 | down |
| Pnpla3 | 0.270005 | -1.888943 | 0.006716 | 0.027582 | down |
| Car9 | 2.348151 | 1.2315253 | 0.006736 | 0.027651 | up |
| Cadm4 | 2.29066 | 1.1957637 | 0.006744 | 0.027671 | up |
| LOC120101644 | 2.229887 | 1.1569709 | 0.006902 | 0.02817 | up |
| ENSRNOG00000064389 | 0.065901 | -3.92355 | 0.006985 | 0.028459 | down |
| ENSRNOG00000062545 | 0.42885 | -1.221456 | 0.007087 | 0.028792 | down |
| Rpl6-ps1 | 2.15061 | 1.1047459 | 0.007145 | 0.029011 | up |

| | | | | | |
|---|---|---|---|---|---|
| ENSRNOG00000066122 | 2.693512 | 1.4294885 | 0.007175 | 0.029095 | up |
| Ccdc150 | 2.042308 | 1.0302005 | 0.0073 | 0.029504 | up |
| Poteg | 0.005095 | -7.616607 | 0.00734 | 0.029654 | down |
| ENSRNOG00000063857 | 2.492704 | 1.3177114 | 0.007392 | 0.029849 | up |
| ENSRNOG00000064064 | 0.332776 | -1.587375 | 0.007545 | 0.030383 | down |
| Adgrl2 | 4.448863 | 2.1534365 | 0.007562 | 0.030433 | up |
| Tp63 | 0.395756 | -1.337318 | 0.007589 | 0.03052 | down |
| ENSRNOG00000069140 | 0.281274 | -1.829951 | 0.007637 | 0.030669 | down |
| ENSRNOG00000063050 | 0.391227 | -1.353921 | 0.007644 | 0.03069 | down |
| ENSRNOG00000066903 | 0.495077 | -1.014274 | 0.007706 | 0.030877 | down |
| Rasgef1c | 7.549812 | 2.9164407 | 0.007742 | 0.030983 | up |
| Il24 | 2.595403 | 1.3759587 | 0.007769 | 0.031087 | up |
| Hs3st4 | 12.07163 | 3.5935483 | 0.007774 | 0.031101 | up |
| AABR07072761.1 | 2.557052 | 1.3544813 | 0.007847 | 0.031378 | up |
| ENSRNOG00000069708 | 2.383281 | 1.2529492 | 0.007977 | 0.031788 | up |
| Prg2 | 3.528917 | 1.8192257 | 0.007998 | 0.031859 | up |
| AABR07058124.2 | 0.000284 | -11.78322 | 0.008058 | 0.032056 | down |
| LOC120094453 | 2.827578 | 1.499567 | 0.00812 | 0.032265 | up |
| LOC683313 | 695.288 | 9.4414669 | 0.008406 | 0.033269 | up |
| Epx | 2.741821 | 1.4551346 | 0.00851 | 0.03363 | up |
| Trdn | 0.491354 | -1.025166 | 0.008564 | 0.033816 | down |
| ENSRNOG00000063033 | 0.426544 | -1.229235 | 0.008573 | 0.033844 | down |
| Synpo2l | 3.355724 | 1.746624 | 0.008583 | 0.033879 | up |
| Adcyap1r1 | 0.38213 | -1.387863 | 0.008633 | 0.034032 | down |
| ENSRNOG00000069130 | 3.522476 | 1.8165898 | 0.008783 | 0.034514 | up |
| ENSRNOG00000066935 | 0.369822 | -1.435097 | 0.008934 | 0.035011 | down |
| Mirlet7d | 3.371557 | 1.753415 | 0.008937 | 0.035016 | up |
| Nrap | 0.346357 | -1.529666 | 0.008955 | 0.035075 | down |
| Siglec15 | 4.891951 | 2.2904098 | 0.009044 | 0.035372 | up |
| Tm4sf4 | 0.311237 | -1.683915 | 0.00908 | 0.035495 | down |
| Hist1h3b | 2.088825 | 1.0626916 | 0.009094 | 0.035546 | up |
| Fgf13 | 0.41482 | -1.269442 | 0.009149 | 0.035743 | down |
| ENSRNOG00000067879 | 5.041278 | 2.3337894 | 0.009185 | 0.035863 | up |
| Prss29 | 0.000826 | -10.24132 | 0.009237 | 0.036027 | down |
| Slc12a1 | 0.338148 | -1.564272 | 0.009267 | 0.036131 | down |
| Rims1 | 0.132627 | -2.914549 | 0.009284 | 0.036173 | down |
| ENSRNOG00000065203 | 0.003393 | -8.203352 | 0.009387 | 0.036495 | down |
| Scn10a | 3.272036 | 1.7101886 | 0.009393 | 0.036509 | up |
| Ldlrad2 | 4.479113 | 2.1632129 | 0.009455 | 0.036718 | up |
| Snap91 | 2.217503 | 1.1489361 | 0.009469 | 0.03676 | up |
| Rhebl1 | 2.084645 | 1.0598017 | 0.009627 | 0.037273 | up |

| | | | | | |
|---|---|---|---|---|---|
| Cryba2 | 1734.02 | 10.759905 | 0.009628 | 0.037273 | up |
| Krt81 | 328.4445 | 8.3595057 | 0.009683 | 0.037455 | up |
| Hpcal4 | 0.465465 | -1.103255 | 0.009691 | 0.037481 | down |
| St8sia3 | 0.001818 | -9.103529 | 0.009724 | 0.037584 | down |
| Stk32b | 8.04389 | 3.0078933 | 0.009746 | 0.037659 | up |
| ENSRNOG00000070051 | 0.000589 | -10.7299 | 0.009782 | 0.037773 | down |
| ENSRNOG00000066281 | 0.090102 | -3.472305 | 0.009931 | 0.038224 | down |
| Snord27 | 2.964655 | 1.5678643 | 0.01 | 0.038436 | up |
| Cpa3 | 2.021709 | 1.0155751 | 0.010062 | 0.038649 | up |
| Calcr | 0.002928 | -8.416071 | 0.010067 | 0.03866 | down |
| Klhl33 | 0.387956 | -1.366034 | 0.010271 | 0.039309 | down |
| R3hdml | 32.33642 | 5.0150879 | 0.010361 | 0.039584 | up |
| Dhrs7c | 0.273048 | -1.872772 | 0.010419 | 0.039781 | down |
| Myoz2 | 0.308251 | -1.697825 | 0.010425 | 0.039796 | down |
| LOC100909660 | 2629.15 | 11.360381 | 0.010463 | 0.039909 | up |
| AABR07019088.1 | 0.494762 | -1.015193 | 0.010685 | 0.040635 | down |
| Esrrb | 0.457893 | -1.126917 | 0.010706 | 0.04071 | down |
| ENSRNOG00000069233 | 3.182449 | 1.6701373 | 0.010837 | 0.041136 | up |
| Draxin | 2.317843 | 1.2127826 | 0.01088 | 0.041294 | up |
| Ctxnd1 | 78.37756 | 6.2923688 | 0.0111 | 0.042058 | up |
| Arl14 | 1177.294 | 10.201259 | 0.011153 | 0.042236 | up |
| Exoc1l | 0.356886 | -1.486465 | 0.011168 | 0.04228 | down |
| Gng13 | 0.35588 | -1.490538 | 0.01118 | 0.042305 | down |
| Dpp6 | 2.322592 | 1.2157356 | 0.011316 | 0.042769 | up |
| ENSRNOG00000063598 | 0.397069 | -1.332538 | 0.011345 | 0.042843 | down |
| Kcnv2 | 2.033537 | 1.0239911 | 0.01139 | 0.042991 | up |
| B3gnt3 | 2.093471 | 1.0658968 | 0.011398 | 0.043006 | up |
| Eda2r | 4.091837 | 2.0327488 | 0.011522 | 0.043385 | up |
| Txlnb | 0.315249 | -1.665435 | 0.011528 | 0.043385 | down |
| Ndst3 | 2.19524 | 1.1343789 | 0.01155 | 0.043448 | up |
| AABR07065789.1 | 0.102787 | -3.28227 | 0.011579 | 0.043543 | down |
| Slc2a5 | 3.615404 | 1.8541568 | 0.011616 | 0.043652 | up |
| ENSRNOG00000066124 | 0.40379 | -1.308324 | 0.011679 | 0.043836 | down |
| LOC120094450 | 4.016818 | 2.0060532 | 0.011789 | 0.044176 | up |
| Tnfrsf13c | 0.499273 | -1.002099 | 0.011794 | 0.04418 | down |
| AABR07034928.1 | 11.17459 | 3.4821496 | 0.011943 | 0.044671 | up |
| Rps12l4 | 2.006841 | 1.0049261 | 0.011988 | 0.044817 | up |
| Bpifc | 4.025297 | 2.0090952 | 0.012012 | 0.044882 | up |
| Clec6a-ps1 | 3406.209 | 11.733951 | 0.012045 | 0.044961 | up |
| Cd177 | 0.409363 | -1.288548 | 0.012069 | 0.045009 | down |
| Efnb3 | 0.338155 | -1.564243 | 0.012071 | 0.045009 | down |

| | | | | | |
|---|---|---|---|---|---|
| Has2 | 2.787531 | 1.4789877 | 0.012308 | 0.045728 | up |
| Cdnf | 0.445208 | -1.167449 | 0.012343 | 0.04582 | down |
| RGD1565653 | 0.359399 | -1.476341 | 0.01242 | 0.046061 | down |
| Glt1d1 | 2.585007 | 1.3701682 | 0.012462 | 0.04618 | up |
| LOC120094464 | 2.322982 | 1.2159782 | 0.012464 | 0.04618 | up |
| Tssk4 | 3.368979 | 1.7523115 | 0.01252 | 0.046351 | up |
| Tenm1 | 5.73048 | 2.5186561 | 0.012639 | 0.046726 | up |
| Ddah2 | 2.034535 | 1.0246993 | 0.012719 | 0.046976 | up |
| Zg16b | 0.460791 | -1.117817 | 0.01275 | 0.047065 | down |
| Lilrb3 | 4.557472 | 2.1882337 | 0.012842 | 0.047349 | up |
| Efcab5 | 11.05137 | 3.4661533 | 0.012883 | 0.047443 | up |
| Cidea | 0.462876 | -1.111301 | 0.012888 | 0.047443 | down |
| Irgc | 3.765497 | 1.9128404 | 0.012902 | 0.04748 | up |
| Cacng1 | 16.48608 | 4.0431769 | 0.013011 | 0.047826 | up |
| Alx1 | 0.346483 | -1.529145 | 0.013045 | 0.047919 | down |
| Dlk2 | 0.159174 | -2.651319 | 0.01325 | 0.048554 | down |
| Syt6 | 0.319732 | -1.645067 | 0.013479 | 0.049273 | down |
| Hormad1 | 381.2117 | 8.5744486 | 0.013588 | 0.049599 | up |
| Tph1 | 2.371547 | 1.2458285 | 0.013723 | 0.049995 | up |

**Table 2. Male BLEO vs Male SHAM mRNA-seq**

Significantly Differentially Expressed Genes ($Log_2FC > |1|$; qvalue < 0.05)

| Gene | fc | $log_2(fc)$ | pval | qval | regulation |
|---|---|---|---|---|---|
| Ptprv | 38.21173 | 5.2559438 | 7.14E-47 | 1.66E-42 | up |
| Tnfrsf22 | 11.56455 | 3.5316378 | 2.65E-44 | 3.07E-40 | up |
| Syt13 | 16.04864 | 4.004379 | 3.54E-41 | 2.74E-37 | up |
| C1qa | 14.62869 | 3.8707284 | 9.78E-41 | 5.67E-37 | up |
| Plod2 | 5.265177 | 2.3964821 | 1.86E-39 | 8.63E-36 | up |
| Ces1d | 0.162992 | -2.617128 | 1.97E-38 | 7.61E-35 | down |
| Cyp2b1 | 0.174042 | -2.522494 | 8.29E-37 | 2.75E-33 | down |
| Prrx1 | 4.505815 | 2.1717882 | 8.74E-36 | 2.53E-32 | up |
| C1qb | 11.2934 | 3.4974078 | 1.51E-35 | 3.88E-32 | up |
| Slc26a4 | 21.19505 | 4.4056554 | 3.4E-35 | 7.88E-32 | up |
| Thbs2 | 12.48172 | 3.6417449 | 1.51E-33 | 3.18E-30 | up |
| Cyp2b2 | 0.152594 | -2.712232 | 2.87E-33 | 5.55E-30 | down |
| Fmo1 | 0.272737 | -1.874417 | 2.57E-32 | 4.32E-29 | down |
| Rab7b | 3.755035 | 1.9088263 | 2.61E-32 | 4.32E-29 | up |
| Cthrc1 | 7.516432 | 2.910048 | 4.1E-31 | 6.34E-28 | up |
| Col5a3 | 7.608985 | 2.9277039 | 5.11E-31 | 7.4E-28 | up |
| Retnla | 15.79393 | 3.9812982 | 1.27E-30 | 1.73E-27 | up |
| Mybl2 | 6.168133 | 2.6248338 | 2.85E-30 | 3.67E-27 | up |
| Clec10a | 10.80354 | 3.4334316 | 4.62E-30 | 5.64E-27 | up |
| Ptprn | 20.83921 | 4.381229 | 6.28E-30 | 7.29E-27 | up |
| C1qc | 8.926906 | 3.1581603 | 7.4E-30 | 8.17E-27 | up |
| Gldn | 20.31527 | 4.344493 | 1.86E-29 | 1.96E-26 | up |
| Synm | 0.358724 | -1.479056 | 5.83E-29 | 5.87E-26 | down |
| Cenpt | 3.672615 | 1.8768078 | 2.37E-28 | 2.29E-25 | up |
| Itgam | 4.202937 | 2.071398 | 3.8E-28 | 3.52E-25 | up |
| Folr2 | 16.90075 | 4.0790155 | 5.03E-28 | 4.48E-25 | up |
| Pclaf | 6.60242 | 2.7229949 | 6.31E-28 | 5.42E-25 | up |
| Tmem176b | 3.828415 | 1.9367472 | 7.92E-28 | 6.56E-25 | up |
| Bank1 | 0.154461 | -2.694684 | 1.64E-27 | 1.31E-24 | down |
| Mmp12 | 8.192056 | 3.0342256 | 2.48E-27 | 1.91E-24 | up |
| Arhgef18 | 0.465426 | -1.103375 | 2.58E-27 | 1.93E-24 | down |
| Dpp4 | 0.368308 | -1.441015 | 3.98E-27 | 2.88E-24 | down |
| Ctsk | 6.679166 | 2.7396679 | 4.28E-27 | 3E-24 | up |
| Nipsnap3a | 3.835522 | 1.9394231 | 9.56E-27 | 6.52E-24 | up |
| ENSRNOG00000067254 | 5.10759 | 2.3526428 | 1.2E-26 | 7.92E-24 | up |
| Sapcd2 | 6.244901 | 2.6426786 | 1.76E-26 | 1.13E-23 | up |
| Fn1 | 4.312426 | 2.1084996 | 8.82E-26 | 5.53E-23 | up |

| | | | | | |
|---|---|---|---|---|---|
| Cenph | 6.126178 | 2.6149873 | 1.09E-25 | 6.67E-23 | up |
| Trem2 | 15.12842 | 3.9191895 | 2.53E-25 | 1.51E-22 | up |
| Ccn4 | 10.60617 | 3.4068321 | 2.23E-24 | 1.29E-21 | up |
| Vcan | 3.287151 | 1.7168375 | 4.12E-24 | 2.33E-21 | up |
| Tubb3 | 12.70956 | 3.6678423 | 4.98E-24 | 2.75E-21 | up |
| Ptn | 5.893584 | 2.5591453 | 5.58E-24 | 3.01E-21 | up |
| Krt76 | 0.224827 | -2.153114 | 8.41E-24 | 4.43E-21 | down |
| Col7a1 | 8.045833 | 3.0082417 | 9.14E-24 | 4.71E-21 | up |
| Rps27l | 2.632513 | 1.3964406 | 1.3E-23 | 6.41E-21 | up |
| Siglec1 | 4.313531 | 2.1088695 | 1.3E-23 | 6.41E-21 | up |
| Spc24 | 6.082151 | 2.6045817 | 2.89E-23 | 1.39E-20 | up |
| Stab1 | 8.50994 | 3.089149 | 2.94E-23 | 1.39E-20 | up |
| Matk | 9.628718 | 3.2673437 | 1.35E-22 | 6.24E-20 | up |
| Timp1 | 9.681973 | 3.275301 | 2.57E-22 | 1.17E-19 | up |
| Mafb | 5.107628 | 2.3526535 | 3.32E-22 | 1.48E-19 | up |
| Msc | 7.222708 | 2.8525399 | 4.67E-22 | 2.04E-19 | up |
| Rem1 | 5.273845 | 2.3988551 | 4.92E-22 | 2.11E-19 | up |
| Cemip | 7.33891 | 2.8755658 | 5.49E-22 | 2.31E-19 | up |
| Itpkb | 0.388422 | -1.364304 | 5.89E-22 | 2.44E-19 | down |
| Troap | 6.156504 | 2.6221113 | 6.06E-22 | 2.46E-19 | up |
| Birc5 | 6.095579 | 2.6077633 | 1.13E-21 | 4.52E-19 | up |
| Otulinl | 3.048603 | 1.6081481 | 1.38E-21 | 5.42E-19 | up |
| Ms4a7 | 14.08997 | 3.8165964 | 2.68E-21 | 1.04E-18 | up |
| Cyp4b1 | 0.247522 | -2.014374 | 1.07E-20 | 4.07E-18 | down |
| Osmr | 3.574044 | 1.8375576 | 1.16E-20 | 4.35E-18 | up |
| P2ry6 | 5.432215 | 2.4415406 | 1.65E-20 | 5.96E-18 | up |
| Atg9b | 41.43365 | 5.3727311 | 1.92E-20 | 6.84E-18 | up |
| Mt2A | 12.41768 | 3.6343236 | 2.47E-20 | 8.67E-18 | up |
| Spred3 | 5.880147 | 2.5558523 | 5.84E-20 | 2.02E-17 | up |
| Pttg1 | 5.549745 | 2.4724215 | 8.3E-20 | 2.79E-17 | up |
| Prss22 | 11.55929 | 3.5309811 | 1.17E-19 | 3.87E-17 | up |
| Gckr | 0.162817 | -2.618679 | 1.72E-19 | 5.63E-17 | down |
| Serpine1 | 9.694423 | 3.277155 | 1.88E-19 | 6.05E-17 | up |
| Slamf8 | 2.937654 | 1.5546645 | 2.04E-19 | 6.49E-17 | up |
| Hjurp | 3.253497 | 1.7019912 | 2.51E-19 | 7.88E-17 | up |
| Reln | 5.004386 | 2.3231931 | 2.66E-19 | 8.21E-17 | up |
| Ska1 | 6.810414 | 2.7677426 | 2.87E-19 | 8.63E-17 | up |
| Slc28a3 | 0.386162 | -1.372723 | 3.03E-19 | 9.01E-17 | down |
| Ccnb2-ps2 | 4.291682 | 2.1015433 | 3.81E-19 | 1.12E-16 | up |
| Nr3c2 | 0.292709 | -1.772461 | 3.9E-19 | 1.13E-16 | down |
| Knstrn | 4.91693 | 2.2977579 | 4.36E-19 | 1.25E-16 | up |

| | | | | | |
|---|---|---|---|---|---|
| Gem | 3.142332 | 1.6518358 | 6.42E-19 | 1.82E-16 | up |
| Ptafr | 3.226803 | 1.6901057 | 1.15E-18 | 3.18E-16 | up |
| Cd86 | 3.212862 | 1.6838592 | 1.33E-18 | 3.64E-16 | up |
| Clec4a | 12.27905 | 3.6181274 | 1.77E-18 | 4.76E-16 | up |
| Zfp365 | 0.439208 | -1.187023 | 1.88E-18 | 5E-16 | down |
| Sirpb3 | 0.053115 | -4.234724 | 2.2E-18 | 5.8E-16 | down |
| Krt6a | 5915.272 | 12.530229 | 2.88E-18 | 7.5E-16 | up |
| Kcnn3 | 0.187315 | -2.416458 | 3.68E-18 | 9.47E-16 | down |
| Cdkn3 | 5.894583 | 2.5593897 | 4.54E-18 | 1.16E-15 | up |
| Sec14l3 | 0.227684 | -2.134893 | 4.84E-18 | 1.22E-15 | down |
| Calcb | 7.650357 | 2.9355271 | 5.18E-18 | 1.29E-15 | up |
| LOC681182 | 0.203867 | -2.2943 | 5.79E-18 | 1.41E-15 | down |
| Clec4a3 | 5.985919 | 2.5815728 | 6.27E-18 | 1.51E-15 | up |
| Mxd3 | 6.27094 | 2.6486816 | 6.45E-18 | 1.54E-15 | up |
| ENSRNOG00000064136 | 3.951069 | 1.9822432 | 6.69E-18 | 1.58E-15 | up |
| Cd163 | 19.2104 | 4.2638154 | 6.99E-18 | 1.64E-15 | up |
| Sphk1 | 6.408801 | 2.6800545 | 7.2E-18 | 1.67E-15 | up |
| LOC100359539 | 5.836868 | 2.5451945 | 7.48E-18 | 1.72E-15 | up |
| Il2ra | 4.906541 | 2.2947063 | 7.98E-18 | 1.81E-15 | up |
| Tnfaip6 | 6.750781 | 2.7550544 | 8.07E-18 | 1.82E-15 | up |
| Lif | 5.183409 | 2.3739012 | 1.06E-17 | 2.34E-15 | up |
| Ccne1 | 6.955182 | 2.7980883 | 1.27E-17 | 2.78E-15 | up |
| Antxr1 | 0.31902 | -1.648283 | 1.33E-17 | 2.89E-15 | down |
| Ms4a6a | 4.759155 | 2.2507053 | 1.38E-17 | 2.96E-15 | up |
| Ms4a12 | 14876.44 | 13.860742 | 2.28E-17 | 4.8E-15 | up |
| Uhrf1 | 3.773749 | 1.9159987 | 2.3E-17 | 4.81E-15 | up |
| Spib | 0.23575 | -2.084672 | 2.36E-17 | 4.88E-15 | down |
| Maf | 3.773181 | 1.9157815 | 2.4E-17 | 4.91E-15 | up |
| Tmem40 | 6.962491 | 2.7996036 | 2.49E-17 | 5.07E-15 | up |
| Saal1 | 4.230964 | 2.0809865 | 2.54E-17 | 5.12E-15 | up |
| Acp3 | 0.366758 | -1.4471 | 2.62E-17 | 5.23E-15 | down |
| Slamf9 | 8.477528 | 3.0836437 | 2.88E-17 | 5.71E-15 | up |
| Hba-a3 | 0.073041 | -3.775141 | 3.94E-17 | 7.75E-15 | down |
| Ptx3 | 5.934128 | 2.5690361 | 4.01E-17 | 7.81E-15 | up |
| Vipr1 | 0.205433 | -2.283259 | 4.28E-17 | 8.26E-15 | down |
| Itgb3 | 0.271592 | -1.88049 | 4.42E-17 | 8.46E-15 | down |
| ENSRNOG00000062356 | 4.49E-05 | -14.44261 | 4.61E-17 | 8.76E-15 | down |
| Cgref1 | 4.968945 | 2.3129396 | 5E-17 | 9.27E-15 | up |
| Mmrn1 | 2.853859 | 1.5129141 | 6.85E-17 | 1.26E-14 | up |
| Hbb | 0.068537 | -3.866963 | 7.18E-17 | 1.31E-14 | down |
| Slc39a14 | 2.429523 | 1.2806733 | 1.17E-16 | 2.13E-14 | up |

| | | | | | |
|---|---|---|---|---|---|
| ENSRNOG00000071081 | 21623 | 14.400279 | 1.2E-16 | 2.15E-14 | up |
| Fcrl2 | 56.15836 | 5.8114288 | 1.21E-16 | 2.16E-14 | up |
| Dio3 | 46.91624 | 5.5520153 | 1.34E-16 | 2.37E-14 | up |
| Ccna2 | 5.131708 | 2.3594392 | 1.37E-16 | 2.4E-14 | up |
| Tubb6 | 3.824263 | 1.9351816 | 1.38E-16 | 2.4E-14 | up |
| Plekhg4 | 31.15628 | 4.9614512 | 1.43E-16 | 2.47E-14 | up |
| Ube2c | 5.803691 | 2.5369708 | 1.6E-16 | 2.74E-14 | up |
| Kif20a | 3.256912 | 1.7035046 | 1.6E-16 | 2.74E-14 | up |
| Esm1 | 0.225188 | -2.1508 | 1.75E-16 | 2.95E-14 | down |
| Rab32 | 2.657809 | 1.4102372 | 1.76E-16 | 2.95E-14 | up |
| Aldh1l2 | 8.287445 | 3.0509274 | 1.93E-16 | 3.23E-14 | up |
| Ssc5d | 3.388032 | 1.7604474 | 2.36E-16 | 3.89E-14 | up |
| Tmem86a | 5.359009 | 2.4219662 | 2.37E-16 | 3.89E-14 | up |
| Tril | 0.317295 | -1.656103 | 2.43E-16 | 3.96E-14 | down |
| Ncam1 | 3.826755 | 1.9361214 | 3.4E-16 | 5.52E-14 | up |
| Myo7a | 3.299446 | 1.7222238 | 3.59E-16 | 5.75E-14 | up |
| Olfml2b | 3.990578 | 1.9965978 | 3.59E-16 | 5.75E-14 | up |
| Rcc1 | 2.16376 | 1.1135404 | 3.86E-16 | 6.13E-14 | up |
| Asgr2 | 14.91203 | 3.8984052 | 3.89E-16 | 6.13E-14 | up |
| Ddias | 4.199195 | 2.0701128 | 4.56E-16 | 7.14E-14 | up |
| Arhgap19 | 2.4313 | 1.2817277 | 4.67E-16 | 7.26E-14 | up |
| Cenpa | 5.967851 | 2.5772115 | 4.79E-16 | 7.4E-14 | up |
| Aif1 | 4.428365 | 2.1467743 | 4.91E-16 | 7.53E-14 | up |
| Tubb2b | 23953.43 | 14.547945 | 7.22E-16 | 1.09E-13 | up |
| Apobec1 | 9.910281 | 3.3089259 | 7.45E-16 | 1.11E-13 | up |
| Prnd | 10.18132 | 3.3478522 | 8.14E-16 | 1.21E-13 | up |
| Gas7 | 2.90218 | 1.5371368 | 8.56E-16 | 1.26E-13 | up |
| RGD1565367 | 3.573209 | 1.8372203 | 9.48E-16 | 1.39E-13 | up |
| Fhl2 | 3.296149 | 1.7207813 | 1.26E-15 | 1.83E-13 | up |
| Pou2f3 | 0.188491 | -2.407435 | 1.29E-15 | 1.87E-13 | down |
| Asf1b | 4.307173 | 2.1067414 | 1.47E-15 | 2.11E-13 | up |
| Cep55 | 4.854701 | 2.2793825 | 1.69E-15 | 2.42E-13 | up |
| Gpr88 | 2.80336 | 1.487157 | 1.86E-15 | 2.63E-13 | up |
| Atp2a3 | 0.376366 | -1.409792 | 2E-15 | 2.82E-13 | down |
| Ccl9 | 2.133541 | 1.0932499 | 2.13E-15 | 2.97E-13 | up |
| Scin | 0.08216 | -3.605417 | 2.4E-15 | 3.34E-13 | down |
| Ms4a4a | 3.945242 | 1.9801138 | 2.44E-15 | 3.36E-13 | up |
| Ccnb2 | 5.369319 | 2.4247392 | 2.45E-15 | 3.36E-13 | up |
| Plpp3 | 0.41182 | -1.279914 | 2.68E-15 | 3.66E-13 | down |
| Ccdc198 | 0.143079 | -2.805112 | 2.71E-15 | 3.68E-13 | down |
| Cavin2 | 0.325624 | -1.618721 | 3.24E-15 | 4.37E-13 | down |

| | | | | | |
|---|---|---|---|---|---|
| Ccdc3 | 0.302758 | -1.723764 | 3.35E-15 | 4.49E-13 | down |
| Esrrb | 0.162347 | -2.622848 | 3.48E-15 | 4.64E-13 | down |
| ENSRNOG00000069144 | 248.6953 | 7.9582356 | 3.51E-15 | 4.65E-13 | up |
| Cercam | 3.168539 | 1.6638178 | 3.76E-15 | 4.96E-13 | up |
| Recql4 | 3.269613 | 1.7091197 | 4.18E-15 | 5.48E-13 | up |
| Efemp2 | 3.269239 | 1.7089547 | 4.32E-15 | 5.62E-13 | up |
| Kif2c | 3.975075 | 1.9909819 | 4.37E-15 | 5.66E-13 | up |
| Fap | 15.06179 | 3.9128212 | 4.44E-15 | 5.72E-13 | up |
| Sat1 | 2.617393 | 1.3881308 | 4.64E-15 | 5.95E-13 | up |
| Apoe | 9.456409 | 3.2412925 | 5.08E-15 | 6.47E-13 | up |
| Mup4 | 0.000289 | -11.75695 | 5.74E-15 | 7.2E-13 | down |
| Vxn | 6.500088 | 2.7004592 | 5.74E-15 | 7.2E-13 | up |
| Bub1b | 4.271009 | 2.0945768 | 6.45E-15 | 8.01E-13 | up |
| Slc5a1 | 0.106772 | -3.227392 | 6.46E-15 | 8.01E-13 | down |
| Gfra2 | 0.421471 | -1.246494 | 6.89E-15 | 8.49E-13 | down |
| Dpyd | 0.451426 | -1.147439 | 7.06E-15 | 8.65E-13 | down |
| Bmal2 | 33.7423 | 5.0764866 | 7.09E-15 | 8.65E-13 | up |
| Fcgr2b | 4.420555 | 2.1442274 | 7.67E-15 | 9.31E-13 | up |
| Mnda | 5.113476 | 2.3543044 | 8.01E-15 | 9.67E-13 | up |
| Fgf23 | 592.3928 | 9.2104102 | 8.35E-15 | 1E-12 | up |
| Spag5 | 3.926493 | 1.9732414 | 8.71E-15 | 1.04E-12 | up |
| Amdhd2 | 3.041718 | 1.6048866 | 9.69E-15 | 1.15E-12 | up |
| Osr1 | 3.162824 | 1.6612135 | 9.95E-15 | 1.18E-12 | up |
| Cdca3 | 4.393991 | 2.135532 | 1.07E-14 | 1.25E-12 | up |
| Azgp1 | 0.301377 | -1.730356 | 1.07E-14 | 1.25E-12 | down |
| Ckap2 | 4.301113 | 2.10471 | 1.1E-14 | 1.28E-12 | up |
| Cdc6 | 4.716249 | 2.23764 | 1.12E-14 | 1.3E-12 | up |
| Cpxm1 | 7.450676 | 2.8973713 | 1.16E-14 | 1.34E-12 | up |
| Plk1 | 3.611929 | 1.8527694 | 1.18E-14 | 1.35E-12 | up |
| Bcl3 | 3.172383 | 1.6655672 | 1.29E-14 | 1.47E-12 | up |
| Tbpl1 | 11219.87 | 13.453768 | 1.31E-14 | 1.49E-12 | up |
| Wdr62 | 2.645977 | 1.4038004 | 1.36E-14 | 1.53E-12 | up |
| Ppic | 2.441554 | 1.2877999 | 1.36E-14 | 1.53E-12 | up |
| Anln | 4.743293 | 2.2458891 | 1.45E-14 | 1.63E-12 | up |
| Pon3 | 0.429389 | -1.219643 | 1.52E-14 | 1.7E-12 | down |
| Cdc7 | 3.803315 | 1.9272575 | 1.58E-14 | 1.75E-12 | up |
| Cxcl10 | 9.081103 | 3.1828675 | 1.71E-14 | 1.88E-12 | up |
| Kcnk13 | 6.753055 | 2.7555403 | 1.76E-14 | 1.93E-12 | up |
| Hspa12a | 0.32433 | -1.624468 | 1.82E-14 | 1.97E-12 | down |
| Cks1b | 3.02196 | 1.5954844 | 1.88E-14 | 2.02E-12 | up |
| Erfe | 8.493431 | 3.0863474 | 1.95E-14 | 2.08E-12 | up |

| | | | | | |
|---|---|---|---|---|---|
| Alpk3 | 0.312089 | -1.679971 | 1.96E-14 | 2.08E-12 | down |
| Sh2d7 | 0.174258 | -2.520706 | 2.07E-14 | 2.19E-12 | down |
| Reck | 0.462988 | -1.110955 | 2.11E-14 | 2.22E-12 | down |
| Gpt | 0.265623 | -1.912546 | 2.14E-14 | 2.24E-12 | down |
| Nuf2 | 4.716374 | 2.2376781 | 2.2E-14 | 2.3E-12 | up |
| Hpgd | 0.261834 | -1.933275 | 2.34E-14 | 2.43E-12 | down |
| Bcat1 | 4.145071 | 2.0513968 | 2.98E-14 | 3.08E-12 | up |
| Mcm10 | 3.835999 | 1.9396025 | 3.75E-14 | 3.87E-12 | up |
| Mybl1 | 5.161073 | 2.3676711 | 4.05E-14 | 4.14E-12 | up |
| Mindy2 | 0.310736 | -1.686238 | 4.18E-14 | 4.25E-12 | down |
| Tmem256 | 2.288345 | 1.1943048 | 4.19E-14 | 4.25E-12 | up |
| Ccl12 | 28.7593 | 4.8459569 | 4.25E-14 | 4.28E-12 | up |
| Trh | 583.7213 | 9.1891359 | 4.27E-14 | 4.28E-12 | up |
| Ankrd6 | 0.477327 | -1.066949 | 4.88E-14 | 4.87E-12 | down |
| Sfn | 6.487418 | 2.6976443 | 5.06E-14 | 5.04E-12 | up |
| Kifc1 | 4.071997 | 2.0257363 | 5.96E-14 | 5.91E-12 | up |
| Chaf1a | 2.181965 | 1.1256281 | 7.09E-14 | 7E-12 | up |
| Trip13 | 4.40646 | 2.1396201 | 7.17E-14 | 7.05E-12 | up |
| Melk | 4.530279 | 2.1796 | 7.63E-14 | 7.47E-12 | up |
| Plcg2 | 0.414627 | -1.270114 | 7.7E-14 | 7.5E-12 | down |
| Tp53bp2 | 0.495029 | -1.014415 | 7.9E-14 | 7.66E-12 | down |
| Shisal1 | 7.289324 | 2.8657851 | 8.32E-14 | 8E-12 | up |
| Gpr183 | 3.864158 | 1.9501542 | 8.95E-14 | 8.57E-12 | up |
| LOC683313 | 3308.098 | 11.691786 | 8.98E-14 | 8.57E-12 | up |
| Mki67 | 3.196148 | 1.676334 | 9.1E-14 | 8.64E-12 | up |
| Cd80 | 4.890284 | 2.2899184 | 9.87E-14 | 9.34E-12 | up |
| Cenpq | 4.494348 | 2.1681117 | 1.1E-13 | 1.03E-11 | up |
| Abracl | 2.546548 | 1.348543 | 1.13E-13 | 1.06E-11 | up |
| Adamts4 | 8.909625 | 3.1553648 | 1.19E-13 | 1.12E-11 | up |
| Rtkn2 | 0.176101 | -2.505528 | 1.25E-13 | 1.17E-11 | down |
| ENSRNOG00000069750 | 23787.24 | 14.5379 | 1.26E-13 | 1.17E-11 | up |
| Rad51 | 3.797198 | 1.9249353 | 1.41E-13 | 1.3E-11 | up |
| AABR07030544.1 | 0.349655 | -1.515998 | 1.42E-13 | 1.31E-11 | down |
| Nlrp12 | 0.185263 | -2.432355 | 1.47E-13 | 1.34E-11 | down |
| Eme1 | 6.119313 | 2.6133698 | 1.57E-13 | 1.43E-11 | up |
| Smad9 | 0.346911 | -1.527363 | 1.58E-13 | 1.43E-11 | down |
| Des | 4.509737 | 2.1730432 | 1.61E-13 | 1.46E-11 | up |
| Ppp1r14b | 2.701678 | 1.4338558 | 1.65E-13 | 1.48E-11 | up |
| Prrx2 | 3.820522 | 1.9337697 | 1.66E-13 | 1.49E-11 | up |
| Tuba8 | 9.76001 | 3.2868826 | 1.72E-13 | 1.54E-11 | up |
| Gpsm2 | 2.495127 | 1.319113 | 1.73E-13 | 1.54E-11 | up |

| | | | | | |
|---|---|---|---|---|---|
| Fancd2 | 3.349298 | 1.7438589 | 1.74E-13 | 1.55E-11 | up |
| Spc25 | 3.378617 | 1.7564329 | 1.78E-13 | 1.58E-11 | up |
| Sema6b | 3.545719 | 1.8260782 | 1.88E-13 | 1.66E-11 | up |
| Capn8 | 15.25319 | 3.931039 | 1.91E-13 | 1.68E-11 | up |
| Adcy9 | 0.492372 | -1.022178 | 1.96E-13 | 1.71E-11 | down |
| Espl1 | 3.770726 | 1.9148422 | 2.02E-13 | 1.76E-11 | up |
| Cadm4 | 4.240319 | 2.0841729 | 2.04E-13 | 1.77E-11 | up |
| Enpep | 0.263729 | -1.92287 | 2.56E-13 | 2.21E-11 | down |
| Scube2 | 0.179584 | -2.477271 | 2.76E-13 | 2.38E-11 | down |
| Uchl1 | 5.485365 | 2.4555876 | 2.89E-13 | 2.48E-11 | up |
| Prss30 | 10.07148 | 3.3322033 | 3.11E-13 | 2.66E-11 | up |
| Rgs6 | 0.306883 | -1.704239 | 3.16E-13 | 2.69E-11 | down |
| Pbk | 4.763136 | 2.2519117 | 3.42E-13 | 2.9E-11 | up |
| Kctd4 | 15.5667 | 3.9603914 | 3.66E-13 | 3.08E-11 | up |
| Hepacam2 | 0.18594 | -2.427093 | 3.69E-13 | 3.1E-11 | down |
| Cxcl6 | 10.60097 | 3.4061245 | 3.92E-13 | 3.28E-11 | up |
| Iqgap3 | 3.225219 | 1.6893971 | 4.35E-13 | 3.62E-11 | up |
| Pkmyt1 | 3.169161 | 1.6641009 | 4.38E-13 | 3.64E-11 | up |
| AABR07019403.1 | 3.53953 | 1.8235577 | 4.77E-13 | 3.95E-11 | up |
| Glrx | 2.484284 | 1.3128301 | 5.18E-13 | 4.27E-11 | up |
| Flrt2 | 3.576358 | 1.8384913 | 5.24E-13 | 4.3E-11 | up |
| Tgfbr3 | 0.35847 | -1.480077 | 5.36E-13 | 4.39E-11 | down |
| Hba-a1 | 0.093682 | -3.416091 | 5.67E-13 | 4.63E-11 | down |
| Clec4a2 | 4.56317 | 2.1900365 | 6.01E-13 | 4.89E-11 | up |
| C1qtnf6 | 3.721124 | 1.8957384 | 6.05E-13 | 4.91E-11 | up |
| Tgfbi | 3.445686 | 1.7847914 | 6.41E-13 | 5.18E-11 | up |
| Cenpw | 7.000073 | 2.8073701 | 6.46E-13 | 5.2E-11 | up |
| Tnfsf10 | 0.23149 | -2.110978 | 6.5E-13 | 5.21E-11 | down |
| Tmem176a | 3.861915 | 1.9493162 | 6.71E-13 | 5.37E-11 | up |
| Inf2 | 2.63881 | 1.3998876 | 6.98E-13 | 5.56E-11 | up |
| Vgf | 13.94471 | 3.8016457 | 7.14E-13 | 5.66E-11 | up |
| Cav1 | 0.365539 | -1.451902 | 7.42E-13 | 5.87E-11 | down |
| Meis3 | 2.649721 | 1.4058403 | 8.09E-13 | 6.38E-11 | up |
| Bmpr2 | 0.296664 | -1.753096 | 8.29E-13 | 6.52E-11 | down |
| Bax | 2.528078 | 1.338041 | 8.78E-13 | 6.88E-11 | up |
| Siglec10 | 0.39037 | -1.357085 | 8.86E-13 | 6.92E-11 | down |
| Wwp1 | 0.41206 | -1.279074 | 9.12E-13 | 7.09E-11 | down |
| LOC291863 | 0.332875 | -1.586949 | 9.23E-13 | 7.16E-11 | down |
| Cxcl13 | 15.06321 | 3.9129575 | 9.74E-13 | 7.52E-11 | up |
| Htra3 | 2.705685 | 1.4359938 | 1.01E-12 | 7.73E-11 | up |
| Il24 | 10.42837 | 3.3824412 | 1.02E-12 | 7.77E-11 | up |

| | | | | | |
|---|---|---|---|---|---|
| Dbn1 | 2.61198 | 1.3851436 | 1.11E-12 | 8.49E-11 | up |
| LOC691995 | 3.832 | 1.9380976 | 1.18E-12 | 8.95E-11 | up |
| Cdca8 | 3.505463 | 1.8096049 | 1.18E-12 | 8.95E-11 | up |
| B4galnt4 | 3.445759 | 1.7848218 | 1.19E-12 | 9E-11 | up |
| Rhoc | 2.283765 | 1.1914145 | 1.2E-12 | 9E-11 | up |
| Igdcc4 | 2.772635 | 1.4712575 | 1.27E-12 | 9.53E-11 | up |
| Rrm2 | 4.525235 | 2.1779928 | 1.39E-12 | 1.04E-10 | up |
| Ticrr | 3.427248 | 1.7770506 | 1.42E-12 | 1.06E-10 | up |
| Lgals1 | 3.417921 | 1.7731192 | 1.47E-12 | 1.09E-10 | up |
| Tacc3 | 2.916085 | 1.5440329 | 1.48E-12 | 1.09E-10 | up |
| Ndc80 | 3.345588 | 1.7422596 | 1.54E-12 | 1.13E-10 | up |
| Tpx2 | 4.177274 | 2.0625616 | 1.56E-12 | 1.14E-10 | up |
| Plppr2 | 3.06907 | 1.6178017 | 1.59E-12 | 1.16E-10 | up |
| Slc15a2 | 0.173041 | -2.530814 | 1.66E-12 | 1.21E-10 | down |
| Oip5 | 4.712121 | 2.2363766 | 1.8E-12 | 1.3E-10 | up |
| Adam12 | 4.931597 | 2.3020549 | 1.89E-12 | 1.36E-10 | up |
| E2f8 | 5.045102 | 2.3348834 | 1.89E-12 | 1.36E-10 | up |
| Aoah | 0.411968 | -1.279397 | 1.94E-12 | 1.39E-10 | down |
| Upk3bl1 | 49.79026 | 5.6377916 | 1.94E-12 | 1.39E-10 | up |
| Ptprb | 0.355199 | -1.493299 | 2.03E-12 | 1.45E-10 | down |
| Numbl | 3.206244 | 1.6808844 | 2.12E-12 | 1.5E-10 | up |
| AABR07061385.2 | 25.87473 | 4.6934719 | 2.13E-12 | 1.5E-10 | up |
| Avil | 0.18473 | -2.436513 | 2.77E-12 | 1.94E-10 | down |
| Chek2 | 2.869474 | 1.5207863 | 2.89E-12 | 2.01E-10 | up |
| Twist1 | 5.36896 | 2.4246426 | 3.14E-12 | 2.18E-10 | up |
| Itprid1 | 0.226814 | -2.140418 | 3.21E-12 | 2.22E-10 | down |
| Top2a | 3.387138 | 1.7600668 | 3.22E-12 | 2.22E-10 | up |
| Fxyd2 | 5.467106 | 2.4507773 | 3.65E-12 | 2.51E-10 | up |
| Haus4 | 2.912779 | 1.5423964 | 3.72E-12 | 2.55E-10 | up |
| Bub1 | 3.36227 | 1.7494357 | 3.72E-12 | 2.55E-10 | up |
| Fmo2 | 0.274367 | -1.865822 | 3.99E-12 | 2.71E-10 | down |
| Ninj1 | 3.229453 | 1.6912897 | 4.18E-12 | 2.83E-10 | up |
| Grik5 | 2.138605 | 1.09667 | 4.4E-12 | 2.97E-10 | up |
| LOC100359539 | 5.295101 | 2.4046581 | 4.49E-12 | 3.02E-10 | up |
| Olfm5 | 5606.914 | 12.452991 | 5E-12 | 3.36E-10 | up |
| Pkib | 2.818897 | 1.4951305 | 5.03E-12 | 3.37E-10 | up |
| Ms4a14 | 92.63268 | 6.5334493 | 5.13E-12 | 3.42E-10 | up |
| Cnn1 | 3.081506 | 1.6236354 | 5.21E-12 | 3.46E-10 | up |
| Apobec3 | 2.301794 | 1.2027584 | 5.3E-12 | 3.51E-10 | up |
| Depdc1b | 3.292209 | 1.7190557 | 5.32E-12 | 3.51E-10 | up |
| Cped1 | 0.319624 | -1.645554 | 5.35E-12 | 3.53E-10 | down |

| | | | | | |
|---|---|---|---|---|---|
| Edil3 | 0.244003 | -2.035028 | 5.53E-12 | 3.63E-10 | down |
| Rgs10 | 2.921977 | 1.546945 | 6.18E-12 | 4.04E-10 | up |
| Lctl | 2053.936 | 11.004175 | 6.24E-12 | 4.06E-10 | up |
| A2m | 51.75623 | 5.6936607 | 6.44E-12 | 4.18E-10 | up |
| Cgnl1 | 0.273715 | -1.869255 | 6.53E-12 | 4.22E-10 | down |
| Adgrf5 | 0.34943 | -1.516923 | 6.56E-12 | 4.22E-10 | down |
| Rad51ap1 | 3.17622 | 1.6673109 | 7.04E-12 | 4.51E-10 | up |
| Perp | 0.461953 | -1.114182 | 7.13E-12 | 4.55E-10 | down |
| Racgap1 | 3.419289 | 1.7736965 | 7.23E-12 | 4.59E-10 | up |
| Arhgef25 | 2.781775 | 1.4760057 | 7.74E-12 | 4.9E-10 | up |
| C1s | 4.878332 | 2.2863879 | 8.02E-12 | 5.06E-10 | up |
| Adgrg3 | 3.860271 | 1.948702 | 8.05E-12 | 5.07E-10 | up |
| Ccnb1 | 4.360253 | 2.124412 | 8.19E-12 | 5.15E-10 | up |
| Tek | 0.438023 | -1.190922 | 8.3E-12 | 5.2E-10 | down |
| Rad54l | 2.772635 | 1.4712576 | 8.48E-12 | 5.3E-10 | up |
| Bmp15 | 0.088807 | -3.493187 | 9.03E-12 | 5.6E-10 | down |
| Dlgap5 | 3.166167 | 1.6627373 | 9.32E-12 | 5.76E-10 | up |
| Cadps2 | 0.410397 | -1.284907 | 9.79E-12 | 6.02E-10 | down |
| Pou2af2 | 0.220573 | -2.180673 | 9.81E-12 | 6.02E-10 | down |
| Stambpl1 | 2.3761 | 1.2485957 | 1.02E-11 | 6.24E-10 | up |
| Smtn | 2.556256 | 1.3540321 | 1.03E-11 | 6.27E-10 | up |
| Rpl22l1 | 2.120076 | 1.0841161 | 1.07E-11 | 6.52E-10 | up |
| Osbpl6 | 0.388015 | -1.365817 | 1.07E-11 | 6.52E-10 | down |
| Actg2 | 4.157744 | 2.0558009 | 1.08E-11 | 6.52E-10 | up |
| Nfatc4 | 2.561261 | 1.3568545 | 1.12E-11 | 6.73E-10 | up |
| Phyhipl | 0.207667 | -2.267658 | 1.16E-11 | 6.97E-10 | down |
| Gins2 | 3.572512 | 1.8369388 | 1.17E-11 | 7.03E-10 | up |
| Ccr5 | 5.156389 | 2.366361 | 1.24E-11 | 7.4E-10 | up |
| Pcdha5 | 0.280072 | -1.836132 | 1.26E-11 | 7.48E-10 | down |
| Fanca | 3.488958 | 1.8027962 | 1.27E-11 | 7.51E-10 | up |
| Stra6l | 4.498567 | 2.1694655 | 1.33E-11 | 7.88E-10 | up |
| Cenpe | 5.010815 | 2.3250452 | 1.44E-11 | 8.47E-10 | up |
| Insl3 | 2.808382 | 1.4897394 | 1.44E-11 | 8.47E-10 | up |
| Col1a1 | 3.26891 | 1.7088098 | 1.45E-11 | 8.47E-10 | up |
| Adamts6 | 2.990371 | 1.5803245 | 1.45E-11 | 8.47E-10 | up |
| Ccnf | 3.463433 | 1.7922028 | 1.45E-11 | 8.47E-10 | up |
| Slc41a2 | 2.499842 | 1.3218371 | 1.46E-11 | 8.5E-10 | up |
| Myo1a | 0.25952 | -1.946082 | 1.53E-11 | 8.87E-10 | down |
| Fjx1 | 2.669362 | 1.416495 | 1.57E-11 | 9.08E-10 | up |
| Pcolce | 4.113753 | 2.0404552 | 1.58E-11 | 9.15E-10 | up |
| Commd4 | 2.050673 | 1.0360972 | 1.6E-11 | 9.24E-10 | up |

| | | | | | |
|---|---|---|---|---|---|
| Fcrlb | 7.635761 | 2.932772 | 1.66E-11 | 9.51E-10 | up |
| P2ry12 | 2.908362 | 1.5402068 | 1.88E-11 | 1.07E-09 | up |
| Tead2 | 2.883997 | 1.5280696 | 1.88E-11 | 1.07E-09 | up |
| AABR07032097.1 | 6.444906 | 2.6881594 | 1.9E-11 | 1.07E-09 | up |
| Ntrk2 | 4.9534 | 2.3084192 | 1.95E-11 | 1.1E-09 | up |
| RGD1559482 | 2.994125 | 1.5821347 | 2.03E-11 | 1.14E-09 | up |
| Incenp | 2.327854 | 1.2190008 | 2.05E-11 | 1.15E-09 | up |
| Ier5l | 3.760345 | 1.910865 | 2.12E-11 | 1.18E-09 | up |
| Depdc1 | 5.55296 | 2.4732569 | 2.21E-11 | 1.23E-09 | up |
| Plk5 | 3.879862 | 1.9560052 | 2.25E-11 | 1.24E-09 | up |
| Mthfd2 | 2.705988 | 1.4361556 | 2.26E-11 | 1.25E-09 | up |
| Fbln7 | 3.732209 | 1.9000296 | 2.31E-11 | 1.27E-09 | up |
| Xkr5 | 4.038335 | 2.0137606 | 2.41E-11 | 1.32E-09 | up |
| Hemgn | 0.143624 | -2.799633 | 2.59E-11 | 1.41E-09 | down |
| Ccl2 | 12.95616 | 3.6955664 | 2.62E-11 | 1.43E-09 | up |
| Exo1 | 3.896135 | 1.9620438 | 2.62E-11 | 1.43E-09 | up |
| Lpar3 | 0.391962 | -1.351215 | 2.7E-11 | 1.47E-09 | down |
| Abcc4 | 0.430654 | -1.215399 | 2.95E-11 | 1.6E-09 | down |
| Akap12 | 0.493114 | -1.020006 | 3E-11 | 1.62E-09 | down |
| ENSRNOG00000069439 | 0.022359 | -5.482984 | 3.16E-11 | 1.7E-09 | down |
| Epb41l4b | 0.499015 | -1.002846 | 3.18E-11 | 1.71E-09 | down |
| Dscc1 | 4.134601 | 2.0477482 | 3.2E-11 | 1.71E-09 | up |
| Cenpi | 5.080363 | 2.3449315 | 3.38E-11 | 1.81E-09 | up |
| Peli2 | 0.402723 | -1.312141 | 3.49E-11 | 1.86E-09 | down |
| Nsl1 | 3.11754 | 1.6404079 | 3.6E-11 | 1.92E-09 | up |
| Sbno2 | 2.418135 | 1.2738946 | 3.69E-11 | 1.95E-09 | up |
| C4b | 6.159823 | 2.6228888 | 3.77E-11 | 1.99E-09 | up |
| Arhgef39 | 6.806102 | 2.7668289 | 3.83E-11 | 2.02E-09 | up |
| Cenpo | 2.235901 | 1.160856 | 3.85E-11 | 2.03E-09 | up |
| Areg | 7.993267 | 2.9987853 | 4.1E-11 | 2.15E-09 | up |
| Pdzrn3 | 2.242732 | 1.1652569 | 4.14E-11 | 2.16E-09 | up |
| Paox | 2.196947 | 1.1355003 | 4.16E-11 | 2.17E-09 | up |
| Cep72 | 2.988787 | 1.5795598 | 4.19E-11 | 2.18E-09 | up |
| Eno2 | 2.91163 | 1.5418269 | 4.5E-11 | 2.33E-09 | up |
| Cdca5 | 3.936862 | 1.9770461 | 4.51E-11 | 2.33E-09 | up |
| Ncapd2 | 2.345859 | 1.230116 | 4.55E-11 | 2.35E-09 | up |
| Erbb4 | 0.115957 | -3.108339 | 4.76E-11 | 2.45E-09 | down |
| Aurka | 2.594085 | 1.3752257 | 4.86E-11 | 2.49E-09 | up |
| Alas2 | 0.071379 | -3.808364 | 5.16E-11 | 2.64E-09 | down |
| Fignl1 | 3.659027 | 1.87146 | 5.21E-11 | 2.66E-09 | up |
| Batf | 3.401601 | 1.7662138 | 5.23E-11 | 2.66E-09 | up |

| | | | | | |
|---|---|---|---|---|---|
| Hmmr | 3.795421 | 1.9242598 | 5.27E-11 | 2.68E-09 | up |
| Thsd4 | 0.274092 | -1.86727 | 5.38E-11 | 2.72E-09 | down |
| Flrt3 | 0.234491 | -2.092397 | 5.48E-11 | 2.77E-09 | down |
| Proser2 | 0.445458 | -1.16664 | 5.57E-11 | 2.81E-09 | down |
| Mmp23 | 3.549674 | 1.8276864 | 5.58E-11 | 2.81E-09 | up |
| Ren | 2693.67 | 11.395358 | 5.78E-11 | 2.9E-09 | up |
| Pstpip2 | 0.396875 | -1.333243 | 5.82E-11 | 2.91E-09 | down |
| Pdzd2 | 0.245055 | -2.02882 | 5.87E-11 | 2.93E-09 | down |
| Diaph3 | 3.924997 | 1.9726914 | 6.03E-11 | 3.01E-09 | up |
| Izumo1r | 4235.812 | 12.048423 | 6.46E-11 | 3.21E-09 | up |
| Lrrn3 | 0.283122 | -1.820502 | 6.56E-11 | 3.25E-09 | down |
| Ccl21 | 4.825097 | 2.2705579 | 6.85E-11 | 3.39E-09 | up |
| Olfml2a | 0.469181 | -1.091783 | 6.97E-11 | 3.44E-09 | down |
| Gpr173 | 2.528251 | 1.3381396 | 7.29E-11 | 3.59E-09 | up |
| Sting1 | 2.457301 | 1.2970745 | 7.65E-11 | 3.75E-09 | up |
| Col5a1 | 3.040737 | 1.6044212 | 8.03E-11 | 3.93E-09 | up |
| Serpinb10 | 0.097688 | -3.355678 | 8.15E-11 | 3.98E-09 | down |
| Dpp7 | 2.497263 | 1.320348 | 8.21E-11 | 4E-09 | up |
| Tmem106a | 2.550235 | 1.3506302 | 8.41E-11 | 4.09E-09 | up |
| Tnn | 24.81823 | 4.6333285 | 8.54E-11 | 4.14E-09 | up |
| Tnc | 5.401423 | 2.4333396 | 8.58E-11 | 4.15E-09 | up |
| Lifr | 0.171862 | -2.540681 | 8.62E-11 | 4.16E-09 | down |
| Cyp4a8 | 0.047782 | -4.387394 | 8.94E-11 | 4.3E-09 | down |
| Phex | 0.412056 | -1.279087 | 9.03E-11 | 4.33E-09 | down |
| Crct1 | 22525.54 | 14.459274 | 9.66E-11 | 4.63E-09 | up |
| Tfpi2 | 5.695436 | 2.5098062 | 9.85E-11 | 4.71E-09 | up |
| Tspan13 | 0.49078 | -1.026852 | 9.99E-11 | 4.76E-09 | down |
| Mt1 | 4.814039 | 2.2672478 | 1.06E-10 | 5.03E-09 | up |
| Pdlim7 | 3.033334 | 1.6009045 | 1.07E-10 | 5.1E-09 | up |
| C11h3orf70 | 2.0407 | 1.0290639 | 1.18E-10 | 5.58E-09 | up |
| Pilrb | 0.148281 | -2.753598 | 1.21E-10 | 5.71E-09 | down |
| Htr2b | 3.482408 | 1.8000852 | 1.28E-10 | 6.05E-09 | up |
| Spata13 | 0.379953 | -1.396109 | 1.29E-10 | 6.06E-09 | down |
| Hk3 | 2.374452 | 1.2475946 | 1.3E-10 | 6.09E-09 | up |
| Aen | 2.085635 | 1.060487 | 1.32E-10 | 6.19E-09 | up |
| Lrfn4 | 2.159219 | 1.1105096 | 1.32E-10 | 6.19E-09 | up |
| Samd11 | 3.251973 | 1.7013153 | 1.42E-10 | 6.6E-09 | up |
| Syt8 | 5.650566 | 2.4983954 | 1.45E-10 | 6.68E-09 | up |
| Hipk2 | 0.220491 | -2.181208 | 1.46E-10 | 6.72E-09 | down |
| Stard8 | 0.451676 | -1.146641 | 1.49E-10 | 6.84E-09 | down |
| ENSRNOG00000069152 | 3.260125 | 1.7049275 | 1.49E-10 | 6.84E-09 | up |

| | | | | | |
|---|---|---|---|---|---|
| Ly49i7 | 7163.854 | 12.80652 | 1.5E-10 | 6.84E-09 | up |
| Selenbp1 | 0.268919 | -1.894754 | 1.51E-10 | 6.91E-09 | down |
| Cct6b | 0.228012 | -2.132821 | 1.52E-10 | 6.94E-09 | down |
| Exph5 | 0.2301 | -2.119669 | 1.55E-10 | 7.05E-09 | down |
| ENSRNOG00000068254 | 2.71576 | 1.441356 | 1.56E-10 | 7.07E-09 | up |
| Tmsb10 | 2.797915 | 1.4843523 | 1.6E-10 | 7.2E-09 | up |
| Ccl22 | 9.937947 | 3.3129479 | 1.6E-10 | 7.21E-09 | up |
| Fblim1 | 2.507889 | 1.3264734 | 1.66E-10 | 7.46E-09 | up |
| LOC120102992 | 2.230294 | 1.1572336 | 1.68E-10 | 7.55E-09 | up |
| Cdkn2a | 6.920383 | 2.7908519 | 1.79E-10 | 8E-09 | up |
| Slc4a1 | 0.232404 | -2.105294 | 1.8E-10 | 8.02E-09 | down |
| Ect2 | 3.839477 | 1.9409097 | 1.89E-10 | 8.4E-09 | up |
| Gbx2 | 2577.552 | 11.331786 | 1.96E-10 | 8.71E-09 | up |
| Cavin1 | 0.482714 | -1.050761 | 1.97E-10 | 8.72E-09 | down |
| Noxa1 | 6.405951 | 2.6794127 | 2E-10 | 8.82E-09 | up |
| Polr2h | 2.557162 | 1.3545435 | 2.05E-10 | 9.03E-09 | up |
| Gnai1 | 0.346911 | -1.527363 | 2.2E-10 | 9.65E-09 | down |
| Zc3h12a | 2.397506 | 1.2615346 | 2.21E-10 | 9.67E-09 | up |
| Lgals5 | 0.065822 | -3.925283 | 2.27E-10 | 9.91E-09 | down |
| Nrip3 | 3.30924 | 1.7265 | 2.28E-10 | 9.94E-09 | up |
| Mfap3l | 0.31142 | -1.683065 | 2.29E-10 | 9.94E-09 | down |
| Rarb | 0.449297 | -1.15426 | 2.42E-10 | 1.05E-08 | down |
| Nkain1 | 6.628252 | 2.7286284 | 2.84E-10 | 1.22E-08 | up |
| Tm4sf19 | 3.38436 | 1.7588831 | 2.88E-10 | 1.23E-08 | up |
| Kcnmb2 | 0.173193 | -2.529548 | 2.9E-10 | 1.24E-08 | down |
| Rnft2 | 3.343688 | 1.7414402 | 2.97E-10 | 1.26E-08 | up |
| Hhipl1 | 2.482918 | 1.3120366 | 3.06E-10 | 1.3E-08 | up |
| Fam110c | 2.541178 | 1.3454975 | 3.1E-10 | 1.31E-08 | up |
| Rap1gap | 0.392883 | -1.347829 | 3.24E-10 | 1.37E-08 | down |
| Pabir2 | 0.315342 | -1.665011 | 3.41E-10 | 1.43E-08 | down |
| Slfn13 | 5.085535 | 2.3463995 | 3.75E-10 | 1.57E-08 | up |
| RT1-DMa | 2.913487 | 1.5427467 | 3.83E-10 | 1.6E-08 | up |
| Mthfd1l | 2.743223 | 1.4558721 | 3.84E-10 | 1.6E-08 | up |
| Col1a2 | 2.915589 | 1.5437873 | 3.88E-10 | 1.62E-08 | up |
| Shcbp1 | 3.749161 | 1.9065678 | 3.92E-10 | 1.63E-08 | up |
| Tgfb3 | 2.452432 | 1.294213 | 3.93E-10 | 1.63E-08 | up |
| Slc16a12 | 0.205402 | -2.283477 | 3.93E-10 | 1.63E-08 | down |
| Plaur | 3.581067 | 1.8403894 | 3.95E-10 | 1.63E-08 | up |
| Chaf1b | 2.9027 | 1.5373955 | 3.97E-10 | 1.64E-08 | up |
| Slc39a5 | 2.273423 | 1.1848659 | 4.13E-10 | 1.69E-08 | up |
| Pole2 | 3.070004 | 1.6182406 | 4.18E-10 | 1.71E-08 | up |

| | | | | | |
|---|---|---|---|---|---|
| Susd3 | 2.608473 | 1.3832056 | 4.35E-10 | 1.77E-08 | up |
| Clec14a | 0.331344 | -1.593596 | 4.37E-10 | 1.78E-08 | down |
| Misp3 | 0.418551 | -1.256525 | 4.49E-10 | 1.82E-08 | down |
| Cxcl11 | 5.600569 | 2.4855734 | 4.5E-10 | 1.82E-08 | up |
| Gins1 | 3.55299 | 1.8290336 | 4.71E-10 | 1.9E-08 | up |
| H2az1 | 2.051889 | 1.0369526 | 4.76E-10 | 1.92E-08 | up |
| Gpr153 | 2.779296 | 1.4747194 | 4.78E-10 | 1.92E-08 | up |
| Ccl7 | 13.47102 | 3.7517872 | 4.83E-10 | 1.94E-08 | up |
| Ido1 | 5689.266 | 12.474027 | 5.04E-10 | 2.02E-08 | up |
| Ccl24 | 5.962538 | 2.5759265 | 5.23E-10 | 2.09E-08 | up |
| Tmod1 | 0.428675 | -1.222042 | 5.48E-10 | 2.19E-08 | down |
| Mmp19 | 2.891866 | 1.5320005 | 5.65E-10 | 2.25E-08 | up |
| Kif15 | 2.963269 | 1.5671898 | 5.82E-10 | 2.31E-08 | up |
| Nnmt | 2.656838 | 1.4097104 | 5.88E-10 | 2.33E-08 | up |
| Bmp3 | 0.251925 | -1.988933 | 5.94E-10 | 2.35E-08 | down |
| Jph2 | 2.301975 | 1.2028724 | 6.19E-10 | 2.44E-08 | up |
| Cenpn | 3.057332 | 1.6122731 | 6.25E-10 | 2.45E-08 | up |
| Col6a4 | 0.180292 | -2.471596 | 6.27E-10 | 2.46E-08 | down |
| Cdh11 | 2.305238 | 1.2049159 | 6.34E-10 | 2.48E-08 | up |
| Vwa1 | 3.342322 | 1.7408505 | 6.7E-10 | 2.62E-08 | up |
| Tedc2 | 2.302487 | 1.2031929 | 6.87E-10 | 2.67E-08 | up |
| Ppm1m | 2.547661 | 1.3491732 | 6.87E-10 | 2.67E-08 | up |
| Aunip | 3.91959 | 1.9707026 | 7.03E-10 | 2.73E-08 | up |
| Dmxl2 | 0.484921 | -1.044179 | 7.25E-10 | 2.81E-08 | down |
| Kank4 | 0.49676 | -1.00938 | 7.3E-10 | 2.82E-08 | down |
| Ncald | 0.384301 | -1.379691 | 7.38E-10 | 2.85E-08 | down |
| Ncaph | 4.456823 | 2.1560158 | 7.78E-10 | 2.99E-08 | up |
| Tagln | 2.741289 | 1.4548542 | 8.13E-10 | 3.12E-08 | up |
| Traip | 4.575399 | 2.1938975 | 8.2E-10 | 3.14E-08 | up |
| Aopep | 2.012547 | 1.0090228 | 8.32E-10 | 3.17E-08 | up |
| Etv4 | 2.500144 | 1.3220112 | 8.39E-10 | 3.19E-08 | up |
| Kdr | 0.30124 | -1.731013 | 8.45E-10 | 3.21E-08 | down |
| Acsm5 | 0.315139 | -1.66594 | 9.34E-10 | 3.52E-08 | down |
| Ntng2 | 2.664408 | 1.4138148 | 9.52E-10 | 3.58E-08 | up |
| Arhgap11a | 2.895931 | 1.5340273 | 9.6E-10 | 3.6E-08 | up |
| LOC120093742 | 88.08734 | 6.4608629 | 1.01E-09 | 3.78E-08 | up |
| Cdt1 | 2.755596 | 1.4623646 | 1.11E-09 | 4.14E-08 | up |
| Rtn4rl2 | 8.336899 | 3.0595108 | 1.2E-09 | 4.43E-08 | up |
| Cdk1 | 4.09423 | 2.0335921 | 1.21E-09 | 4.49E-08 | up |
| Mcm5 | 3.079229 | 1.6225692 | 1.23E-09 | 4.55E-08 | up |
| Wt1 | 7.657122 | 2.9368023 | 1.28E-09 | 4.73E-08 | up |

| | | | | | |
|---|---|---|---|---|---|
| Cip2a | 4.070268 | 2.0251238 | 1.35E-09 | 4.95E-08 | up |
| Dgkg | 0.317193 | -1.656565 | 1.53E-09 | 5.6E-08 | down |
| Sct | 5.529501 | 2.4671494 | 1.57E-09 | 5.76E-08 | up |
| Cds2 | 0.453518 | -1.140767 | 1.6E-09 | 5.83E-08 | down |
| Cfd | 9.41288 | 3.2346362 | 1.61E-09 | 5.85E-08 | up |
| Wwc1 | 0.415307 | -1.267751 | 1.65E-09 | 6.02E-08 | down |
| Mmp14 | 2.508075 | 1.3265804 | 1.67E-09 | 6.07E-08 | up |
| Map3k7cl | 4.364677 | 2.1258749 | 1.68E-09 | 6.09E-08 | up |
| Arhgap31 | 0.34117 | -1.551436 | 1.8E-09 | 6.53E-08 | down |
| Btn2a2 | 2.090667 | 1.0639635 | 1.86E-09 | 6.7E-08 | up |
| Dmp1 | 18.26165 | 4.1907448 | 1.91E-09 | 6.87E-08 | up |
| Esco2 | 4.023935 | 2.0086071 | 1.99E-09 | 7.15E-08 | up |
| Adgrv1 | 0.443931 | -1.171594 | 2.08E-09 | 7.46E-08 | down |
| Bid | 2.165392 | 1.114628 | 2.12E-09 | 7.58E-08 | up |
| Epb41l5 | 0.489375 | -1.030986 | 2.16E-09 | 7.69E-08 | down |
| Sctr | 8.011493 | 3.0020711 | 2.17E-09 | 7.69E-08 | up |
| Nusap1 | 3.190396 | 1.6737353 | 2.17E-09 | 7.69E-08 | up |
| Rpa3 | 2.057826 | 1.041121 | 2.17E-09 | 7.71E-08 | up |
| LOC100909964 | 0.232872 | -2.102389 | 2.21E-09 | 7.83E-08 | down |
| Pidd1 | 2.82264 | 1.4970454 | 2.29E-09 | 8.08E-08 | up |
| C1qtnf5 | 3.089361 | 1.6273083 | 2.29E-09 | 8.08E-08 | up |
| Clic5 | 0.289035 | -1.790685 | 2.35E-09 | 8.25E-08 | down |
| Basp1 | 2.848693 | 1.5103003 | 2.41E-09 | 8.45E-08 | up |
| Cdkl5 | 0.211884 | -2.238657 | 2.46E-09 | 8.62E-08 | down |
| Clec4a1 | 3.83333 | 1.9385982 | 2.49E-09 | 8.7E-08 | up |
| Dmrtc2 | 0.373095 | -1.422385 | 2.54E-09 | 8.88E-08 | down |
| Gna14 | 0.474101 | -1.076734 | 2.59E-09 | 9.03E-08 | down |
| Ttk | 4.984194 | 2.3173601 | 2.62E-09 | 9.12E-08 | up |
| Sptbn1 | 0.395856 | -1.336952 | 2.72E-09 | 9.43E-08 | down |
| Duox2 | 9.690248 | 3.2765336 | 2.76E-09 | 9.57E-08 | up |
| ENSRNOG00000062648 | 5.337449 | 2.4161504 | 2.84E-09 | 9.78E-08 | up |
| Adcy1 | 0.186806 | -2.420387 | 2.85E-09 | 9.81E-08 | down |
| Rasd2 | 0.275162 | -1.861645 | 2.88E-09 | 9.87E-08 | down |
| Apln | 0.210395 | -2.248829 | 2.9E-09 | 9.94E-08 | down |
| Etv5 | 0.423471 | -1.239665 | 2.95E-09 | 1.01E-07 | down |
| Ptrh1 | 2.722208 | 1.4447773 | 3.2E-09 | 1.09E-07 | up |
| Pole | 2.538042 | 1.3437159 | 3.36E-09 | 1.13E-07 | up |
| Cpt1c | 2.683495 | 1.424113 | 3.41E-09 | 1.15E-07 | up |
| Adamts3 | 3.320352 | 1.7313363 | 3.47E-09 | 1.17E-07 | up |
| Ninl | 3.850726 | 1.9451304 | 3.47E-09 | 1.17E-07 | up |
| Cd33 | 2.506847 | 1.3258738 | 3.64E-09 | 1.22E-07 | up |

| | | | | | |
|---|---|---|---|---|---|
| Ankrd39 | 2.883574 | 1.5278583 | 3.64E-09 | 1.22E-07 | up |
| Sema3g | 0.278863 | -1.842373 | 3.71E-09 | 1.24E-07 | down |
| Tmem120a | 2.005999 | 1.0043209 | 3.88E-09 | 1.29E-07 | up |
| Pla2g10 | 14.93293 | 3.9004252 | 3.9E-09 | 1.3E-07 | up |
| Omd | 0.39762 | -1.330539 | 3.94E-09 | 1.31E-07 | down |
| Thy1 | 3.264723 | 1.7069607 | 3.95E-09 | 1.31E-07 | up |
| Plgrkt | 2.13775 | 1.0960931 | 4.01E-09 | 1.32E-07 | up |
| Tnfrsf12a | 4.321954 | 2.1116838 | 4.22E-09 | 1.4E-07 | up |
| RGD1309104 | 2.109185 | 1.0766854 | 4.48E-09 | 1.47E-07 | up |
| Aldh1a1 | 0.230587 | -2.116614 | 4.51E-09 | 1.48E-07 | down |
| Hsd11b2 | 0.359038 | -1.477794 | 4.79E-09 | 1.57E-07 | down |
| Rab13 | 2.212951 | 1.1459716 | 4.88E-09 | 1.6E-07 | up |
| LOC120094021 | 2.656504 | 1.409529 | 4.9E-09 | 1.6E-07 | up |
| Fanci | 2.574372 | 1.3642208 | 5.09E-09 | 1.66E-07 | up |
| Nipal3 | 0.462683 | -1.111904 | 5.12E-09 | 1.67E-07 | down |
| Rps28-ps1 | 2.888217 | 1.5301793 | 5.31E-09 | 1.73E-07 | up |
| Adm2 | 5.207199 | 2.3805076 | 5.39E-09 | 1.75E-07 | up |
| Itga7 | 2.56963 | 1.3615608 | 5.43E-09 | 1.76E-07 | up |
| Has2 | 15.04985 | 3.9116776 | 5.58E-09 | 1.81E-07 | up |
| Gpx8 | 2.02538 | 1.0181924 | 5.79E-09 | 1.87E-07 | up |
| L1cam | 0.458016 | -1.126531 | 5.9E-09 | 1.9E-07 | down |
| Grem1 | 10.74923 | 3.4261621 | 6.01E-09 | 1.92E-07 | up |
| Thrb | 0.377852 | -1.404106 | 6.15E-09 | 1.96E-07 | down |
| Adcyap1r1 | 0.167433 | -2.578343 | 6.58E-09 | 2.09E-07 | down |
| R3hdml | 63.55786 | 5.9899986 | 7.05E-09 | 2.23E-07 | up |
| Bmp6 | 0.391149 | -1.354211 | 7.22E-09 | 2.28E-07 | down |
| Pals1 | 0.44013 | -1.183998 | 7.65E-09 | 2.41E-07 | down |
| Batf3 | 2.816145 | 1.4937217 | 7.77E-09 | 2.44E-07 | up |
| Lepr | 0.303611 | -1.719705 | 7.82E-09 | 2.46E-07 | down |
| ENSRNOG00000070640 | 0.330199 | -1.59859 | 7.85E-09 | 2.46E-07 | down |
| ENSRNOG00000063857 | 4.196642 | 2.0692353 | 7.94E-09 | 2.49E-07 | up |
| Afap1l1 | 0.466184 | -1.101029 | 7.98E-09 | 2.5E-07 | down |
| Chst15 | 0.487291 | -1.037144 | 8.04E-09 | 2.51E-07 | down |
| Tnfsf9 | 5.116319 | 2.3551061 | 8.17E-09 | 2.55E-07 | up |
| Plekhh2 | 0.340425 | -1.55459 | 8.37E-09 | 2.6E-07 | down |
| Gpr176 | 4.240485 | 2.0842294 | 8.62E-09 | 2.67E-07 | up |
| Igfbp4 | 2.530713 | 1.3395441 | 8.7E-09 | 2.69E-07 | up |
| Tac4 | 8.929577 | 3.1585918 | 8.7E-09 | 2.69E-07 | up |
| B3gnt7 | 2.047407 | 1.0337979 | 8.72E-09 | 2.69E-07 | up |
| Sectm1a | 5.613252 | 2.4888368 | 8.76E-09 | 2.7E-07 | up |
| Slc6a4 | 0.426724 | -1.228623 | 8.81E-09 | 2.72E-07 | down |

| | | | | | |
|---|---|---|---|---|---|
| Tk1 | 5.462655 | 2.4496023 | 8.89E-09 | 2.73E-07 | up |
| Tnfrsf11a | 2.423262 | 1.2769506 | 9.03E-09 | 2.77E-07 | up |
| Cxcl14 | 0.324838 | -1.622209 | 9.09E-09 | 2.78E-07 | down |
| Slc16a9 | 0.437693 | -1.192008 | 9.15E-09 | 2.8E-07 | down |
| Tnfrsf4 | 4.605107 | 2.2032347 | 9.16E-09 | 2.8E-07 | up |
| Pld4 | 3.099276 | 1.6319314 | 9.23E-09 | 2.82E-07 | up |
| Cdc45 | 3.498766 | 1.8068462 | 9.41E-09 | 2.86E-07 | up |
| Nap1l3 | 0.418655 | -1.256165 | 9.81E-09 | 2.98E-07 | down |
| Retnlg | 4.848687 | 2.2775943 | 9.9E-09 | 3E-07 | up |
| AABR07044460.2 | 0.200018 | -2.321798 | 1.04E-08 | 3.14E-07 | down |
| Ctss | 2.609736 | 1.383904 | 1.06E-08 | 3.2E-07 | up |
| Add2 | 0.141201 | -2.82418 | 1.06E-08 | 3.2E-07 | down |
| Ceacam16 | 1448.494 | 10.500338 | 1.11E-08 | 3.33E-07 | up |
| Ripk3 | 2.378659 | 1.2501484 | 1.12E-08 | 3.34E-07 | up |
| Spata20 | 12.63738 | 3.6596258 | 1.14E-08 | 3.4E-07 | up |
| Mmp7 | 60.4555 | 5.9178016 | 1.15E-08 | 3.43E-07 | up |
| Fen1 | 2.353271 | 1.2346675 | 1.17E-08 | 3.49E-07 | up |
| Plcd4 | 3.458592 | 1.7901848 | 1.19E-08 | 3.55E-07 | up |
| Sertad4 | 3.806746 | 1.9285583 | 1.2E-08 | 3.55E-07 | up |
| Limch1 | 0.263166 | -1.925955 | 1.2E-08 | 3.57E-07 | down |
| LOC120098812 | 3.759676 | 1.9106085 | 1.25E-08 | 3.69E-07 | up |
| Smad6 | 0.344969 | -1.535461 | 1.29E-08 | 3.81E-07 | down |
| Nlgn2 | 3.060228 | 1.6136389 | 1.29E-08 | 3.81E-07 | up |
| Slc2a5 | 8.409183 | 3.0719657 | 1.31E-08 | 3.87E-07 | up |
| Celf3 | 4.542772 | 2.1835728 | 1.32E-08 | 3.9E-07 | up |
| Mcm4 | 2.210161 | 1.1441512 | 1.35E-08 | 3.97E-07 | up |
| Ank2 | 0.296163 | -1.755537 | 1.37E-08 | 4.03E-07 | down |
| C1r | 2.572073 | 1.3629318 | 1.38E-08 | 4.04E-07 | up |
| Pdlim4 | 3.878146 | 1.9553672 | 1.39E-08 | 4.06E-07 | up |
| Sgo2 | 4.775235 | 2.2555718 | 1.44E-08 | 4.19E-07 | up |
| Gpnmb | 2.92076 | 1.5463438 | 1.44E-08 | 4.2E-07 | up |
| Nexmif | 0.405593 | -1.301894 | 1.45E-08 | 4.2E-07 | down |
| Ebf4 | 2.343267 | 1.2285214 | 1.45E-08 | 4.2E-07 | up |
| Rnase2 | 20.62709 | 4.3664681 | 1.45E-08 | 4.2E-07 | up |
| Abcb4 | 2.433205 | 1.282858 | 1.45E-08 | 4.2E-07 | up |
| Kif18b | 3.44768 | 1.7856261 | 1.5E-08 | 4.33E-07 | up |
| Slpi | 6.566682 | 2.7151646 | 1.54E-08 | 4.43E-07 | up |
| Clstn2 | 0.283221 | -1.82 | 1.55E-08 | 4.46E-07 | down |
| Cyth4 | 3.177951 | 1.6680969 | 1.56E-08 | 4.48E-07 | up |
| Micall2 | 2.262075 | 1.1776467 | 1.58E-08 | 4.53E-07 | up |
| Gnat3 | 0.249104 | -2.005179 | 1.61E-08 | 4.61E-07 | down |

| | | | | | |
|---|---|---|---|---|---|
| Rab3il1 | 2.836002 | 1.5038585 | 1.72E-08 | 4.9E-07 | up |
| Ppp1r1b | 0.308035 | -1.698834 | 1.84E-08 | 5.24E-07 | down |
| Klra5 | 19.28282 | 4.2692442 | 1.86E-08 | 5.28E-07 | up |
| Steap3 | 3.020412 | 1.5947455 | 1.87E-08 | 5.3E-07 | up |
| Tpm2 | 2.731159 | 1.4495131 | 1.9E-08 | 5.36E-07 | up |
| Lilrb4 | 6.578961 | 2.7178598 | 1.95E-08 | 5.49E-07 | up |
| Tac1 | 0.000372 | -11.39326 | 1.99E-08 | 5.61E-07 | down |
| ENSRNOG00000068856 | 0.493186 | -1.019795 | 2.06E-08 | 5.8E-07 | down |
| Pask | 2.116196 | 1.0814734 | 2.1E-08 | 5.89E-07 | up |
| Sema6a | 0.419125 | -1.254546 | 2.14E-08 | 5.99E-07 | down |
| RGD1307182 | 5.2947 | 2.4045489 | 2.18E-08 | 6.1E-07 | up |
| Qprt | 3.820256 | 1.9336694 | 2.29E-08 | 6.4E-07 | up |
| Orc1 | 4.693314 | 2.2306071 | 2.34E-08 | 6.54E-07 | up |
| Mcm6 | 2.303732 | 1.2039731 | 2.38E-08 | 6.65E-07 | up |
| Speg | 2.752033 | 1.4604978 | 2.39E-08 | 6.66E-07 | up |
| Rps20-ps10 | 2.011447 | 1.0082339 | 2.5E-08 | 6.94E-07 | up |
| Cenpu | 3.085244 | 1.6253846 | 2.53E-08 | 7.02E-07 | up |
| Tcfl5 | 7.515448 | 2.909859 | 2.54E-08 | 7.03E-07 | up |
| Chadl | 0.479 | -1.061903 | 2.55E-08 | 7.06E-07 | down |
| Mfsd10 | 2.040245 | 1.0287427 | 2.61E-08 | 7.2E-07 | up |
| Amelx | 0.015314 | -6.029041 | 2.61E-08 | 7.21E-07 | down |
| Lgi1 | 0.253685 | -1.978889 | 2.7E-08 | 7.44E-07 | down |
| Tdrd5 | 601.6774 | 9.2328465 | 2.73E-08 | 7.51E-07 | up |
| F11 | 0.311451 | -1.682922 | 2.74E-08 | 7.52E-07 | down |
| Sox9 | 0.322685 | -1.6318 | 2.74E-08 | 7.52E-07 | down |
| Hells | 3.029445 | 1.5990537 | 2.75E-08 | 7.53E-07 | up |
| Tns1 | 0.32 | -1.643856 | 2.85E-08 | 7.79E-07 | down |
| Slc37a2 | 2.773027 | 1.4714618 | 2.86E-08 | 7.79E-07 | up |
| S100a10 | 2.290652 | 1.1957582 | 2.87E-08 | 7.8E-07 | up |
| C15h8orf74 | 1313.95 | 10.359695 | 2.88E-08 | 7.83E-07 | up |
| Gprin3 | 0.284958 | -1.811177 | 2.91E-08 | 7.89E-07 | down |
| Tmco5b | 1939.15 | 10.921209 | 2.92E-08 | 7.92E-07 | up |
| LOC120099015 | 0.302749 | -1.723804 | 3.02E-08 | 8.17E-07 | down |
| Slc29a3 | 2.281801 | 1.1901733 | 3.22E-08 | 8.71E-07 | up |
| Prkce | 0.295139 | -1.760531 | 3.26E-08 | 8.79E-07 | down |
| ENSRNOG00000067089 | 2.304342 | 1.2043546 | 3.32E-08 | 8.94E-07 | up |
| Ppp1r9a | 0.329974 | -1.599577 | 3.36E-08 | 9.05E-07 | down |
| Crb2 | 4.793008 | 2.2609314 | 3.39E-08 | 9.11E-07 | up |
| Cygb | 2.306537 | 1.2057286 | 3.42E-08 | 9.19E-07 | up |
| Ifi30 | 2.657829 | 1.4102484 | 3.48E-08 | 9.34E-07 | up |
| Rcn3 | 2.487119 | 1.3144757 | 3.55E-08 | 9.5E-07 | up |

| | | | | | |
|---|---|---|---|---|---|
| Fzd8 | 0.409461 | -1.288203 | 3.77E-08 | 1.01E-06 | down |
| LOC681458 | 659.8342 | 9.3659598 | 3.78E-08 | 1.01E-06 | up |
| ENSRNOG00000064142 | 2.955198 | 1.5632547 | 3.79E-08 | 1.01E-06 | up |
| Pla2g7 | 3.532938 | 1.8208686 | 3.86E-08 | 1.03E-06 | up |
| Gal3st3 | 0.308835 | -1.69509 | 3.89E-08 | 1.03E-06 | down |
| Mmp8 | 0.180041 | -2.473599 | 4.04E-08 | 1.07E-06 | down |
| Has1 | 8.890958 | 3.1523389 | 4.09E-08 | 1.08E-06 | up |
| Slc38a6 | 2.146954 | 1.1022914 | 4.1E-08 | 1.08E-06 | up |
| LOC103691165 | 3.247136 | 1.6991677 | 4.11E-08 | 1.08E-06 | up |
| Sgo1 | 4.092227 | 2.0328861 | 4.25E-08 | 1.12E-06 | up |
| Sele | 4.149834 | 2.0530536 | 4.29E-08 | 1.13E-06 | up |
| Itgal | 0.45266 | -1.143502 | 4.31E-08 | 1.13E-06 | down |
| Dnm1 | 2.991709 | 1.58097 | 4.49E-08 | 1.17E-06 | up |
| Irgc | 4.194811 | 2.0686059 | 4.54E-08 | 1.18E-06 | up |
| Bgn | 2.777984 | 1.4740381 | 4.77E-08 | 1.25E-06 | up |
| Pycr1 | 2.788901 | 1.4796965 | 4.81E-08 | 1.25E-06 | up |
| Luzp2 | 0.163447 | -2.613105 | 4.83E-08 | 1.26E-06 | down |
| ENSRNOG00000065682 | 2.160939 | 1.1116584 | 5E-08 | 1.3E-06 | up |
| Cobl | 0.481072 | -1.055675 | 5.01E-08 | 1.3E-06 | down |
| Acoxl | 0.187733 | -2.413248 | 5.07E-08 | 1.31E-06 | down |
| Pif1 | 5.342715 | 2.4175731 | 5.22E-08 | 1.35E-06 | up |
| Yipf6 | 0.460841 | -1.117659 | 5.26E-08 | 1.36E-06 | down |
| Fam222a | 0.420786 | -1.248843 | 5.27E-08 | 1.36E-06 | down |
| Tppp | 0.282261 | -1.824899 | 5.34E-08 | 1.37E-06 | down |
| Angptl4 | 4.780295 | 2.2570996 | 5.42E-08 | 1.39E-06 | up |
| Hormad2 | 5.102515 | 2.3512085 | 5.51E-08 | 1.41E-06 | up |
| Mab21l3 | 3.634656 | 1.8618189 | 5.61E-08 | 1.44E-06 | up |
| Cpvl | 0.227947 | -2.133232 | 5.69E-08 | 1.45E-06 | down |
| Dll4 | 0.264876 | -1.916609 | 5.69E-08 | 1.45E-06 | down |
| ENSRNOG00000070500 | 6.622408 | 2.7273559 | 5.9E-08 | 1.5E-06 | up |
| Kif11 | 3.141671 | 1.6515321 | 5.91E-08 | 1.5E-06 | up |
| Alox15 | 0.157958 | -2.662383 | 6.04E-08 | 1.53E-06 | down |
| Haus7 | 2.049189 | 1.0350532 | 6.31E-08 | 1.6E-06 | up |
| Kif14 | 3.435634 | 1.7805762 | 6.39E-08 | 1.61E-06 | up |
| Ppp1r16b | 0.427349 | -1.226513 | 6.47E-08 | 1.63E-06 | down |
| Bora | 2.387179 | 1.2553067 | 6.61E-08 | 1.66E-06 | up |
| ENSRNOG00000063499 | 0.29719 | -1.750544 | 6.75E-08 | 1.69E-06 | down |
| Aspm | 2.029734 | 1.0212909 | 6.87E-08 | 1.72E-06 | up |
| Cma1 | 5.757079 | 2.525337 | 6.89E-08 | 1.72E-06 | up |
| Pus7l | 2.124507 | 1.0871283 | 6.96E-08 | 1.74E-06 | up |
| Plce1 | 0.483548 | -1.04827 | 7E-08 | 1.75E-06 | down |

| | | | | | |
|---|---|---|---|---|---|
| Ccl17 | 7.731824 | 2.9508087 | 7.05E-08 | 1.76E-06 | up |
| Ly49si2 | 4.405349 | 2.1392562 | 7.06E-08 | 1.76E-06 | up |
| Olfm1 | 0.4371 | -1.193963 | 7.16E-08 | 1.78E-06 | down |
| Tspan4 | 2.331821 | 1.2214573 | 7.34E-08 | 1.82E-06 | up |
| Pirb | 5.752808 | 2.5242663 | 7.41E-08 | 1.83E-06 | up |
| Svop | 20.58277 | 4.363365 | 7.51E-08 | 1.85E-06 | up |
| Spon2 | 0.319416 | -1.646491 | 7.58E-08 | 1.87E-06 | down |
| Phlda2 | 2.631524 | 1.3958988 | 7.68E-08 | 1.89E-06 | up |
| Prkaa2 | 0.263181 | -1.92587 | 7.88E-08 | 1.93E-06 | down |
| Cacnb3 | 2.006855 | 1.0049363 | 7.89E-08 | 1.93E-06 | up |
| Srd5a2 | 60.23208 | 5.9124603 | 8.05E-08 | 1.96E-06 | up |
| P4ha3 | 2.951827 | 1.5616081 | 8.16E-08 | 1.99E-06 | up |
| Phactr2 | 0.434455 | -1.202722 | 8.5E-08 | 2.06E-06 | down |
| Lgmn | 2.129744 | 1.0906803 | 8.69E-08 | 2.11E-06 | up |
| Daam1 | 0.416595 | -1.263282 | 8.78E-08 | 2.13E-06 | down |
| Cd200r1l | 0.420594 | -1.2495 | 8.82E-08 | 2.14E-06 | down |
| Wdr4 | 2.156841 | 1.1089197 | 9.25E-08 | 2.23E-06 | up |
| Serpinb9 | 0.499127 | -1.002522 | 9.5E-08 | 2.28E-06 | down |
| Arhgap5 | 0.418498 | -1.256709 | 9.82E-08 | 2.35E-06 | down |
| Jmy | 0.420402 | -1.25016 | 1.01E-07 | 2.43E-06 | down |
| LOC680920 | 9.510404 | 3.2495067 | 1.06E-07 | 2.53E-06 | up |
| Nsmf | 2.114888 | 1.0805811 | 1.07E-07 | 2.54E-06 | up |
| Sstr4 | 0.382797 | -1.385347 | 1.07E-07 | 2.54E-06 | down |
| Jaml | 2.374656 | 1.2477185 | 1.09E-07 | 2.58E-06 | up |
| Kctd8 | 0.318264 | -1.651704 | 1.12E-07 | 2.64E-06 | down |
| Il6 | 20.52239 | 4.3591266 | 1.12E-07 | 2.64E-06 | up |
| Gsn | 0.385921 | -1.373623 | 1.12E-07 | 2.65E-06 | down |
| Fam163a | 0.25992 | -1.943861 | 1.15E-07 | 2.7E-06 | down |
| Ifi27l2b | 2.851364 | 1.5116522 | 1.15E-07 | 2.71E-06 | up |
| Klhl30 | 2.617755 | 1.3883302 | 1.16E-07 | 2.73E-06 | up |
| Cxcr3 | 2.734788 | 1.4514289 | 1.2E-07 | 2.81E-06 | up |
| Tspan7 | 0.467185 | -1.097934 | 1.22E-07 | 2.85E-06 | down |
| Cyp21a1 | 11.73113 | 3.5522707 | 1.23E-07 | 2.88E-06 | up |
| Pygo1 | 0.385275 | -1.37604 | 1.24E-07 | 2.88E-06 | down |
| Mgp | 2.973882 | 1.5723472 | 1.27E-07 | 2.95E-06 | up |
| Kcnk2 | 0.466683 | -1.099487 | 1.32E-07 | 3.07E-06 | down |
| Prdm5 | 2.200865 | 1.1380703 | 1.34E-07 | 3.11E-06 | up |
| Foxp3 | 3.774898 | 1.9164375 | 1.38E-07 | 3.19E-06 | up |
| Icam5 | 4.100784 | 2.0358998 | 1.38E-07 | 3.19E-06 | up |
| 4933403O08Rik | 0.000557 | -10.81082 | 1.43E-07 | 3.29E-06 | down |
| ENSRNOG00000070543 | 7.903243 | 2.9824448 | 1.44E-07 | 3.31E-06 | up |

| | | | | | |
|---|---|---|---|---|---|
| Hecw2 | 0.44094 | -1.181345 | 1.46E-07 | 3.35E-06 | down |
| Lum | 3.824142 | 1.9351361 | 1.48E-07 | 3.39E-06 | up |
| Bpifb2 | 14.31447 | 3.8394024 | 1.48E-07 | 3.39E-06 | up |
| Sh2d6 | 0.304612 | -1.714954 | 1.5E-07 | 3.42E-06 | down |
| Ablim3 | 0.411271 | -1.281838 | 1.51E-07 | 3.44E-06 | down |
| Icam2 | 0.496115 | -1.011255 | 1.55E-07 | 3.52E-06 | down |
| Arl11 | 2.513809 | 1.3298751 | 1.57E-07 | 3.56E-06 | up |
| Mastl | 3.265088 | 1.7071217 | 1.57E-07 | 3.56E-06 | up |
| Dctpp1 | 2.563781 | 1.3582731 | 1.59E-07 | 3.6E-06 | up |
| Rad51c | 2.148476 | 1.1033134 | 1.64E-07 | 3.71E-06 | up |
| ENSRNOG00000065206 | 7.964984 | 2.9936714 | 1.64E-07 | 3.71E-06 | up |
| Slc7a5 | 2.924783 | 1.5483296 | 1.64E-07 | 3.71E-06 | up |
| Mcm3 | 2.372759 | 1.2465655 | 1.67E-07 | 3.76E-06 | up |
| Slc43a1 | 2.424393 | 1.2776235 | 1.68E-07 | 3.77E-06 | up |
| Loxl2 | 2.176839 | 1.1222349 | 1.69E-07 | 3.79E-06 | up |
| Col23a1 | 0.478704 | -1.062795 | 1.7E-07 | 3.81E-06 | down |
| Siglec15 | 23.89683 | 4.5787473 | 1.75E-07 | 3.91E-06 | up |
| Ggh | 0.499777 | -1.000645 | 1.75E-07 | 3.91E-06 | down |
| Snx8 | 2.166287 | 1.1152244 | 1.76E-07 | 3.93E-06 | up |
| Gna15 | 2.482782 | 1.3119577 | 1.84E-07 | 4.09E-06 | up |
| Fhl3 | 2.56653 | 1.3598192 | 1.85E-07 | 4.12E-06 | up |
| Tmem270 | 47.36762 | 5.5658292 | 1.85E-07 | 4.12E-06 | up |
| Efhd1 | 0.278189 | -1.845863 | 1.87E-07 | 4.15E-06 | down |
| Zfp385a | 2.098242 | 1.0691811 | 1.89E-07 | 4.18E-06 | up |
| Zfhx4 | 0.353409 | -1.50059 | 1.89E-07 | 4.19E-06 | down |
| LOC102547700 | 2.150649 | 1.104772 | 1.9E-07 | 4.21E-06 | up |
| Hmgb2 | 2.518239 | 1.3324155 | 1.92E-07 | 4.24E-06 | up |
| Lmo1 | 3.692056 | 1.8844245 | 1.93E-07 | 4.25E-06 | up |
| Gpr34 | 5.036286 | 2.3323601 | 1.93E-07 | 4.25E-06 | up |
| Plxna2 | 0.316089 | -1.661596 | 1.97E-07 | 4.32E-06 | down |
| P2ry10 | 0.396302 | -1.335328 | 2.02E-07 | 4.43E-06 | down |
| Clec12a | 3.040276 | 1.6042023 | 2.03E-07 | 4.44E-06 | up |
| Cav2 | 0.400198 | -1.321215 | 2.05E-07 | 4.48E-06 | down |
| Tifab | 2.276438 | 1.1867783 | 2.12E-07 | 4.63E-06 | up |
| Phyhd1 | 2.112097 | 1.0786762 | 2.13E-07 | 4.63E-06 | up |
| Rbp2 | 33.4737 | 5.0649563 | 2.18E-07 | 4.74E-06 | up |
| Gsta4 | 0.460141 | -1.119853 | 2.26E-07 | 4.91E-06 | down |
| Nedd9 | 0.359122 | -1.477455 | 2.28E-07 | 4.95E-06 | down |
| Atrnl1 | 0.463074 | -1.110684 | 2.31E-07 | 5E-06 | down |
| Inha | 2.785413 | 1.4778913 | 2.33E-07 | 5.04E-06 | up |
| Cdsn | 3.646998 | 1.8667092 | 2.34E-07 | 5.05E-06 | up |

| | | | | | |
|---|---|---|---|---|---|
| Edn3 | 0.375784 | -1.412026 | 2.34E-07 | 5.05E-06 | down |
| Clec4g | 0.218811 | -2.192245 | 2.36E-07 | 5.08E-06 | down |
| LOC120093104 | 5.383515 | 2.4285484 | 2.37E-07 | 5.1E-06 | up |
| Fxyd4 | 5.850234 | 2.5484943 | 2.4E-07 | 5.16E-06 | up |
| Pstpip1 | 2.943927 | 1.557742 | 2.46E-07 | 5.28E-06 | up |
| Naaa | 0.432488 | -1.209269 | 2.47E-07 | 5.3E-06 | down |
| Mad2l1 | 2.433879 | 1.2832572 | 2.55E-07 | 5.45E-06 | up |
| Adamts17 | 0.339663 | -1.557826 | 2.56E-07 | 5.48E-06 | down |
| Anlnl1 | 3.570581 | 1.8361589 | 2.62E-07 | 5.6E-06 | up |
| Slc26a10 | 9.151357 | 3.1939856 | 2.66E-07 | 5.66E-06 | up |
| Frrs1l | 0.10102 | -3.307288 | 2.67E-07 | 5.69E-06 | down |
| Csf1 | 2.008537 | 1.0061454 | 2.68E-07 | 5.7E-06 | up |
| C7 | 2.147209 | 1.1024628 | 2.69E-07 | 5.72E-06 | up |
| Npas2 | 0.429048 | -1.220787 | 2.7E-07 | 5.74E-06 | down |
| Ube2ql1 | 7.079333 | 2.8236133 | 2.7E-07 | 5.74E-06 | up |
| Cytl1 | 0.313086 | -1.675371 | 2.71E-07 | 5.75E-06 | down |
| Cst7 | 3.369342 | 1.7524669 | 2.77E-07 | 5.87E-06 | up |
| Setbp1 | 0.415972 | -1.265441 | 2.86E-07 | 6.05E-06 | down |
| Krt82 | 1109.636 | 10.11587 | 2.87E-07 | 6.05E-06 | up |
| Wdhd1 | 2.054545 | 1.0388189 | 2.92E-07 | 6.15E-06 | up |
| Chek1 | 2.363221 | 1.2407545 | 2.93E-07 | 6.17E-06 | up |
| Cdc42bpg | 0.490663 | -1.027195 | 2.93E-07 | 6.17E-06 | down |
| Lrrc25 | 2.29118 | 1.1960905 | 2.97E-07 | 6.25E-06 | up |
| Phactr1 | 0.241736 | -2.048497 | 2.99E-07 | 6.27E-06 | down |
| Cotl1 | 2.481942 | 1.3114697 | 3E-07 | 6.29E-06 | up |
| Ror2 | 3.324099 | 1.7329632 | 3.01E-07 | 6.31E-06 | up |
| Cpxm2 | 2.246292 | 1.1675453 | 3.03E-07 | 6.35E-06 | up |
| Phgdh | 3.574145 | 1.8375982 | 3.05E-07 | 6.38E-06 | up |
| Pde8b | 0.306093 | -1.707959 | 3.07E-07 | 6.41E-06 | down |
| Gpr75 | 0.37814 | -1.403009 | 3.09E-07 | 6.44E-06 | down |
| Igfbp5 | 0.446883 | -1.162031 | 3.1E-07 | 6.45E-06 | down |
| Il3ra | 2.313316 | 1.2099621 | 3.1E-07 | 6.45E-06 | up |
| Podnl1 | 5.653773 | 2.4992138 | 3.11E-07 | 6.47E-06 | up |
| Tmem47 | 0.444372 | -1.170159 | 3.3E-07 | 6.82E-06 | down |
| Utrn | 0.356353 | -1.48862 | 3.3E-07 | 6.82E-06 | down |
| Pimreg | 7.5563 | 2.91768 | 3.31E-07 | 6.82E-06 | up |
| Arhgef26 | 0.460062 | -1.1201 | 3.38E-07 | 6.95E-06 | down |
| Camk2a | 0.310929 | -1.685342 | 3.4E-07 | 6.99E-06 | down |
| ENSRNOG00000063938 | 4625.446 | 12.175377 | 3.49E-07 | 7.17E-06 | up |
| Spdl1 | 2.208349 | 1.1429681 | 3.51E-07 | 7.19E-06 | up |
| Cdc20 | 2.939493 | 1.5555673 | 3.52E-07 | 7.22E-06 | up |

| | | | | | |
|---|---|---|---|---|---|
| Lingo1 | 0.359349 | -1.476542 | 3.55E-07 | 7.27E-06 | down |
| Pde3a | 0.368716 | -1.439417 | 3.59E-07 | 7.33E-06 | down |
| Nrxn1 | 0.257357 | -1.958158 | 3.66E-07 | 7.46E-06 | down |
| Cldn9 | 3.604703 | 1.8498804 | 3.7E-07 | 7.53E-06 | up |
| Cbx2 | 2.299316 | 1.201205 | 3.73E-07 | 7.58E-06 | up |
| Glt8d2 | 2.492325 | 1.317492 | 3.75E-07 | 7.61E-06 | up |
| Clip3 | 3.146407 | 1.6537052 | 3.78E-07 | 7.66E-06 | up |
| ENSRNOG00000067297 | 0.263244 | -1.925525 | 3.82E-07 | 7.74E-06 | down |
| Col18a1 | 2.123041 | 1.0861324 | 3.9E-07 | 7.88E-06 | up |
| Ankrd29 | 0.427848 | -1.224831 | 3.93E-07 | 7.95E-06 | down |
| Tlr2 | 2.282312 | 1.190496 | 4.1E-07 | 8.26E-06 | up |
| Ripor3 | 4.939463 | 2.3043541 | 4.15E-07 | 8.35E-06 | up |
| Gpc1 | 2.582007 | 1.3684929 | 4.15E-07 | 8.35E-06 | up |
| Pianp | 2.198974 | 1.1368308 | 4.22E-07 | 8.45E-06 | up |
| Thbs1 | 3.145565 | 1.6533191 | 4.22E-07 | 8.45E-06 | up |
| Xpnpep2 | 0.402133 | -1.314254 | 4.24E-07 | 8.48E-06 | down |
| C6 | 4.869702 | 2.2838335 | 4.33E-07 | 8.63E-06 | up |
| Itgb3bp | 2.121732 | 1.0852425 | 4.34E-07 | 8.65E-06 | up |
| Stk19 | 2.096985 | 1.0683165 | 4.67E-07 | 9.28E-06 | up |
| RGD1359290 | 2.136721 | 1.0953986 | 4.76E-07 | 9.45E-06 | up |
| LOC100359951 | 2.047842 | 1.0341047 | 4.8E-07 | 9.52E-06 | up |
| Fcgr1a | 2.450915 | 1.2933204 | 4.9E-07 | 9.68E-06 | up |
| Padi3 | 15.8263 | 3.9842517 | 4.91E-07 | 9.7E-06 | up |
| Ezh2 | 2.122969 | 1.0860835 | 5.03E-07 | 9.9E-06 | up |
| Slco4a1 | 3.055755 | 1.6115286 | 5.15E-07 | 1.01E-05 | up |
| Camk2b | 0.471861 | -1.083566 | 5.21E-07 | 1.02E-05 | down |
| Kcnn4 | 3.697869 | 1.8866943 | 5.22E-07 | 1.02E-05 | up |
| Tnfrsf9 | 5.096627 | 2.3495427 | 5.28E-07 | 1.03E-05 | up |
| Aldh6a1 | 0.42637 | -1.229821 | 5.33E-07 | 1.04E-05 | down |
| Gsta3 | 0.36498 | -1.454112 | 5.54E-07 | 1.08E-05 | down |
| Ankrd50 | 0.438147 | -1.190513 | 5.66E-07 | 1.09E-05 | down |
| Vegfd | 3.613899 | 1.8535562 | 5.7E-07 | 1.1E-05 | up |
| AABR07047835.1 | 0.287603 | -1.79785 | 5.81E-07 | 1.12E-05 | down |
| Uhmk1 | 0.394251 | -1.342814 | 5.92E-07 | 1.14E-05 | down |
| Cth | 0.471228 | -1.085504 | 6.2E-07 | 1.18E-05 | down |
| Tnip3 | 3.136804 | 1.6492956 | 6.24E-07 | 1.19E-05 | up |
| Cldn1 | 0.319696 | -1.645228 | 6.26E-07 | 1.19E-05 | down |
| Adgrg5 | 3.791113 | 1.9226216 | 6.28E-07 | 1.2E-05 | up |
| Trim47 | 2.084407 | 1.0596368 | 6.43E-07 | 1.22E-05 | up |
| Pcsk1 | 0.329753 | -1.60054 | 6.47E-07 | 1.23E-05 | down |
| Gnao1 | 0.435359 | -1.199724 | 6.47E-07 | 1.23E-05 | down |

| | | | | | |
|---|---|---|---|---|---|
| Lmnb1 | 2.164274 | 1.1138834 | 6.58E-07 | 1.25E-05 | up |
| Ankrd37 | 2.725673 | 1.4466127 | 6.97E-07 | 1.32E-05 | up |
| C4a | 3.850156 | 1.944917 | 7E-07 | 1.32E-05 | up |
| Col3a1 | 2.074721 | 1.0529172 | 7.02E-07 | 1.32E-05 | up |
| Fsbp | 2.915168 | 1.5435791 | 7.08E-07 | 1.33E-05 | up |
| Irs3 | 0.36846 | -1.440419 | 7.17E-07 | 1.34E-05 | down |
| Lyve1 | 0.413272 | -1.274837 | 7.2E-07 | 1.35E-05 | down |
| Sox7 | 0.484395 | -1.045745 | 7.26E-07 | 1.36E-05 | down |
| Magi3 | 0.424766 | -1.23526 | 7.35E-07 | 1.38E-05 | down |
| LOC689757 | 11.47206 | 3.5200529 | 7.44E-07 | 1.39E-05 | up |
| Slco1a4 | 0.336097 | -1.573048 | 7.48E-07 | 1.4E-05 | down |
| Abcd4 | 2.365415 | 1.2420933 | 7.58E-07 | 1.41E-05 | up |
| Mtmr11 | 2.614011 | 1.3862654 | 7.61E-07 | 1.42E-05 | up |
| Epas1 | 0.391022 | -1.354678 | 7.65E-07 | 1.43E-05 | down |
| Padi4 | 2.706907 | 1.4366454 | 7.88E-07 | 1.46E-05 | up |
| Cmtm7 | 2.3454 | 1.2298341 | 7.92E-07 | 1.47E-05 | up |
| Cenpk | 4.386609 | 2.133106 | 7.93E-07 | 1.47E-05 | up |
| Ednrb | 0.244895 | -2.029764 | 8.21E-07 | 1.52E-05 | down |
| Tox3 | 0.440676 | -1.182209 | 8.46E-07 | 1.56E-05 | down |
| Tpbgl | 2.275464 | 1.1861609 | 8.7E-07 | 1.6E-05 | up |
| Ctrc | 2.956276 | 1.5637811 | 8.89E-07 | 1.64E-05 | up |
| Bend7 | 0.354736 | -1.495184 | 9.33E-07 | 1.71E-05 | down |
| Crmp1 | 3.553294 | 1.8291569 | 9.39E-07 | 1.72E-05 | up |
| Tmem59l | 3.343132 | 1.7412002 | 9.41E-07 | 1.72E-05 | up |
| Nrp1 | 0.342707 | -1.544953 | 9.55E-07 | 1.75E-05 | down |
| Cers6 | 0.463489 | -1.109392 | 9.79E-07 | 1.79E-05 | down |
| Oaz3 | 72.65006 | 6.182892 | 1E-06 | 1.83E-05 | up |
| Tmem255a | 0.302061 | -1.727087 | 1.03E-06 | 1.87E-05 | down |
| Usp31 | 0.460183 | -1.119721 | 1.03E-06 | 1.88E-05 | down |
| Adgre4 | 0.218499 | -2.194304 | 1.03E-06 | 1.88E-05 | down |
| Ldb2 | 0.498537 | -1.004226 | 1.04E-06 | 1.89E-05 | down |
| Tmprss4 | 2.871421 | 1.5217647 | 1.05E-06 | 1.9E-05 | up |
| Slc8b1 | 2.309162 | 1.2073691 | 1.08E-06 | 1.95E-05 | up |
| Foxo1 | 0.47492 | -1.074243 | 1.08E-06 | 1.95E-05 | down |
| Arhgef12 | 0.480508 | -1.057367 | 1.09E-06 | 1.97E-05 | down |
| Gjb4 | 25.92226 | 4.6961198 | 1.09E-06 | 1.97E-05 | up |
| Epb41 | 0.471817 | -1.0837 | 1.1E-06 | 1.97E-05 | down |
| Cd68 | 2.385569 | 1.2543337 | 1.12E-06 | 2.01E-05 | up |
| Smim29 | 2.037833 | 1.0270356 | 1.12E-06 | 2.02E-05 | up |
| Ackr2 | 3.988665 | 1.9959058 | 1.12E-06 | 2.02E-05 | up |
| Lrrc15 | 4.50596 | 2.1718345 | 1.13E-06 | 2.03E-05 | up |

| | | | | | |
|---|---|---|---|---|---|
| Htr2c | 0.105143 | -3.249572 | 1.14E-06 | 2.04E-05 | down |
| Klhl29 | 3.240898 | 1.6963936 | 1.14E-06 | 2.04E-05 | up |
| ENSRNOG00000063200 | 11.1852 | 3.4835194 | 1.15E-06 | 2.06E-05 | up |
| LOC120097031 | 0.181858 | -2.459119 | 1.15E-06 | 2.06E-05 | down |
| Rrlt | 3.492511 | 1.8042648 | 1.16E-06 | 2.08E-05 | up |
| LOC120102993 | 2.193231 | 1.1330578 | 1.2E-06 | 2.14E-05 | up |
| Sfxn5 | 0.446624 | -1.162866 | 1.23E-06 | 2.18E-05 | down |
| Sh3rf3 | 3.500798 | 1.8076837 | 1.25E-06 | 2.21E-05 | up |
| Bmpr1a | 0.497123 | -1.008324 | 1.28E-06 | 2.25E-05 | down |
| RT1-DMb | 2.236517 | 1.1612535 | 1.31E-06 | 2.3E-05 | up |
| Chtf18 | 2.248588 | 1.1690193 | 1.31E-06 | 2.31E-05 | up |
| Dlg2 | 2.094647 | 1.0667068 | 1.32E-06 | 2.32E-05 | up |
| Znf750 | 0.389869 | -1.358937 | 1.39E-06 | 2.43E-05 | down |
| Psrc1 | 4.405068 | 2.1391642 | 1.39E-06 | 2.43E-05 | up |
| Celf4 | 9.721942 | 3.2812445 | 1.4E-06 | 2.45E-05 | up |
| Syt17 | 0.481067 | -1.055691 | 1.4E-06 | 2.45E-05 | down |
| Ercc6l | 2.817829 | 1.4945839 | 1.45E-06 | 2.52E-05 | up |
| Slc7a10 | 0.261871 | -1.933073 | 1.45E-06 | 2.52E-05 | down |
| Gnaq | 0.436172 | -1.197032 | 1.47E-06 | 2.55E-05 | down |
| LOC102549592 | 2.452585 | 1.2943032 | 1.47E-06 | 2.55E-05 | up |
| Nme2 | 2.056541 | 1.04022 | 1.49E-06 | 2.57E-05 | up |
| Pax8 | 3.719493 | 1.8951058 | 1.5E-06 | 2.59E-05 | up |
| Epha3 | 0.273274 | -1.871581 | 1.51E-06 | 2.62E-05 | down |
| Kntc1 | 2.815094 | 1.4931833 | 1.52E-06 | 2.63E-05 | up |
| Foxs1 | 3.208514 | 1.6819055 | 1.55E-06 | 2.66E-05 | up |
| RT1-Db1 | 2.485778 | 1.3136977 | 1.55E-06 | 2.67E-05 | up |
| Syt6 | 0.244256 | -2.033536 | 1.56E-06 | 2.68E-05 | down |
| Myo10 | 0.397978 | -1.329241 | 1.59E-06 | 2.73E-05 | down |
| Cyp7b1 | 2.249816 | 1.1698068 | 1.59E-06 | 2.73E-05 | up |
| Chia | 4.780155 | 2.2570575 | 1.61E-06 | 2.75E-05 | up |
| Tmcc2 | 0.372911 | -1.423096 | 1.62E-06 | 2.77E-05 | down |
| Ctsb | 2.118984 | 1.0833728 | 1.64E-06 | 2.8E-05 | up |
| Tshr | 0.478557 | -1.063237 | 1.64E-06 | 2.8E-05 | down |
| Pmf1 | 2.27291 | 1.1845404 | 1.64E-06 | 2.8E-05 | up |
| Cnr1 | 0.249272 | -2.004206 | 1.66E-06 | 2.83E-05 | down |
| AABR07066753.1 | 35.76664 | 5.1605428 | 1.67E-06 | 2.83E-05 | up |
| Sdcbp2 | 3.49837 | 1.8066828 | 1.67E-06 | 2.83E-05 | up |
| Tnfrsf19 | 0.198886 | -2.329989 | 1.67E-06 | 2.83E-05 | down |
| Donson | 2.425698 | 1.2784001 | 1.68E-06 | 2.85E-05 | up |
| Zbtb8b | 0.432747 | -1.208404 | 1.74E-06 | 2.95E-05 | down |
| Gstm7 | 0.362023 | -1.465847 | 1.74E-06 | 2.95E-05 | down |

| | | | | | |
|---|---|---|---|---|---|
| Ankle1 | 11.58568 | 3.5342705 | 1.75E-06 | 2.95E-05 | up |
| Dok2 | 3.437691 | 1.7814397 | 1.78E-06 | 3E-05 | up |
| Rmnd5a | 0.488948 | -1.032246 | 1.79E-06 | 3.01E-05 | down |
| Fmn1 | 0.454858 | -1.136513 | 1.8E-06 | 3.03E-05 | down |
| Gal3st4 | 2.432885 | 1.2826683 | 1.81E-06 | 3.04E-05 | up |
| Lmo7 | 0.478522 | -1.063343 | 1.81E-06 | 3.04E-05 | down |
| Rps20 | 2.031802 | 1.02276 | 1.82E-06 | 3.06E-05 | up |
| Tmsb10-ps6 | 2.713371 | 1.4400864 | 1.84E-06 | 3.09E-05 | up |
| Skp2 | 2.756903 | 1.4630485 | 1.87E-06 | 3.12E-05 | up |
| Tekt5 | 0.288998 | -1.790868 | 1.88E-06 | 3.13E-05 | down |
| Fam25a | 46.00812 | 5.5238166 | 1.88E-06 | 3.13E-05 | up |
| Doc2b | 0.459134 | -1.123014 | 1.9E-06 | 3.15E-05 | down |
| Prelid3a | 4.126533 | 2.0449302 | 1.9E-06 | 3.16E-05 | up |
| Clec7a | 0.499758 | -1.000698 | 1.91E-06 | 3.17E-05 | down |
| Sstr1 | 0.249436 | -2.003258 | 1.93E-06 | 3.21E-05 | down |
| C3ar1 | 3.758551 | 1.9101764 | 1.94E-06 | 3.21E-05 | up |
| Cdca7l | 2.125474 | 1.0877848 | 1.95E-06 | 3.23E-05 | up |
| Mlana | 2428.24 | 11.245695 | 1.99E-06 | 3.28E-05 | up |
| Cit | 3.287834 | 1.7171374 | 2.04E-06 | 3.35E-05 | up |
| Arhgap29 | 0.492743 | -1.021093 | 2.06E-06 | 3.37E-05 | down |
| Gpihbp1 | 0.443358 | -1.173456 | 2.06E-06 | 3.37E-05 | down |
| ENSRNOG00000067232 | 2.139856 | 1.0975139 | 2.19E-06 | 3.58E-05 | up |
| Mmp2 | 2.535019 | 1.3419964 | 2.2E-06 | 3.59E-05 | up |
| Rmi2 | 2.850379 | 1.5111538 | 2.21E-06 | 3.59E-05 | up |
| Psapl1 | 4.939924 | 2.3044889 | 2.22E-06 | 3.61E-05 | up |
| Mab21l4 | 0.486928 | -1.038218 | 2.24E-06 | 3.65E-05 | down |
| Vps9d1 | 2.148472 | 1.1033108 | 2.25E-06 | 3.66E-05 | up |
| Irak3 | 2.406535 | 1.2669573 | 2.32E-06 | 3.77E-05 | up |
| Atp5mc1 | 3.869693 | 1.9522191 | 2.33E-06 | 3.78E-05 | up |
| Cdca7 | 2.448648 | 1.2919855 | 2.37E-06 | 3.83E-05 | up |
| Fxyd5 | 2.328275 | 1.2192616 | 2.41E-06 | 3.89E-05 | up |
| ENSRNOG00000066155 | 3.134403 | 1.6481906 | 2.43E-06 | 3.92E-05 | up |
| Vil1 | 0.348249 | -1.521809 | 2.44E-06 | 3.92E-05 | down |
| Slc41a3 | 2.965564 | 1.5683063 | 2.46E-06 | 3.95E-05 | up |
| Myh3 | 83.57509 | 6.3850012 | 2.46E-06 | 3.95E-05 | up |
| Acta2 | 2.896657 | 1.5343886 | 2.47E-06 | 3.96E-05 | up |
| Reg3b | 98.50495 | 6.6221243 | 2.52E-06 | 4.04E-05 | up |
| Lrp8 | 2.658353 | 1.4105326 | 2.55E-06 | 4.07E-05 | up |
| Tcirg1 | 2.412457 | 1.270503 | 2.56E-06 | 4.08E-05 | up |
| PCOLCE2 | 0.27683 | -1.852928 | 2.58E-06 | 4.12E-05 | down |
| LOC498555 | 2.104193 | 1.0732669 | 2.61E-06 | 4.15E-05 | up |

| | | | | | |
|---|---|---|---|---|---|
| Tnik | 0.423365 | -1.240025 | 2.61E-06 | 4.15E-05 | down |
| AABR07004881.1 | 0.454574 | -1.137414 | 2.62E-06 | 4.17E-05 | down |
| Slco4c1 | 0.293637 | -1.767894 | 2.64E-06 | 4.19E-05 | down |
| Cpne7 | 6.45776 | 2.6910338 | 2.68E-06 | 4.25E-05 | up |
| Brca1 | 2.20093 | 1.1381135 | 2.73E-06 | 4.32E-05 | up |
| Prex2 | 0.393831 | -1.344351 | 2.78E-06 | 4.39E-05 | down |
| Mdk | 2.162557 | 1.1127382 | 2.79E-06 | 4.41E-05 | up |
| Eef1akmt4 | 2.22013 | 1.1506442 | 2.81E-06 | 4.43E-05 | up |
| Serping1 | 2.359817 | 1.2386752 | 2.84E-06 | 4.46E-05 | up |
| Slc25a23 | 0.478743 | -1.062677 | 2.84E-06 | 4.46E-05 | down |
| Klf8 | 0.298003 | -1.7466 | 2.85E-06 | 4.47E-05 | down |
| Ncmap | 2.994463 | 1.5822972 | 2.86E-06 | 4.49E-05 | up |
| Crim1 | 0.473762 | -1.077767 | 2.96E-06 | 4.63E-05 | down |
| Susd2 | 0.353411 | -1.50058 | 2.98E-06 | 4.65E-05 | down |
| Reg3a | 5697.056 | 12.476001 | 2.98E-06 | 4.65E-05 | up |
| Tmem202 | 15.25263 | 3.9309864 | 3.07E-06 | 4.78E-05 | up |
| Angptl2 | 2.065641 | 1.0465898 | 3.21E-06 | 4.99E-05 | up |
| Lcn3 | 1041.198 | 10.024029 | 3.23E-06 | 5E-05 | up |
| Slc26a11 | 2.012193 | 1.0087686 | 3.24E-06 | 5.02E-05 | up |
| Pla2g3 | 6.870319 | 2.780377 | 3.27E-06 | 5.05E-05 | up |
| Nefh | 5.165876 | 2.3690131 | 3.31E-06 | 5.11E-05 | up |
| Mms22l | 2.304095 | 1.2042004 | 3.37E-06 | 5.19E-05 | up |
| Il17c | 3052.83 | 11.575932 | 3.39E-06 | 5.21E-05 | up |
| Aif1l | 2.569164 | 1.3612992 | 3.39E-06 | 5.21E-05 | up |
| Cbfa2t3 | 0.483025 | -1.04983 | 3.39E-06 | 5.21E-05 | down |
| Kcnj10 | 0.320942 | -1.639615 | 3.41E-06 | 5.25E-05 | down |
| Kng1 | 4.124065 | 2.0440671 | 3.43E-06 | 5.26E-05 | up |
| Fer1l6 | 0.181487 | -2.462064 | 3.45E-06 | 5.3E-05 | down |
| Adgrd1 | 3.095237 | 1.6300499 | 3.49E-06 | 5.35E-05 | up |
| Tstd3 | 2.061895 | 1.0439711 | 3.5E-06 | 5.37E-05 | up |
| Tshz3 | 2.055761 | 1.0396724 | 3.52E-06 | 5.38E-05 | up |
| Shisa9 | 0.198352 | -2.333863 | 3.56E-06 | 5.43E-05 | down |
| Ube2t | 3.055826 | 1.6115626 | 3.56E-06 | 5.43E-05 | up |
| Hey1 | 0.4074 | -1.295481 | 3.59E-06 | 5.47E-05 | down |
| Birc3 | 2.313259 | 1.209927 | 3.59E-06 | 5.48E-05 | up |
| Cd8b | 3.015115 | 1.592213 | 3.63E-06 | 5.52E-05 | up |
| Cd84 | 2.883833 | 1.5279877 | 3.63E-06 | 5.52E-05 | up |
| Slurp1 | 8.710871 | 3.122817 | 3.64E-06 | 5.52E-05 | up |
| Kcnd1 | 2.495063 | 1.319076 | 3.68E-06 | 5.58E-05 | up |
| Lrp6 | 0.454656 | -1.137153 | 3.71E-06 | 5.63E-05 | down |
| Bcl2a1 | 2.084177 | 1.0594777 | 3.72E-06 | 5.64E-05 | up |

| | | | | | |
|---|---|---|---|---|---|
| Tmprss9 | 3.496308 | 1.8058322 | 3.75E-06 | 5.67E-05 | up |
| ENSRNOG00000068920 | 2.161144 | 1.1117951 | 3.78E-06 | 5.71E-05 | up |
| ENSRNOG00000064568 | 3.049555 | 1.6085988 | 3.89E-06 | 5.86E-05 | up |
| Cyp2c23 | 0.383868 | -1.381319 | 3.9E-06 | 5.88E-05 | down |
| Rps28 | 2.131903 | 1.0921418 | 3.91E-06 | 5.88E-05 | up |
| Ackr4 | 0.413394 | -1.27441 | 3.98E-06 | 5.98E-05 | down |
| Rps27a | 3.387488 | 1.7602158 | 4.08E-06 | 6.11E-05 | up |
| Grin3a | 24.95296 | 4.6411392 | 4.15E-06 | 6.21E-05 | up |
| Fbxl17 | 0.499448 | -1.001593 | 4.26E-06 | 6.35E-05 | down |
| Cnih2 | 3.21008 | 1.6826091 | 4.28E-06 | 6.38E-05 | up |
| Col6a1 | 2.634242 | 1.3973879 | 4.3E-06 | 6.4E-05 | up |
| Tead1 | 0.422742 | -1.242152 | 4.36E-06 | 6.47E-05 | down |
| Asic1 | 0.420807 | -1.24877 | 4.38E-06 | 6.48E-05 | down |
| Hhip | 0.234408 | -2.092909 | 4.38E-06 | 6.49E-05 | down |
| Igsf3 | 2.010264 | 1.0073851 | 4.4E-06 | 6.51E-05 | up |
| Diaph2 | 0.326818 | -1.613441 | 4.51E-06 | 6.64E-05 | down |
| Sapcd1 | 5.080727 | 2.3450348 | 4.56E-06 | 6.72E-05 | up |
| Gnmt | 0.357639 | -1.483422 | 4.62E-06 | 6.79E-05 | down |
| Dmkn | 2164.433 | 11.079773 | 4.64E-06 | 6.81E-05 | up |
| Efnb2 | 0.324065 | -1.625644 | 4.67E-06 | 6.84E-05 | down |
| Myl9 | 2.070832 | 1.0502105 | 4.76E-06 | 6.96E-05 | up |
| Cybrd1 | 0.475064 | -1.073805 | 4.81E-06 | 7.02E-05 | down |
| Gsta1 | 0.464935 | -1.104899 | 4.85E-06 | 7.07E-05 | down |
| Cyp19a1 | 665.4109 | 9.3781016 | 4.9E-06 | 7.11E-05 | up |
| Tspan11 | 2.688932 | 1.427033 | 4.93E-06 | 7.14E-05 | up |
| Kif20b | 3.609949 | 1.8519786 | 5.04E-06 | 7.29E-05 | up |
| AABR07046657.1 | 0.49091 | -1.02647 | 5.06E-06 | 7.3E-05 | down |
| Tnip2 | 2.195708 | 1.1346865 | 5.15E-06 | 7.43E-05 | up |
| Arhgap32 | 0.471897 | -1.083456 | 5.17E-06 | 7.45E-05 | down |
| Scn3a | 5.06546 | 2.3406932 | 5.24E-06 | 7.55E-05 | up |
| Irf5 | 2.537708 | 1.3435259 | 5.29E-06 | 7.61E-05 | up |
| Rps20-ps11 | 2.02513 | 1.0180149 | 5.4E-06 | 7.76E-05 | up |
| Mcemp1 | 0.354505 | -1.496124 | 5.44E-06 | 7.81E-05 | down |
| Hykk | 0.337609 | -1.566576 | 5.45E-06 | 7.82E-05 | down |
| Ncapg | 2.281779 | 1.1901593 | 5.48E-06 | 7.86E-05 | up |
| Gins3 | 2.454271 | 1.2952946 | 5.55E-06 | 7.94E-05 | up |
| LOC120094258 | 0.487255 | -1.037252 | 5.56E-06 | 7.95E-05 | down |
| Calcrl | 0.324278 | -1.624696 | 5.62E-06 | 8.02E-05 | down |
| Scgb1a1 | 0.274871 | -1.863172 | 5.64E-06 | 8.04E-05 | down |
| Zyg11b | 0.429411 | -1.219569 | 5.76E-06 | 8.2E-05 | down |
| Myh13 | 0.283394 | -1.819119 | 5.77E-06 | 8.21E-05 | down |

| | | | | | |
|---|---|---|---|---|---|
| Kl | 0.470728 | -1.087036 | 5.84E-06 | 8.31E-05 | down |
| Sell | 0.440322 | -1.183368 | 6E-06 | 8.5E-05 | down |
| Slc26a6 | 2.878913 | 1.5255241 | 6.05E-06 | 8.57E-05 | up |
| Plk4 | 2.548045 | 1.3493907 | 6.08E-06 | 8.6E-05 | up |
| Spink8 | 2.684838 | 1.424835 | 6.12E-06 | 8.66E-05 | up |
| Dsn1 | 3.551471 | 1.8284169 | 6.16E-06 | 8.7E-05 | up |
| Klf15 | 0.375211 | -1.414228 | 6.19E-06 | 8.74E-05 | down |
| Rasa4 | 3.093899 | 1.6294259 | 6.3E-06 | 8.86E-05 | up |
| Pkp1 | 2.287978 | 1.1940731 | 6.41E-06 | 9E-05 | up |
| ENSRNOG00000070694 | 3.560344 | 1.8320166 | 6.45E-06 | 9.04E-05 | up |
| Igfbpl1 | 0.345906 | -1.531549 | 6.46E-06 | 9.04E-05 | down |
| Serpine2 | 2.788907 | 1.4797 | 6.48E-06 | 9.07E-05 | up |
| Lpp | 0.461148 | -1.116698 | 6.57E-06 | 9.18E-05 | down |
| RGD1565058 | 2669.896 | 11.382568 | 6.72E-06 | 9.37E-05 | up |
| Cdh5 | 0.41519 | -1.268157 | 6.77E-06 | 9.43E-05 | down |
| H2ax | 2.046716 | 1.0333108 | 6.77E-06 | 9.43E-05 | up |
| ENSRNOG00000071219 | 2.644212 | 1.4028379 | 6.93E-06 | 9.63E-05 | up |
| Mfap5 | 2.20178 | 1.13867 | 6.97E-06 | 9.66E-05 | up |
| Cd14 | 2.311525 | 1.208845 | 7.12E-06 | 9.85E-05 | up |
| Ces5a | 651.082 | 9.3466954 | 7.12E-06 | 9.85E-05 | up |
| Zfp704 | 0.377895 | -1.403944 | 7.13E-06 | 9.86E-05 | down |
| Espn | 0.400932 | -1.318571 | 7.14E-06 | 9.86E-05 | down |
| Dsg1 | 15.59619 | 3.9631215 | 7.3E-06 | 0.000101 | up |
| Tnfrsf8 | 0.484807 | -1.044518 | 7.37E-06 | 0.000101 | down |
| Csprs | 0.18483 | -2.43573 | 7.38E-06 | 0.000102 | down |
| Zfp106 | 0.466393 | -1.100382 | 7.38E-06 | 0.000102 | down |
| ENSRNOG00000065195 | 2.532943 | 1.3408147 | 7.5E-06 | 0.000103 | up |
| Rrad | 2.544995 | 1.3476629 | 7.65E-06 | 0.000105 | up |
| Met | 0.450477 | -1.150473 | 7.68E-06 | 0.000105 | down |
| Plpp1 | 2.141253 | 1.0984556 | 7.87E-06 | 0.000107 | up |
| Nab2 | 2.097128 | 1.0684147 | 7.89E-06 | 0.000108 | up |
| Trip11 | 0.469891 | -1.089603 | 7.97E-06 | 0.000109 | down |
| Msh5 | 4.19983 | 2.0703311 | 8.08E-06 | 0.00011 | up |
| LOC100912041 | 3.301579 | 1.7231563 | 8.11E-06 | 0.00011 | up |
| RGD1565355 | 0.482713 | -1.050763 | 8.12E-06 | 0.00011 | down |
| Hs3st3b1 | 0.322153 | -1.634181 | 8.17E-06 | 0.000111 | down |
| Mt1-ps3 | 3.808968 | 1.9294002 | 8.28E-06 | 0.000112 | up |
| Miga1 | 0.490068 | -1.028948 | 8.32E-06 | 0.000113 | down |
| Fkbp10 | 2.368221 | 1.2438039 | 8.36E-06 | 0.000113 | up |
| Muc5ac | 4.391047 | 2.134565 | 8.79E-06 | 0.000118 | up |
| Chp2 | 4.0235 | 2.008451 | 8.88E-06 | 0.000119 | up |

| | | | | | |
|---|---|---|---|---|---|
| Rasgrf1 | 8.263337 | 3.0467245 | 8.89E-06 | 0.000119 | up |
| Meox1 | 3.403801 | 1.7671469 | 9.13E-06 | 0.000122 | up |
| Slc28a2 | 2.987503 | 1.57894 | 9.15E-06 | 0.000123 | up |
| Mchr1 | 31.77383 | 4.9897671 | 9.16E-06 | 0.000123 | up |
| ENSRNOG00000066162 | 0.350463 | -1.512664 | 9.23E-06 | 0.000123 | down |
| Pde7b | 0.366011 | -1.450041 | 9.42E-06 | 0.000126 | down |
| LOC100911413 | 4.529579 | 2.179377 | 9.47E-06 | 0.000126 | up |
| Me3 | 0.458409 | -1.125292 | 9.51E-06 | 0.000127 | down |
| Vash1 | 2.028223 | 1.0202166 | 9.53E-06 | 0.000127 | up |
| Erbin | 0.401934 | -1.314971 | 9.61E-06 | 0.000128 | down |
| Hspb7 | 2.48233 | 1.3116952 | 9.63E-06 | 0.000128 | up |
| Dip2c | 0.465627 | -1.102754 | 9.75E-06 | 0.00013 | down |
| Sp4 | 0.429481 | -1.219335 | 9.77E-06 | 0.00013 | down |
| Bmpr1b | 0.270423 | -1.88671 | 9.78E-06 | 0.00013 | down |
| Sbsn | 6.410123 | 2.680352 | 9.86E-06 | 0.00013 | up |
| ENSRNOG00000063513 | 2.77656 | 1.4732984 | 9.97E-06 | 0.000132 | up |
| Greb1 | 3.944082 | 1.9796896 | 9.99E-06 | 0.000132 | up |
| Gria3 | 2.896468 | 1.5342945 | 1.01E-05 | 0.000133 | up |
| Hoxd8 | 4.919462 | 2.2985006 | 1.02E-05 | 0.000134 | up |
| Mslnl | 2.868735 | 1.5204147 | 1.02E-05 | 0.000134 | up |
| Pabpn1 | 2.042733 | 1.0305009 | 1.09E-05 | 0.000142 | up |
| Atp10b | 0.325021 | -1.621397 | 1.1E-05 | 0.000143 | down |
| Rexo5 | 2.781604 | 1.475917 | 1.1E-05 | 0.000144 | up |
| Opcml | 0.162879 | -2.61813 | 1.12E-05 | 0.000146 | down |
| Guca2a | 5.326209 | 2.413109 | 1.14E-05 | 0.000148 | up |
| Amotl1 | 0.437135 | -1.19385 | 1.15E-05 | 0.000149 | down |
| Xcr1 | 4.705056 | 2.2342117 | 1.17E-05 | 0.000151 | up |
| Bmx | 0.423059 | -1.241071 | 1.18E-05 | 0.000152 | down |
| Npy4r | 0.355122 | -1.493611 | 1.18E-05 | 0.000152 | down |
| Pcdh12 | 0.404586 | -1.305483 | 1.19E-05 | 0.000154 | down |
| Slc7a3 | 49.23404 | 5.6215841 | 1.19E-05 | 0.000154 | up |
| Gstm2 | 0.45497 | -1.136157 | 1.2E-05 | 0.000155 | down |
| Nfe2 | 0.361161 | -1.469284 | 1.2E-05 | 0.000155 | down |
| Plekhs1 | 0.319507 | -1.646082 | 1.22E-05 | 0.000157 | down |
| Sim1 | 0.109138 | -3.19577 | 1.22E-05 | 0.000157 | down |
| Sox5 | 0.40909 | -1.289508 | 1.22E-05 | 0.000157 | down |
| Gpr137 | 2.049113 | 1.0349993 | 1.23E-05 | 0.000157 | up |
| Jph1 | 0.438784 | -1.188418 | 1.23E-05 | 0.000158 | down |
| FAM120C | 0.482501 | -1.051395 | 1.25E-05 | 0.000159 | down |
| Fcamr | 0.294223 | -1.765018 | 1.25E-05 | 0.00016 | down |
| LOC120100233 | 3.179223 | 1.6686742 | 1.25E-05 | 0.00016 | up |

| | | | | | |
|---|---|---|---|---|---|
| Rbp1 | 2.392328 | 1.2584152 | 1.26E-05 | 0.000161 | up |
| Ttbk2 | 0.421326 | -1.24699 | 1.27E-05 | 0.000162 | down |
| Fam72a | 2.823891 | 1.4976843 | 1.27E-05 | 0.000162 | up |
| Wwc3 | 0.45039 | -1.150752 | 1.28E-05 | 0.000163 | down |
| Card19 | 2.100676 | 1.0708538 | 1.29E-05 | 0.000164 | up |
| Col6a2 | 2.57656 | 1.3654463 | 1.29E-05 | 0.000164 | up |
| Xpr1 | 0.489097 | -1.031807 | 1.31E-05 | 0.000166 | down |
| Snord86 | 4.394149 | 2.1355837 | 1.31E-05 | 0.000166 | up |
| Osm | 2.754813 | 1.4619542 | 1.31E-05 | 0.000166 | up |
| Plvap | 0.301731 | -1.728666 | 1.32E-05 | 0.000167 | down |
| Fut7 | 3.797268 | 1.9249617 | 1.33E-05 | 0.000168 | up |
| Gjc2 | 2.540772 | 1.3452668 | 1.36E-05 | 0.000171 | up |
| AABR07029262.1 | 2121.838 | 11.051099 | 1.38E-05 | 0.000174 | up |
| Rps21 | 2.07462 | 1.0528469 | 1.41E-05 | 0.000176 | up |
| Lilrb3 | 29.22763 | 4.869261 | 1.41E-05 | 0.000177 | up |
| Meiob | 6.468706 | 2.6934772 | 1.42E-05 | 0.000177 | up |
| Zfp462 | 0.409921 | -1.286581 | 1.44E-05 | 0.00018 | down |
| Spred2 | 0.483243 | -1.04918 | 1.46E-05 | 0.000182 | down |
| Rgs13 | 0.428832 | -1.221515 | 1.47E-05 | 0.000183 | down |
| Cpne3 | 0.495778 | -1.012233 | 1.48E-05 | 0.000184 | down |
| Padi2 | 0.427997 | -1.224326 | 1.48E-05 | 0.000184 | down |
| Mme | 0.348779 | -1.519616 | 1.5E-05 | 0.000185 | down |
| Ugt8 | 0.355045 | -1.493924 | 1.52E-05 | 0.000188 | down |
| Cilp | 2.2958 | 1.198997 | 1.54E-05 | 0.00019 | up |
| Hmx3 | 0.291973 | -1.776094 | 1.55E-05 | 0.000191 | down |
| Cdkn1a | 2.434455 | 1.2835986 | 1.56E-05 | 0.000192 | up |
| Vash2 | 3.182153 | 1.6700032 | 1.56E-05 | 0.000192 | up |
| Itga1 | 0.315628 | -1.663702 | 1.56E-05 | 0.000192 | down |
| Socs3 | 3.387807 | 1.7603518 | 1.57E-05 | 0.000193 | up |
| Gjb3 | 2.216076 | 1.1480072 | 1.58E-05 | 0.000194 | up |
| Ppm1j | 3.680767 | 1.8800065 | 1.6E-05 | 0.000196 | up |
| Alkal1 | 3584.908 | 11.80772 | 1.62E-05 | 0.000198 | up |
| Gypa | 0.141546 | -2.82066 | 1.64E-05 | 0.000201 | down |
| Btnl9 | 6.861664 | 2.7785585 | 1.64E-05 | 0.000201 | up |
| Orc6 | 2.715047 | 1.4409773 | 1.66E-05 | 0.000203 | up |
| Raver2 | 0.415223 | -1.268042 | 1.68E-05 | 0.000205 | down |
| Stxbp6 | 0.282903 | -1.821621 | 1.68E-05 | 0.000206 | down |
| ENSRNOG00000070938 | 2.286184 | 1.1929414 | 1.69E-05 | 0.000206 | up |
| Emilin1 | 2.276156 | 1.1865996 | 1.7E-05 | 0.000207 | up |
| Ttc16 | 2.287672 | 1.19388 | 1.7E-05 | 0.000207 | up |
| ENSRNOG00000069855 | 272.796 | 8.0916787 | 1.72E-05 | 0.00021 | up |

| | | | | | |
|---|---|---|---|---|---|
| Lilrc2 | 0.372705 | -1.423894 | 1.73E-05 | 0.000211 | down |
| Pof1b | 0.408349 | -1.292125 | 1.74E-05 | 0.000212 | down |
| ENSRNOG00000068949 | 2.165016 | 1.1143776 | 1.75E-05 | 0.000212 | up |
| Polq | 3.654787 | 1.8697872 | 1.8E-05 | 0.000217 | up |
| Pirt | 2.618001 | 1.3884655 | 1.81E-05 | 0.000218 | up |
| Ccdc183 | 20.42305 | 4.3521263 | 1.81E-05 | 0.000219 | up |
| Clmn | 0.409862 | -1.286789 | 1.84E-05 | 0.000222 | down |
| Fgd2 | 2.845805 | 1.5088367 | 1.85E-05 | 0.000222 | up |
| Rnf125 | 0.4364 | -1.196276 | 1.86E-05 | 0.000224 | down |
| Pla2g2a | 10.5444 | 3.398405 | 1.86E-05 | 0.000224 | up |
| Klra2 | 2.751471 | 1.4602033 | 1.89E-05 | 0.000227 | up |
| Tmem200a | 3.438231 | 1.7816666 | 1.9E-05 | 0.000228 | up |
| Sel1l3 | 2.065069 | 1.0461898 | 1.92E-05 | 0.000229 | up |
| LOC120095690 | 3.740971 | 1.903413 | 1.93E-05 | 0.000231 | up |
| Taf1d | 2.524882 | 1.3362161 | 1.93E-05 | 0.000231 | up |
| Fndc1 | 2.680574 | 1.4225421 | 1.96E-05 | 0.000233 | up |
| hist1h2ail2 | 7.379797 | 2.8835812 | 1.96E-05 | 0.000233 | up |
| Kcnh1 | 5.339332 | 2.4166593 | 2E-05 | 0.000237 | up |
| Kcna6 | 0.346789 | -1.527871 | 2.05E-05 | 0.000242 | down |
| RGD1308065 | 7.341366 | 2.8760486 | 2.05E-05 | 0.000243 | up |
| Ckap2l | 2.558368 | 1.3552236 | 2.07E-05 | 0.000245 | up |
| Mir214 | 23.46101 | 4.5521933 | 2.12E-05 | 0.000251 | up |
| Abcd2 | 0.245121 | -2.028435 | 2.13E-05 | 0.000251 | down |
| Faxdc2 | 0.447963 | -1.15855 | 2.13E-05 | 0.000252 | down |
| ENSRNOG00000062480 | 2.430763 | 1.2814092 | 2.14E-05 | 0.000252 | up |
| Resp18 | 2.754539 | 1.4618108 | 2.14E-05 | 0.000252 | up |
| Lmbrd2 | 0.39684 | -1.33337 | 2.15E-05 | 0.000252 | down |
| Syt15 | 0.483279 | -1.049071 | 2.17E-05 | 0.000254 | down |
| Sult1a1 | 0.329368 | -1.602226 | 2.18E-05 | 0.000255 | down |
| Gcnt3 | 13.03576 | 3.7044024 | 2.19E-05 | 0.000256 | up |
| LOC100363991 | 4.946123 | 2.3062981 | 2.19E-05 | 0.000257 | up |
| Hipk3 | 0.352556 | -1.504077 | 2.23E-05 | 0.000261 | down |
| Tulp4 | 0.48023 | -1.058203 | 2.24E-05 | 0.000262 | down |
| Gfpt2 | 2.846303 | 1.5090891 | 2.25E-05 | 0.000263 | up |
| Scn5a | 2.685324 | 1.425096 | 2.25E-05 | 0.000263 | up |
| Mid1 | 0.292444 | -1.773766 | 2.28E-05 | 0.000266 | down |
| Prx | 0.311942 | -1.680649 | 2.29E-05 | 0.000266 | down |
| ENSRNOG00000065398 | 2.336786 | 1.2245259 | 2.3E-05 | 0.000268 | up |
| Grin2d | 0.49192 | -1.023504 | 2.31E-05 | 0.000268 | down |
| Pdzd4 | 2.939979 | 1.5558058 | 2.31E-05 | 0.000268 | up |
| Fcna | 2.904125 | 1.5381035 | 2.33E-05 | 0.00027 | up |

| | | | | | |
|---|---|---|---|---|---|
| LOC120100725 | 4.561603 | 2.189541 | 2.33E-05 | 0.00027 | up |
| Fbxo5 | 2.784195 | 1.4772602 | 2.35E-05 | 0.000272 | up |
| Kcnc4 | 5.314118 | 2.4098303 | 2.36E-05 | 0.000272 | up |
| Wscd1 | 0.462032 | -1.113936 | 2.36E-05 | 0.000272 | down |
| Gfod1 | 0.496367 | -1.010521 | 2.36E-05 | 0.000272 | down |
| Rhbdl2 | 2.376142 | 1.2486211 | 2.37E-05 | 0.000274 | up |
| Slc26a2 | 0.334749 | -1.578847 | 2.39E-05 | 0.000275 | down |
| Shroom4 | 0.315257 | -1.665401 | 2.39E-05 | 0.000275 | down |
| Pla2g2d | 0.406326 | -1.299291 | 2.4E-05 | 0.000276 | down |
| Dpy30 | 0.045581 | -4.455434 | 2.4E-05 | 0.000277 | down |
| Ppp2r3b | 2.797333 | 1.4840523 | 2.47E-05 | 0.000283 | up |
| Kcnd3 | 0.483461 | -1.048527 | 2.53E-05 | 0.00029 | down |
| Aspn | 2.979272 | 1.5749598 | 2.63E-05 | 0.0003 | up |
| Ace | 0.473893 | -1.077367 | 2.64E-05 | 0.000301 | down |
| AABR07021213.1 | 33.63906 | 5.0720656 | 2.67E-05 | 0.000304 | up |
| Ghrhr | 10.24532 | 3.3568934 | 2.71E-05 | 0.000308 | up |
| Rpp25 | 2.975337 | 1.5730531 | 2.71E-05 | 0.000308 | up |
| Acer2 | 0.392631 | -1.348755 | 2.72E-05 | 0.000309 | down |
| Kif22 | 8.554992 | 3.0967666 | 2.73E-05 | 0.000309 | up |
| Atp1a3 | 3.191604 | 1.6742819 | 2.73E-05 | 0.00031 | up |
| Ro60 | 0.311984 | -1.680455 | 2.76E-05 | 0.000312 | down |
| Cryba4 | 5.275765 | 2.3993803 | 2.77E-05 | 0.000312 | up |
| Tlr12 | 4.059049 | 2.0211418 | 2.8E-05 | 0.000315 | up |
| Kcnj4 | 0.389943 | -1.358663 | 2.82E-05 | 0.000317 | down |
| Kcns2 | 0.139982 | -2.836688 | 2.84E-05 | 0.000319 | down |
| Cdh24 | 3.428205 | 1.7774534 | 2.85E-05 | 0.00032 | up |
| Cers4 | 0.447061 | -1.161457 | 2.85E-05 | 0.00032 | down |
| Poln | 2.25854 | 1.1753907 | 2.85E-05 | 0.00032 | up |
| Spp1 | 17.98424 | 4.1686612 | 2.88E-05 | 0.000323 | up |
| Syp | 3.379825 | 1.7569486 | 2.94E-05 | 0.000329 | up |
| E2f7 | 2.659235 | 1.4110112 | 2.95E-05 | 0.00033 | up |
| Nebl | 0.469873 | -1.089657 | 2.96E-05 | 0.000331 | down |
| ENSRNOG00000064361 | 0.429424 | -1.219525 | 2.96E-05 | 0.000331 | down |
| LOC102550737 | 2.734337 | 1.4511912 | 3E-05 | 0.000334 | up |
| Gp6 | 0.21635 | -2.208559 | 3.02E-05 | 0.000337 | down |
| Rarres2 | 2.026373 | 1.0189 | 3.05E-05 | 0.00034 | up |
| Eda2r | 13.03967 | 3.7048354 | 3.09E-05 | 0.000344 | up |
| ENSRNOG00000067610 | 3.901483 | 1.9640226 | 3.09E-05 | 0.000344 | up |
| Mettl24 | 0.462262 | -1.113219 | 3.11E-05 | 0.000345 | down |
| ENSRNOG00000068491 | 2.599944 | 1.3784803 | 3.12E-05 | 0.000346 | up |
| Mdfi | 3.113781 | 1.6386673 | 3.16E-05 | 0.00035 | up |

| | | | | | |
|---|---|---|---|---|---|
| Ccr2 | 3.101886 | 1.6331455 | 3.16E-05 | 0.00035 | up |
| Col16a1 | 2.412422 | 1.2704825 | 3.18E-05 | 0.000352 | up |
| Aff3 | 0.455917 | -1.133157 | 3.21E-05 | 0.000354 | down |
| Mmp3 | 1489.952 | 10.54105 | 3.23E-05 | 0.000357 | up |
| Percc1 | 0.376397 | -1.409675 | 3.32E-05 | 0.000365 | down |
| Mier2 | 2.021925 | 1.0157296 | 3.33E-05 | 0.000366 | up |
| Gpr87 | 24.79093 | 4.6317404 | 3.35E-05 | 0.000368 | up |
| Cyyr1 | 0.449735 | -1.152852 | 3.38E-05 | 0.000371 | down |
| Slc7a2 | 0.30828 | -1.697684 | 3.45E-05 | 0.000377 | down |
| Wscd2 | 3.77024 | 1.9146562 | 3.45E-05 | 0.000377 | up |
| Tmem120b | 2.00086 | 1.0006203 | 3.53E-05 | 0.000384 | up |
| Akap5 | 0.318443 | -1.650893 | 3.56E-05 | 0.000387 | down |
| Hgf | 0.381029 | -1.392028 | 3.6E-05 | 0.000391 | down |
| Cenpp | 2.476241 | 1.3081518 | 3.61E-05 | 0.000393 | up |
| LOC102553726 | 1248.394 | 10.285858 | 3.61E-05 | 0.000393 | up |
| Aoc1 | 7.83239 | 2.9694526 | 3.68E-05 | 0.000399 | up |
| AABR07037995.1 | 0.482285 | -1.052044 | 3.87E-05 | 0.000418 | down |
| Cldn10 | 0.339236 | -1.559639 | 3.88E-05 | 0.000419 | down |
| ENSRNOG00000065424 | 2.070294 | 1.0498356 | 3.91E-05 | 0.000422 | up |
| Krt78 | 2.562784 | 1.3577121 | 3.95E-05 | 0.000426 | up |
| Mnd1 | 3.010353 | 1.5899328 | 3.98E-05 | 0.000429 | up |
| Sdc3 | 0.474615 | -1.075172 | 4.01E-05 | 0.000431 | down |
| AABR07021591.1 | 32.15543 | 5.0069906 | 4.02E-05 | 0.000432 | up |
| Rpl30 | 2.214415 | 1.1469258 | 4.02E-05 | 0.000432 | up |
| Snora64 | 2.956824 | 1.5640482 | 4.04E-05 | 0.000433 | up |
| Btbd7 | 0.484955 | -1.044077 | 4.04E-05 | 0.000434 | down |
| ENSRNOG00000064300 | 6.161908 | 2.6233771 | 4.05E-05 | 0.000434 | up |
| Ibsp | 33.40373 | 5.0619372 | 4.06E-05 | 0.000435 | up |
| Rnf144a | 0.486106 | -1.040657 | 4.09E-05 | 0.000438 | down |
| Dtl | 2.898851 | 1.5354812 | 4.14E-05 | 0.000443 | up |
| Ajap1 | 17.10089 | 4.0959993 | 4.32E-05 | 0.000461 | up |
| Ckap4 | 2.033129 | 1.0237015 | 4.34E-05 | 0.000462 | up |
| Fos | 2.300909 | 1.202204 | 4.37E-05 | 0.000465 | up |
| Ypel4 | 4.247655 | 2.0866667 | 4.41E-05 | 0.000469 | up |
| Ptprk | 0.43274 | -1.208429 | 4.44E-05 | 0.000471 | down |
| Tha1 | 2.002062 | 1.0014866 | 4.44E-05 | 0.000471 | up |
| Lrrc17 | 3.110326 | 1.6370656 | 4.47E-05 | 0.000474 | up |
| Wfikkn2 | 3.351973 | 1.7450104 | 4.48E-05 | 0.000475 | up |
| Nkx2-8 | 0.214388 | -2.221705 | 4.51E-05 | 0.000477 | down |
| Gucy1a2 | 0.316052 | -1.661766 | 4.51E-05 | 0.000477 | down |
| Ier3 | 2.180867 | 1.1249018 | 4.6E-05 | 0.000485 | up |

| | | | | | |
|---|---|---|---|---|---|
| Aqp5 | 0.385241 | -1.376168 | 4.62E-05 | 0.000486 | down |
| ENSRNOG00000065587 | 2.655174 | 1.4088063 | 4.64E-05 | 0.000488 | up |
| Megf9 | 0.396651 | -1.334057 | 4.67E-05 | 0.00049 | down |
| Bhlhe40 | 2.200065 | 1.1375462 | 4.67E-05 | 0.00049 | up |
| Gcgr | 3.932824 | 1.9755658 | 4.7E-05 | 0.000492 | up |
| Plekhn1 | 3.181063 | 1.669509 | 4.72E-05 | 0.000495 | up |
| Scara5 | 2.14112 | 1.0983655 | 4.75E-05 | 0.000497 | up |
| ENSRNOG00000067337 | 2.335031 | 1.2234417 | 4.8E-05 | 0.000501 | up |
| Hdac10 | 2.05527 | 1.0393276 | 4.89E-05 | 0.00051 | up |
| Enpp6 | 0.220838 | -2.178942 | 4.9E-05 | 0.00051 | down |
| Fzd4 | 0.425796 | -1.231765 | 4.98E-05 | 0.000517 | down |
| Mtus2 | 2.741739 | 1.455091 | 4.99E-05 | 0.000518 | up |
| Dph7 | 2.079383 | 1.0561555 | 5.08E-05 | 0.000526 | up |
| Elovl7 | 0.470923 | -1.086437 | 5.09E-05 | 0.000527 | down |
| Pde5a | 0.348822 | -1.519438 | 5.1E-05 | 0.000528 | down |
| Lrrc43 | 2.837251 | 1.5044938 | 5.13E-05 | 0.00053 | up |
| Adamtsl2 | 2.527663 | 1.3378044 | 5.32E-05 | 0.000548 | up |
| Foxn3 | 0.410601 | -1.28419 | 5.39E-05 | 0.000554 | down |
| Prss23 | 0.453443 | -1.141007 | 5.39E-05 | 0.000554 | down |
| Tmem119 | 2.236648 | 1.1613385 | 5.43E-05 | 0.000556 | up |
| LOC100362751 | 2.198626 | 1.1366022 | 5.48E-05 | 0.00056 | up |
| Ptpn4 | 0.443304 | -1.173633 | 5.49E-05 | 0.000562 | down |
| Sgcg | 2.261315 | 1.1771619 | 5.51E-05 | 0.000563 | up |
| Metrnl | 2.272654 | 1.1843784 | 5.53E-05 | 0.000564 | up |
| LOC102550221 | 60.49654 | 5.9187808 | 5.6E-05 | 0.000571 | up |
| Lrfn5 | 0.001334 | -9.550466 | 5.63E-05 | 0.000574 | down |
| Znrf3 | 0.480095 | -1.058609 | 5.67E-05 | 0.000577 | down |
| Shh | 0.362507 | -1.463917 | 5.67E-05 | 0.000577 | down |
| Hic1 | 2.333554 | 1.222529 | 5.72E-05 | 0.000581 | up |
| Rgs1 | 2.733435 | 1.450715 | 5.72E-05 | 0.000581 | up |
| Tmem100 | 0.33536 | -1.576215 | 5.74E-05 | 0.000583 | down |
| Gpr63 | 2.412157 | 1.2703239 | 5.74E-05 | 0.000583 | up |
| Sds | 2.870403 | 1.5212535 | 5.76E-05 | 0.000584 | up |
| Rasef | 0.396793 | -1.333542 | 5.79E-05 | 0.000586 | down |
| Bean1 | 0.360373 | -1.472437 | 5.79E-05 | 0.000586 | down |
| Sash1 | 0.499386 | -1.001773 | 5.82E-05 | 0.000588 | down |
| AABR07054614.1 | 2.732455 | 1.4501976 | 5.83E-05 | 0.000589 | up |
| Cldn4 | 2.404446 | 1.2657044 | 5.88E-05 | 0.000593 | up |
| Ucn2 | 8.262495 | 3.0465776 | 5.92E-05 | 0.000596 | up |
| ENSRNOG00000069550 | 2.655231 | 1.4088371 | 5.93E-05 | 0.000597 | up |
| Atp8a1 | 0.401614 | -1.316117 | 5.94E-05 | 0.000598 | down |

| | | | | | |
|---|---|---|---|---|---|
| Tesmin | 3.320965 | 1.7316027 | 6.09E-05 | 0.000612 | up |
| Grn | 2.070405 | 1.0499132 | 6.1E-05 | 0.000613 | up |
| LOC120093457 | 317.2776 | 8.3096019 | 6.15E-05 | 0.000616 | up |
| Cabp1 | 2.978464 | 1.5745683 | 6.17E-05 | 0.000618 | up |
| ENSRNOG00000064431 | 24.184 | 4.5959809 | 6.18E-05 | 0.000619 | up |
| Arrb1 | 0.404269 | -1.306611 | 6.19E-05 | 0.00062 | down |
| Serf1 | 2.008188 | 1.0058942 | 6.32E-05 | 0.000632 | up |
| Pard6a | 2.164641 | 1.1141277 | 6.36E-05 | 0.000635 | up |
| Sgk2 | 2.07441 | 1.0527014 | 6.57E-05 | 0.000653 | up |
| LOC497940 | 2.065269 | 1.04633 | 6.64E-05 | 0.000659 | up |
| Trim58 | 0.001815 | -9.105741 | 6.78E-05 | 0.000673 | down |
| Rnf38 | 0.453246 | -1.141633 | 6.79E-05 | 0.000674 | down |
| Klhl11 | 0.437828 | -1.191563 | 6.8E-05 | 0.000674 | down |
| Chrd | 3.158392 | 1.6591904 | 6.88E-05 | 0.000681 | up |
| Capg | 2.098021 | 1.0690293 | 6.95E-05 | 0.000687 | up |
| Faim2 | 0.404453 | -1.305957 | 7.04E-05 | 0.000695 | down |
| Zbtb20 | 0.444483 | -1.169801 | 7.1E-05 | 0.0007 | down |
| Amtn | 4.185687 | 2.0654645 | 7.15E-05 | 0.000705 | up |
| Trpc7 | 333.4306 | 8.3812426 | 7.22E-05 | 0.00071 | up |
| Dnmt3b | 2.000807 | 1.0005822 | 7.22E-05 | 0.00071 | up |
| ENSRNOG00000067151 | 2.703394 | 1.4347717 | 7.25E-05 | 0.000712 | up |
| Ppargc1a | 0.308913 | -1.694727 | 7.31E-05 | 0.000719 | down |
| Tymp | 2.595015 | 1.3757431 | 7.33E-05 | 0.00072 | up |
| Rab27b | 0.345214 | -1.534437 | 7.43E-05 | 0.000728 | down |
| Dmpk | 2.830512 | 1.5010632 | 7.45E-05 | 0.000729 | up |
| Wfdc8 | 0.234356 | -2.093225 | 7.68E-05 | 0.000749 | down |
| Ccl4 | 2.744745 | 1.4566719 | 7.73E-05 | 0.000753 | up |
| Il17b | 5.193719 | 2.376768 | 7.99E-05 | 0.000776 | up |
| Col17a1 | 0.444798 | -1.168778 | 8.02E-05 | 0.000778 | down |
| Rasgrf2 | 0.400218 | -1.321142 | 8.05E-05 | 0.00078 | down |
| Atp7a | 0.35354 | -1.500055 | 8.16E-05 | 0.00079 | down |
| Mcc | 0.497671 | -1.006735 | 8.2E-05 | 0.000794 | down |
| Snora2l2 | 2.220834 | 1.1511015 | 8.25E-05 | 0.000798 | up |
| Slc15a1 | 21.33611 | 4.4152255 | 8.29E-05 | 0.000801 | up |
| AABR07039648.1 | 2.258797 | 1.1755549 | 8.35E-05 | 0.000806 | up |
| C3h9orf50 | 3.578542 | 1.839372 | 8.42E-05 | 0.000813 | up |
| Gpr39 | 2.317429 | 1.2125254 | 8.42E-05 | 0.000813 | up |
| Plau | 2.139712 | 1.0974164 | 8.5E-05 | 0.000819 | up |
| LOC100360856 | 412.4914 | 8.6882203 | 8.67E-05 | 0.000832 | up |
| Thsd7a | 0.417933 | -1.258657 | 8.72E-05 | 0.000837 | down |
| Zbtb41 | 0.371318 | -1.429275 | 8.73E-05 | 0.000837 | down |

| | | | | | |
|---|---|---|---|---|---|
| Kcnmb1 | 2.424387 | 1.2776202 | 8.79E-05 | 0.000841 | up |
| Ddx11 | 2.241742 | 1.16462 | 8.83E-05 | 0.000844 | up |
| Rab6b | 0.396022 | -1.336348 | 8.85E-05 | 0.000846 | down |
| Efcc1 | 0.465401 | -1.103453 | 8.93E-05 | 0.000852 | down |
| Rasgef1c | 9.750407 | 3.2854624 | 8.99E-05 | 0.000857 | up |
| Rnf24 | 0.263012 | -1.926799 | 9.04E-05 | 0.000861 | down |
| Mycn | 3.833729 | 1.9387482 | 9.05E-05 | 0.000862 | up |
| Vmo1 | 3.83218 | 1.9381655 | 9.14E-05 | 0.00087 | up |
| Thbs4 | 7.130066 | 2.8339153 | 9.24E-05 | 0.000879 | up |
| Ltbp2 | 2.500092 | 1.3219815 | 9.34E-05 | 0.000887 | up |
| Egfl6 | 0.380611 | -1.393609 | 9.36E-05 | 0.000889 | down |
| ENSRNOG00000063207 | 2.014062 | 1.0101079 | 9.38E-05 | 0.00089 | up |
| Unc93b1 | 2.040078 | 1.0286245 | 9.46E-05 | 0.000895 | up |
| Kit | 0.460296 | -1.119365 | 9.51E-05 | 0.000899 | down |
| RT1-CE3 | 21.88767 | 4.4520462 | 9.51E-05 | 0.000899 | up |
| Rtl9 | 6.822372 | 2.7702735 | 9.55E-05 | 0.000901 | up |
| Tmod2 | 0.458038 | -1.12646 | 9.55E-05 | 0.000901 | down |
| Cck | 13.71286 | 3.7774571 | 9.59E-05 | 0.000904 | up |
| Mpp7 | 0.427478 | -1.226079 | 9.59E-05 | 0.000904 | down |
| Zfp513 | 2.228602 | 1.1561392 | 9.74E-05 | 0.000916 | up |
| RGD1564854 | 0.45119 | -1.148194 | 9.81E-05 | 0.000923 | down |
| ENSRNOG00000069427 | 170.5886 | 7.4143775 | 9.87E-05 | 0.000927 | up |
| Amot | 0.377634 | -1.404938 | 0.0001 | 0.000938 | down |
| Kcnt1 | 3.758064 | 1.9099897 | 0.0001 | 0.000939 | up |
| Zfp862 | 2.105055 | 1.0738578 | 0.000101 | 0.000948 | up |
| Raet1e | 1752.606 | 10.775286 | 0.000102 | 0.000952 | up |
| Cntfr | 0.471475 | -1.084748 | 0.000103 | 0.000958 | down |
| Stil | 2.586556 | 1.3710324 | 0.000103 | 0.000959 | up |
| Itga4 | 0.405572 | -1.301972 | 0.000105 | 0.000974 | down |
| Fancb | 2.571797 | 1.3627769 | 0.000106 | 0.00098 | up |
| Zfp575 | 4.631721 | 2.2115484 | 0.000106 | 0.00098 | up |
| LOC120093064 | 3.396365 | 1.7639916 | 0.000108 | 0.001001 | up |
| Scamp5 | 2.24404 | 1.1660983 | 0.00011 | 0.00102 | up |
| Ager | 0.324775 | -1.622488 | 0.000112 | 0.001032 | down |
| Tnfrsf18 | 2.566933 | 1.3600454 | 0.000112 | 0.001033 | up |
| Kcne4 | 2.372062 | 1.2461415 | 0.000112 | 0.001034 | up |
| Arid5a | 2.094648 | 1.0667078 | 0.000113 | 0.001036 | up |
| Elapor1 | 0.392916 | -1.347707 | 0.000113 | 0.001038 | down |
| Knl1 | 2.796512 | 1.4836285 | 0.000114 | 0.001047 | up |
| Doxl2 | 69.42182 | 6.1173173 | 0.000115 | 0.00106 | up |
| LOC685793 | 385.5945 | 8.5909408 | 0.000115 | 0.00106 | up |

| | | | | | |
|---|---|---|---|---|---|
| Lhx6 | 2.554825 | 1.3532246 | 0.000116 | 0.00106 | up |
| Lrrc8b | 0.496781 | -1.009319 | 0.000116 | 0.001063 | down |
| Catsperb | 3.128116 | 1.6452942 | 0.000116 | 0.001064 | up |
| Nek2l1 | 2.175897 | 1.1216102 | 0.000117 | 0.001067 | up |
| Dusp8 | 0.455952 | -1.133048 | 0.000117 | 0.001068 | down |
| ENSRNOG00000069272 | 33.66002 | 5.0729639 | 0.000117 | 0.00107 | up |
| Treml4 | 0.271736 | -1.879721 | 0.000117 | 0.00107 | down |
| Cd300e | 0.360369 | -1.472454 | 0.000118 | 0.001072 | down |
| Prkn | 0.325526 | -1.619157 | 0.000119 | 0.001086 | down |
| Akap2 | 0.413285 | -1.274792 | 0.00012 | 0.001094 | down |
| Hpse2 | 2.500158 | 1.322019 | 0.00012 | 0.001094 | up |
| H2az1-ps1 | 2.078721 | 1.0556963 | 0.00012 | 0.001094 | up |
| Rspo4 | 28.21125 | 4.8181988 | 0.000121 | 0.001096 | up |
| Dpysl4 | 4.758222 | 2.2504225 | 0.000123 | 0.001112 | up |
| AC131483.1 | 0.472637 | -1.081197 | 0.000124 | 0.001117 | down |
| Sema3e | 0.385085 | -1.376752 | 0.000125 | 0.00113 | down |
| Lvrn | 3.590902 | 1.8443462 | 0.000125 | 0.001131 | up |
| Btbd3 | 0.416778 | -1.262648 | 0.000126 | 0.00114 | down |
| Mroh6 | 16.40706 | 4.0362447 | 0.000127 | 0.001142 | up |
| Frem2 | 0.429397 | -1.219616 | 0.000127 | 0.001144 | down |
| Rbm47 | 0.48621 | -1.040347 | 0.000128 | 0.001155 | down |
| Lmntd2 | 2.468565 | 1.3036724 | 0.000129 | 0.001156 | up |
| Lsmem2 | 0.455686 | -1.133889 | 0.00013 | 0.001163 | down |
| Plcd3 | 2.161485 | 1.1120227 | 0.000131 | 0.001176 | up |
| Mutyh | 2.557893 | 1.3549561 | 0.000132 | 0.001178 | up |
| Rnf150 | 2.04234 | 1.0302227 | 0.000134 | 0.001198 | up |
| Blnk | 2.406408 | 1.2668816 | 0.000134 | 0.0012 | up |
| Wfdc18 | 50230.44 | 15.616274 | 0.000135 | 0.001203 | up |
| Col6a3 | 2.13332 | 1.0931003 | 0.000136 | 0.001209 | up |
| Golga3 | 0.154516 | -2.694168 | 0.000136 | 0.00121 | down |
| Ms4a6c | 2.291772 | 1.1964634 | 0.000136 | 0.00121 | up |
| Gpr158 | 0.24475 | -2.030621 | 0.000136 | 0.001213 | down |
| Comp | 4.575938 | 2.1940674 | 0.000139 | 0.001231 | up |
| Cwh43 | 0.254351 | -1.975105 | 0.000139 | 0.001232 | down |
| Pmfbp1 | 5.86425 | 2.5519465 | 0.000139 | 0.001233 | up |
| Cd83 | 2.107635 | 1.0756248 | 0.000139 | 0.001233 | up |
| Ms4a1 | 0.24248 | -2.044064 | 0.000141 | 0.001249 | down |
| ENSRNOG00000069326 | 0.403859 | -1.308077 | 0.000142 | 0.001256 | down |
| Ly86 | 2.199816 | 1.1373832 | 0.000143 | 0.001263 | up |
| Plet1 | 0.43838 | -1.189746 | 0.000144 | 0.001264 | down |
| Pdcd1 | 3.031298 | 1.5999356 | 0.000145 | 0.001276 | up |

| | | | | | |
|---|---|---|---|---|---|
| LOC102548506 | 6.900271 | 2.786653 | 0.000145 | 0.001279 | up |
| ENSRNOG00000063737 | 12.58502 | 3.6536355 | 0.000147 | 0.001289 | up |
| Mib1 | 0.392006 | -1.351054 | 0.000147 | 0.001289 | down |
| Necab2 | 3.228913 | 1.6910484 | 0.000149 | 0.001302 | up |
| Map2 | 0.310857 | -1.685676 | 0.00015 | 0.001311 | down |
| Arhgef38 | 0.461193 | -1.116559 | 0.00015 | 0.001312 | down |
| Kcng2 | 2.770179 | 1.4699791 | 0.000151 | 0.001319 | up |
| Asic4 | 2.964584 | 1.5678298 | 0.000151 | 0.001319 | up |
| Cd300a | 2.354079 | 1.2351628 | 0.000151 | 0.001319 | up |
| Tnr | 0.229585 | -2.1229 | 0.000151 | 0.001322 | down |
| Olfm4 | 3.082572 | 1.6241345 | 0.000153 | 0.001334 | up |
| Pcdh20 | 0.13927 | -2.844039 | 0.000153 | 0.001335 | down |
| ENSRNOG00000063856 | 2.961061 | 1.566114 | 0.000153 | 0.001336 | up |
| Kmt5c | 2.543733 | 1.3469471 | 0.000153 | 0.001336 | up |
| ENSRNOG00000063597 | 16.46515 | 4.0413436 | 0.000154 | 0.001342 | up |
| Nppb | 2301.536 | 11.168381 | 0.000154 | 0.001344 | up |
| Chst5 | 2.909863 | 1.5409511 | 0.000156 | 0.001357 | up |
| ENSRNOG00000069490 | 4.237654 | 2.0832659 | 0.000157 | 0.001365 | up |
| Pex5l | 0.2905 | -1.783391 | 0.000159 | 0.001374 | down |
| Il25 | 0.1288 | -2.956797 | 0.000163 | 0.001406 | down |
| Cntn1 | 0.298008 | -1.746577 | 0.000164 | 0.001412 | down |
| LOC681325 | 0.443068 | -1.174401 | 0.000164 | 0.001413 | down |
| Hist1h4m | 5.169285 | 2.3699648 | 0.000168 | 0.001442 | up |
| Plcxd2 | 2.290274 | 1.19552 | 0.000168 | 0.001445 | up |
| Klhdc7a | 0.412033 | -1.279168 | 0.000169 | 0.001451 | down |
| Hmox1 | 2.381818 | 1.2520632 | 0.000169 | 0.001452 | up |
| Ptger3 | 2.580966 | 1.3679113 | 0.00017 | 0.001462 | up |
| Cd200r1 | 2.304073 | 1.2041864 | 0.000171 | 0.001463 | up |
| Pag1 | 0.319842 | -1.644568 | 0.000176 | 0.001499 | down |
| Slc7a11 | 3.455694 | 1.7889756 | 0.000179 | 0.001524 | up |
| Steap4 | 3.331188 | 1.7360369 | 0.000179 | 0.001525 | up |
| Slc16a6 | 2.42195 | 1.2761691 | 0.000179 | 0.001527 | up |
| ENSRNOG00000066804 | 30.61083 | 4.9359703 | 0.00018 | 0.001535 | up |
| Htr7 | 6.217066 | 2.6362339 | 0.00018 | 0.001535 | up |
| Kcnj15 | 0.367139 | -1.445602 | 0.000183 | 0.001553 | down |
| Slfn5 | 0.465522 | -1.103078 | 0.000183 | 0.001553 | down |
| Catsper4 | 14.28142 | 3.8360679 | 0.000186 | 0.001577 | up |
| ENSRNOG00000062509 | 2.39806 | 1.261868 | 0.000186 | 0.001577 | up |
| Mia | 2.526332 | 1.3370444 | 0.000188 | 0.001585 | up |
| Tnfaip8l2 | 2.085899 | 1.0606691 | 0.000188 | 0.001591 | up |
| Arg1 | 2.23388 | 1.1595516 | 0.000189 | 0.001593 | up |

| | | | | | |
|---|---|---|---|---|---|
| Neil3 | 16.45346 | 4.040319 | 0.000189 | 0.001593 | up |
| Pdgfrl | 2.108584 | 1.0762748 | 0.000189 | 0.001596 | up |
| Stxbp4 | 0.446502 | -1.163262 | 0.000192 | 0.001619 | down |
| Mirlet7c2 | 5.865029 | 2.5521382 | 0.000196 | 0.00165 | up |
| Krt79 | 0.387577 | -1.367447 | 0.000197 | 0.001656 | down |
| Kdm7a | 0.453317 | -1.141408 | 0.000198 | 0.001661 | down |
| Anxa10 | 3621.088 | 11.822208 | 0.0002 | 0.001674 | up |
| Ppm1n | 2.03854 | 1.0275365 | 0.0002 | 0.001676 | up |
| Ebi3 | 4.058062 | 2.0207909 | 0.000202 | 0.001689 | up |
| LOC120094784 | 3.768399 | 1.9139519 | 0.000203 | 0.001701 | up |
| Kif5a | 3.185631 | 1.671579 | 0.000206 | 0.00172 | up |
| Gjb5 | 3.489227 | 1.8029073 | 0.000209 | 0.001737 | up |
| Nckap5 | 0.325109 | -1.621005 | 0.000211 | 0.001755 | down |
| Gmnn | 2.033835 | 1.0242028 | 0.000211 | 0.001757 | up |
| Hmx1 | 579.724 | 9.1792224 | 0.000212 | 0.001762 | up |
| LOC102552404 | 14.56478 | 3.8644117 | 0.000212 | 0.001762 | up |
| Slc38a5 | 0.401969 | -1.314845 | 0.000216 | 0.001785 | down |
| Arrdc3 | 0.460193 | -1.119688 | 0.000219 | 0.001812 | down |
| Haghl | 2.311341 | 1.20873 | 0.00022 | 0.001813 | up |
| Matcap1 | 2.155129 | 1.1077744 | 0.00022 | 0.001813 | up |
| RGD1564409 | 274.5408 | 8.1008769 | 0.00022 | 0.001817 | up |
| Diras2 | 3.056212 | 1.6117447 | 0.000222 | 0.001826 | up |
| C9h6orf141 | 7.023535 | 2.8121972 | 0.000223 | 0.001836 | up |
| Pinlyp | 4.364662 | 2.1258699 | 0.000225 | 0.001854 | up |
| Serpinf1 | 2.455626 | 1.2960907 | 0.000229 | 0.001879 | up |
| Acr | 2.842779 | 1.507302 | 0.000233 | 0.001905 | up |
| Stmn4 | 2.632836 | 1.3966178 | 0.000237 | 0.001932 | up |
| ENSRNOG00000067579 | 4.64114 | 2.2144793 | 0.000238 | 0.001942 | up |
| Garin5b | 2.518251 | 1.3324221 | 0.000239 | 0.001952 | up |
| Gp1ba | 0.480636 | -1.056984 | 0.000239 | 0.001952 | down |
| Gstt3 | 0.459332 | -1.12239 | 0.000242 | 0.001965 | down |
| Cd300le | 2.569019 | 1.3612175 | 0.000245 | 0.001992 | up |
| Soga1 | 0.383788 | -1.381617 | 0.000245 | 0.001993 | down |
| Fbn2 | 2.902084 | 1.5370894 | 0.00025 | 0.002023 | up |
| Rad51b | 2.610854 | 1.3845218 | 0.00025 | 0.002024 | up |
| Dock3 | 2.637177 | 1.3989942 | 0.00025 | 0.002025 | up |
| Kbtbd6 | 0.304772 | -1.714198 | 0.00025 | 0.002026 | down |
| Gpr68 | 2.552883 | 1.3521276 | 0.000253 | 0.002041 | up |
| Ranbp17 | 2.161288 | 1.111891 | 0.000254 | 0.00205 | up |
| Chst13 | 0.37727 | -1.406332 | 0.000261 | 0.002104 | down |
| Nupr1l1 | 2.267235 | 1.1809338 | 0.000262 | 0.002109 | up |

| | | | | | |
|---|---|---|---|---|---|
| Clec2e | 4.680491 | 2.2266598 | 0.000264 | 0.00212 | up |
| Cacng4 | 2.201494 | 1.138483 | 0.000265 | 0.002132 | up |
| Ramp3 | 2.154298 | 1.1072179 | 0.000271 | 0.002173 | up |
| Ada | 2.095732 | 1.067454 | 0.000272 | 0.002181 | up |
| Nupr1 | 2.228019 | 1.1557616 | 0.000272 | 0.002183 | up |
| P3h2 | 0.458912 | -1.12371 | 0.000273 | 0.002188 | down |
| Gli3 | 0.402562 | -1.312718 | 0.000274 | 0.002193 | down |
| Agmat | 3.361597 | 1.7491468 | 0.000277 | 0.002211 | up |
| Ddx6 | 0.457008 | -1.129708 | 0.000277 | 0.002213 | down |
| ENSRNOG00000063035 | 8.54E-05 | -13.51481 | 0.000282 | 0.00225 | down |
| Hmgb2l1 | 3.186802 | 1.6721095 | 0.000282 | 0.00225 | up |
| Tnnt1 | 7.093209 | 2.8264385 | 0.000283 | 0.002256 | up |
| LOC102555109 | 2.373686 | 1.2471291 | 0.000284 | 0.002266 | up |
| St6galnac1 | 37.64317 | 5.2343164 | 0.000287 | 0.002286 | up |
| Olfm2 | 3.412794 | 1.7709534 | 0.000289 | 0.002299 | up |
| Epsti1 | 2.263503 | 1.178557 | 0.00029 | 0.002308 | up |
| Eln | 2.965764 | 1.568404 | 0.000291 | 0.002309 | up |
| Poc1a | 2.39182 | 1.2581086 | 0.000291 | 0.002309 | up |
| Clec11a | 2.346331 | 1.2304064 | 0.000291 | 0.002311 | up |
| ENSRNOG00000067050 | 5158.258 | 12.332668 | 0.000293 | 0.002326 | up |
| Sptbn4 | 2.410174 | 1.2691376 | 0.000296 | 0.002346 | up |
| ENSRNOG00000063179 | 0.362036 | -1.465796 | 0.0003 | 0.002373 | down |
| Rnf11l2 | 0.386103 | -1.372941 | 0.000305 | 0.002402 | down |
| Ndst1 | 0.391188 | -1.354067 | 0.000307 | 0.002415 | down |
| Ipo11-ps1 | 0.473183 | -1.079531 | 0.000309 | 0.002431 | down |
| Dpf1 | 2.897367 | 1.5347423 | 0.000309 | 0.002434 | up |
| Arhgap9 | 2.093281 | 1.0657662 | 0.00031 | 0.002438 | up |
| LOC100364116 | 2.261511 | 1.1772868 | 0.000314 | 0.002462 | up |
| Sult4a1 | 8.402074 | 3.0707455 | 0.000314 | 0.002464 | up |
| LOC100912568 | 0.350186 | -1.513807 | 0.000315 | 0.002469 | down |
| Reps2 | 0.332782 | -1.587353 | 0.000315 | 0.002469 | down |
| Ccn2 | 2.858407 | 1.5152115 | 0.00032 | 0.002506 | up |
| Slc7a13 | 0.26882 | -1.895289 | 0.000321 | 0.002511 | down |
| RGD1563578 | 593.354 | 9.2127493 | 0.000323 | 0.002522 | up |
| Snai3 | 2.475543 | 1.3077447 | 0.000324 | 0.002524 | up |
| Cnih3 | 5.302011 | 2.4065396 | 0.000326 | 0.002542 | up |
| Klf13 | 0.470177 | -1.088723 | 0.000328 | 0.002553 | down |
| ENSRNOG00000063720 | 2.667317 | 1.4153891 | 0.000331 | 0.002569 | up |
| Asb11 | 0.150723 | -2.730027 | 0.000333 | 0.002579 | down |
| Tgfb1i1 | 2.409913 | 1.2689813 | 0.000336 | 0.002603 | up |
| Arl5b | 0.496486 | -1.010175 | 0.000337 | 0.002604 | down |

| | | | | | |
|---|---|---|---|---|---|
| Ccr8 | 5.388322 | 2.4298361 | 0.00034 | 0.002627 | up |
| Fndc7 | 0.385414 | -1.375521 | 0.000347 | 0.002675 | down |
| Ephb1 | 2.189712 | 1.130741 | 0.000352 | 0.0027 | up |
| Cxcl1 | 2.281549 | 1.1900138 | 0.000353 | 0.002708 | up |
| Trpm5 | 0.383338 | -1.383313 | 0.000354 | 0.002713 | down |
| N4bp2l1 | 2.563217 | 1.3579558 | 0.000354 | 0.002713 | up |
| Cd79a | 0.357064 | -1.485746 | 0.000354 | 0.002716 | down |
| Ifit1bl | 0.340421 | -1.554608 | 0.000355 | 0.002722 | down |
| LOC120093683 | 127.7253 | 6.996901 | 0.000358 | 0.002741 | up |
| ENSRNOG00000063687 | 0.450548 | -1.150246 | 0.000361 | 0.002757 | down |
| Cyp2f4 | 0.162986 | -2.617181 | 0.000363 | 0.002772 | down |
| Ccnl2 | 2.085557 | 1.0604328 | 0.000363 | 0.002772 | up |
| Foxf1 | 0.430907 | -1.214553 | 0.000366 | 0.002792 | down |
| Naaladl2 | 0.212836 | -2.232189 | 0.000367 | 0.002803 | down |
| Gas2l3 | 3.464073 | 1.7924693 | 0.00037 | 0.002817 | up |
| Omp | 2.420082 | 1.2750561 | 0.00037 | 0.002817 | up |
| Dcaf17 | 0.415379 | -1.267499 | 0.00037 | 0.002817 | down |
| Il1rl1 | 2.017998 | 1.012925 | 0.00037 | 0.002817 | up |
| Fendrr | 0.473194 | -1.079495 | 0.000371 | 0.002819 | down |
| Ccne2 | 2.293834 | 1.1977607 | 0.000372 | 0.002826 | up |
| C2cd4b | 2.102554 | 1.0721428 | 0.000373 | 0.002833 | up |
| LOC120097771 | 0.469382 | -1.091166 | 0.000375 | 0.002843 | down |
| Adgrg2 | 3.757147 | 1.9096378 | 0.000376 | 0.002852 | up |
| Pgf | 2.446365 | 1.2906398 | 0.000378 | 0.002858 | up |
| Emp2 | 0.442764 | -1.175389 | 0.000379 | 0.002869 | down |
| Lama1 | 9.052383 | 3.1782976 | 0.000379 | 0.002869 | up |
| Mapk11 | 2.311652 | 1.2089245 | 0.00038 | 0.002874 | up |
| Clgn | 16.66018 | 4.0583317 | 0.000381 | 0.002876 | up |
| Fscn3 | 2.737497 | 1.4528576 | 0.000386 | 0.002911 | up |
| Hcrtr2 | 0.121197 | -3.044578 | 0.000389 | 0.002932 | down |
| Mpo | 9.33481 | 3.2226207 | 0.000395 | 0.00297 | up |
| Kcne2 | 0.399598 | -1.323379 | 0.0004 | 0.002996 | down |
| Snord52 | 3.50507 | 1.8094432 | 0.000401 | 0.003008 | up |
| Inpp4b | 0.43082 | -1.214843 | 0.000402 | 0.003009 | down |
| Serpina3a | 10.01607 | 3.3242445 | 0.000403 | 0.003013 | up |
| Dio1 | 0.259194 | -1.947897 | 0.000403 | 0.003017 | down |
| Adora3 | 2.739775 | 1.4540576 | 0.000406 | 0.003032 | up |
| Cdkn1c | 0.492974 | -1.020417 | 0.000408 | 0.003047 | down |
| Tsx | 0.429084 | -1.220667 | 0.000411 | 0.003065 | down |
| AABR07044362.1 | 5.749098 | 2.5233356 | 0.000414 | 0.003081 | up |
| Mest | 2.118566 | 1.0830884 | 0.000414 | 0.003082 | up |

| | | | | | |
|---|---|---|---|---|---|
| Slc38a4 | 2.308916 | 1.2072155 | 0.000417 | 0.0031 | up |
| Anks4b | 0.39665 | -1.334063 | 0.000417 | 0.0031 | down |
| Rgs16 | 3.521265 | 1.8160936 | 0.00042 | 0.003118 | up |
| Acsm2 | 0.333437 | -1.584513 | 0.00042 | 0.003118 | down |
| AC117058.1 | 6.375038 | 2.6724339 | 0.00042 | 0.003118 | up |
| AC112568.1 | 4.264122 | 2.0922489 | 0.000423 | 0.003137 | up |
| Cmtm4 | 0.415562 | -1.266863 | 0.000424 | 0.003143 | down |
| Klf7 | 0.434185 | -1.203618 | 0.000428 | 0.003168 | down |
| LOC120094464 | 4.155945 | 2.0551767 | 0.000431 | 0.003183 | up |
| Gcsam | 2.788223 | 1.4793458 | 0.000435 | 0.003208 | up |
| Ncoa2 | 0.454278 | -1.138354 | 0.000435 | 0.003208 | down |
| Ciita | 2.229864 | 1.156956 | 0.000435 | 0.003208 | up |
| Sprr2g | 0.000297 | -11.71584 | 0.000437 | 0.003219 | down |
| Capn12 | 4.730368 | 2.2419523 | 0.000437 | 0.003219 | up |
| Optc | 2.721338 | 1.4443163 | 0.000446 | 0.003273 | up |
| AABR07008309.1 | 4.760229 | 2.251031 | 0.000449 | 0.003292 | up |
| Lcor | 0.438572 | -1.189114 | 0.000449 | 0.003293 | down |
| LOC687707 | 3.781026 | 1.9187776 | 0.00045 | 0.003297 | up |
| Mroh4 | 357.998 | 8.4838077 | 0.000452 | 0.003305 | up |
| Scd | 0.411093 | -1.282463 | 0.000454 | 0.003318 | down |
| Rundc3b | 0.363135 | -1.461424 | 0.000455 | 0.003319 | down |
| Spock2 | 0.423607 | -1.239203 | 0.00046 | 0.003352 | down |
| Ngf | 4.314149 | 2.109076 | 0.000466 | 0.00339 | up |
| Ebf3 | 4.23321 | 2.0817521 | 0.000468 | 0.003401 | up |
| Kcnq5 | 0.456684 | -1.130731 | 0.000476 | 0.003454 | down |
| Krt71 | 15.04011 | 3.9107428 | 0.000478 | 0.003463 | up |
| Ston2 | 0.474662 | -1.075028 | 0.000483 | 0.0035 | down |
| ENSRNOG00000065234 | 7.586702 | 2.9234729 | 0.000485 | 0.00351 | up |
| ENSRNOG00000068121 | 2.809165 | 1.4901416 | 0.000488 | 0.003529 | up |
| Mfge8 | 3.370249 | 1.7528551 | 0.000497 | 0.003583 | up |
| Car9 | 4.944077 | 2.3057013 | 0.000498 | 0.003585 | up |
| Pasd1 | 0.463277 | -1.110052 | 0.0005 | 0.003597 | down |
| Cpz | 2.675682 | 1.4199067 | 0.0005 | 0.003597 | up |
| LOC120094453 | 3.753424 | 1.9082073 | 0.000503 | 0.003616 | up |
| Baiap2l2 | 2.969715 | 1.5703247 | 0.000508 | 0.003648 | up |
| ENSRNOG00000062522 | 0.319715 | -1.645143 | 0.000511 | 0.00367 | down |
| Tbl1x | 0.488994 | -1.032112 | 0.000514 | 0.003688 | down |
| Sfnl1 | 7.599564 | 2.9259167 | 0.000518 | 0.003709 | up |
| Ccl20 | 0.492142 | -1.022854 | 0.000519 | 0.003712 | down |
| Tanc2 | 0.445048 | -1.167966 | 0.00052 | 0.003721 | down |
| S100a4 | 2.014226 | 1.0102253 | 0.000531 | 0.00379 | up |

| | | | | | |
|---|---|---|---|---|---|
| Hmbox1 | 0.446441 | -1.16346 | 0.000531 | 0.00379 | down |
| Car11 | 2.703597 | 1.4348804 | 0.000533 | 0.003801 | up |
| Gabra1 | 0.328515 | -1.605969 | 0.000533 | 0.003801 | down |
| Nrcam | 0.491209 | -1.025592 | 0.000533 | 0.003803 | down |
| Slc47a2 | 5.44575 | 2.4451308 | 0.000539 | 0.003833 | up |
| Catsperd | 2.252462 | 1.1715028 | 0.000541 | 0.003846 | up |
| Mroh7 | 5.085448 | 2.3463748 | 0.000542 | 0.003851 | up |
| Adamtsl5 | 2.448114 | 1.2916709 | 0.000556 | 0.003943 | up |
| Clec1b | 0.434359 | -1.20304 | 0.000561 | 0.003969 | down |
| Sostdc1 | 0.306726 | -1.704978 | 0.000565 | 0.003999 | down |
| Gdf6 | 22.47659 | 4.490351 | 0.000566 | 0.004001 | up |
| Adam33 | 3.062327 | 1.6146284 | 0.000567 | 0.004007 | up |
| Hoxb7 | 2.672403 | 1.4181378 | 0.000569 | 0.004016 | up |
| Npw | 0.415206 | -1.268102 | 0.000574 | 0.004049 | down |
| Gen1 | 2.834588 | 1.5031392 | 0.000576 | 0.004057 | up |
| Samd12 | 0.406527 | -1.298578 | 0.000576 | 0.004061 | down |
| Disp2 | 2.2797 | 1.1888437 | 0.000578 | 0.004069 | up |
| LOC100911002 | 23.10309 | 4.5300139 | 0.000582 | 0.004096 | up |
| AABR07032255.1 | 3.728482 | 1.8985882 | 0.000583 | 0.004102 | up |
| Tlr11 | 3.970349 | 1.9892659 | 0.000588 | 0.004124 | up |
| Tac3 | 7.004419 | 2.8082654 | 0.000589 | 0.004132 | up |
| Irf8 | 2.21564 | 1.1477234 | 0.000589 | 0.004132 | up |
| Sox11 | 4.851399 | 2.2784008 | 0.000596 | 0.004176 | up |
| Nr2c2 | 0.437027 | -1.194204 | 0.000598 | 0.004188 | down |
| Slc36a3 | 4.518926 | 2.1759799 | 0.000599 | 0.00419 | up |
| Draxin | 2.39653 | 1.2609467 | 0.000602 | 0.004207 | up |
| Cd209d | 49.04906 | 5.6161537 | 0.000605 | 0.004226 | up |
| Tnks | 0.481467 | -1.054491 | 0.000611 | 0.004258 | down |
| Slc6a17 | 2.489647 | 1.3159414 | 0.000612 | 0.004267 | up |
| Rbp7 | 6.809878 | 2.7676289 | 0.000613 | 0.004272 | up |
| Gphn | 0.220097 | -2.183791 | 0.000619 | 0.004306 | down |
| Emp3 | 2.035278 | 1.0252259 | 0.000623 | 0.004329 | up |
| Wasf1 | 2.49161 | 1.3170785 | 0.000626 | 0.004352 | up |
| ENSRNOG00000065113 | 6.077477 | 2.6034726 | 0.00063 | 0.004376 | up |
| Cited4 | 0.412879 | -1.276208 | 0.000631 | 0.00438 | down |
| Gpx7 | 2.029915 | 1.021419 | 0.000637 | 0.004411 | up |
| Cryaa | 22.41017 | 4.4860816 | 0.000638 | 0.004419 | up |
| LOC102548682 | 2.704934 | 1.4355936 | 0.000639 | 0.004422 | up |
| Piezo2 | 0.476318 | -1.070004 | 0.000639 | 0.004422 | down |
| ENSRNOG00000067697 | 1915.038 | 10.903157 | 0.000641 | 0.004429 | up |
| St6galnac5 | 0.038142 | -4.712473 | 0.000641 | 0.004433 | down |

| | | | | | |
|---|---|---|---|---|---|
| LOC120098113 | 2.633166 | 1.3967986 | 0.000642 | 0.004435 | up |
| Arl5c | 2.940945 | 1.5562797 | 0.00065 | 0.004485 | up |
| Cep350 | 0.462346 | -1.112955 | 0.000655 | 0.004512 | down |
| Mgarp | 22.94016 | 4.5198034 | 0.000655 | 0.004514 | up |
| Myoz1 | 0.273306 | -1.87141 | 0.000657 | 0.004523 | down |
| Ccdc15 | 0.357414 | -1.484333 | 0.000658 | 0.004525 | down |
| Icos | 2.456558 | 1.2966383 | 0.000668 | 0.004577 | up |
| Zfp385b | 0.495786 | -1.01221 | 0.000673 | 0.00461 | down |
| Col4a4 | 0.413899 | -1.27265 | 0.000674 | 0.004618 | down |
| Serpind1 | 0.435854 | -1.198083 | 0.000675 | 0.004618 | down |
| Manf | 5.706016 | 2.5124838 | 0.000695 | 0.004738 | up |
| Pced1a | 2.137628 | 1.0960111 | 0.000697 | 0.004745 | up |
| ENSRNOG00000071161 | 0.000889 | -10.13596 | 0.000719 | 0.004876 | down |
| Cd52 | 0.249314 | -2.003965 | 0.000729 | 0.004943 | down |
| Zc3h12d | 3.126839 | 1.6447049 | 0.000732 | 0.004959 | up |
| Snord123 | 7.960147 | 2.9927952 | 0.000733 | 0.004962 | up |
| Ptprh | 0.41625 | -1.264477 | 0.000734 | 0.004969 | down |
| Col6a5 | 0.487021 | -1.037944 | 0.000741 | 0.005007 | down |
| Kcnip2 | 0.388925 | -1.362436 | 0.000749 | 0.005056 | down |
| ENSRNOG00000064012 | 2.772397 | 1.4711339 | 0.000751 | 0.005063 | up |
| Adamts7 | 2.050853 | 1.0362241 | 0.000752 | 0.005065 | up |
| ENSRNOG00000069186 | 3.324591 | 1.7331767 | 0.000753 | 0.00507 | up |
| ENSRNOG00000066981 | 10.4595 | 3.3867417 | 0.000757 | 0.005099 | up |
| Prf1 | 0.480637 | -1.056982 | 0.000758 | 0.0051 | down |
| Nodal | 905.808 | 9.8230615 | 0.000769 | 0.005164 | up |
| ENSRNOG00000063848 | 0.318088 | -1.652501 | 0.000774 | 0.005186 | down |
| Tmem273 | 2.63962 | 1.40033 | 0.000783 | 0.005241 | up |
| ENSRNOG00000065335 | 3.610742 | 1.8522954 | 0.000792 | 0.005298 | up |
| Slc34a2 | 0.437923 | -1.19125 | 0.000792 | 0.0053 | down |
| Clec2dl1 | 4.352809 | 2.1219469 | 0.000808 | 0.005397 | up |
| LOC300024 | 0.423333 | -1.240136 | 0.000811 | 0.005411 | down |
| C2h4orf46 | 2.107675 | 1.0756525 | 0.000814 | 0.005424 | up |
| Pate5 | 0.258264 | -1.953081 | 0.000817 | 0.005437 | down |
| Tmem64 | 0.472947 | -1.080249 | 0.000817 | 0.005437 | down |
| Cbl | 0.469749 | -1.090038 | 0.000819 | 0.005449 | down |
| Calca | 2.667442 | 1.4154567 | 0.00082 | 0.005449 | up |
| Fam124b | 0.328919 | -1.604198 | 0.000825 | 0.005481 | down |
| Ptgfr | 0.476913 | -1.068201 | 0.000826 | 0.005486 | down |
| Nxpe1 | 2.009104 | 1.006552 | 0.000831 | 0.005518 | up |
| LOC102551724 | 10.47582 | 3.388992 | 0.000836 | 0.005542 | up |
| Popdc2 | 0.421818 | -1.245307 | 0.000836 | 0.005542 | down |

| | | | | | |
|---|---|---|---|---|---|
| Rest | 0.485319 | -1.042995 | 0.000838 | 0.00555 | down |
| Aqp12a | 9.397945 | 3.2323453 | 0.000841 | 0.005568 | up |
| Il10 | 12.02963 | 3.5885202 | 0.000842 | 0.005577 | up |
| LOC683961 | 2.013897 | 1.0099902 | 0.000849 | 0.005615 | up |
| Pde4d | 0.49129 | -1.025352 | 0.000855 | 0.005652 | down |
| Rnase1l1 | 8.552195 | 3.0962948 | 0.000864 | 0.005693 | up |
| Smim44 | 2.354288 | 1.2352908 | 0.000865 | 0.005694 | up |
| Scg2 | 3.075405 | 1.6207764 | 0.000866 | 0.005704 | up |
| LOC108351472 | 3.30385 | 1.7241484 | 0.000867 | 0.005706 | up |
| AC118772.2 | 2.182531 | 1.1260018 | 0.00087 | 0.005724 | up |
| Foxq1 | 2.190121 | 1.1310104 | 0.000872 | 0.005731 | up |
| Cd8a | 2.276145 | 1.1865923 | 0.000875 | 0.005755 | up |
| Epor | 3.175348 | 1.6669149 | 0.00088 | 0.005779 | up |
| Palb2 | 2.14939 | 1.103927 | 0.000882 | 0.005792 | up |
| Tnf | 2.456304 | 1.296489 | 0.000883 | 0.005793 | up |
| Zkscan1 | 0.378737 | -1.40073 | 0.000892 | 0.005845 | down |
| Slc16a14 | 0.310202 | -1.688721 | 0.000897 | 0.005873 | down |
| ENSRNOG00000065144 | 2.726705 | 1.4471586 | 0.000899 | 0.005883 | up |
| Adgre1 | 2.247663 | 1.168426 | 0.000899 | 0.005883 | up |
| Pla2g5 | 3.831464 | 1.9378958 | 0.0009 | 0.005885 | up |
| Wnk3 | 0.440575 | -1.18254 | 0.000917 | 0.005987 | down |
| Psca | 4.341223 | 2.1181016 | 0.000924 | 0.006019 | up |
| Tmsb15b2 | 2.019055 | 1.01368 | 0.000925 | 0.006022 | up |
| Dpysl5 | 2.289859 | 1.1952589 | 0.000942 | 0.00612 | up |
| Lrrc71 | 2.149986 | 1.1043275 | 0.000948 | 0.006157 | up |
| LOC108352070 | 11.47074 | 3.519886 | 0.000951 | 0.006175 | up |
| Tet3 | 0.484367 | -1.045828 | 0.000961 | 0.006236 | down |
| Efcab8 | 0.321185 | -1.638525 | 0.000973 | 0.006307 | down |
| Xiap | 0.381642 | -1.389709 | 0.000984 | 0.006362 | down |
| Upk1a | 0.45729 | -1.12882 | 0.000992 | 0.006407 | down |
| Hmgb2-ps6 | 2.507113 | 1.3260273 | 0.000995 | 0.00642 | up |
| Ctla4 | 7.296681 | 2.8672405 | 0.000998 | 0.006437 | up |
| Car12 | 3.430307 | 1.7783378 | 0.001 | 0.006445 | up |
| Per2 | 2.57388 | 1.363945 | 0.001003 | 0.006454 | up |
| Scg3 | 2.71976 | 1.4434795 | 0.001029 | 0.006585 | up |
| Ccdc154 | 0.449465 | -1.153718 | 0.00103 | 0.006593 | down |
| Angptl8 | 2.683498 | 1.4241149 | 0.001032 | 0.006601 | up |
| ENSRNOG00000069106 | 0.458959 | -1.123564 | 0.001033 | 0.006604 | down |
| Plppr3 | 2.138141 | 1.0963568 | 0.001033 | 0.006604 | up |
| Csrnp3 | 0.304539 | -1.715302 | 0.001034 | 0.006604 | down |
| Clec2d | 12.26484 | 3.6164565 | 0.001034 | 0.006604 | up |

| | | | | | |
|---|---|---|---|---|---|
| Cracd | 0.481754 | -1.05363 | 0.001037 | 0.006617 | down |
| LOC100361226 | 19.45627 | 4.2821631 | 0.001042 | 0.006651 | up |
| LOC100360491 | 2.015846 | 1.0113852 | 0.001044 | 0.006662 | up |
| Tab3 | 0.472447 | -1.081775 | 0.001049 | 0.006685 | down |
| Timd4 | 3.483576 | 1.8005692 | 0.001053 | 0.006708 | up |
| Magee2 | 0.231496 | -2.110942 | 0.001055 | 0.006716 | down |
| Il23r | 3.247123 | 1.6991618 | 0.001061 | 0.00675 | up |
| Nog | 2.679899 | 1.4221785 | 0.001062 | 0.006754 | up |
| Erc1 | 0.464059 | -1.107621 | 0.001071 | 0.006797 | down |
| Smim5 | 3.411327 | 1.7703329 | 0.00108 | 0.006848 | up |
| AABR07049134.1 | 0.376342 | -1.409883 | 0.001086 | 0.006874 | down |
| Ccdc85a | 0.492766 | -1.021024 | 0.001106 | 0.006998 | down |
| Chrne | 2.578681 | 1.3666331 | 0.001119 | 0.007068 | up |
| Arc | 3.410022 | 1.769781 | 0.001121 | 0.007073 | up |
| Zc3h12c | 0.449328 | -1.15416 | 0.001127 | 0.007092 | down |
| Gria1 | 0.32915 | -1.603181 | 0.001129 | 0.007104 | down |
| Lyl1 | 2.28805 | 1.1941184 | 0.00114 | 0.007162 | up |
| Cyp27b1 | 11.11018 | 3.4738103 | 0.00115 | 0.007214 | up |
| Adig | 4.812865 | 2.2668958 | 0.001155 | 0.00724 | up |
| Ace3 | 0.474803 | -1.074598 | 0.001165 | 0.007302 | down |
| Arid5b | 0.491269 | -1.025416 | 0.00117 | 0.007329 | down |
| Zfp692 | 2.773434 | 1.4716733 | 0.00119 | 0.00744 | up |
| Lat2 | 2.29766 | 1.2001655 | 0.001194 | 0.007458 | up |
| Tnfrsf13b | 0.371236 | -1.429593 | 0.001201 | 0.007496 | down |
| Cyp2s1 | 0.49072 | -1.027028 | 0.00121 | 0.007538 | down |
| Spaca6 | 2.544876 | 1.3475951 | 0.001214 | 0.007557 | up |
| LOC120096111 | 2.799215 | 1.4850225 | 0.001219 | 0.007586 | up |
| Ffar3 | 13.26048 | 3.7290611 | 0.00122 | 0.007591 | up |
| Rgs7 | 0.436735 | -1.195169 | 0.001229 | 0.007636 | down |
| Mreg | 2.073869 | 1.0523247 | 0.001236 | 0.00767 | up |
| Dach1 | 0.455125 | -1.135666 | 0.001243 | 0.007705 | down |
| Bbx | 0.390153 | -1.357887 | 0.001247 | 0.007732 | down |
| C1galt1 | 0.481539 | -1.054276 | 0.001249 | 0.007736 | down |
| LOC102549694 | 2.339112 | 1.2259611 | 0.001249 | 0.007736 | up |
| Slc39a2 | 0.25027 | -1.998444 | 0.001249 | 0.007736 | down |
| Stx1a | 2.300198 | 1.2017584 | 0.001265 | 0.00782 | up |
| Sftpc | 0.446944 | -1.161834 | 0.001266 | 0.007824 | down |
| LOC108348070 | 0.473476 | -1.078637 | 0.001271 | 0.007846 | down |
| Slc35e2b | 0.421767 | -1.245483 | 0.001288 | 0.00794 | down |
| Prss46 | 2494.308 | 11.284424 | 0.001295 | 0.007977 | up |
| ENSRNOG00000064211 | 2.166018 | 1.1150454 | 0.001296 | 0.007981 | up |

| | | | | | |
|---|---|---|---|---|---|
| Bcl2l15 | 10.30597 | 3.3654082 | 0.001306 | 0.008026 | up |
| Cxcl2 | 3.347042 | 1.7428865 | 0.001306 | 0.008026 | up |
| Bard1 | 2.839126 | 1.5054469 | 0.001316 | 0.008078 | up |
| Prokr1 | 0.25458 | -1.973808 | 0.00132 | 0.008097 | down |
| Fras1 | 0.391093 | -1.354417 | 0.001333 | 0.008161 | down |
| Prr11 | 3.357562 | 1.7474141 | 0.001344 | 0.008216 | up |
| Efna2 | 3.37199 | 1.7536001 | 0.001346 | 0.008223 | up |
| Or10aa3 | 5.364323 | 2.4233961 | 0.001351 | 0.008248 | up |
| Mgam | 0.224692 | -2.153979 | 0.001356 | 0.00827 | down |
| ENSRNOG00000065200 | 0.374435 | -1.417212 | 0.001361 | 0.008297 | down |
| ENSRNOG00000064711 | 2.349553 | 1.2323861 | 0.001369 | 0.008342 | up |
| Rundc3a | 2.105764 | 1.0743434 | 0.001373 | 0.008358 | up |
| Ly49s7 | 10.91395 | 3.4481018 | 0.001374 | 0.008358 | up |
| Hsd17b1 | 2.727406 | 1.4475293 | 0.001381 | 0.008396 | up |
| Sncg | 3.875524 | 1.9543913 | 0.001381 | 0.008396 | up |
| Nell1 | 0.432711 | -1.208524 | 0.001383 | 0.008406 | down |
| LOC120103274 | 2.849841 | 1.5108812 | 0.001386 | 0.00842 | up |
| Fbxl2 | 2.182098 | 1.1257161 | 0.001393 | 0.008463 | up |
| Btla | 0.422696 | -1.242308 | 0.001394 | 0.008469 | down |
| Cpne5 | 2.561386 | 1.3569249 | 0.001407 | 0.008532 | up |
| Gnat2 | 0.466988 | -1.098542 | 0.001408 | 0.008537 | down |
| Flywch2 | 2.122613 | 1.0858417 | 0.001414 | 0.008569 | up |
| ENSRNOG00000063514 | 2.062663 | 1.0445083 | 0.001417 | 0.008586 | up |
| LOC103690064 | 0.391693 | -1.352206 | 0.001419 | 0.008593 | down |
| Gpr84 | 4.792221 | 2.2606946 | 0.001423 | 0.008618 | up |
| Ptpn5 | 405.722 | 8.6643477 | 0.001428 | 0.00864 | up |
| Mrtfb | 0.356911 | -1.486365 | 0.001444 | 0.008726 | down |
| Fabp4 | 8.901576 | 3.1540608 | 0.001445 | 0.008731 | up |
| Gja3 | 3.801458 | 1.9265528 | 0.00145 | 0.008759 | up |
| Matn4 | 0.467579 | -1.096718 | 0.001455 | 0.008782 | down |
| LOC685203 | 0.394744 | -1.341011 | 0.001463 | 0.008814 | down |
| Card14 | 2.435971 | 1.2844967 | 0.001467 | 0.008839 | up |
| Actl10 | 2.209896 | 1.1439782 | 0.00147 | 0.008843 | up |
| Rnase6 | 0.416237 | -1.264523 | 0.001478 | 0.008889 | down |
| Ubap1l | 2.319931 | 1.2140822 | 0.00149 | 0.008955 | up |
| Galns | 2.096061 | 1.0676804 | 0.00149 | 0.008956 | up |
| Ctrb1 | 8.983132 | 3.1672185 | 0.001494 | 0.008979 | up |
| Cep295 | 2.196605 | 1.1352754 | 0.001504 | 0.009025 | up |
| LOC120097581 | 3.134365 | 1.6481733 | 0.001506 | 0.009037 | up |
| Scn1b | 2.166612 | 1.1154406 | 0.001525 | 0.009138 | up |
| Galnt16 | 2.117086 | 1.0820798 | 0.001529 | 0.009158 | up |

| | | | | | |
|---|---|---|---|---|---|
| Glis1 | 3.029617 | 1.5991355 | 0.001531 | 0.009164 | up |
| Slc29a1 | 0.389887 | -1.35887 | 0.001533 | 0.009167 | down |
| ENSRNOG00000067632 | 2.144606 | 1.1007124 | 0.001538 | 0.009195 | up |
| Snord54 | 7.898763 | 2.9816267 | 0.001541 | 0.009204 | up |
| Gli1 | 2.089985 | 1.0634924 | 0.001547 | 0.00923 | up |
| Agps | 0.450686 | -1.149807 | 0.001567 | 0.00934 | down |
| Chrdl2 | 8.362687 | 3.0639666 | 0.001574 | 0.009376 | up |
| Gzmb | 0.473842 | -1.077521 | 0.001591 | 0.009466 | down |
| Anxa13 | 714.9805 | 9.48176 | 0.001593 | 0.009479 | up |
| Gpc2 | 2.510194 | 1.3277987 | 0.0016 | 0.009509 | up |
| Fam13a | 0.427082 | -1.227415 | 0.0016 | 0.009509 | down |
| Fmo3 | 0.433511 | -1.205858 | 0.001607 | 0.009547 | down |
| Folr1 | 2.266304 | 1.1803415 | 0.001616 | 0.009596 | up |
| Myh8 | 0.001967 | -8.990055 | 0.001624 | 0.009639 | down |
| ENSRNOG00000067625 | 0.333341 | -1.58493 | 0.001632 | 0.009681 | down |
| Unc5cl | 2.187974 | 1.1295957 | 0.001637 | 0.00971 | up |
| Macroh2a2 | 2.368436 | 1.2439345 | 0.001638 | 0.009713 | up |
| Dnase1l2 | 3.363173 | 1.749823 | 0.001648 | 0.009768 | up |
| ENSRNOG00000065112 | 2.630797 | 1.3954997 | 0.001683 | 0.00994 | up |
| Fam205a | 3.262301 | 1.70589 | 0.001692 | 0.009982 | up |
| Flt3 | 2.316458 | 1.2119203 | 0.001701 | 0.01003 | up |
| Ch25h | 2.325619 | 1.2176146 | 0.001702 | 0.010032 | up |
| Fam107a | 0.429848 | -1.218101 | 0.00172 | 0.010119 | down |
| LOC102546903 | 3.706795 | 1.8901723 | 0.001722 | 0.010127 | up |
| LOC103692025 | 0.408383 | -1.292006 | 0.001727 | 0.010154 | down |
| Fam229a | 2.79223 | 1.4814179 | 0.001743 | 0.010233 | up |
| AABR07065812.2 | 70.59795 | 6.1415544 | 0.00175 | 0.010271 | up |
| Kcnj13 | 0.496282 | -1.010767 | 0.001759 | 0.010308 | down |
| Angpt4 | 2.566695 | 1.3599118 | 0.00177 | 0.01037 | up |
| Mc5r | 2.897702 | 1.5349092 | 0.001773 | 0.010382 | up |
| Iffo1 | 2.053909 | 1.0383724 | 0.001797 | 0.010506 | up |
| St6galnac2 | 3.251352 | 1.7010396 | 0.001808 | 0.010568 | up |
| Brsk2 | 11.60495 | 3.5366686 | 0.001812 | 0.010584 | up |
| Usp4 | 0.415502 | -1.267072 | 0.001826 | 0.01065 | down |
| ENSRNOG00000068268 | 0.240668 | -2.054883 | 0.001826 | 0.01065 | down |
| Snord46 | 2.465171 | 1.301688 | 0.001838 | 0.010718 | up |
| Trnp1 | 3.244899 | 1.6981734 | 0.001846 | 0.010751 | up |
| Tubb1 | 0.358885 | -1.478405 | 0.001862 | 0.010838 | down |
| Hes2 | 5.033796 | 2.3316467 | 0.001874 | 0.010878 | up |
| RGD1562462 | 11.96681 | 3.5809663 | 0.001877 | 0.010893 | up |
| Cacng3 | 0.083966 | -3.574048 | 0.001879 | 0.010903 | down |

| | | | | | |
|---|---|---|---|---|---|
| LOC108349010 | 0.477849 | -1.065374 | 0.001884 | 0.010926 | down |
| Homer2 | 2.033297 | 1.0238209 | 0.001945 | 0.011233 | up |
| ENSRNOG00000069456 | 1102.066 | 10.105995 | 0.00195 | 0.011252 | up |
| LOC100909660 | 124.8359 | 6.9638892 | 0.00196 | 0.011294 | up |
| Dtx1 | 2.099944 | 1.0703505 | 0.001987 | 0.01142 | up |
| Reep2 | 2.497576 | 1.3205283 | 0.001993 | 0.011456 | up |
| Ar | 0.088714 | -3.494689 | 0.002028 | 0.011644 | down |
| Klrc3 | 2.779072 | 1.4746032 | 0.002034 | 0.011674 | up |
| Irf3 | 2.139318 | 1.0971513 | 0.002036 | 0.011681 | up |
| ENSRNOG00000067112 | 2770.16 | 11.435754 | 0.002037 | 0.011681 | up |
| Lypd1 | 2.561157 | 1.3567959 | 0.00205 | 0.011747 | up |
| Nsun7 | 0.418202 | -1.257729 | 0.002063 | 0.011816 | down |
| ENSRNOG00000065967 | 3.801192 | 1.9264518 | 0.002127 | 0.012132 | up |
| Dbp | 2.771166 | 1.4704933 | 0.002131 | 0.012155 | up |
| Olr1585 | 2.664465 | 1.4138458 | 0.002133 | 0.012159 | up |
| Ooep | 0.383456 | -1.382868 | 0.002134 | 0.012164 | down |
| Ftl1 | 2.05498 | 1.0391246 | 0.002141 | 0.01219 | up |
| Ddah2 | 2.284318 | 1.1917633 | 0.002165 | 0.012299 | up |
| Socs1 | 2.061609 | 1.0437709 | 0.002185 | 0.012397 | up |
| Wfdc21 | 4.356228 | 2.1230796 | 0.002186 | 0.012397 | up |
| Fam83b | 0.407763 | -1.294198 | 0.002186 | 0.012397 | down |
| Strip2 | 2.387388 | 1.2554333 | 0.002192 | 0.01242 | up |
| Slc13a3 | 13.90709 | 3.7977485 | 0.002212 | 0.012517 | up |
| Serp2 | 3.532268 | 1.8205947 | 0.002212 | 0.012517 | up |
| Col4a3 | 0.331939 | -1.591009 | 0.002216 | 0.012536 | down |
| Gfap | 3.123603 | 1.643211 | 0.002227 | 0.012584 | up |
| Gpr50 | 0.128095 | -2.96471 | 0.002231 | 0.012601 | down |
| Ern1 | 0.467625 | -1.096576 | 0.002242 | 0.012657 | down |
| Veph1 | 0.388362 | -1.364526 | 0.002244 | 0.012665 | down |
| Negr1 | 0.426002 | -1.231066 | 0.002254 | 0.012708 | down |
| Arxes2 | 8.176272 | 3.0314431 | 0.002257 | 0.01272 | up |
| Pla2g4b | 2.296136 | 1.1992079 | 0.002267 | 0.012745 | up |
| Bdkrb1 | 11.1107 | 3.4738772 | 0.002308 | 0.012941 | up |
| Cd79b | 0.361826 | -1.466634 | 0.00231 | 0.012947 | down |
| Fam193b | 2.31252 | 1.2094658 | 0.002322 | 0.013003 | up |
| Ppp1r1a | 2.609518 | 1.3837834 | 0.002322 | 0.013003 | up |
| ENSRNOG00000063458 | 0.497477 | -1.007298 | 0.002331 | 0.013042 | down |
| LOC102553350 | 6.733723 | 2.7514043 | 0.002333 | 0.01305 | up |
| ENSRNOG00000064302 | 0.432626 | -1.208809 | 0.002338 | 0.013071 | down |
| Tnfrsf25 | 2.618562 | 1.3887748 | 0.002342 | 0.013091 | up |
| Snord12 | 3.919007 | 1.9704883 | 0.002354 | 0.013151 | up |

| | | | | | |
|---|---|---|---|---|---|
| Ptk6 | 16.66993 | 4.059176 | 0.002371 | 0.013236 | up |
| Drp2 | 2.410791 | 1.2695067 | 0.002372 | 0.013236 | up |
| Nfib | 0.452653 | -1.143523 | 0.002378 | 0.013261 | down |
| Gfra4 | 8.358374 | 3.0632223 | 0.002384 | 0.013285 | up |
| Snord30 | 3.187502 | 1.6724264 | 0.002393 | 0.013327 | up |
| ENSRNOG00000062535 | 3.437388 | 1.7813129 | 0.002401 | 0.013362 | up |
| Pcdh19 | 0.343459 | -1.541789 | 0.002416 | 0.013439 | down |
| Gpr162 | 3.915495 | 1.9691946 | 0.002426 | 0.013482 | up |
| Tnnc2 | 0.036344 | -4.782148 | 0.002475 | 0.013719 | down |
| LOC120100346 | 5.123851 | 2.3572285 | 0.002483 | 0.013757 | up |
| P2ry13 | 2.312605 | 1.209519 | 0.002485 | 0.013761 | up |
| Tigd3 | 2.358686 | 1.2379833 | 0.002517 | 0.013913 | up |
| Atp2b3 | 0.451287 | -1.147884 | 0.002531 | 0.013978 | down |
| Atp8b3 | 2.581471 | 1.3681931 | 0.002532 | 0.013978 | up |
| Hoxb8 | 3.207132 | 1.6812836 | 0.002534 | 0.013987 | up |
| Kremen2 | 606.368 | 9.2440498 | 0.002537 | 0.013994 | up |
| Lrp2 | 0.435983 | -1.197658 | 0.002542 | 0.014018 | down |
| Cntnap1 | 2.158577 | 1.1100808 | 0.002544 | 0.014026 | up |
| H1f8 | 921.678 | 9.848119 | 0.002574 | 0.014178 | up |
| ENSRNOG00000065178 | 2.963742 | 1.56742 | 0.002607 | 0.014334 | up |
| ENSRNOG00000071137 | 6.127388 | 2.6152723 | 0.002632 | 0.01442 | up |
| Minar2 | 2.152969 | 1.1063279 | 0.002633 | 0.014422 | up |
| ENSRNOG00000064855 | 3.261182 | 1.7053948 | 0.002636 | 0.014428 | up |
| Pax2 | 714.6073 | 9.4810069 | 0.002637 | 0.014431 | up |
| ENSRNOG00000067032 | 39.03574 | 5.2867238 | 0.002659 | 0.014534 | up |
| Nipal4 | 2.91948 | 1.5457113 | 0.002696 | 0.01471 | up |
| Brinp2 | 2.380619 | 1.251337 | 0.002702 | 0.014728 | up |
| LOC102550644 | 0.460336 | -1.119242 | 0.002705 | 0.01474 | down |
| Fst | 2.069414 | 1.0492224 | 0.002719 | 0.014807 | up |
| Arhgap40 | 352.034 | 8.459571 | 0.002723 | 0.014825 | up |
| Slc12a5 | 2.436956 | 1.28508 | 0.002739 | 0.0149 | up |
| Pwwp4 | 3.549016 | 1.8274192 | 0.002742 | 0.014914 | up |
| AC112557.1 | 2.369408 | 1.2445268 | 0.002798 | 0.015176 | up |
| Tmc3 | 20.38084 | 4.3491416 | 0.00284 | 0.015371 | up |
| Adora1 | 2.519687 | 1.3332443 | 0.002841 | 0.015371 | up |
| Ogdhl | 2.180253 | 1.1244958 | 0.002849 | 0.015406 | up |
| ENSRNOG00000066321 | 2.359458 | 1.2384553 | 0.002855 | 0.015433 | up |
| Ctsg | 1591.556 | 10.636222 | 0.002872 | 0.015503 | up |
| Dennd1c | 2.006351 | 1.004574 | 0.002872 | 0.015503 | up |
| Fbxl19 | 2.299526 | 1.2013364 | 0.002875 | 0.015515 | up |
| Snrpel1 | 2.02538 | 1.0181928 | 0.002876 | 0.015515 | up |

| | | | | | |
|---|---|---|---|---|---|
| Col11a1 | 8.610203 | 3.1060473 | 0.002884 | 0.015547 | up |
| Hist1h2ao | 6.739851 | 2.7527166 | 0.00289 | 0.015578 | up |
| Prss32 | 10.32998 | 3.3687657 | 0.002904 | 0.015634 | up |
| Nabp1 | 2.370875 | 1.2454196 | 0.002908 | 0.015648 | up |
| ENSRNOG00000065800 | 2.736618 | 1.4523939 | 0.002963 | 0.015906 | up |
| Chrna4 | 0.477273 | -1.067113 | 0.002964 | 0.015907 | down |
| Msx3 | 0.392896 | -1.347782 | 0.002975 | 0.01595 | down |
| Krt88 | 30.30463 | 4.9214663 | 0.002993 | 0.016028 | up |
| Pclo | 0.41276 | -1.276625 | 0.003023 | 0.016179 | down |
| Pltp | 2.11517 | 1.0807733 | 0.003037 | 0.016233 | up |
| Il22ra2 | 0.391475 | -1.35301 | 0.003043 | 0.01626 | down |
| ENSRNOG00000070797 | 3.879582 | 1.9559012 | 0.003047 | 0.016278 | up |
| Pnoc | 21.16307 | 4.4034772 | 0.003052 | 0.016301 | up |
| LOC120101643 | 2.115158 | 1.0807657 | 0.003055 | 0.016308 | up |
| Trex2 | 1238.856 | 10.274793 | 0.003066 | 0.016361 | up |
| ENSRNOG00000068419 | 2.632307 | 1.3963276 | 0.003075 | 0.016393 | up |
| LOC120099441 | 2.443372 | 1.2888733 | 0.003095 | 0.016486 | up |
| LOC102554746 | 4.735425 | 2.2434938 | 0.0031 | 0.016503 | up |
| Cyp2d3 | 2.547674 | 1.3491807 | 0.003108 | 0.016528 | up |
| Micb | 0.398538 | -1.327209 | 0.003133 | 0.016642 | down |
| Evi2a | 2.010235 | 1.0073641 | 0.003143 | 0.016681 | up |
| Itgad | 2.495751 | 1.319474 | 0.003147 | 0.016694 | up |
| ENSRNOG00000063332 | 5.99352 | 2.5834035 | 0.003183 | 0.016857 | up |
| Nlrp3 | 2.417308 | 1.2734013 | 0.003188 | 0.016875 | up |
| Mro | 4.716275 | 2.2376478 | 0.003204 | 0.016944 | up |
| Tcf15 | 4.68717 | 2.2287172 | 0.003205 | 0.016944 | up |
| Tmem125 | 5.120788 | 2.3563659 | 0.003205 | 0.016944 | up |
| Btn1a1 | 4.980648 | 2.3163333 | 0.003228 | 0.017053 | up |
| Mid2 | 0.459702 | -1.121229 | 0.003228 | 0.017053 | down |
| Scart1 | 5.004551 | 2.3232406 | 0.003235 | 0.017082 | up |
| Muc2 | 8.46321 | 3.081205 | 0.00324 | 0.017097 | up |
| Elane | 2506.664 | 11.291553 | 0.003244 | 0.017117 | up |
| Ereg | 3.855318 | 1.9468499 | 0.003246 | 0.017123 | up |
| ENSRNOG00000067737 | 0.384151 | -1.380256 | 0.003262 | 0.017177 | down |
| ENSRNOG00000065416 | 3.257571 | 1.7037966 | 0.003266 | 0.017191 | up |
| Pdcd1lg2 | 7.611507 | 2.9281821 | 0.003273 | 0.017226 | up |
| ENSRNOG00000071088 | 2.387493 | 1.2554962 | 0.003287 | 0.01728 | up |
| Wnt9b | 0.427954 | -1.224474 | 0.003288 | 0.01728 | down |
| Atp2a1 | 0.077019 | -3.698649 | 0.0033 | 0.017327 | down |
| Lhfpl4 | 2.081816 | 1.0578424 | 0.003325 | 0.017439 | up |
| Ptprn2 | 2.058249 | 1.0414172 | 0.003329 | 0.017453 | up |

| | | | | | |
|---|---|---|---|---|---|
| Slc4a11 | 2.132614 | 1.092623 | 0.003331 | 0.017454 | up |
| ENSRNOG00000069366 | 2.301135 | 1.2023458 | 0.003333 | 0.01746 | up |
| LOC108352411 | 2.926424 | 1.549139 | 0.003334 | 0.017461 | up |
| ENSRNOG00000069879 | 3.396337 | 1.7639795 | 0.003375 | 0.017623 | up |
| ENSRNOG00000065655 | 9.249324 | 3.2093479 | 0.00338 | 0.017632 | up |
| ENSRNOG00000069208 | 2.647115 | 1.4044209 | 0.003443 | 0.01789 | up |
| Gabbr2 | 4.935729 | 2.3032631 | 0.003447 | 0.017903 | up |
| Galnt14 | 0.300389 | -1.735097 | 0.003448 | 0.017906 | down |
| Tlr10 | 2.932737 | 1.5522475 | 0.003459 | 0.01796 | up |
| Car7 | 0.19903 | -2.328941 | 0.00348 | 0.018057 | down |
| Tmem61 | 2.546312 | 1.3484091 | 0.003497 | 0.018127 | up |
| Otog | 107.614 | 6.749722 | 0.003507 | 0.018169 | up |
| Snap91 | 2.488315 | 1.3151692 | 0.003543 | 0.018328 | up |
| Rasal1 | 2.441464 | 1.2877467 | 0.003551 | 0.018361 | up |
| ENSRNOG00000070740 | 2.302264 | 1.2030534 | 0.003566 | 0.018428 | up |
| RGD1309808 | 2.567091 | 1.3601342 | 0.00357 | 0.018443 | up |
| Lcn2 | 5.70844 | 2.5130966 | 0.003574 | 0.018464 | up |
| Gbp6 | 665.0008 | 9.3772123 | 0.003596 | 0.018553 | up |
| Orm1 | 2.769829 | 1.4697971 | 0.003617 | 0.018639 | up |
| Cd70 | 1364.072 | 10.413704 | 0.003637 | 0.018719 | up |
| Spata48 | 2.488604 | 1.3153364 | 0.003649 | 0.018771 | up |
| Sphkap | 0.348611 | -1.520312 | 0.003661 | 0.018818 | down |
| Steap1 | 2.409199 | 1.2685536 | 0.00367 | 0.018859 | up |
| Brip1 | 2.385375 | 1.2542161 | 0.003712 | 0.019043 | up |
| Npbwr1 | 0.124883 | -3.001346 | 0.003719 | 0.019072 | down |
| Sema4g | 2.170819 | 1.1182397 | 0.003744 | 0.019173 | up |
| ENSRNOG00000064824 | 0.380939 | -1.392367 | 0.003751 | 0.01919 | down |
| Clnk | 2.231165 | 1.1577973 | 0.003764 | 0.019248 | up |
| Havcr2 | 2.32547 | 1.2175225 | 0.003774 | 0.019286 | up |
| Mis18a | 3.204861 | 1.6802616 | 0.003801 | 0.019402 | up |
| Snai1 | 2.07811 | 1.0552722 | 0.003814 | 0.019449 | up |
| LOC685767 | 0.207225 | -2.270728 | 0.003846 | 0.019575 | down |
| LOC120094450 | 7.688434 | 2.9426898 | 0.003881 | 0.019745 | up |
| Ankrd1 | 2.653204 | 1.4077353 | 0.003904 | 0.01984 | up |
| Fign | 0.46802 | -1.095357 | 0.003921 | 0.019895 | down |
| Dmbt1 | 10.60534 | 3.4067185 | 0.003975 | 0.020119 | up |
| ENSRNOG00000068511 | 0.38689 | -1.370005 | 0.003976 | 0.020122 | down |
| Sptbn5 | 2.65181 | 1.4069771 | 0.003982 | 0.020146 | up |
| Cracr2a | 0.435703 | -1.198582 | 0.004001 | 0.020213 | down |
| Prodh2 | 0.427469 | -1.226108 | 0.004024 | 0.020309 | down |
| ENSRNOG00000071037 | 23.58322 | 4.5596891 | 0.004025 | 0.020309 | up |

| | | | | | |
|---|---|---|---|---|---|
| Muc3a | 11.18093 | 3.4829682 | 0.00411 | 0.020688 | up |
| Pcbd1 | 12224.67 | 13.577508 | 0.004128 | 0.020767 | up |
| AABR07053509.1 | 2.007087 | 1.0051032 | 0.004158 | 0.020895 | up |
| Sfmbt2 | 2.969625 | 1.5702807 | 0.004178 | 0.020986 | up |
| ENSRNOG00000064675 | 3811.554 | 11.896164 | 0.004183 | 0.021004 | up |
| Fxyd7 | 4.133636 | 2.0474113 | 0.004183 | 0.021004 | up |
| Lta | 5.304583 | 2.4072394 | 0.004215 | 0.021143 | up |
| Trpv1 | 2.37432 | 1.2475141 | 0.004217 | 0.021147 | up |
| Rasl11b | 2.34492 | 1.2295385 | 0.004223 | 0.021166 | up |
| S100g | 0.374939 | -1.415272 | 0.004244 | 0.021262 | down |
| LOC120094528 | 3.865967 | 1.9508291 | 0.004253 | 0.021305 | up |
| Zbp1 | 2.505531 | 1.3251162 | 0.004293 | 0.021472 | up |
| ENSRNOG00000066580 | 2.144492 | 1.1006357 | 0.004307 | 0.021531 | up |
| B3gnt3 | 2.747298 | 1.4580136 | 0.004323 | 0.021595 | up |
| Rnf112 | 4.111887 | 2.0398007 | 0.004363 | 0.021769 | up |
| Kcnh4 | 0.353623 | -1.499716 | 0.004365 | 0.021773 | down |
| Ugt2b | 0.139553 | -2.84112 | 0.00437 | 0.021793 | down |
| Wnt7a | 0.496933 | -1.008876 | 0.004374 | 0.0218 | down |
| ENSRNOG00000068718 | 3.26331 | 1.7063359 | 0.0044 | 0.021916 | up |
| C5 | 0.157998 | -2.662018 | 0.004413 | 0.021962 | down |
| Slc14a2 | 437.6461 | 8.7736209 | 0.004458 | 0.022164 | up |
| Umodl1 | 2.623244 | 1.3913519 | 0.004468 | 0.0222 | up |
| ENSRNOG00000070561 | 3.101479 | 1.6329562 | 0.004477 | 0.022223 | up |
| Snord58b | 2.996737 | 1.5833925 | 0.004496 | 0.022297 | up |
| Abcg8 | 201.786 | 7.6566823 | 0.004552 | 0.022521 | up |
| Ms4a4c-ps1 | 0.373854 | -1.419454 | 0.004589 | 0.022642 | down |
| Tnfrsf14 | 2.807123 | 1.4890922 | 0.004658 | 0.022957 | up |
| AC131360.2 | 0.387437 | -1.367967 | 0.004699 | 0.023145 | down |
| Lpar6 | 2.05952 | 1.0423083 | 0.004716 | 0.023219 | up |
| LOC120094888 | 4.542608 | 2.1835209 | 0.00472 | 0.023234 | up |
| LOC120100232 | 3.834093 | 1.9388853 | 0.004746 | 0.023326 | up |
| Rpl36a-ps7 | 2.235265 | 1.1604458 | 0.004772 | 0.023431 | up |
| Pak1ip1 | 2.040112 | 1.0286487 | 0.004787 | 0.02348 | up |
| AABR07015941.1 | 6869.824 | 12.746057 | 0.0048 | 0.023534 | up |
| Or2b11 | 433.694 | 8.7605337 | 0.004811 | 0.023581 | up |
| ENSRNOG00000067218 | 2.835606 | 1.503657 | 0.004826 | 0.023624 | up |
| ENSRNOG00000063395 | 2.283768 | 1.1914162 | 0.004831 | 0.02363 | up |
| LOC102550841 | 0.458033 | -1.126478 | 0.004837 | 0.023653 | down |
| Lingo2 | 0.469424 | -1.091038 | 0.004848 | 0.023699 | down |
| Or2y1g | 5.217267 | 2.3832943 | 0.004858 | 0.023735 | up |
| Atf7ip2 | 2.100695 | 1.0708669 | 0.004894 | 0.023881 | up |

| | | | | | |
|---|---|---|---|---|---|
| Mir25 | 3.435052 | 1.7803321 | 0.004916 | 0.023954 | up |
| Cxcr1 | 0.319778 | -1.644857 | 0.004918 | 0.023957 | down |
| Tmem229a | 0.255137 | -1.970658 | 0.004933 | 0.024014 | down |
| Tex36 | 0.296106 | -1.755817 | 0.004977 | 0.024196 | down |
| Gpbp1l2 | 2.163398 | 1.1132988 | 0.004985 | 0.024223 | up |
| ENSRNOG00000068486 | 3.209635 | 1.6824093 | 0.00499 | 0.024235 | up |
| Atp6v1e2 | 4.857809 | 2.2803058 | 0.004992 | 0.024242 | up |
| Duoxa2 | 2.772112 | 1.4709857 | 0.005028 | 0.024356 | up |
| Kcp | 2.578402 | 1.3664774 | 0.005067 | 0.024531 | up |
| ENSRNOG00000066240 | 3.478663 | 1.798533 | 0.00508 | 0.02457 | up |
| LOC120095461 | 3.862357 | 1.9494815 | 0.005104 | 0.024663 | up |
| Pkd1l2 | 2.178878 | 1.1235856 | 0.005123 | 0.024734 | up |
| Cyp1a1 | 0.23632 | -2.081187 | 0.00514 | 0.0248 | down |
| ENSRNOG00000063488 | 3.422024 | 1.7748499 | 0.005163 | 0.024888 | up |
| ENSRNOG00000063568 | 3.588548 | 1.8434004 | 0.005174 | 0.024936 | up |
| Atp6v0d2 | 4.264596 | 2.092409 | 0.005183 | 0.024975 | up |
| Tspan10 | 15.26608 | 3.9322582 | 0.005203 | 0.025059 | up |
| ENSRNOG00000063519 | 7.645113 | 2.9345378 | 0.005232 | 0.025187 | up |
| Cadm1 | 0.496595 | -1.009857 | 0.005257 | 0.025286 | down |
| Chodl | 0.37374 | -1.419895 | 0.005332 | 0.025557 | down |
| Npr3 | 0.359273 | -1.476848 | 0.005335 | 0.025564 | down |
| RT1-DOa | 2.001101 | 1.0007938 | 0.005357 | 0.025655 | up |
| LOC102556098 | 2.196425 | 1.1351573 | 0.005364 | 0.025675 | up |
| LOC120100091 | 9.638756 | 3.2688469 | 0.005373 | 0.025708 | up |
| ENSRNOG00000068351 | 10.28904 | 3.3630366 | 0.005419 | 0.025904 | up |
| C13h1orf116 | 0.445363 | -1.166945 | 0.005433 | 0.025961 | down |
| Klf9 | 0.447941 | -1.158618 | 0.005451 | 0.026028 | down |
| LOC120094441 | 3.220859 | 1.6874454 | 0.005454 | 0.026033 | up |
| Car4 | 0.467042 | -1.098376 | 0.005483 | 0.026142 | down |
| Tslp | 2.881823 | 1.5269819 | 0.005492 | 0.026176 | up |
| Lcn5 | 999.0065 | 9.9643502 | 0.005527 | 0.026339 | up |
| Slc35f1 | 0.372968 | -1.422874 | 0.005549 | 0.026423 | down |
| ENSRNOG00000064147 | 0.112303 | -3.154529 | 0.005607 | 0.026673 | down |
| ENSRNOG00000064058 | 2.847684 | 1.5097891 | 0.005626 | 0.026753 | up |
| Myh7b | 2.345816 | 1.2300898 | 0.005638 | 0.026784 | up |
| ENSRNOG00000069437 | 3.974467 | 1.9907614 | 0.0057 | 0.027011 | up |
| Hist1h4m | 2.511293 | 1.3284301 | 0.005704 | 0.027015 | up |
| Oscar | 2.329958 | 1.2203037 | 0.005713 | 0.027043 | up |
| Hcn4 | 0.491203 | -1.025608 | 0.005751 | 0.027191 | down |
| Itga10 | 10.22442 | 3.3539465 | 0.005774 | 0.027261 | up |
| Cep85l | 0.493622 | -1.018521 | 0.005784 | 0.027293 | down |

| | | | | | |
|---|---|---|---|---|---|
| Slc29a4 | 2.729302 | 1.4485321 | 0.00579 | 0.027305 | up |
| ENSRNOG00000068207 | 0.383266 | -1.383582 | 0.005804 | 0.027362 | down |
| ENSRNOG00000069947 | 5.372257 | 2.4255283 | 0.005832 | 0.027477 | up |
| Phip | 0.449102 | -1.154886 | 0.005879 | 0.02768 | down |
| Atp7b | 0.269951 | -1.889233 | 0.005906 | 0.027778 | down |
| LOC679368 | 0.440765 | -1.181917 | 0.005907 | 0.027778 | down |
| Or13c7e | 3.722869 | 1.8964149 | 0.005911 | 0.027788 | up |
| Slc6a16 | 3.632594 | 1.8610002 | 0.005919 | 0.027816 | up |
| Gpr65 | 2.233579 | 1.1593576 | 0.005926 | 0.027831 | up |
| Espnl | 2.293646 | 1.1976427 | 0.005956 | 0.027945 | up |
| Nmur2 | 6.631187 | 2.7292672 | 0.006022 | 0.028205 | up |
| ENSRNOG00000071110 | 2.444224 | 1.2893767 | 0.006033 | 0.028247 | up |
| LOC100134871 | 0.008987 | -6.797962 | 0.006046 | 0.028293 | down |
| ENSRNOG00000065658 | 5.959287 | 2.5751397 | 0.006062 | 0.028346 | up |
| Flrt1 | 3.073299 | 1.6197883 | 0.006088 | 0.028433 | up |
| Sntb1 | 0.460958 | -1.117292 | 0.006119 | 0.028558 | down |
| Rdh12 | 5.894677 | 2.5594128 | 0.006133 | 0.028602 | up |
| Slc22a14 | 450.352 | 8.8149093 | 0.006146 | 0.028644 | up |
| Sertm1 | 0.455593 | -1.134184 | 0.006202 | 0.028866 | down |
| Vip | 5.400353 | 2.4330537 | 0.006208 | 0.028886 | up |
| Penk | 2.317947 | 1.2128477 | 0.006322 | 0.029352 | up |
| Slc1a7 | 2.067959 | 1.0482074 | 0.00639 | 0.029603 | up |
| Scn10a | 3.002909 | 1.5863607 | 0.006394 | 0.029616 | up |
| Lefty2 | 13.15165 | 3.7171723 | 0.006444 | 0.029806 | up |
| LOC108352885 | 14.45797 | 3.8537933 | 0.006501 | 0.030038 | up |
| Ccdc184 | 2.65087 | 1.4064661 | 0.006502 | 0.03004 | up |
| Gadd45g | 2.337389 | 1.2248981 | 0.006504 | 0.030043 | up |
| Slc38a8 | 2.603369 | 1.3803801 | 0.006508 | 0.030049 | up |
| Tigit | 6.255222 | 2.645061 | 0.006536 | 0.030152 | up |
| Aurkb | 3.717093 | 1.8941748 | 0.006605 | 0.030394 | up |
| Prss53 | 2.171065 | 1.1184027 | 0.006614 | 0.030418 | up |
| ENSRNOG00000066873 | 3.080144 | 1.6229977 | 0.006661 | 0.030597 | up |
| ENSRNOG00000066146 | 2.140948 | 1.0982501 | 0.006723 | 0.030856 | up |
| Tmem132e | 2.373774 | 1.2471824 | 0.006728 | 0.030876 | up |
| ENSRNOG00000069950 | 0.464292 | -1.106897 | 0.006754 | 0.030976 | down |
| LOC497796 | 4.154179 | 2.0545634 | 0.006756 | 0.030976 | up |
| Cyp2e1 | 0.387936 | -1.366109 | 0.006768 | 0.031019 | down |
| ENSRNOG00000068375 | 0.292044 | -1.775743 | 0.00679 | 0.031102 | down |
| Dhrs9 | 2.457975 | 1.29747 | 0.006814 | 0.031187 | up |
| Hist1h2bg | 6.713618 | 2.7470904 | 0.00683 | 0.031226 | up |
| Bpifb5 | 6.166534 | 2.6244599 | 0.006838 | 0.031251 | up |

| | | | | | |
|---|---|---|---|---|---|
| LOC120100339 | 2.982989 | 1.5767586 | 0.006846 | 0.031274 | up |
| Faxc | 2.07416 | 1.052527 | 0.006863 | 0.031342 | up |
| Hist1h3b | 2.376443 | 1.2488038 | 0.006874 | 0.031372 | up |
| ENSRNOG00000065283 | 0.065742 | -3.927036 | 0.006883 | 0.031397 | down |
| Mir27b | 3.342774 | 1.7410458 | 0.006891 | 0.031421 | up |
| Rd3 | 2.827581 | 1.4995684 | 0.006909 | 0.031479 | up |
| Snca | 0.12171 | -3.038478 | 0.006922 | 0.031527 | down |
| LOC500959 | 2.302778 | 1.2033752 | 0.006944 | 0.031594 | up |
| Stk32c | 2.269828 | 1.1825829 | 0.00698 | 0.031733 | up |
| Moxd1 | 0.431642 | -1.212091 | 0.007047 | 0.031976 | down |
| LOC120103476 | 6.749545 | 2.7547903 | 0.007057 | 0.032009 | up |
| Ncan | 5.011329 | 2.3251932 | 0.007086 | 0.032109 | up |
| Cdhr2 | 0.450252 | -1.151195 | 0.007088 | 0.032114 | down |
| Ciart | 2.702304 | 1.4341901 | 0.007099 | 0.032139 | up |
| ENSRNOG00000068345 | 0.438196 | -1.190351 | 0.007107 | 0.032166 | down |
| Sytl5 | 0.490529 | -1.027591 | 0.007111 | 0.032175 | down |
| Slc1a6 | 86.52939 | 6.4351183 | 0.007165 | 0.032394 | up |
| Esrrg | 0.389396 | -1.360689 | 0.007194 | 0.032496 | down |
| Frmpd1 | 2.539768 | 1.3446965 | 0.007201 | 0.032524 | up |
| Susd5 | 3.004638 | 1.5871912 | 0.007242 | 0.032688 | up |
| ENSRNOG00000070864 | 2.799592 | 1.4852163 | 0.007245 | 0.032691 | up |
| Glod5 | 2.322015 | 1.2153775 | 0.007274 | 0.032808 | up |
| ENSRNOG00000067573 | 4.986147 | 2.3179253 | 0.007303 | 0.032895 | up |
| Oas1e | 4.127936 | 2.0454207 | 0.007307 | 0.032905 | up |
| Mss51 | 2.104052 | 1.07317 | 0.00739 | 0.033207 | up |
| ENSRNOG00000066901 | 4.141673 | 2.0502135 | 0.007414 | 0.033308 | up |
| Lrrc3b | 3.24841 | 1.6997339 | 0.007417 | 0.033318 | up |
| Slc23a1 | 0.485703 | -1.041854 | 0.007423 | 0.033336 | down |
| Slc4a8 | 0.49281 | -1.020896 | 0.007433 | 0.033369 | down |
| AC118772.1 | 0.112309 | -3.154458 | 0.00746 | 0.033471 | down |
| Sh2d5 | 2.926406 | 1.5491298 | 0.007572 | 0.033909 | up |
| Rgs18 | 2.070841 | 1.0502168 | 0.007585 | 0.033961 | up |
| Pla2g4f | 2.571578 | 1.3626538 | 0.007654 | 0.034231 | up |
| ENSRNOG00000066766 | 3.262268 | 1.7058754 | 0.007673 | 0.034294 | up |
| Lyc2 | 0.276817 | -1.852996 | 0.007685 | 0.034333 | down |
| Tafa4 | 15.94434 | 3.9949727 | 0.007701 | 0.034367 | up |
| Arhgap4 | 2.049653 | 1.0353795 | 0.007717 | 0.034426 | up |
| ENSRNOG00000067516 | 0.140267 | -2.833752 | 0.007748 | 0.03453 | down |
| ENSRNOG00000063847 | 3.780464 | 1.9185632 | 0.007757 | 0.034558 | up |
| Btg4 | 739.3033 | 9.5300225 | 0.007762 | 0.034569 | up |
| Stab2 | 3.810193 | 1.9298641 | 0.007765 | 0.034573 | up |

| | | | | | |
|---|---|---|---|---|---|
| Hist1h3b | 14.69961 | 3.8777058 | 0.007785 | 0.034649 | up |
| Sec31b | 3.134588 | 1.648276 | 0.00781 | 0.034738 | up |
| AABR07071383.1 | 509.264 | 8.9922699 | 0.007847 | 0.034885 | up |
| RGD1560559 | 12.18182 | 3.6066577 | 0.007892 | 0.035052 | up |
| Best4 | 6.867529 | 2.7797911 | 0.007897 | 0.035055 | up |
| RGD1563917 | 2.083311 | 1.0588786 | 0.007951 | 0.035267 | up |
| Olr59 | 2.575513 | 1.3648596 | 0.007967 | 0.035321 | up |
| ENSRNOG00000064050 | 0.184222 | -2.440484 | 0.008012 | 0.035509 | down |
| Epb42 | 0.263336 | -1.925024 | 0.008054 | 0.035665 | down |
| Adra1a | 0.47374 | -1.077832 | 0.008073 | 0.035717 | down |
| AABR07006685.1 | 0.349029 | -1.518582 | 0.008101 | 0.035813 | down |
| Fam178b | 2.161619 | 1.112112 | 0.008222 | 0.036259 | up |
| ENSRNOG00000062961 | 0.20552 | -2.282647 | 0.00824 | 0.036315 | down |
| Fpr3 | 0.331341 | -1.593612 | 0.008328 | 0.036642 | down |
| Psd | 2.283472 | 1.1912289 | 0.008333 | 0.036645 | up |
| Pax5 | 0.362244 | -1.464967 | 0.008368 | 0.036768 | down |
| Pvrig | 3.968913 | 1.9887438 | 0.008413 | 0.036946 | up |
| ENSRNOG00000064063 | 0.495043 | -1.014375 | 0.008438 | 0.037033 | down |
| Nxph3 | 2.078094 | 1.0552611 | 0.00854 | 0.03741 | up |
| ENSRNOG00000069704 | 4.603791 | 2.2028223 | 0.008552 | 0.037448 | up |
| Snord12b | 3.634427 | 1.8617279 | 0.008562 | 0.037487 | up |
| LOC120097742 | 3.842879 | 1.9421874 | 0.008693 | 0.037991 | up |
| Rnase11 | 22.46416 | 4.4895531 | 0.008703 | 0.03801 | up |
| Pon1 | 0.413251 | -1.274909 | 0.00874 | 0.038137 | down |
| Caln1 | 0.238043 | -2.070707 | 0.008802 | 0.038362 | down |
| Rpl6-ps1 | 2.307578 | 1.2063796 | 0.00884 | 0.038487 | up |
| Il21r | 2.282921 | 1.190881 | 0.008856 | 0.03855 | up |
| ENSRNOG00000068005 | 0.204731 | -2.288202 | 0.008915 | 0.038776 | down |
| Pdia2 | 3.034996 | 1.6016945 | 0.00893 | 0.038827 | up |
| Kif4b | 9.756973 | 3.2864336 | 0.009131 | 0.039529 | up |
| Gpr35 | 2.589397 | 1.3726163 | 0.009162 | 0.039634 | up |
| Celf5 | 2.031194 | 1.0223278 | 0.009209 | 0.039822 | up |
| Slc46a2 | 0.328877 | -1.604381 | 0.009259 | 0.04001 | down |
| ENSRNOG00000068753 | 7.13935 | 2.8357926 | 0.009266 | 0.040031 | up |
| Myf6 | 0.000622 | -10.65086 | 0.009274 | 0.040053 | down |
| LOC690507 | 12.05685 | 3.5917814 | 0.00928 | 0.040071 | up |
| ENSRNOG00000063527 | 2.515546 | 1.3308716 | 0.009339 | 0.040266 | up |
| Ccdc188 | 354.7922 | 8.4708305 | 0.009396 | 0.040463 | up |
| Rspo3 | 5.79332 | 2.5343905 | 0.00944 | 0.040627 | up |
| Garin3 | 379.364 | 8.567439 | 0.009463 | 0.040698 | up |
| Bpifc | 2.314425 | 1.2106538 | 0.009501 | 0.040811 | up |

| | | | | | |
|---|---|---|---|---|---|
| Cenatac | 2.0932 | 1.06571 | 0.009534 | 0.040924 | up |
| ENSRNOG00000068830 | 2.249353 | 1.1695098 | 0.00959 | 0.041108 | up |
| ENSRNOG00000070742 | 2.907071 | 1.5395662 | 0.009705 | 0.041548 | up |
| Hist1h4m | 2.270765 | 1.1831785 | 0.009812 | 0.041912 | up |
| Fam199x | 0.484912 | -1.044206 | 0.009813 | 0.041912 | down |
| Apbb3 | 2.44378 | 1.2891144 | 0.009845 | 0.042009 | up |
| LOC120098677 | 2.49951 | 1.3216456 | 0.009862 | 0.042067 | up |
| Meltf | 2.372276 | 1.246272 | 0.009939 | 0.042327 | up |
| Chrm1 | 2.339446 | 1.2261666 | 0.009954 | 0.042373 | up |
| Gpbar1 | 0.1884 | -2.408128 | 0.010071 | 0.042778 | down |
| Aim2 | 2.489976 | 1.316132 | 0.010092 | 0.042833 | up |
| Efna3 | 3.456925 | 1.7894895 | 0.010109 | 0.042898 | up |
| LOC120101240 | 2.714757 | 1.4408231 | 0.010129 | 0.04297 | up |
| Gadl1 | 0.459108 | -1.123093 | 0.010259 | 0.043414 | down |
| Tmem145 | 610.806 | 9.2545704 | 0.010314 | 0.043595 | up |
| Mir21 | 3.55188 | 1.8285828 | 0.010321 | 0.043615 | up |
| Nr4a3 | 2.614686 | 1.3866378 | 0.010335 | 0.043666 | up |
| RGD1565410 | 2.095381 | 1.0672129 | 0.010337 | 0.043667 | up |
| Slc36a2 | 13.40457 | 3.7446532 | 0.010485 | 0.044205 | up |
| LOC120099984 | 6.054336 | 2.5979687 | 0.010493 | 0.04422 | up |
| Gpr33-ps1 | 8.324474 | 3.0573591 | 0.010516 | 0.04431 | up |
| LOC120094470 | 2.788663 | 1.4795734 | 0.010589 | 0.044545 | up |
| Rpl10l | 0.472935 | -1.080286 | 0.010608 | 0.044607 | down |
| LOC100359656 | 55.31285 | 5.7895426 | 0.010624 | 0.044649 | up |
| Ly49si3 | 2.002137 | 1.0015406 | 0.010625 | 0.044649 | up |
| ENSRNOG00000065037 | 4.195368 | 2.0687975 | 0.010636 | 0.044684 | up |
| Kyat1 | 0.37722 | -1.40652 | 0.010669 | 0.044791 | down |
| RGD1560775 | 5.57137 | 2.478032 | 0.010745 | 0.045069 | up |
| Nxf7 | 25.62464 | 4.6794596 | 0.010889 | 0.045576 | up |
| Rnase9 | 0.270883 | -1.884256 | 0.010968 | 0.045806 | down |
| Alas1 | 0.475582 | -1.072235 | 0.010989 | 0.045877 | down |
| ENSRNOG00000064415 | 0.000253 | -11.94612 | 0.010994 | 0.045888 | down |
| ENSRNOG00000063439 | 2.436349 | 1.2847208 | 0.011013 | 0.045962 | up |
| Cyp2a3 | 0.156889 | -2.672186 | 0.011017 | 0.045969 | down |
| Spopfm2l1 | 4.134294 | 2.0476411 | 0.011259 | 0.046817 | up |
| Tdrd1 | 394.57 | 8.6241375 | 0.011266 | 0.04683 | up |
| Acer1 | 2.576723 | 1.3655376 | 0.011298 | 0.04694 | up |
| ENSRNOG00000062934 | 9.528654 | 3.2522725 | 0.011303 | 0.046951 | up |
| ENSRNOG00000070735 | 0.495594 | -1.01277 | 0.011333 | 0.047044 | down |
| Dusp15 | 4.289518 | 2.1008156 | 0.011333 | 0.047044 | up |
| S100a3 | 2.633878 | 1.3971884 | 0.011344 | 0.047072 | up |

| | | | | | |
|---|---|---|---|---|---|
| Hoxd9 | 5.440092 | 2.4436311 | 0.011354 | 0.047104 | up |
| Olah | 2.368831 | 1.2441755 | 0.011483 | 0.047514 | up |
| Mdga1 | 2.028228 | 1.0202201 | 0.011499 | 0.047552 | up |
| Hes7 | 2.201 | 1.1381594 | 0.011501 | 0.047552 | up |
| AABR07043115.1 | 15.11136 | 3.9175617 | 0.011628 | 0.047966 | up |
| LOC120097209 | 4.960135 | 2.3103793 | 0.01163 | 0.047966 | up |
| Entrep2 | 2.822529 | 1.4969883 | 0.011772 | 0.048476 | up |
| Rhbdl1 | 2.143004 | 1.0996344 | 0.011869 | 0.048804 | up |
| Snora53 | 2.294041 | 1.1978911 | 0.011897 | 0.048887 | up |
| Tm4sf4 | 5.11762 | 2.355473 | 0.011973 | 0.049144 | up |
| Krt36 | 3.186461 | 1.6719552 | 0.01204 | 0.049307 | up |
| Snord45a | 2.174841 | 1.1209102 | 0.012056 | 0.049362 | up |
| Crnn | 813.8 | 9.6685305 | 0.01211 | 0.049534 | up |
| RGD1308564 | 0.111157 | -3.169324 | 0.012146 | 0.049642 | down |
| Aurkc | 2.028631 | 1.0205067 | 0.012176 | 0.049739 | up |
| Slc23a3 | 2.473905 | 1.30679 | 0.012196 | 0.049788 | up |
| Tex52 | 3.120849 | 1.6419384 | 0.012196 | 0.049788 | up |
| ENSRNOG00000070944 | 0.426749 | -1.228539 | 0.012196 | 0.049788 | down |
| Kel | 0.040358 | -4.631017 | 0.01221 | 0.049835 | down |

**Table 3. Female BLEO vs Male BLEO mRNA-seq**
Significantly Differentially Expressed Genes ($Log_2FC > |1|$; qvalue < 0.05)

| Gene | fc | log₂(fc) | pval | qval | regulation |
|---|---|---|---|---|---|
| Uty | 4.9E-05 | -14.31572 | 2.96E-38 | 6.88E-34 | down |
| AABR07061385.2 | 0.011865 | -6.397128 | 1.06E-23 | 1.23E-19 | down |
| Rgs4 | 0.192572 | -2.376527 | 6.28E-21 | 4.86E-17 | down |
| Uba1y | 0.000242 | -12.01236 | 1.69E-19 | 9.79E-16 | down |
| ENSRNOG00000071081 | 4.62E-05 | -14.40028 | 1.32E-17 | 6.15E-14 | down |
| ENSRNOG00000069947 | 0.033693 | -4.891392 | 5.79E-15 | 2.24E-11 | down |
| ENSRNOG00000069319 | 0.000209 | -12.22562 | 4.21E-14 | 1.1E-10 | down |
| LOC103694557 | 0.000339 | -11.52809 | 4.26E-14 | 1.1E-10 | down |
| Lcmt2 | 1899.192 | 10.89117 | 1.02E-12 | 2.38E-09 | up |
| AABR07053830.1 | 0.001344 | -9.538825 | 1.2E-12 | 2.44E-09 | down |
| Eif2s3y | 0.000675 | -10.53319 | 1.5E-12 | 2.68E-09 | down |
| Bank1 | 4.107096 | 2.0381186 | 2.88E-12 | 4.77E-09 | up |
| Serpina3a | 0.075269 | -3.731808 | 8.7E-12 | 1.35E-08 | down |
| Col7a1 | 0.308033 | -1.698841 | 1.77E-11 | 2.56E-08 | down |
| Btn2a2 | 0.496271 | -1.010801 | 2.25E-11 | 3.08E-08 | down |
| Fmo1 | 2.071227 | 1.0504858 | 4.88E-11 | 6.3E-08 | up |
| Calca | 0.19829 | -2.334315 | 5.33E-11 | 6.52E-08 | down |
| Pcdha5 | 2.778934 | 1.4745316 | 6.99E-11 | 8.12E-08 | up |
| Smad9 | 2.405245 | 1.2661837 | 1.05E-10 | 1.11E-07 | up |
| Rpl10l | 4.488106 | 2.1661068 | 1.51E-10 | 1.53E-07 | up |
| Tmem202 | 0.198175 | -2.335156 | 3.04E-10 | 2.94E-07 | down |
| Gckr | 4.38915 | 2.1339415 | 3.79E-10 | 3.47E-07 | up |
| Cldn4 | 0.245533 | -2.026013 | 6.11E-10 | 5.26E-07 | down |
| Cyp2b1 | 2.347794 | 1.231306 | 6.97E-10 | 5.58E-07 | up |
| Kdm5d | 0.002786 | -8.487429 | 1.66E-09 | 1.29E-06 | down |
| LOC681182 | 2.683037 | 1.4238671 | 1.8E-09 | 1.35E-06 | up |
| Vipr1 | 3.201204 | 1.6786144 | 1.95E-09 | 1.42E-06 | up |
| Apobec1 | 0.222709 | -2.166767 | 2.32E-09 | 1.64E-06 | down |
| Kdr | 3.107549 | 1.6357769 | 2.77E-09 | 1.89E-06 | up |
| Kng1 | 0.258994 | -1.949009 | 3.05E-09 | 2.03E-06 | down |
| Susd2 | 3.815432 | 1.9318465 | 4.06E-09 | 2.62E-06 | up |
| Alpk3 | 2.380005 | 1.2509649 | 5.85E-09 | 3.61E-06 | up |
| Cpne7 | 0.201397 | -2.311885 | 5.91E-09 | 3.61E-06 | down |
| Upk3bl1 | 0.103494 | -3.272385 | 8.52E-09 | 5.06E-06 | down |
| Cct6b | 4.294487 | 2.1024857 | 8.72E-09 | 5.06E-06 | up |
| C15h8orf74 | 0.000761 | -10.3597 | 1.09E-08 | 6.18E-06 | down |
| Cysltr2 | 2.24707 | 1.1680448 | 1.34E-08 | 7.42E-06 | up |
| Cyp2b2 | 2.505548 | 1.3251263 | 1.41E-08 | 7.61E-06 | up |

| | | | | | |
|---|---|---|---|---|---|
| Atg9b | 0.108402 | -3.205539 | 1.49E-08 | 7.88E-06 | down |
| ENSRNOG00000037911 | 50.96355 | 5.6713939 | 1.66E-08 | 8.57E-06 | up |
| Phyhipl | 3.684174 | 1.8813412 | 1.71E-08 | 8.61E-06 | up |
| AY172581.19 | 0.496346 | -1.010581 | 2.6E-08 | 1.26E-05 | down |
| Hpx | 0.222436 | -2.168537 | 2.88E-08 | 1.37E-05 | down |
| Asgr2 | 0.224934 | -2.152425 | 3.47E-08 | 1.61E-05 | down |
| Rem1 | 0.357252 | -1.484985 | 6.79E-08 | 2.98E-05 | down |
| ENSRNOG00000065796 | 2.136605 | 1.0953202 | 7.1E-08 | 3.05E-05 | up |
| C1qc | 0.343765 | -1.540504 | 7.85E-08 | 3.26E-05 | down |
| Cst6 | 0.261948 | -1.93265 | 7.92E-08 | 3.26E-05 | down |
| Slc5a1 | 4.028086 | 2.0100945 | 9.18E-08 | 3.67E-05 | up |
| Kcnn3 | 3.081195 | 1.6234899 | 9.61E-08 | 3.69E-05 | up |
| Ankrd13d | 0.476694 | -1.068865 | 9.7E-08 | 3.69E-05 | down |
| ENSRNOG00000067297 | 3.688502 | 1.8830352 | 1.07E-07 | 4.01E-05 | up |
| C1qa | 0.294436 | -1.763972 | 1.19E-07 | 4.34E-05 | down |
| Hspa12a | 2.006881 | 1.0049548 | 1.2E-07 | 4.34E-05 | up |
| Fam110c | 0.394385 | -1.342325 | 1.27E-07 | 4.46E-05 | down |
| 4933403O08Rik | 2307.565 | 11.172156 | 1.37E-07 | 4.68E-05 | up |
| ENSRNOG00000069272 | 0.000524 | -10.89905 | 1.56E-07 | 5.1E-05 | down |
| Rnf112 | 0.104331 | -3.260763 | 1.56E-07 | 5.1E-05 | down |
| Resp18 | 0.237556 | -2.073663 | 1.65E-07 | 5.33E-05 | down |
| Rnf152 | 2.437581 | 1.28545 | 2.17E-07 | 6.67E-05 | up |
| C1qb | 0.342181 | -1.547168 | 2.18E-07 | 6.67E-05 | down |
| Ntrk2 | 0.345078 | -1.535004 | 2.35E-07 | 6.98E-05 | down |
| Vnn1 | 2.40777 | 1.2676978 | 2.41E-07 | 7.1E-05 | up |
| Capn8 | 0.108477 | -3.204537 | 2.46E-07 | 7.15E-05 | down |
| Prss22 | 0.233343 | -2.099478 | 2.69E-07 | 7.73E-05 | down |
| Ssc5d | 0.488506 | -1.033551 | 3E-07 | 8.5E-05 | down |
| LOC681325 | 2.669961 | 1.4168189 | 3.11E-07 | 8.7E-05 | up |
| Adm2 | 0.195823 | -2.352378 | 3.18E-07 | 8.78E-05 | down |
| Flrt3 | 2.830854 | 1.5012373 | 3.59E-07 | 9.6E-05 | up |
| Scg2 | 0.167169 | -2.580621 | 3.67E-07 | 9.6E-05 | down |
| Degs2 | 0.36344 | -1.46021 | 3.7E-07 | 9.6E-05 | down |
| Plppr3 | 0.386077 | -1.373041 | 3.71E-07 | 9.6E-05 | down |
| Rtkn2 | 3.232311 | 1.692566 | 3.72E-07 | 9.6E-05 | up |
| Prss30 | 0.162311 | -2.623169 | 4.03E-07 | 0.000103 | down |
| Ddx3 | 0.001852 | -9.077073 | 4.53E-07 | 0.000113 | down |
| Slc27a3 | 0.402627 | -1.312483 | 5.6E-07 | 0.000137 | down |
| Cpt1c | 0.447224 | -1.160931 | 5.71E-07 | 0.000138 | down |
| Noxa1 | 0.25284 | -1.983703 | 5.85E-07 | 0.000139 | down |
| Rgs6 | 2.275805 | 1.1863768 | 5.86E-07 | 0.000139 | up |

| | | | | | |
|---|---|---|---|---|---|
| Dnm1 | 0.375842 | -1.411804 | 6.33E-07 | 0.000147 | down |
| Cryaa | 0.000545 | -10.84184 | 6.44E-07 | 0.000147 | down |
| B4galnt4 | 0.392455 | -1.349402 | 6.46E-07 | 0.000147 | down |
| Ubash3b | 2.012005 | 1.0086342 | 6.67E-07 | 0.00015 | up |
| Arhgef26 | 2.196871 | 1.1354501 | 6.73E-07 | 0.00015 | up |
| Klf8 | 3.544344 | 1.8255186 | 6.91E-07 | 0.000153 | up |
| Cdkl5 | 4.233149 | 2.0817312 | 7.1E-07 | 0.000156 | up |
| Clec10a | 0.362989 | -1.462002 | 7.24E-07 | 0.000157 | down |
| Mlana | 0.000412 | -11.2457 | 8.29E-07 | 0.000178 | down |
| Ces2c | 0.213605 | -2.226982 | 8.47E-07 | 0.000179 | down |
| Esm1 | 3.502547 | 1.8084043 | 8.48E-07 | 0.000179 | up |
| Srd5a2 | 0.096692 | -3.370465 | 8.56E-07 | 0.000179 | down |
| Cavin2 | 2.203777 | 1.1399783 | 8.67E-07 | 0.00018 | up |
| Matn3 | 4.139286 | 2.0493819 | 9.2E-07 | 0.000189 | up |
| Il27ra | 0.054306 | -4.202734 | 9.84E-07 | 0.0002 | down |
| Il24 | 0.20433 | -2.291029 | 1.2E-06 | 0.000238 | down |
| Sstr1 | 3.851289 | 1.9453412 | 1.2E-06 | 0.000238 | up |
| Cxcl10 | 0.313842 | -1.67189 | 1.2E-06 | 0.000238 | down |
| Plekhh2 | 2.25138 | 1.1708097 | 1.26E-06 | 0.000249 | up |
| Slc36a2 | 0.111035 | -3.170914 | 1.49E-06 | 0.000286 | down |
| Lcn3 | 0.00096 | -10.02403 | 1.51E-06 | 0.000286 | down |
| Slc16a12 | 3.337651 | 1.7388331 | 1.52E-06 | 0.000286 | up |
| Edil3 | 2.793931 | 1.4822965 | 1.54E-06 | 0.000288 | up |
| Itgb3 | 2.367773 | 1.243531 | 1.55E-06 | 0.000288 | up |
| Susd5 | 0.19296 | -2.373623 | 1.6E-06 | 0.000293 | down |
| Magi3 | 2.318863 | 1.2134176 | 1.6E-06 | 0.000293 | up |
| Lrrc3b | 0.209328 | -2.256162 | 1.7E-06 | 0.000306 | down |
| Acoxl | 5.064944 | 2.3405463 | 1.73E-06 | 0.000309 | up |
| Dusp8 | 2.534829 | 1.3418885 | 1.8E-06 | 0.000314 | up |
| Bmp15 | 6.234649 | 2.6403083 | 1.8E-06 | 0.000314 | up |
| Flt1 | 2.445747 | 1.2902751 | 1.84E-06 | 0.000319 | up |
| Sirpb3 | 5.315326 | 2.4101582 | 1.88E-06 | 0.000322 | up |
| Vxn | 0.289388 | -1.788924 | 1.93E-06 | 0.000326 | down |
| Vmo1 | 0.212869 | -2.231961 | 2.11E-06 | 0.000345 | down |
| Mchr1 | 0.042739 | -4.548304 | 2.17E-06 | 0.000352 | down |
| Lsmem2 | 2.900156 | 1.5361307 | 2.28E-06 | 0.000365 | up |
| AY172581.9 | 0.487445 | -1.036689 | 2.47E-06 | 0.000383 | down |
| Kcnmb2 | 3.954927 | 1.9836511 | 2.52E-06 | 0.000385 | up |
| Nrsn1 | 0.00076 | -10.36233 | 2.57E-06 | 0.000391 | down |
| Ntm | 0.117902 | -3.084344 | 2.62E-06 | 0.000396 | down |
| Usp4 | 2.885804 | 1.5289734 | 2.66E-06 | 0.000396 | up |

| | | | | | |
|---|---|---|---|---|---|
| P2ry6 | 0.397957 | -1.329316 | 2.78E-06 | 0.000412 | down |
| RGD1565058 | 0.000375 | -11.38257 | 2.92E-06 | 0.000427 | down |
| Peli2 | 2.033624 | 1.0240531 | 2.99E-06 | 0.000433 | up |
| ENSRNOG00000064007 | 0.240747 | -2.05441 | 3.04E-06 | 0.000435 | down |
| Enpep | 2.513215 | 1.3295344 | 3.09E-06 | 0.000439 | up |
| AABR07008309.1 | 0.135259 | -2.886207 | 3.19E-06 | 0.00045 | down |
| Aif1 | 0.403346 | -1.309911 | 3.3E-06 | 0.000458 | down |
| Cyp2d5 | 0.330196 | -1.598607 | 3.32E-06 | 0.000458 | down |
| ENSRNOG00000065206 | 0.200121 | -2.321053 | 3.33E-06 | 0.000458 | down |
| Unc5a | 2.447094 | 1.2910697 | 3.4E-06 | 0.000464 | up |
| LOC103691165 | 0.371232 | -1.429605 | 3.53E-06 | 0.000474 | down |
| Tekt5 | 3.653988 | 1.8694717 | 3.87E-06 | 0.00051 | up |
| Smad6 | 2.711562 | 1.4391243 | 3.89E-06 | 0.00051 | up |
| Bmpr2 | 2.592436 | 1.3743085 | 3.98E-06 | 0.00052 | up |
| Map3k7cl | 0.429716 | -1.218543 | 4.19E-06 | 0.000535 | down |
| Rgs10 | 0.485098 | -1.043653 | 4.41E-06 | 0.000557 | down |
| Obp3 | 0.084752 | -3.560603 | 4.63E-06 | 0.000574 | down |
| Mafb | 0.401764 | -1.31558 | 4.75E-06 | 0.00058 | down |
| Tmem271 | 2.919386 | 1.5456648 | 4.78E-06 | 0.000582 | up |
| Tsbp1 | 0.00161 | -9.278411 | 4.9E-06 | 0.000593 | down |
| Cdc42bpg | 2.021468 | 1.0154036 | 5.21E-06 | 0.000619 | up |
| Fcrl2 | 0.214741 | -2.219328 | 5.27E-06 | 0.000621 | down |
| Slco1a4 | 2.904482 | 1.538281 | 5.31E-06 | 0.000623 | up |
| Gucy2e | 0.00277 | -8.496101 | 5.4E-06 | 0.00063 | down |
| Foxn1 | 0.003645 | -8.09971 | 5.56E-06 | 0.000646 | down |
| Xcr1 | 0.218969 | -2.191199 | 5.63E-06 | 0.000647 | down |
| Cfd | 0.168915 | -2.565628 | 5.66E-06 | 0.000648 | down |
| Nrp1 | 2.663435 | 1.413288 | 5.76E-06 | 0.000656 | up |
| Mastl | 2.184591 | 1.1273629 | 6E-06 | 0.00068 | up |
| Ptprv | 0.146184 | -2.774144 | 6.04E-06 | 0.000681 | down |
| Vgf | 0.227326 | -2.137165 | 6.09E-06 | 0.000683 | down |
| Klra5 | 0.142523 | -2.810729 | 6.13E-06 | 0.000684 | down |
| Ncapg | 2.040364 | 1.0288266 | 6.23E-06 | 0.000693 | up |
| ENSRNOG00000063035 | 10211.14 | 13.317857 | 6.46E-06 | 0.000715 | up |
| A2m | 0.204307 | -2.291191 | 6.6E-06 | 0.000727 | down |
| Wwc2 | 2.058933 | 1.041897 | 6.9E-06 | 0.00075 | up |
| Pdzd4 | 0.341907 | -1.548326 | 6.95E-06 | 0.00075 | down |
| Cybrd1 | 2.253348 | 1.1720704 | 6.97E-06 | 0.00075 | up |
| Gulo | 0.040948 | -4.610074 | 6.98E-06 | 0.00075 | down |
| Gldn | 0.340253 | -1.555321 | 7.26E-06 | 0.00077 | down |
| Ptprt | 0.08233 | -3.602439 | 7.55E-06 | 0.000793 | down |

| | | | | | |
|---|---|---|---|---|---|
| LOC120098812 | 0.346589 | -1.528704 | 7.6E-06 | 0.000793 | down |
| Albfm1 | 0.291561 | -1.77813 | 7.66E-06 | 0.000794 | down |
| Kcnk13 | 0.380243 | -1.395006 | 7.85E-06 | 0.000807 | down |
| Gjb5 | 0.287839 | -1.796667 | 7.95E-06 | 0.000812 | down |
| Ms4a4a | 0.490519 | -1.02762 | 7.97E-06 | 0.000812 | down |
| Apln | 3.541189 | 1.8242339 | 8.02E-06 | 0.000813 | up |
| Clic5 | 2.577004 | 1.365695 | 8.14E-06 | 0.00082 | up |
| Ifi27l2b | 0.374995 | -1.415057 | 8.19E-06 | 0.00082 | down |
| Zfp367 | 2.368855 | 1.24419 | 8.29E-06 | 0.000827 | up |
| Il1a | 4.16102 | 2.0569374 | 8.48E-06 | 0.000838 | up |
| Akap5 | 3.579682 | 1.8398315 | 8.63E-06 | 0.00085 | up |
| Tmem40 | 0.435663 | -1.198716 | 8.75E-06 | 0.000852 | down |
| Qprt | 0.375713 | -1.412295 | 8.92E-06 | 0.000863 | down |
| Pilrb | 4.305426 | 2.106156 | 9.39E-06 | 0.000905 | up |
| Chrne | 0.343084 | -1.543367 | 9.49E-06 | 0.000911 | down |
| Zdhhc2 | 2.432099 | 1.2822019 | 9.85E-06 | 0.000934 | up |
| Add2 | 4.675678 | 2.2251755 | 1.01E-05 | 0.000947 | up |
| Ednrb | 3.822785 | 1.934624 | 1.01E-05 | 0.000947 | up |
| Bdnf | 3.34698 | 1.74286 | 1.08E-05 | 0.001 | up |
| Col5a3 | 0.284738 | -1.812294 | 1.1E-05 | 0.001018 | down |
| Ankrd29 | 2.229145 | 1.1564905 | 1.18E-05 | 0.001075 | up |
| Cgnl1 | 2.451637 | 1.2937454 | 1.19E-05 | 0.001076 | up |
| Wnt7b | 0.470607 | -1.087405 | 1.19E-05 | 0.001076 | down |
| LOC120093457 | 9.7E-05 | -13.33224 | 1.2E-05 | 0.001081 | down |
| Mcc | 2.203018 | 1.1394814 | 1.31E-05 | 0.001158 | up |
| Fxyd4 | 0.241432 | -2.050314 | 1.37E-05 | 0.001193 | down |
| Atp1b2 | 0.445912 | -1.165168 | 1.45E-05 | 0.001252 | down |
| Ablim3 | 2.240255 | 1.1636633 | 1.46E-05 | 0.001252 | up |
| Epha3 | 2.970603 | 1.5707558 | 1.47E-05 | 0.001258 | up |
| ENSRNOG00000071219 | 0.405946 | -1.30064 | 1.47E-05 | 0.001258 | down |
| Pabir2 | 2.183382 | 1.1265643 | 1.48E-05 | 0.001258 | up |
| Podnl1 | 0.225556 | -2.148444 | 1.54E-05 | 0.001301 | down |
| Phactr1 | 3.443257 | 1.783774 | 1.55E-05 | 0.001301 | up |
| Ly49si2 | 0.341317 | -1.550816 | 1.56E-05 | 0.001308 | down |
| Kcnk4 | 0.272039 | -1.878112 | 1.57E-05 | 0.001309 | down |
| Dll4 | 2.845976 | 1.5089237 | 1.59E-05 | 0.001316 | up |
| Adh6 | 0.429037 | -1.220826 | 1.59E-05 | 0.001316 | down |
| Thy1 | 0.368511 | -1.440222 | 1.68E-05 | 0.001381 | down |
| Ccl6 | 2.231951 | 1.1583057 | 1.76E-05 | 0.001433 | up |
| Ptprz1 | 0.4068 | -1.29761 | 1.77E-05 | 0.001433 | down |
| Aqp3 | 0.14532 | -2.782695 | 1.78E-05 | 0.001433 | down |

| | | | | | |
|---|---|---|---|---|---|
| Cnr1 | 3.576113 | 1.8383924 | 1.81E-05 | 0.001448 | up |
| Tnfrsf22 | 0.386551 | -1.371271 | 1.99E-05 | 0.001569 | down |
| Tnfsf10 | 2.765266 | 1.4674182 | 2E-05 | 0.001573 | up |
| Chst5 | 0.353752 | -1.499188 | 2.01E-05 | 0.001579 | down |
| Atrnl1 | 2.012145 | 1.008734 | 2.03E-05 | 0.001582 | up |
| Col2a1 | 0.001771 | -9.141481 | 2.17E-05 | 0.001658 | down |
| Olfm4 | 0.32205 | -1.634645 | 2.18E-05 | 0.001658 | down |
| Cd177 | 0.21651 | -2.207492 | 2.27E-05 | 0.001714 | down |
| Tnr | 4.194758 | 2.0685876 | 2.29E-05 | 0.00172 | up |
| Celf4 | 0.25869 | -1.950704 | 2.32E-05 | 0.00174 | down |
| Ptprb | 2.059981 | 1.0426311 | 2.35E-05 | 0.00175 | up |
| Tmcc2 | 2.53189 | 1.3402146 | 2.35E-05 | 0.00175 | up |
| Xpnpep2 | 2.37026 | 1.2450454 | 2.38E-05 | 0.001763 | up |
| Egfl6 | 2.931029 | 1.5514072 | 2.39E-05 | 0.001763 | up |
| Bmp6 | 2.098324 | 1.0692373 | 2.39E-05 | 0.001763 | up |
| Pinlyp | 0.181743 | -2.460028 | 2.4E-05 | 0.001765 | down |
| Col9a2 | 0.301366 | -1.730412 | 2.42E-05 | 0.001774 | down |
| Ppp1r16b | 2.038489 | 1.0274999 | 2.47E-05 | 0.001801 | up |
| Agmat | 0.298784 | -1.742824 | 2.73E-05 | 0.001967 | down |
| Foxf1 | 2.420784 | 1.2754745 | 2.75E-05 | 0.001972 | up |
| Cabp1 | 0.34334 | -1.542292 | 2.81E-05 | 0.002008 | down |
| Jakmip3 | 0.162378 | -2.62257 | 2.87E-05 | 0.002034 | down |
| Nxph3 | 0.358344 | -1.480585 | 2.87E-05 | 0.002034 | down |
| Pdlim4 | 0.358236 | -1.481017 | 2.92E-05 | 0.002052 | down |
| Enpp6 | 3.97884 | 1.9923479 | 2.96E-05 | 0.002063 | up |
| Sytl3 | 0.454627 | -1.137243 | 2.96E-05 | 0.002063 | down |
| Ms4a6a | 0.493283 | -1.019512 | 2.98E-05 | 0.002063 | down |
| Chrdl2 | 0.211346 | -2.242319 | 2.98E-05 | 0.002063 | down |
| Tram2 | 2.602327 | 1.379802 | 2.99E-05 | 0.002063 | up |
| Lpal2 | 0.00075 | -10.38073 | 2.99E-05 | 0.002063 | down |
| Kcna2 | 0.242137 | -2.046106 | 3.06E-05 | 0.002098 | down |
| Cdhr4 | 0.365755 | -1.45105 | 3.18E-05 | 0.002139 | down |
| Rpp14 | 5.149646 | 2.3644733 | 3.19E-05 | 0.002141 | up |
| Mettl24 | 2.172926 | 1.1196393 | 3.2E-05 | 0.002141 | up |
| LOC687707 | 0.249939 | -2.000353 | 3.22E-05 | 0.002141 | down |
| Eng | 2.313825 | 1.2102795 | 3.23E-05 | 0.002141 | up |
| Bend7 | 2.40813 | 1.2679132 | 3.24E-05 | 0.002141 | up |
| Pld4 | 0.484805 | -1.044522 | 3.27E-05 | 0.002141 | down |
| Sema6a | 2.008557 | 1.0061595 | 3.27E-05 | 0.002141 | up |
| Asic1 | 2.180887 | 1.1249147 | 3.33E-05 | 0.002166 | up |
| Cxcl13 | 0.203352 | -2.297949 | 3.36E-05 | 0.002174 | down |

| | | | | | |
|---|---|---|---|---|---|
| Itga1 | 3.113486 | 1.6385309 | 3.42E-05 | 0.00221 | up |
| ENSRNOG00000062648 | 0.234284 | -2.093672 | 3.48E-05 | 0.002236 | down |
| Tmprss9 | 0.276934 | -1.852388 | 3.49E-05 | 0.002238 | down |
| RGD1564162 | 0.380109 | -1.395514 | 3.5E-05 | 0.002239 | down |
| Kntc1 | 2.026475 | 1.0189723 | 3.53E-05 | 0.002247 | up |
| Tecta | 0.0053 | -7.559782 | 3.61E-05 | 0.002286 | down |
| Amigo2 | 2.476406 | 1.3082478 | 3.66E-05 | 0.002305 | up |
| Cyp2c12 | 0.000657 | -10.57231 | 3.87E-05 | 0.002402 | down |
| Tnn | 0.175457 | -2.510812 | 3.96E-05 | 0.002433 | down |
| Pglyrp2 | 0.252739 | -1.984281 | 4.02E-05 | 0.002467 | down |
| Gcgr | 0.282698 | -1.822667 | 4.05E-05 | 0.002472 | down |
| Bmp8a | 5.171323 | 2.3705334 | 4.07E-05 | 0.002472 | up |
| Naaa | 2.154898 | 1.1076194 | 4.11E-05 | 0.002484 | up |
| Ptprn | 0.259489 | -1.946254 | 4.12E-05 | 0.002487 | down |
| RT1-Db1 | 0.409181 | -1.289189 | 4.14E-05 | 0.002491 | down |
| AABR07044631.2 | 0.440366 | -1.183226 | 4.25E-05 | 0.002538 | down |
| Trarg1 | 0.338062 | -1.564641 | 4.29E-05 | 0.002555 | down |
| Prss23 | 2.236664 | 1.1613488 | 4.38E-05 | 0.002585 | up |
| Itga10 | 0.263477 | -1.924249 | 4.63E-05 | 0.002711 | down |
| Pygo1 | 2.062181 | 1.0441712 | 5.06E-05 | 0.002929 | up |
| Ninj1 | 0.49119 | -1.025646 | 5.19E-05 | 0.002983 | down |
| LOC120099971 | 0.309601 | -1.691519 | 5.21E-05 | 0.002991 | down |
| Tgfbi | 0.470266 | -1.08845 | 5.3E-05 | 0.003021 | down |
| Plcd4 | 0.41182 | -1.279913 | 5.32E-05 | 0.003021 | down |
| AC112568.1 | 0.284401 | -1.814003 | 5.47E-05 | 0.003078 | down |
| Thrsp | 0.021483 | -5.540688 | 5.71E-05 | 0.003179 | down |
| Tmem26 | 2.116673 | 1.0817982 | 5.91E-05 | 0.003262 | up |
| Dnah17 | 0.489745 | -1.029897 | 6.02E-05 | 0.003298 | down |
| Slc26a10 | 0.165539 | -2.59476 | 6.24E-05 | 0.003379 | down |
| Batf3 | 0.436053 | -1.197423 | 6.28E-05 | 0.003387 | down |
| Radil | 2.184138 | 1.1270637 | 6.39E-05 | 0.003425 | up |
| Gfod1 | 2.190476 | 1.1312444 | 6.41E-05 | 0.003429 | up |
| Kit | 2.28578 | 1.1926867 | 6.64E-05 | 0.003547 | up |
| Rasd2 | 2.558518 | 1.3553085 | 6.73E-05 | 0.003581 | up |
| Nipa1 | 2.098555 | 1.0693963 | 6.89E-05 | 0.003649 | up |
| Sh3rf2 | 2.784946 | 1.4776492 | 6.97E-05 | 0.003681 | up |
| Rab6b | 2.432057 | 1.2821773 | 7.01E-05 | 0.003681 | up |
| Plg | 0.204379 | -2.290679 | 7.03E-05 | 0.003686 | down |
| Apoc1 | 0.141916 | -2.816889 | 7.11E-05 | 0.003709 | down |
| Nlgn2 | 0.438704 | -1.18868 | 7.19E-05 | 0.003737 | down |
| ENSRNOG00000063615 | 77.46604 | 6.275492 | 7.37E-05 | 0.003814 | up |

| | | | | | |
|---|---|---|---|---|---|
| Slco4c1 | 2.924577 | 1.548228 | 7.49E-05 | 0.003839 | up |
| Angptl4 | 0.355321 | -1.492803 | 7.6E-05 | 0.003862 | down |
| Ppef2 | 2.51895 | 1.3328222 | 7.63E-05 | 0.003871 | up |
| Tns1 | 2.413868 | 1.2713466 | 7.66E-05 | 0.003878 | up |
| Ptcra | 0.000952 | -10.03679 | 7.75E-05 | 0.003897 | down |
| Gja3 | 3.719633 | 1.8951604 | 7.79E-05 | 0.00391 | up |
| Cyp2b21 | 0.000789 | -10.30748 | 8.07E-05 | 0.004003 | down |
| Clec14a | 2.339117 | 1.2259642 | 8.17E-05 | 0.004022 | up |
| Kcnj4 | 2.508267 | 1.3266907 | 8.45E-05 | 0.004097 | up |
| Fendrr | 2.612917 | 1.3856615 | 8.45E-05 | 0.004097 | up |
| Lta | 0.106054 | -3.237123 | 8.57E-05 | 0.004138 | down |
| Dgki | 2.717594 | 1.4423302 | 8.73E-05 | 0.004188 | up |
| Atp10b | 3.161254 | 1.660497 | 8.75E-05 | 0.004188 | up |
| Pmfbp1 | 0.276983 | -1.852132 | 8.76E-05 | 0.004188 | down |
| AY172581.22 | 0.471402 | -1.08497 | 8.95E-05 | 0.004251 | down |
| Cd72 | 0.255782 | -1.967015 | 8.98E-05 | 0.004252 | down |
| Gabra1 | 3.448578 | 1.7860015 | 9.16E-05 | 0.004299 | up |
| Krt42 | 0.0007 | -10.48059 | 9.23E-05 | 0.004315 | down |
| Prkce | 2.573333 | 1.3636379 | 9.33E-05 | 0.004352 | up |
| Plvap | 3.052885 | 1.6101735 | 9.4E-05 | 0.004373 | up |
| Adrb3 | 0.056971 | -4.133616 | 9.43E-05 | 0.00438 | down |
| Trem2 | 0.305008 | -1.713079 | 9.5E-05 | 0.004404 | down |
| Cd8b | 0.341137 | -1.551577 | 9.57E-05 | 0.004426 | down |
| Rmi2 | 2.049236 | 1.035086 | 9.59E-05 | 0.004427 | up |
| Lgals3bp | 0.479357 | -1.060828 | 9.61E-05 | 0.004428 | down |
| Wnk3 | 2.959885 | 1.5655411 | 9.63E-05 | 0.004428 | up |
| Cps1 | 0.159832 | -2.64537 | 9.73E-05 | 0.004467 | down |
| Map2 | 3.382814 | 1.758224 | 9.83E-05 | 0.004484 | up |
| Gphn | 6.780007 | 2.7612868 | 9.94E-05 | 0.004522 | up |
| F2 | 0.357101 | -1.485595 | 9.97E-05 | 0.004522 | down |
| LOC108348213 | 0.390333 | -1.357222 | 0.0001 | 0.004536 | down |
| Tnnt1 | 0.303134 | -1.721972 | 0.000101 | 0.004551 | down |
| Fam229a | 0.326663 | -1.614126 | 0.000101 | 0.004557 | down |
| Wnt3 | 0.243055 | -2.040644 | 0.000102 | 0.004596 | down |
| Pck1 | 0.038418 | -4.702073 | 0.000103 | 0.004607 | down |
| Ace | 2.159401 | 1.110631 | 0.000103 | 0.004607 | up |
| Kl | 2.071034 | 1.0503514 | 0.000106 | 0.00471 | up |
| Twist1 | 0.342495 | -1.545845 | 0.000106 | 0.004734 | down |
| AY172581.18 | 0.459917 | -1.120555 | 0.000108 | 0.004798 | down |
| Sdc3 | 2.079899 | 1.0565133 | 0.000109 | 0.004816 | up |
| Zfp462 | 2.182721 | 1.1261275 | 0.00011 | 0.004851 | up |

| | | | | | |
|---|---|---|---|---|---|
| Sema3e | 2.695204 | 1.4303944 | 0.000111 | 0.004856 | up |
| LOC100909964 | 2.762261 | 1.4658494 | 0.000115 | 0.005014 | up |
| Hba-a3 | 3.986016 | 1.9949475 | 0.000115 | 0.005023 | up |
| Slc18a1 | 0.000985 | -9.987423 | 0.000119 | 0.005146 | down |
| Limch1 | 2.558607 | 1.3553587 | 0.00012 | 0.005179 | up |
| Slc2a4 | 0.250318 | -1.998164 | 0.000121 | 0.005183 | down |
| Pcdh12 | 2.095265 | 1.0671327 | 0.000121 | 0.005183 | up |
| Amotl1 | 2.14731 | 1.1025303 | 0.000123 | 0.005257 | up |
| F10 | 0.269795 | -1.890066 | 0.000126 | 0.005386 | down |
| Plxdc1 | 0.443918 | -1.171635 | 0.000132 | 0.005565 | down |
| Exph5 | 2.46371 | 1.3008326 | 0.000135 | 0.005603 | up |
| Csrnp3 | 2.378343 | 1.249957 | 0.000135 | 0.005603 | up |
| Hdx | 2.607317 | 1.3825662 | 0.000135 | 0.005617 | up |
| Kif14 | 2.146342 | 1.1018798 | 0.000136 | 0.005639 | up |
| Lyzl4 | 0.291544 | -1.778216 | 0.000136 | 0.005639 | down |
| Rarres1 | 0.313631 | -1.672858 | 0.000137 | 0.005639 | down |
| C1qtnf6 | 0.445473 | -1.16659 | 0.000137 | 0.005645 | down |
| Htr7 | 0.323465 | -1.628317 | 0.000137 | 0.005646 | down |
| Ankrd1 | 3.74968 | 1.9067673 | 0.000139 | 0.00568 | up |
| Ly49s6 | 0.237038 | -2.076809 | 0.000139 | 0.00568 | down |
| Shroom4 | 2.970158 | 1.5705396 | 0.000141 | 0.005695 | up |
| Thsd4 | 2.273279 | 1.1847748 | 0.000141 | 0.005695 | up |
| Rps7-ps23 | 0.485074 | -1.043722 | 0.000142 | 0.005721 | down |
| AABR07021213.1 | 0.062411 | -4.002065 | 0.000143 | 0.00573 | down |
| Ptprn2 | 0.411717 | -1.280276 | 0.000145 | 0.005785 | down |
| Ptprg | 2.107681 | 1.0756563 | 0.000145 | 0.005785 | up |
| Tnfrsf9 | 0.419402 | -1.253594 | 0.00015 | 0.005944 | down |
| Duox2 | 0.189598 | -2.398984 | 0.00015 | 0.005944 | down |
| Stoml3 | 0.247088 | -2.016902 | 0.00015 | 0.005944 | down |
| Mslnl | 0.407508 | -1.295098 | 0.000151 | 0.005944 | down |
| Matk | 0.322338 | -1.633355 | 0.000151 | 0.005957 | down |
| Nefh | 0.141453 | -2.821602 | 0.000152 | 0.005969 | down |
| Mapk8ip2 | 0.300484 | -1.734641 | 0.000152 | 0.005971 | down |
| Arhgap31 | 2.053599 | 1.0381542 | 0.000153 | 0.005982 | up |
| Tmem86a | 0.359176 | -1.477236 | 0.000153 | 0.005988 | down |
| Cldn9 | 0.332488 | -1.588627 | 0.000154 | 0.005996 | down |
| ENSRNOG00000066091 | 0.130591 | -2.936876 | 0.000154 | 0.006017 | down |
| Bmp3 | 2.397515 | 1.2615396 | 0.000155 | 0.006034 | up |
| Ly49i7 | 0.053898 | -4.213622 | 0.000157 | 0.006096 | down |
| Cpxm1 | 0.323099 | -1.62995 | 0.00016 | 0.006172 | down |
| Tmem100 | 2.820204 | 1.4957993 | 0.000162 | 0.00621 | up |

| | | | | | |
|---|---|---|---|---|---|
| Sds | 0.441822 | -1.178462 | 0.000162 | 0.00621 | down |
| Cxcl11 | 0.240601 | -2.055285 | 0.000164 | 0.006221 | down |
| Col4a4 | 2.800454 | 1.485661 | 0.000164 | 0.006221 | up |
| Kirrel3 | 0.394459 | -1.342052 | 0.000165 | 0.006221 | down |
| Cpn2 | 0.001083 | -9.850209 | 0.000165 | 0.006221 | down |
| Cyp2d3 | 0.367183 | -1.445431 | 0.000165 | 0.006221 | down |
| Cyp2a2 | 0.041797 | -4.580459 | 0.000166 | 0.006221 | down |
| Nkd1 | 2.136578 | 1.0953021 | 0.000167 | 0.006248 | up |
| Hipk2 | 2.687535 | 1.4262837 | 0.00017 | 0.006339 | up |
| Plxna2 | 2.323275 | 1.2161599 | 0.000172 | 0.006384 | up |
| Lilrb4 | 0.312692 | -1.677188 | 0.000175 | 0.006457 | down |
| RT1-DOa | 0.422781 | -1.242019 | 0.000179 | 0.006573 | down |
| Erbb4 | 4.193814 | 2.068263 | 0.000183 | 0.00667 | up |
| Lox | 2.810836 | 1.4909993 | 0.000186 | 0.00675 | up |
| Clec4a3 | 0.393784 | -1.344523 | 0.000187 | 0.006764 | down |
| Gnaq | 2.025727 | 1.0184399 | 0.000192 | 0.006908 | up |
| Msc | 0.47128 | -1.085344 | 0.000199 | 0.007105 | down |
| Cyth4 | 0.489076 | -1.031869 | 0.0002 | 0.007125 | down |
| Mia | 0.400457 | -1.320279 | 0.000205 | 0.007253 | down |
| Ebi3 | 0.318472 | -1.650762 | 0.000207 | 0.007308 | down |
| Rab37 | 0.496579 | -1.009905 | 0.000207 | 0.007308 | down |
| Ckap2l | 2.269713 | 1.1825097 | 0.000213 | 0.007452 | up |
| Tlcd2 | 2.926608 | 1.5492297 | 0.000214 | 0.007471 | up |
| Hif3a | 0.036648 | -4.770117 | 0.000222 | 0.007681 | down |
| ENSRNOG00000069490 | 0.335687 | -1.574812 | 0.000225 | 0.007757 | down |
| Cdkn2b | 2.535753 | 1.342414 | 0.000228 | 0.007819 | up |
| ENSRNOG00000069186 | 0.140073 | -2.835745 | 0.000229 | 0.007843 | down |
| Lmo3 | 0.398945 | -1.325738 | 0.000231 | 0.007886 | down |
| Shh | 2.523997 | 1.33571 | 0.000233 | 0.007925 | up |
| Cyp3a18 | 0.159095 | -2.652036 | 0.000234 | 0.007925 | down |
| Tfr2 | 0.285011 | -1.81091 | 0.000234 | 0.007925 | down |
| RGD1309808 | 0.347607 | -1.524471 | 0.000235 | 0.007925 | down |
| Ankk1 | 0.154467 | -2.69463 | 0.000235 | 0.007925 | down |
| Cilp | 0.454237 | -1.138482 | 0.000236 | 0.007925 | down |
| Zbp1 | 0.307584 | -1.700948 | 0.000236 | 0.007925 | down |
| Serf1 | 0.473145 | -1.079646 | 0.000239 | 0.007996 | down |
| AY172581.10 | 0.434559 | -1.202375 | 0.000241 | 0.00804 | down |
| Pilra | 2.059664 | 1.0424087 | 0.000242 | 0.00804 | up |
| Shisal1 | 0.377419 | -1.405762 | 0.000242 | 0.00804 | down |
| Il2ra | 0.447323 | -1.160612 | 0.000243 | 0.00804 | down |
| Myh13 | 2.472945 | 1.3062304 | 0.000249 | 0.00817 | up |

| | | | | | |
|---|---|---|---|---|---|
| Angptl8 | 0.322308 | -1.63349 | 0.000249 | 0.00817 | down |
| Pla2g4f | 0.31876 | -1.64946 | 0.000249 | 0.00817 | down |
| Cxcr3 | 0.472347 | -1.082081 | 0.00025 | 0.008191 | down |
| Rapgef4 | 2.399133 | 1.2625134 | 0.000253 | 0.008265 | up |
| Btbd3 | 2.189989 | 1.1309235 | 0.000258 | 0.008364 | up |
| Entpd8 | 0.325415 | -1.619649 | 0.000258 | 0.008367 | down |
| Slc37a2 | 0.463312 | -1.109943 | 0.000262 | 0.008396 | down |
| Cd163 | 0.367854 | -1.442795 | 0.000264 | 0.008446 | down |
| Pate5 | 3.207252 | 1.6813375 | 0.000265 | 0.008484 | up |
| ENSRNOG00000070500 | 0.224059 | -2.158047 | 0.000268 | 0.008535 | down |
| Gdf3 | 0.179441 | -2.478416 | 0.000272 | 0.008609 | down |
| C3ar1 | 0.385046 | -1.376896 | 0.000278 | 0.008728 | down |
| Fut7 | 0.310173 | -1.688854 | 0.000278 | 0.008728 | down |
| Lgals5 | 6.227701 | 2.6386996 | 0.000281 | 0.008782 | up |
| Pdzd2 | 2.312921 | 1.2097159 | 0.000282 | 0.00879 | up |
| Rgs13 | 2.453383 | 1.2947725 | 0.000289 | 0.008941 | up |
| Cdr2 | 2.119039 | 1.08341 | 0.00029 | 0.008956 | up |
| RGD1307182 | 0.313366 | -1.674078 | 0.000294 | 0.009042 | down |
| Slitrk6 | 0.304056 | -1.71759 | 0.000296 | 0.009073 | down |
| Itgad | 0.388823 | -1.362814 | 0.000296 | 0.009073 | down |
| Catsperd | 0.463958 | -1.107934 | 0.000302 | 0.00925 | down |
| Cdkn1a | 0.486948 | -1.038159 | 0.000303 | 0.009258 | down |
| Tmem38a | 0.475597 | -1.072188 | 0.000309 | 0.009376 | down |
| Hpgd | 2.058062 | 1.0412863 | 0.000312 | 0.009414 | up |
| Fap | 0.423031 | -1.241166 | 0.000313 | 0.009447 | down |
| Ppp1r1a | 0.339697 | -1.557681 | 0.000324 | 0.009691 | down |
| Comp | 0.320857 | -1.639998 | 0.000325 | 0.009707 | down |
| Tp63 | 0.199998 | -2.321943 | 0.000328 | 0.009789 | down |
| Ppil6 | 0.427307 | -1.226653 | 0.000335 | 0.009931 | down |
| Mrps6 | 0.428847 | -1.221464 | 0.000336 | 0.009938 | down |
| LOC108352909 | 0.490079 | -1.028914 | 0.000337 | 0.009938 | down |
| Mmp2 | 0.498622 | -1.003981 | 0.000338 | 0.009938 | down |
| Tm6sf2 | 0.487919 | -1.035286 | 0.00034 | 0.009959 | down |
| ENSRNOG00000066155 | 0.427572 | -1.225761 | 0.000343 | 0.010042 | down |
| Snap25 | 0.058323 | -4.099785 | 0.000349 | 0.010192 | down |
| Psapl1 | 0.132724 | -2.9135 | 0.000352 | 0.010211 | down |
| Npr3 | 3.677359 | 1.8786699 | 0.000353 | 0.010238 | up |
| Itih4 | 0.342437 | -1.546089 | 0.000356 | 0.010301 | down |
| Smim5 | 0.312271 | -1.679128 | 0.000362 | 0.010432 | down |
| Tead1 | 2.099195 | 1.0698363 | 0.000367 | 0.010512 | up |
| Gpr156 | 0.288277 | -1.79447 | 0.00037 | 0.010541 | down |

| | | | | | |
|---|---|---|---|---|---|
| Pcdh17 | 2.355505 | 1.2360364 | 0.000377 | 0.010696 | up |
| Wfdc8 | 4.285235 | 2.0993744 | 0.000381 | 0.010779 | up |
| LOC100361226 | 0.073011 | -3.775751 | 0.000383 | 0.010798 | down |
| Clec4a2 | 0.359908 | -1.474298 | 0.000387 | 0.010857 | down |
| Cadm1 | 2.603821 | 1.3806303 | 0.000389 | 0.010868 | up |
| Rasgrf1 | 0.39341 | -1.345895 | 0.000394 | 0.010998 | down |
| Arxes2 | 0.111387 | -3.166346 | 0.0004 | 0.011102 | down |
| N4bp2l1 | 0.447512 | -1.16 | 0.000403 | 0.01113 | down |
| Rab27b | 2.833474 | 1.502572 | 0.000411 | 0.011268 | up |
| Adgrg5 | 0.402484 | -1.312997 | 0.000418 | 0.011386 | down |
| Bard1 | 2.02796 | 1.0200293 | 0.000425 | 0.011491 | up |
| Rgs9bp | 8.661691 | 3.1146488 | 0.000425 | 0.011491 | up |
| Lynx1 | 0.304099 | -1.717387 | 0.000429 | 0.011584 | down |
| ENSRNOG00000068207 | 3.25738 | 1.7037119 | 0.000433 | 0.011632 | up |
| RT1-DMa | 0.483286 | -1.04905 | 0.000441 | 0.011769 | down |
| Clec4a | 0.418653 | -1.256175 | 0.000444 | 0.011822 | down |
| LOC120103148 | 2.591122 | 1.3735771 | 0.000444 | 0.011822 | up |
| ENSRNOG00000063513 | 0.442184 | -1.177282 | 0.000447 | 0.011881 | down |
| Zbtb41 | 2.554436 | 1.3530048 | 0.000451 | 0.011943 | up |
| ENSRNOG00000071156 | 0.00043 | -11.18205 | 0.000469 | 0.012325 | down |
| Spire2 | 0.499803 | -1.000568 | 0.000481 | 0.012551 | down |
| Blnk | 0.449474 | -1.153689 | 0.000483 | 0.012566 | down |
| Esco2 | 2.23469 | 1.1600749 | 0.000484 | 0.012566 | up |
| Tmem273 | 0.417025 | -1.261795 | 0.000487 | 0.012631 | down |
| Clstn2 | 2.195993 | 1.1348733 | 0.000492 | 0.0127 | up |
| Tmem151b | 0.36433 | -1.456682 | 0.000495 | 0.012701 | down |
| Gal3st3 | 2.256138 | 1.1738555 | 0.000495 | 0.012701 | up |
| Sema3g | 2.292994 | 1.1972326 | 0.000496 | 0.012701 | up |
| Calcrl | 2.343253 | 1.228513 | 0.000496 | 0.012701 | up |
| Ppfia3 | 0.447894 | -1.158772 | 0.000505 | 0.01289 | down |
| Megf9 | 2.28089 | 1.1895968 | 0.000512 | 0.012975 | up |
| Ugt2b15 | 0.084567 | -3.563755 | 0.000513 | 0.012991 | down |
| Rhbdl1 | 0.380283 | -1.394854 | 0.000524 | 0.013183 | down |
| Cnih2 | 0.461755 | -1.114802 | 0.00053 | 0.013306 | down |
| Sfn | 0.359997 | -1.473942 | 0.000538 | 0.013457 | down |
| Akr1c3 | 0.045401 | -4.461124 | 0.000549 | 0.013687 | down |
| Pstpip1 | 0.486467 | -1.039587 | 0.000556 | 0.013821 | down |
| Wnt7a | 2.424311 | 1.2775747 | 0.000557 | 0.01383 | up |
| Tmem270 | 0.330725 | -1.596298 | 0.000557 | 0.01383 | down |
| Neurl1 | 0.445316 | -1.167099 | 0.000568 | 0.013984 | down |
| Spock2 | 2.295622 | 1.198885 | 0.000577 | 0.0141 | up |

| | | | | | |
|---|---|---|---|---|---|
| Hs3st3a1 | 2.19589 | 1.1348061 | 0.000577 | 0.0141 | up |
| Apon | 0.000339 | -11.52709 | 0.000578 | 0.014113 | down |
| Bmal2 | 0.246949 | -2.017714 | 0.000581 | 0.014152 | down |
| Avil | 2.517544 | 1.3320171 | 0.000585 | 0.014211 | up |
| Apoe | 0.173672 | -2.525564 | 0.000587 | 0.014238 | down |
| Tmem64 | 2.223256 | 1.1526739 | 0.00059 | 0.014267 | up |
| Stxbp6 | 2.866302 | 1.5191904 | 0.000594 | 0.014331 | up |
| Pla2g10 | 0.307526 | -1.70122 | 0.000596 | 0.014361 | down |
| RT1-Bb | 0.396635 | -1.334115 | 0.000598 | 0.014393 | down |
| AY172581.7 | 0.442385 | -1.176627 | 0.000602 | 0.014459 | down |
| Npy4r | 2.362898 | 1.2405572 | 0.000615 | 0.014735 | up |
| Dlk2 | 0.072385 | -3.78816 | 0.000617 | 0.014757 | down |
| Ubap1l | 0.449178 | -1.154641 | 0.00062 | 0.014776 | down |
| Hnf4a | 0.159682 | -2.646725 | 0.000621 | 0.014784 | down |
| ENSRNOG00000068563 | 47.71544 | 5.5763842 | 0.000623 | 0.014787 | up |
| Itga9 | 2.029597 | 1.0211934 | 0.000626 | 0.014837 | up |
| Timp1 | 0.421457 | -1.246543 | 0.000627 | 0.014837 | down |
| Acer2 | 2.19374 | 1.1333926 | 0.000628 | 0.014837 | up |
| Rsad2 | 0.393686 | -1.344881 | 0.000629 | 0.014837 | down |
| Plin4 | 0.118933 | -3.071776 | 0.00063 | 0.014837 | down |
| Tppp | 2.365119 | 1.2419126 | 0.000636 | 0.014941 | up |
| ENSRNOG00000063337 | 0.357769 | -1.482899 | 0.000643 | 0.015047 | down |
| Piezo2 | 2.16873 | 1.1168505 | 0.000644 | 0.015047 | up |
| RGD1563400 | 0.469743 | -1.090056 | 0.000645 | 0.015047 | down |
| Cd80 | 0.425537 | -1.232645 | 0.000647 | 0.015093 | down |
| Fgd2 | 0.490301 | -1.028262 | 0.000657 | 0.015281 | down |
| Ccn2 | 2.347635 | 1.2312081 | 0.000661 | 0.015348 | up |
| Akap2 | 2.271989 | 1.1839557 | 0.000662 | 0.015359 | up |
| Adipoq | 0.039569 | -4.659472 | 0.000665 | 0.015383 | down |
| Igfals | 0.155511 | -2.684908 | 0.000675 | 0.015543 | down |
| Npw | 2.411248 | 1.26978 | 0.000676 | 0.01555 | up |
| ENSRNOG00000070640 | 2.156862 | 1.1089336 | 0.000679 | 0.015601 | up |
| Ucp1 | 0.000948 | -10.04306 | 0.000687 | 0.01573 | down |
| Tnfaip2 | 0.469121 | -1.091967 | 0.000687 | 0.01573 | down |
| Adss1 | 0.380251 | -1.394976 | 0.00069 | 0.015761 | down |
| Irf7 | 0.479068 | -1.061697 | 0.000696 | 0.015872 | down |
| Cdhr5 | 0.372757 | -1.423691 | 0.000712 | 0.016126 | down |
| ENSRNOG00000063568 | 0.258591 | -1.951254 | 0.000712 | 0.016129 | down |
| Itga11 | 2.150143 | 1.1044324 | 0.000713 | 0.016129 | up |
| Prrg4 | 0.473789 | -1.077684 | 0.000721 | 0.016211 | down |
| ENSRNOG00000063706 | 2.234226 | 1.1597749 | 0.00074 | 0.016534 | up |

| | | | | | |
|---|---|---|---|---|---|
| Lrrc15 | 0.32059 | -1.6412 | 0.000744 | 0.016603 | down |
| Serpinf2 | 0.25149 | -1.991427 | 0.000745 | 0.016603 | down |
| Fbxl2 | 0.479177 | -1.06137 | 0.000746 | 0.016622 | down |
| Itprid1 | 2.134146 | 1.0936591 | 0.000751 | 0.016667 | up |
| Lctl | 0.216087 | -2.210319 | 0.000753 | 0.016682 | down |
| Clec4a1 | 0.472454 | -1.081754 | 0.000755 | 0.016682 | down |
| Tcf15 | 0.378656 | -1.401042 | 0.000759 | 0.016687 | down |
| Dbndd1 | 0.351552 | -1.508192 | 0.000764 | 0.016771 | down |
| Timp3 | 2.742527 | 1.4555057 | 0.000765 | 0.016771 | up |
| Padi4 | 0.429264 | -1.220064 | 0.000767 | 0.016788 | down |
| F7 | 0.31418 | -1.670336 | 0.000769 | 0.01681 | down |
| Nlrp12 | 2.635138 | 1.3978785 | 0.000772 | 0.016842 | up |
| Myh7b | 0.373463 | -1.420965 | 0.000776 | 0.016884 | down |
| Hepacam2 | 2.500256 | 1.3220758 | 0.000782 | 0.016952 | up |
| Irf5 | 0.481285 | -1.055036 | 0.000785 | 0.017007 | down |
| Lepr | 2.225562 | 1.1541696 | 0.000786 | 0.01701 | up |
| Ucn2 | 0.131285 | -2.929229 | 0.000787 | 0.01701 | down |
| Slamf9 | 0.34596 | -1.531323 | 0.0008 | 0.017179 | down |
| Aff3 | 2.015636 | 1.0112351 | 0.000805 | 0.017227 | up |
| Mroh7 | 0.221829 | -2.172481 | 0.000825 | 0.017469 | down |
| Aspg | 0.269524 | -1.891512 | 0.000826 | 0.017469 | down |
| Irf8 | 0.447022 | -1.161582 | 0.000829 | 0.017501 | down |
| Pdcd1 | 0.330699 | -1.596408 | 0.00083 | 0.017503 | down |
| Mx1 | 0.477625 | -1.066049 | 0.00084 | 0.017604 | down |
| Camk2a | 2.193443 | 1.1331971 | 0.000847 | 0.017733 | up |
| Usp18 | 0.395819 | -1.337089 | 0.00085 | 0.017746 | down |
| Fbln7 | 0.434721 | -1.20184 | 0.000854 | 0.017793 | down |
| Habp2 | 0.066183 | -3.917396 | 0.000856 | 0.017793 | down |
| ENSRNOG00000065652 | 7.964659 | 2.9936125 | 0.000868 | 0.017894 | up |
| ENSRNOG00000064431 | 0.069564 | -3.845524 | 0.000868 | 0.017894 | down |
| P3h2 | 2.119517 | 1.0837354 | 0.000873 | 0.017953 | up |
| Car4 | 2.581713 | 1.3683289 | 0.000875 | 0.017986 | up |
| Gdf15 | 0.419443 | -1.253453 | 0.000877 | 0.017998 | down |
| Cyp2d2 | 0.131819 | -2.923372 | 0.000886 | 0.018121 | down |
| LOC102557108 | 0.061902 | -4.013865 | 0.000888 | 0.018145 | down |
| Sh2d7 | 2.509824 | 1.3275864 | 0.000897 | 0.018256 | up |
| RT1-Da | 0.494695 | -1.01539 | 0.000899 | 0.01826 | down |
| Lyl1 | 0.458432 | -1.125219 | 0.000901 | 0.018269 | down |
| Efnb2 | 2.452329 | 1.2941527 | 0.000905 | 0.018324 | up |
| Dao | 0.416876 | -1.262308 | 0.000906 | 0.018331 | down |
| LOC500180 | 0.074127 | -3.75385 | 0.000927 | 0.018682 | down |

| | | | | | |
|---|---|---|---|---|---|
| LOC120093064 | 0.344723 | -1.536491 | 0.000935 | 0.018804 | down |
| Slc5a11 | 0.334534 | -1.579777 | 0.000935 | 0.018804 | down |
| Dpy30 | 18.49872 | 4.2093539 | 0.000937 | 0.018804 | up |
| Dnase1l3 | 0.317117 | -1.656914 | 0.000958 | 0.019157 | down |
| Tmem240 | 0.450247 | -1.151212 | 0.000965 | 0.019285 | down |
| Malrd1 | 0.135492 | -2.883716 | 0.000972 | 0.019392 | down |
| Siglec1 | 0.480195 | -1.058308 | 0.000991 | 0.019652 | down |
| Mcemp1 | 2.192285 | 1.1324356 | 0.000993 | 0.019665 | up |
| Serpinb10 | 3.83744 | 1.9401441 | 0.000994 | 0.019665 | up |
| Scin | 2.936371 | 1.5540344 | 0.001 | 0.01974 | up |
| LOC120100725 | 0.419481 | -1.253321 | 0.001 | 0.01974 | down |
| Tnfrsf18 | 0.455261 | -1.135234 | 0.001003 | 0.019775 | down |
| Crxos1 | 22.79643 | 4.5107362 | 0.001009 | 0.019862 | up |
| AABR07063082.1 | 0.490792 | -1.026815 | 0.00101 | 0.019862 | down |
| Syt13 | 0.402948 | -1.311333 | 0.001022 | 0.020023 | down |
| Icam5 | 0.41348 | -1.274109 | 0.001024 | 0.02006 | down |
| Foxr1 | 0.082307 | -3.602845 | 0.001027 | 0.020088 | down |
| Sfxn4 | 0.466912 | -1.098778 | 0.001036 | 0.020168 | down |
| Hemgn | 2.894188 | 1.5331586 | 0.001037 | 0.020168 | up |
| Pou2f3 | 2.425116 | 1.2780537 | 0.001045 | 0.020292 | up |
| Gpr33-ps1 | 0.246544 | -2.020082 | 0.001045 | 0.020292 | down |
| Cd209d | 0.028021 | -5.157324 | 0.001046 | 0.020292 | down |
| Hpd | 0.094387 | -3.405262 | 0.001058 | 0.020431 | down |
| Trim7 | 0.398695 | -1.326644 | 0.001067 | 0.020564 | down |
| Mrln | 0.000924 | -10.08012 | 0.001069 | 0.020582 | down |
| Adcy1 | 2.629703 | 1.3949 | 0.001071 | 0.020604 | up |
| ENSRNOG00000069261 | 0.000292 | -11.74031 | 0.001076 | 0.020689 | down |
| RGD1560559 | 0.061844 | -4.015218 | 0.001088 | 0.020839 | down |
| Cldn2 | 0.27047 | -1.886457 | 0.001088 | 0.020839 | down |
| ENSRNOG00000069237 | 2.463177 | 1.3005201 | 0.001091 | 0.020867 | up |
| S100a5 | 0.184889 | -2.435266 | 0.001097 | 0.020946 | down |
| Mdga1 | 0.452004 | -1.145591 | 0.001129 | 0.021384 | down |
| LOC679368 | 2.63249 | 1.3964282 | 0.001134 | 0.021427 | up |
| Clip3 | 0.493267 | -1.019559 | 0.001143 | 0.021547 | down |
| Folr2 | 0.371384 | -1.429016 | 0.001159 | 0.021779 | down |
| Kcnt1 | 0.318042 | -1.652711 | 0.001161 | 0.021789 | down |
| Hsd17b13 | 0.496802 | -1.009258 | 0.001178 | 0.021991 | down |
| Ctsw | 0.412075 | -1.279022 | 0.001197 | 0.022273 | down |
| H19 | 0.308506 | -1.69663 | 0.001202 | 0.022344 | down |
| Itih3 | 0.184047 | -2.441853 | 0.001208 | 0.02239 | down |
| Nkx2-8 | 4.550017 | 2.185872 | 0.001224 | 0.02259 | up |

| | | | | | |
|---|---|---|---|---|---|
| Tead4 | 2.098688 | 1.0694877 | 0.001224 | 0.02259 | up |
| Vegfa | 2.210487 | 1.1443641 | 0.001253 | 0.02298 | up |
| Nr5a2 | 3.805967 | 1.9282631 | 0.00126 | 0.02309 | up |
| Slc13a2 | 0.441126 | -1.180736 | 0.001265 | 0.023128 | down |
| Cabp2 | 0.057492 | -4.120504 | 0.001291 | 0.023507 | down |
| Adig | 0.216968 | -2.204443 | 0.001294 | 0.023507 | down |
| Ccl17 | 0.4135 | -1.274039 | 0.001297 | 0.023507 | down |
| Cadm4 | 0.369462 | -1.436503 | 0.001298 | 0.023507 | down |
| Foxp3 | 0.339878 | -1.556912 | 0.001298 | 0.023507 | down |
| Fzd4 | 2.146803 | 1.1021901 | 0.001305 | 0.023582 | up |
| Sncg | 0.264057 | -1.921076 | 0.001305 | 0.023582 | down |
| Adora3 | 0.386195 | -1.3726 | 0.001327 | 0.023887 | down |
| Csprs | 3.24503 | 1.6982319 | 0.001335 | 0.023953 | up |
| ENSRNOG00000067561 | 0.386565 | -1.371218 | 0.001343 | 0.024057 | down |
| Kcp | 0.376021 | -1.411114 | 0.001345 | 0.024068 | down |
| Kcnn4 | 0.395981 | -1.336498 | 0.001351 | 0.02415 | down |
| Dbh | 0.428564 | -1.222418 | 0.001356 | 0.024204 | down |
| Kcnv2 | 2.083035 | 1.0586872 | 0.001365 | 0.02432 | up |
| Vom2r4 | 3.829805 | 1.9372709 | 0.001373 | 0.024384 | up |
| Crct1 | 0.070626 | -3.823665 | 0.001373 | 0.024384 | down |
| Cxcr6 | 0.468908 | -1.092622 | 0.001376 | 0.024398 | down |
| Slc29a1 | 2.728097 | 1.4478948 | 0.001378 | 0.024398 | up |
| Mirlet7c2 | 0.315912 | -1.662404 | 0.001402 | 0.024626 | down |
| Mirlet7b | 0.100838 | -3.309894 | 0.001404 | 0.024626 | down |
| Hmbox1 | 2.296027 | 1.1991399 | 0.001406 | 0.024654 | up |
| LOC102552952 | 896.7616 | 9.8085808 | 0.001422 | 0.024873 | up |
| Diaph2 | 2.281845 | 1.1902006 | 0.00143 | 0.024977 | up |
| Pvrig | 0.159858 | -2.645142 | 0.00144 | 0.025106 | down |
| ENSRNOG00000066146 | 0.354935 | -1.494373 | 0.001449 | 0.025196 | down |
| Ms4a14 | 0.140317 | -2.833243 | 0.001458 | 0.025334 | down |
| Wfikkn2 | 0.493799 | -1.018004 | 0.001467 | 0.025441 | down |
| Prx | 2.49808 | 1.3208195 | 0.001471 | 0.025475 | up |
| C3h9orf50 | 0.357193 | -1.485225 | 0.001472 | 0.025482 | down |
| Gltpd2 | 0.34033 | -1.554994 | 0.001482 | 0.025571 | down |
| Cd300e | 2.503852 | 1.3241492 | 0.001488 | 0.025617 | up |
| Ms4a12 | 0.229228 | -2.125142 | 0.001491 | 0.025645 | down |
| Pltp | 0.420484 | -1.249876 | 0.001492 | 0.025657 | down |
| Adamtsl5 | 0.435487 | -1.199299 | 0.001505 | 0.025722 | down |
| Matn4 | 2.182891 | 1.1262398 | 0.00152 | 0.025882 | up |
| Col6a4 | 3.20264 | 1.6792616 | 0.001522 | 0.025882 | up |
| Pzp | 0.132185 | -2.919367 | 0.001543 | 0.026087 | down |

| | | | | | |
|---|---|---|---|---|---|
| Zc3h12d | 0.40027 | -1.320954 | 0.001543 | 0.026087 | down |
| Dtx1 | 0.493936 | -1.017604 | 0.001553 | 0.026182 | down |
| Lifr | 2.610231 | 1.3841773 | 0.001576 | 0.026405 | up |
| Saal1 | 0.48575 | -1.041715 | 0.001578 | 0.02642 | down |
| Clcn2 | 0.49144 | -1.024912 | 0.001586 | 0.026517 | down |
| F11 | 2.051185 | 1.0364573 | 0.001587 | 0.026517 | up |
| Mpp7 | 2.045506 | 1.0324578 | 0.001597 | 0.026639 | up |
| Sgo2 | 2.004551 | 1.003279 | 0.001599 | 0.026658 | up |
| Tmc5 | 0.375895 | -1.411599 | 0.001622 | 0.026928 | down |
| LOC120099015 | 2.002808 | 1.0020242 | 0.001632 | 0.027022 | up |
| Adam28 | 0.494125 | -1.017053 | 0.001643 | 0.027127 | down |
| Cyp11a1 | 0.270772 | -1.884849 | 0.001651 | 0.027194 | down |
| Hist1h2bc | 0.205174 | -2.285077 | 0.001671 | 0.027374 | down |
| Pax2 | 0.001399 | -9.481007 | 0.00168 | 0.027458 | down |
| LOC100909660 | 0.057466 | -4.121154 | 0.001686 | 0.027526 | down |
| Adam8 | 0.451038 | -1.14868 | 0.001691 | 0.027558 | down |
| Col4a3 | 3.195594 | 1.6760843 | 0.001712 | 0.027744 | up |
| Alas2 | 4.164244 | 2.0580545 | 0.001725 | 0.027789 | up |
| Arhgap40 | 0.002841 | -8.459571 | 0.001725 | 0.027789 | down |
| Cyp3a23-3a1 | 0.063643 | -3.973852 | 0.00173 | 0.027849 | down |
| ENSRNOG00000066162 | 2.083521 | 1.059024 | 0.001733 | 0.027879 | up |
| Prkaa2 | 2.339428 | 1.226156 | 0.001771 | 0.028364 | up |
| Hgf | 2.208996 | 1.1433905 | 0.001786 | 0.02849 | up |
| Lingo1 | 2.106509 | 1.0748542 | 0.001786 | 0.02849 | up |
| Spock3 | 6.266072 | 2.6475614 | 0.00179 | 0.028532 | up |
| Phgdh | 0.476513 | -1.069412 | 0.001792 | 0.028532 | down |
| Rasal1 | 0.383776 | -1.381663 | 0.001799 | 0.028564 | down |
| Slc6a13 | 0.138238 | -2.854773 | 0.001799 | 0.028564 | down |
| Dsg1 | 0.337967 | -1.565045 | 0.001815 | 0.028742 | down |
| Nkapl | 0.002079 | -8.909713 | 0.00182 | 0.028761 | down |
| ENSRNOG00000063594 | 2.14131 | 1.0984937 | 0.00182 | 0.028761 | up |
| P2rx6 | 0.457771 | -1.127301 | 0.001823 | 0.028763 | down |
| Dach1 | 2.132934 | 1.0928392 | 0.001841 | 0.028959 | up |
| Fndc7 | 2.402915 | 1.2647853 | 0.001847 | 0.029021 | up |
| Tac4 | 0.289194 | -1.789891 | 0.001864 | 0.029069 | down |
| Crabp2 | 0.434795 | -1.201592 | 0.001865 | 0.029069 | down |
| Hecw1 | 0.005529 | -7.49874 | 0.001892 | 0.029324 | down |
| ENSRNOG00000063863 | 0.273569 | -1.870023 | 0.001894 | 0.029324 | down |
| Rasa4 | 0.473765 | -1.077757 | 0.001896 | 0.029332 | down |
| Nckap5 | 2.629569 | 1.3948261 | 0.001899 | 0.029332 | up |
| Lrrc37a | 0.46375 | -1.10858 | 0.001909 | 0.029399 | down |

| | | | | | |
|---|---|---|---|---|---|
| Slpi | 0.198357 | -2.333828 | 0.001911 | 0.029399 | down |
| Ccr5 | 0.495716 | -1.012414 | 0.00192 | 0.029399 | down |
| Itga2 | 3.01063 | 1.5900652 | 0.001925 | 0.029442 | up |
| Calcb | 0.151647 | -2.721213 | 0.001927 | 0.029452 | down |
| Iqcf1 | 0.047911 | -4.383513 | 0.001964 | 0.029734 | down |
| ENSRNOG00000068718 | 0.465394 | -1.103476 | 0.001974 | 0.029852 | down |
| Slco2a1 | 2.165848 | 1.1149322 | 0.002004 | 0.030165 | up |
| Atp5mc1 | 0.418843 | -1.255519 | 0.002009 | 0.030192 | down |
| Fam221a | 0.420989 | -1.248146 | 0.002013 | 0.030192 | down |
| ENSRNOG00000069461 | 0.397466 | -1.331098 | 0.002024 | 0.030228 | down |
| Aldob | 0.150917 | -2.728176 | 0.002035 | 0.030363 | down |
| Nmbr | 0.001269 | -9.622217 | 0.002043 | 0.030425 | down |
| Rasgrf2 | 2.048244 | 1.0343873 | 0.002047 | 0.030462 | up |
| Fras1 | 2.539535 | 1.3445643 | 0.002053 | 0.030506 | up |
| Tbx6 | 0.476656 | -1.068979 | 0.002072 | 0.030689 | down |
| Otop1 | 0.020504 | -5.607974 | 0.002099 | 0.031053 | down |
| Kif12 | 0.083155 | -3.588046 | 0.002105 | 0.031089 | down |
| ENSRNOG00000065225 | 0.26089 | -1.938485 | 0.002132 | 0.031389 | down |
| Gpr158 | 3.244206 | 1.6978654 | 0.002133 | 0.031389 | up |
| Aqp5 | 2.085972 | 1.06072 | 0.002153 | 0.031637 | up |
| Fcgr2b | 0.498124 | -1.005424 | 0.002174 | 0.031807 | down |
| Chrnd | 0.001564 | -9.320205 | 0.002188 | 0.031958 | down |
| Rspo2 | 0.295254 | -1.759973 | 0.002212 | 0.032192 | down |
| Pvalb | 0.292133 | -1.775301 | 0.002214 | 0.032192 | down |
| Atp8a1 | 2.168524 | 1.1167137 | 0.002247 | 0.03252 | up |
| Rgs7bp | 2.017531 | 1.0125908 | 0.002261 | 0.032673 | up |
| Vsx1 | 0.172946 | -2.531607 | 0.002265 | 0.032696 | down |
| Otog | 0.009292 | -6.749722 | 0.002274 | 0.032725 | down |
| Car3 | 0.068961 | -3.85807 | 0.002287 | 0.032817 | down |
| Lmbrd2 | 2.099507 | 1.0700508 | 0.002305 | 0.032997 | up |
| Pipox | 0.264596 | -1.918134 | 0.002352 | 0.033501 | down |
| ENSRNOG00000070902 | 3.966285 | 1.9877882 | 0.00236 | 0.033554 | up |
| Ppp4r4 | 0.428681 | -1.222023 | 0.002367 | 0.033563 | down |
| AC131360.2 | 2.798888 | 1.4848535 | 0.002376 | 0.033607 | up |
| Cpt1b | 0.258067 | -1.954184 | 0.002379 | 0.033607 | down |
| Fndc5 | 0.435014 | -1.200867 | 0.002384 | 0.033656 | down |
| Kcns1 | 0.361449 | -1.468134 | 0.002393 | 0.033728 | down |
| Lrrc73 | 0.458299 | -1.125639 | 0.00244 | 0.034238 | down |
| Olfm5 | 0.048927 | -4.353223 | 0.00245 | 0.034271 | down |
| Oasl | 0.45156 | -1.14701 | 0.002452 | 0.034271 | down |
| Xcl1 | 0.225893 | -2.146291 | 0.002475 | 0.034358 | down |

| | | | | | |
|---|---|---|---|---|---|
| Slc12a3 | 0.362492 | -1.463981 | 0.002517 | 0.034863 | down |
| Neto2 | 2.058024 | 1.04126 | 0.002532 | 0.034971 | up |
| Arl5c | 0.425153 | -1.233947 | 0.002537 | 0.03501 | down |
| Slc35f2 | 0.386348 | -1.372029 | 0.002553 | 0.035164 | down |
| ENSRNOG00000064985 | 0.421548 | -1.24623 | 0.002557 | 0.035164 | down |
| Yipf7 | 0.018993 | -5.718359 | 0.002559 | 0.035164 | down |
| Tmem255a | 2.154511 | 1.1073604 | 0.00256 | 0.035164 | up |
| Slc4a11 | 0.461884 | -1.114397 | 0.002585 | 0.03536 | down |
| Clec4g | 2.730546 | 1.4491892 | 0.002595 | 0.035457 | up |
| Tmco5b | 0.095242 | -3.392262 | 0.002607 | 0.035593 | down |
| Slc38a3 | 0.340981 | -1.552237 | 0.002614 | 0.035652 | down |
| Pwwp4 | 0.318083 | -1.652524 | 0.002618 | 0.035652 | down |
| Arrb1 | 2.027713 | 1.0198534 | 0.002631 | 0.035754 | up |
| B4galnt3 | 0.346789 | -1.527869 | 0.002635 | 0.035789 | down |
| Ccdc198 | 2.451695 | 1.2937793 | 0.002686 | 0.036251 | up |
| ENSRNOG00000067737 | 2.643682 | 1.4025486 | 0.00269 | 0.036277 | up |
| Plekhg4 | 0.310157 | -1.688928 | 0.002694 | 0.03629 | down |
| Slc22a1 | 0.464455 | -1.106391 | 0.002752 | 0.036704 | down |
| ENSRNOG00000070735 | 2.374848 | 1.2478351 | 0.002758 | 0.036704 | up |
| Aadac | 0.001919 | -9.025085 | 0.002817 | 0.037319 | down |
| Notum | 0.407558 | -1.294923 | 0.00283 | 0.037437 | down |
| RGD1308564 | 7.58512 | 2.9231721 | 0.002848 | 0.037565 | up |
| Ccdc33 | 0.434928 | -1.20115 | 0.002859 | 0.037615 | down |
| Cdh17 | 0.338112 | -1.564428 | 0.00286 | 0.037615 | down |
| Cxadrl1 | 0.47045 | -1.087887 | 0.002875 | 0.03778 | down |
| ENSRNOG00000069326 | 2.10176 | 1.0715982 | 0.002879 | 0.037787 | up |
| Zfp704 | 2.020615 | 1.0147942 | 0.002927 | 0.038192 | up |
| Car11 | 0.415413 | -1.267381 | 0.002943 | 0.038363 | down |
| Scnn1g | 2.212851 | 1.145906 | 0.002979 | 0.038723 | up |
| Adamts9 | 2.008076 | 1.005814 | 0.002986 | 0.038772 | up |
| Actl10 | 0.412423 | -1.277804 | 0.002988 | 0.038772 | down |
| ENSRNOG00000064997 | 0.169261 | -2.56268 | 0.003011 | 0.038941 | down |
| ENSRNOG00000066627 | 444.956 | 8.7975189 | 0.003013 | 0.038947 | up |
| Togaram2 | 0.456474 | -1.131394 | 0.003036 | 0.03911 | down |
| Slc14a1 | 0.456538 | -1.131194 | 0.003045 | 0.039183 | down |
| ENSRNOG00000063039 | 6.690746 | 2.7421671 | 0.003048 | 0.039205 | up |
| Rtl9 | 0.21552 | -2.214105 | 0.003053 | 0.039221 | down |
| Bean1 | 2.121823 | 1.0853046 | 0.003055 | 0.039221 | up |
| ENSRNOG00000066970 | 0.319962 | -1.644029 | 0.003067 | 0.039275 | down |
| ENSRNOG00000063639 | 0.490501 | -1.027672 | 0.003099 | 0.039572 | down |
| Pde8b | 2.02833 | 1.0202922 | 0.003104 | 0.039589 | up |

| | | | | | |
|---|---|---|---|---|---|
| Tafa5 | 0.39389 | -1.344135 | 0.003116 | 0.039696 | down |
| Ppef1 | 2.24846 | 1.1689373 | 0.003128 | 0.039812 | up |
| Rnf11l2 | 2.241442 | 1.1644274 | 0.003134 | 0.039869 | up |
| Rnf24 | 2.940481 | 1.5560523 | 0.003141 | 0.039906 | up |
| Stra6l | 0.450148 | -1.151529 | 0.003196 | 0.040251 | down |
| Serpinc1 | 0.343739 | -1.540616 | 0.003231 | 0.040584 | down |
| Slc28a2 | 0.478455 | -1.063545 | 0.003235 | 0.040584 | down |
| Plin1 | 0.0068 | -7.200179 | 0.003236 | 0.040584 | down |
| ENSRNOG00000063348 | 0.373477 | -1.420907 | 0.003242 | 0.04061 | down |
| Fabp3 | 0.26607 | -1.91012 | 0.003253 | 0.040712 | down |
| Rdh7 | 0.147782 | -2.758461 | 0.003258 | 0.040743 | down |
| Wfdc21 | 0.240544 | -2.055628 | 0.003306 | 0.041202 | down |
| Lrrc39 | 0.422272 | -1.243755 | 0.003322 | 0.041349 | down |
| LOC120093077 | 0.146139 | -2.77459 | 0.003339 | 0.041509 | down |
| Or10aa3 | 0.270478 | -1.886419 | 0.003363 | 0.041722 | down |
| Mcpt1l1 | 0.358076 | -1.481661 | 0.003372 | 0.041789 | down |
| Klf7 | 2.1578 | 1.1095608 | 0.003385 | 0.041904 | up |
| ENSRNOG00000070740 | 0.437998 | -1.191003 | 0.003396 | 0.042004 | down |
| Sptssb | 0.078245 | -3.675851 | 0.003482 | 0.04268 | down |
| Syt5 | 0.428944 | -1.221139 | 0.003534 | 0.042948 | down |
| Hs3st3b1 | 2.134132 | 1.0936495 | 0.003537 | 0.042948 | up |
| Mir21 | 0.342366 | -1.546389 | 0.003601 | 0.043589 | down |
| Faiml | 0.352333 | -1.504988 | 0.003611 | 0.043656 | down |
| Nwd1 | 0.38198 | -1.388431 | 0.003624 | 0.04373 | down |
| Camkv | 0.160694 | -2.637616 | 0.003624 | 0.04373 | down |
| Aqp8 | 0.24182 | -2.047995 | 0.003635 | 0.043796 | down |
| Slc30a3 | 0.464631 | -1.105843 | 0.003644 | 0.043815 | down |
| Lcn5 | 0.001001 | -9.96435 | 0.00365 | 0.043834 | down |
| Atp1a2 | 0.177818 | -2.49153 | 0.003661 | 0.043923 | down |
| Fam111a | 6.889556 | 2.7844111 | 0.003747 | 0.044701 | up |
| Dio3 | 0.48522 | -1.043288 | 0.003752 | 0.044739 | down |
| Shbg | 0.494841 | -1.014963 | 0.003774 | 0.044928 | down |
| AABR07047835.1 | 2.186475 | 1.1286072 | 0.003775 | 0.044928 | up |
| Ly49s7 | 0.283414 | -1.819019 | 0.003788 | 0.045046 | down |
| Orm1 | 0.410584 | -1.28425 | 0.003864 | 0.045718 | down |
| Lrfn5 | 326.136 | 8.3493299 | 0.00389 | 0.045929 | up |
| Pou2af2 | 2.154056 | 1.1070557 | 0.003916 | 0.046147 | up |
| Smoc1 | 0.257972 | -1.954712 | 0.003963 | 0.046483 | down |
| Nccrp1 | 0.121177 | -3.044817 | 0.003968 | 0.046498 | down |
| Ager | 2.394006 | 1.2594266 | 0.004006 | 0.046877 | up |
| Cd8a | 0.496133 | -1.011202 | 0.004011 | 0.046912 | down |

| | | | | | |
|---|---|---|---|---|---|
| Reps2 | 2.630568 | 1.3953741 | 0.004035 | 0.047076 | up |
| Tph1 | 0.43219 | -1.210262 | 0.004077 | 0.047421 | down |
| Psd | 0.417384 | -1.260552 | 0.004111 | 0.047718 | down |
| Pag1 | 2.332966 | 1.2221654 | 0.004148 | 0.047957 | up |
| C1s | 0.480934 | -1.056089 | 0.00421 | 0.048479 | down |
| ENSRNOG00000066981 | 0.104292 | -3.261296 | 0.004231 | 0.04863 | down |
| LOC120099582 | 0.000453 | -11.10948 | 0.004256 | 0.048836 | down |
| Fga | 0.141785 | -2.818219 | 0.004302 | 0.049315 | down |
| Hhip | 2.645198 | 1.403376 | 0.004325 | 0.049505 | up |

**Table 4. Female SHAM vs Male SHAM mRNA-seq**

Significantly Differentially Expressed Genes ($Log_2FC > |1|$; qvalue < 0.05)

| Gene | fc | log2(fc) | pval | qval | regulation |
|---|---|---|---|---|---|
| Kdm5d | 1.49E-05 | -16.03565 | 1.54E-54 | 3.42E-50 | down |
| Eif2s3y | 7.42E-06 | -17.04014 | 8.97E-54 | 9.98E-50 | down |
| Ddx3 | 8.68E-06 | -16.8131 | 1.79E-51 | 1.33E-47 | down |
| Uty | 4.54E-05 | -14.42834 | 6.5E-43 | 3.61E-39 | down |
| Col1a1 | 0.245467 | -2.026398 | 1.73E-28 | 6.41E-25 | down |
| Mmp2 | 0.479857 | -1.059324 | 9.04E-28 | 2.87E-24 | down |
| Mrps17 | 0.166521 | -2.586221 | 2.49E-22 | 6.91E-19 | down |
| C1qtnf6 | 0.334601 | -1.579486 | 1.91E-21 | 4.72E-18 | down |
| Uba1y | 0.000299 | -11.70758 | 1.78E-19 | 3.6E-16 | down |
| Myl1 | 3.86E-05 | -14.65974 | 8.75E-18 | 1.5E-14 | down |
| Fn1 | 0.454621 | -1.137264 | 1.2E-17 | 1.91E-14 | down |
| Fbn1 | 0.39817 | -1.328544 | 8.09E-17 | 1.2E-13 | down |
| Nid1 | 0.477083 | -1.067688 | 4.83E-16 | 6.71E-13 | down |
| LOC103694557 | 0.000467 | -11.06339 | 1.46E-15 | 1.82E-12 | down |
| Itga2 | 0.312271 | -1.679128 | 1.47E-15 | 1.82E-12 | down |
| Aldh1l2 | 0.137295 | -2.864654 | 4.65E-15 | 5.45E-12 | down |
| Eln | 0.296728 | -1.752789 | 5.27E-15 | 5.86E-12 | down |
| Mup4 | 0.000289 | -11.75695 | 6.24E-15 | 6.61E-12 | down |
| Mylk4 | 0.000518 | -10.91606 | 1.24E-14 | 1.26E-11 | down |
| ENSRNOG00000064849 | 3444.042 | 11.749887 | 1.32E-14 | 1.27E-11 | up |
| Loxl2 | 0.42633 | -1.229957 | 1.55E-14 | 1.43E-11 | down |
| Col1a2 | 0.39687 | -1.33326 | 2.58E-14 | 2.29E-11 | down |
| Usp9y | 0.002706 | -8.529886 | 1.69E-13 | 1.3E-10 | down |
| Col3a1 | 0.352088 | -1.50599 | 1.94E-13 | 1.44E-10 | down |

| | | | | | |
|---|---|---|---|---|---|
| Col5a1 | 0.487092 | -1.037734 | 2.14E-13 | 1.54E-10 | down |
| LOC298116 | 0.000852 | -10.19622 | 3.68E-13 | 2.56E-10 | down |
| Esm1 | 3.4601 | 1.7908137 | 3.98E-13 | 2.68E-10 | up |
| ENSRNOG00000066693 | 3885.094 | 11.923734 | 5.76E-13 | 3.77E-10 | up |
| Ppp1r27 | 0.000308 | -11.6651 | 6.67E-13 | 4.24E-10 | down |
| ENSRNOG00000062326 | 8.52E-05 | -13.51822 | 1.08E-12 | 6.64E-10 | down |
| Col6a2 | 0.490103 | -1.028844 | 1.32E-12 | 7.71E-10 | down |
| Tmem255b | 2.024639 | 1.0176646 | 3.25E-12 | 1.76E-09 | up |
| Opcml | 0.365756 | -1.451048 | 3.91E-12 | 2.01E-09 | down |
| Hao2 | 0.000194 | -12.32845 | 3.99E-12 | 2.01E-09 | down |
| ENSRNOG00000069825 | 864.212 | 9.7552415 | 5.77E-12 | 2.85E-09 | up |
| Pcbd1 | 2846.682 | 11.475066 | 6.86E-12 | 3.25E-09 | up |
| Ltbp2 | 0.489395 | -1.03093 | 1.27E-11 | 5.64E-09 | down |
| Serpinf1 | 0.404291 | -1.306534 | 2.49E-11 | 1.03E-08 | down |
| Nell1 | 0.408306 | -1.292277 | 1.31E-10 | 4.7E-08 | down |
| Cyp4a2 | 0.00068 | -10.52246 | 1.98E-10 | 6.97E-08 | down |
| Ccdc80 | 0.48533 | -1.042963 | 2.15E-10 | 7.46E-08 | down |
| Vcan | 0.421436 | -1.246614 | 2.78E-10 | 9E-08 | down |
| Cyp2a2 | 0.00021 | -12.22008 | 2.79E-10 | 9E-08 | down |
| Stra6 | 0.276084 | -1.85682 | 3.02E-10 | 9.6E-08 | down |
| Sult1c3 | 0.1249 | -3.00116 | 3.13E-10 | 9.73E-08 | down |
| Sgsm1 | 2.957853 | 1.5645505 | 3.18E-10 | 9.73E-08 | up |
| Cyp26b1 | 0.358351 | -1.480554 | 3.19E-10 | 9.73E-08 | down |
| Fndc1 | 0.30147 | -1.729915 | 3.83E-10 | 1.14E-07 | down |
| Sult1e1 | 0.005535 | -7.497196 | 4.24E-10 | 1.24E-07 | down |
| Fscn1 | 0.464425 | -1.106481 | 6.17E-10 | 1.76E-07 | down |
| Cyp2c13 | 6.85E-05 | -13.83384 | 1.05E-09 | 2.95E-07 | down |
| Adamts12 | 0.385648 | -1.374644 | 1.17E-09 | 3.25E-07 | down |
| LOC100360095 | 4.28E-05 | -14.51309 | 2.18E-09 | 5.78E-07 | down |
| Tnc | 0.352128 | -1.50583 | 2.92E-09 | 7.55E-07 | down |
| Dbn1 | 0.480304 | -1.057981 | 2.95E-09 | 7.55E-07 | down |
| LOC681325 | 2.320643 | 1.2145248 | 3.05E-09 | 7.71E-07 | up |
| Tmem59l | 0.490966 | -1.026304 | 3.22E-09 | 8.05E-07 | down |
| Cyp3a23-3a1 | 0.018291 | -5.77273 | 4.86E-09 | 1.2E-06 | down |
| H19 | 0.013254 | -6.237398 | 7.11E-09 | 1.73E-06 | down |
| LOC102548186 | 2.805481 | 1.4882481 | 1.17E-08 | 2.71E-06 | up |
| Col15a1 | 0.378721 | -1.400792 | 1.5E-08 | 3.44E-06 | down |
| Cpz | 0.344491 | -1.537461 | 1.56E-08 | 3.55E-06 | down |
| Adamts17 | 0.416699 | -1.262923 | 1.76E-08 | 3.91E-06 | down |
| Nid2 | 0.495027 | -1.014422 | 2.77E-08 | 5.87E-06 | down |
| Cyp3a2 | 0.002227 | -8.810889 | 4E-08 | 7.95E-06 | down |

| | | | | | |
|---|---|---|---|---|---|
| Ccl9 | 2.040077 | 1.0286235 | 4.85E-08 | 9.47E-06 | up |
| Chl1 | 0.358149 | -1.481367 | 9.57E-08 | 1.72E-05 | down |
| Nrep | 0.464901 | -1.105006 | 1.11E-07 | 1.95E-05 | down |
| ENSRNOG00000069681 | 2.775721 | 1.4728623 | 1.33E-07 | 2.26E-05 | up |
| RGD1564696 | 6.501112 | 2.7006865 | 1.34E-07 | 2.26E-05 | up |
| Bmper | 0.496144 | -1.011169 | 1.51E-07 | 2.49E-05 | down |
| Impg1 | 0.001328 | -9.556549 | 1.67E-07 | 2.63E-05 | down |
| Mall | 0.384889 | -1.377487 | 2.53E-07 | 3.67E-05 | down |
| Dhrs7l1 | 0.050914 | -4.295791 | 4.52E-07 | 6.02E-05 | down |
| Cdh6 | 0.162423 | -2.622175 | 4.72E-07 | 6.21E-05 | down |
| Galnt14 | 4.499583 | 2.1697913 | 4.91E-07 | 6.38E-05 | up |
| Alas2 | 0.418459 | -1.256843 | 5.17E-07 | 6.6E-05 | down |
| Ncam1 | 0.493199 | -1.019759 | 8.67E-07 | 0.000101 | down |
| Etfb | 0.488484 | -1.033617 | 1.04E-06 | 0.00012 | down |
| Pax8 | 2.780849 | 1.4755253 | 1.14E-06 | 0.000129 | up |
| Hmcn1 | 0.498219 | -1.005148 | 1.28E-06 | 0.000142 | down |
| LOC685680 | 2.016861 | 1.0121117 | 2E-06 | 0.000213 | up |
| Serpina1 | 0.421351 | -1.246906 | 2.1E-06 | 0.000222 | down |
| Scn3b | 0.493037 | -1.020233 | 2.42E-06 | 0.000248 | down |
| LOC100912991 | 2.198456 | 1.1364908 | 2.51E-06 | 0.000254 | up |
| Pkhd1 | 0.430134 | -1.217142 | 3.03E-06 | 0.000299 | down |
| Cyp2c11 | 0.024015 | -5.379932 | 3.23E-06 | 0.000317 | down |
| Prox1 | 0.319433 | -1.646413 | 3.57E-06 | 0.000337 | down |
| Orm1 | 0.169743 | -2.558574 | 4.21E-06 | 0.00039 | down |
| Adgrl2 | 0.170457 | -2.552522 | 4.63E-06 | 0.000418 | down |
| Sbspon | 0.46187 | -1.114441 | 4.99E-06 | 0.000442 | down |
| Uchl3 | 0.490027 | -1.029068 | 5.14E-06 | 0.000452 | down |
| Capn3 | 0.201516 | -2.311033 | 5.18E-06 | 0.000453 | down |
| Inmt | 2.339902 | 1.226448 | 5.8E-06 | 0.000496 | up |
| Clec4g | 0.390237 | -1.357578 | 6.25E-06 | 0.000528 | down |
| Col5a3 | 0.452354 | -1.144476 | 7.11E-06 | 0.000592 | down |
| Muc5ac | 0.209613 | -2.254202 | 7.96E-06 | 0.000651 | down |
| Snca | 0.27956 | -1.838771 | 8.04E-06 | 0.000653 | down |
| Acta1 | 0.019318 | -5.693922 | 9.63E-06 | 0.000765 | down |
| Bpifb1 | 0.309 | -1.694321 | 1.15E-05 | 0.000899 | down |
| Capn6 | 0.314658 | -1.668141 | 1.16E-05 | 0.000902 | down |
| Apon | 0.015286 | -6.031682 | 1.35E-05 | 0.000995 | down |
| Gpbp1l2 | 2.391368 | 1.2578362 | 1.49E-05 | 0.001087 | up |
| Hsd3b5 | 0.005274 | -7.567017 | 1.5E-05 | 0.001088 | down |
| Retnlg | 0.348638 | -1.520199 | 1.66E-05 | 0.001192 | down |
| Soga3 | 2.750545 | 1.4597175 | 1.89E-05 | 0.001324 | up |

| | | | | | |
|---|---|---|---|---|---|
| Obp3 | 0.010752 | -6.539251 | 1.92E-05 | 0.00134 | down |
| Mettl7b | 0.057532 | -4.119492 | 1.93E-05 | 0.001341 | down |
| Kcnj12 | 0.483282 | -1.049061 | 2.06E-05 | 0.001417 | down |
| Igsf10 | 0.477107 | -1.067617 | 2.31E-05 | 0.001549 | down |
| Slc28a2 | 0.45844 | -1.125196 | 2.41E-05 | 0.001607 | down |
| Prune2 | 0.341834 | -1.548634 | 2.58E-05 | 0.001684 | down |
| Plcb1 | 0.47708 | -1.067698 | 2.84E-05 | 0.001815 | down |
| Ntrk2 | 0.418846 | -1.25551 | 3.03E-05 | 0.001904 | down |
| Serpine1 | 0.47666 | -1.068968 | 3.04E-05 | 0.001906 | down |
| Ern2 | 0.288008 | -1.795821 | 3.06E-05 | 0.001914 | down |
| Tnnc2 | 0.017046 | -5.874405 | 3.35E-05 | 0.002063 | down |
| Ccl4 | 2.356585 | 1.2366979 | 3.37E-05 | 0.002068 | up |
| Ciart | 0.243449 | -2.038308 | 3.38E-05 | 0.002068 | down |
| Ccl11 | 0.499924 | -1.000219 | 3.68E-05 | 0.002238 | down |
| Cxcl1 | 2.209998 | 1.1440448 | 3.72E-05 | 0.002246 | up |
| Upk1a | 2.51811 | 1.3323412 | 3.86E-05 | 0.00229 | up |
| LOC120099582 | 0.000531 | -10.87845 | 3.88E-05 | 0.00229 | down |
| Tm4sf19 | 3.114848 | 1.6391618 | 3.89E-05 | 0.00229 | up |
| hist1h2ail2 | 3.968754 | 1.9886863 | 4.06E-05 | 0.00236 | up |
| Ccr3 | 0.319614 | -1.645598 | 4.59E-05 | 0.00257 | down |
| Islr2 | 0.40857 | -1.291345 | 4.9E-05 | 0.002699 | down |
| Pcdhb6l | 0.483711 | -1.047784 | 5.45E-05 | 0.002928 | down |
| Tk1 | 0.409659 | -1.287505 | 5.47E-05 | 0.002929 | down |
| Pycr1 | 0.489831 | -1.029645 | 5.88E-05 | 0.003111 | down |
| Adamts14 | 0.482937 | -1.050093 | 6.45E-05 | 0.003324 | down |
| Lingo1 | 0.467317 | -1.097527 | 6.48E-05 | 0.003324 | down |
| Mfap2 | 0.420857 | -1.248597 | 6.49E-05 | 0.003324 | down |
| ENSRNOG00000037911 | 43.53566 | 5.4441255 | 7.1E-05 | 0.003565 | up |
| Dbp | 0.463294 | -1.110001 | 7.77E-05 | 0.003795 | down |
| Dcstamp | 2.190867 | 1.1315018 | 7.78E-05 | 0.003795 | up |
| Mtus2 | 2.028142 | 1.0201584 | 8.15E-05 | 0.003965 | up |
| Samd14 | 0.491244 | -1.025488 | 8.29E-05 | 0.004008 | down |
| ENSRNOG00000062487 | 0.000766 | -10.35114 | 8.64E-05 | 0.004133 | down |
| Slc5a5 | 0.038318 | -4.705825 | 8.8E-05 | 0.004196 | down |
| Urad | 0.000526 | -10.89245 | 9.23E-05 | 0.00434 | down |
| Cbln2 | 0.356965 | -1.486145 | 0.000104 | 0.004746 | down |
| Ebf1 | 0.484919 | -1.044186 | 0.000112 | 0.005042 | down |
| Cpamd8 | 0.091836 | -3.4448 | 0.000114 | 0.005109 | down |
| ENSRNOG00000068078 | 0.303623 | -1.719646 | 0.000131 | 0.005693 | down |
| Cyp3a18 | 0.061512 | -4.022977 | 0.000132 | 0.005706 | down |
| ENSRNOG00000065596 | 4.42E-06 | -17.78837 | 0.000141 | 0.006002 | down |

| | | | | | |
|---|---|---|---|---|---|
| F13b | 0.01153 | -6.438413 | 0.000145 | 0.006098 | down |
| Brinp1 | 0.281571 | -1.828432 | 0.000147 | 0.006175 | down |
| Foxc2 | 0.335088 | -1.577389 | 0.000156 | 0.006537 | down |
| Nkx1-2 | 2021.026 | 10.980872 | 0.000158 | 0.006585 | up |
| ENSRNOG00000071081 | 24819.55 | 14.599189 | 0.000159 | 0.006619 | up |
| Cxcl6 | 3.342925 | 1.7411109 | 0.000162 | 0.006678 | up |
| Sphkap | 0.493052 | -1.020187 | 0.000165 | 0.006767 | down |
| Stac3 | 0.013719 | -6.187716 | 0.000183 | 0.007335 | down |
| Chaf1b | 0.410515 | -1.284494 | 0.000187 | 0.007407 | down |
| Pdzrn4 | 0.378485 | -1.401692 | 0.000188 | 0.007434 | down |
| Lyc2 | 0.307881 | -1.699557 | 0.000201 | 0.007843 | down |
| Dnajc6 | 2.88053 | 1.5263345 | 0.000204 | 0.007895 | up |
| Acsbg1 | 0.410125 | -1.285863 | 0.000207 | 0.00801 | down |
| Ccno | 0.475729 | -1.071788 | 0.00021 | 0.008085 | down |
| Timd2 | 0.319762 | -1.644928 | 0.000212 | 0.008097 | down |
| Apoa5 | 0.153273 | -2.705824 | 0.000236 | 0.008754 | down |
| Pcdhb5 | 0.473657 | -1.078086 | 0.000236 | 0.008754 | down |
| Pcdhb11 | 0.431461 | -1.212697 | 0.000248 | 0.009069 | down |
| Myrfl | 0.232215 | -2.106465 | 0.000259 | 0.009401 | down |
| Cemip | 0.497115 | -1.008347 | 0.000259 | 0.009401 | down |
| Cgref1 | 0.429695 | -1.218615 | 0.0003 | 0.010446 | down |
| ENSRNOG00000062522 | 0.415984 | -1.265402 | 0.000304 | 0.010555 | down |
| Slc4a1 | 0.376117 | -1.410747 | 0.000313 | 0.010767 | down |
| ENSRNOG00000065040 | 9.047689 | 3.1775494 | 0.000324 | 0.011046 | up |
| Ttr | 0.061781 | -4.016686 | 0.00033 | 0.01123 | down |
| Reg3g | 0.128815 | -2.956632 | 0.000335 | 0.011348 | down |
| Serpina3n | 0.037698 | -4.729349 | 0.000336 | 0.011363 | down |
| RT1-CE15 | 2.075207 | 1.053255 | 0.000338 | 0.011376 | up |
| Dpy30 | 0.092314 | -3.437313 | 0.000338 | 0.011378 | down |
| ENSRNOG00000068563 | 52.9007 | 5.7252148 | 0.000398 | 0.0128 | up |
| Nacad | 0.497253 | -1.007949 | 0.000422 | 0.01334 | down |
| Kyat1 | 0.151158 | -2.725867 | 0.000425 | 0.013426 | down |
| Cyp3a23-3a1 | 0.000159 | -12.62043 | 0.00044 | 0.013715 | down |
| Capn8 | 2.783424 | 1.4768605 | 0.000456 | 0.014103 | up |
| Il1rl2 | 2.165031 | 1.1143874 | 0.000457 | 0.014103 | up |
| Serpina3a | 0.121778 | -3.037679 | 0.000475 | 0.014514 | down |
| F2 | 0.323288 | -1.629109 | 0.000534 | 0.015896 | down |
| Frmpd3 | 0.208124 | -2.264483 | 0.000535 | 0.015896 | down |
| Cyp2d3 | 0.411718 | -1.280271 | 0.000536 | 0.015896 | down |
| Ahsg | 0.094905 | -3.397369 | 0.000556 | 0.01641 | down |
| Igfals | 0.085587 | -3.546466 | 0.000572 | 0.016792 | down |

| | | | | | |
|---|---|---|---|---|---|
| Hist1h4m | 2.690755 | 1.4280112 | 0.000584 | 0.017076 | up |
| Fam163a | 0.466935 | -1.098706 | 0.000587 | 0.017121 | down |
| Micb | 0.487426 | -1.036746 | 0.000606 | 0.017398 | down |
| ENSRNOG00000069434 | 0.029437 | -5.086224 | 0.000649 | 0.018395 | down |
| Angpt4 | 0.42638 | -1.229789 | 0.000696 | 0.019311 | down |
| Ucn3 | 0.044821 | -4.479673 | 0.00071 | 0.019607 | down |
| Cpne9 | 3.589341 | 1.8437189 | 0.000725 | 0.019866 | up |
| Foxn4 | 0.398594 | -1.32701 | 0.000734 | 0.020023 | down |
| Apoc1 | 0.177652 | -2.492872 | 0.000757 | 0.020493 | down |
| Atp2a1 | 0.049005 | -4.350922 | 0.000769 | 0.020762 | down |
| Chrna7 | 0.476456 | -1.069584 | 0.00081 | 0.021507 | down |
| ENSRNOG00000067172 | 0.467905 | -1.095713 | 0.000839 | 0.022116 | down |
| Sfrp2 | 0.471372 | -1.085063 | 0.000844 | 0.022245 | down |
| Dnase2b | 2.925721 | 1.548792 | 0.000847 | 0.022274 | up |
| Cep72 | 0.469175 | -1.091803 | 0.000888 | 0.02294 | down |
| Bnip5 | 0.296433 | -1.754221 | 0.000901 | 0.023182 | down |
| Fer1l6 | 0.421843 | -1.245222 | 0.000919 | 0.023518 | down |
| Ces1f | 0.411238 | -1.281954 | 0.000933 | 0.023702 | down |
| Apoa4 | 0.059704 | -4.066038 | 0.000949 | 0.023983 | down |
| Ihh | 2.585949 | 1.3706939 | 0.000978 | 0.024447 | up |
| St6galnac2 | 2.355328 | 1.2359278 | 0.001009 | 0.025037 | up |
| Brinp2 | 0.28228 | -1.824801 | 0.001029 | 0.02529 | down |
| Itln1 | 0.089394 | -3.48367 | 0.001137 | 0.02741 | down |
| Slc26a4 | 2.422315 | 1.2763867 | 0.001153 | 0.02766 | up |
| ENSRNOG00000062708 | 0.053175 | -4.233113 | 0.001158 | 0.02771 | down |
| Slc6a7 | 3.029968 | 1.5993025 | 0.001315 | 0.030547 | up |
| Akr1c12 | 0.198045 | -2.336103 | 0.00134 | 0.030998 | down |
| Clca1 | 0.134296 | -2.896515 | 0.001379 | 0.031674 | down |
| Pak5 | 0.006409 | -7.285735 | 0.001389 | 0.031713 | down |
| Cpn2 | 0.047773 | -4.387671 | 0.00141 | 0.031946 | down |
| Chad | 0.45127 | -1.147937 | 0.001411 | 0.031946 | down |
| ENSRNOG00000063050 | 3.474402 | 1.7967646 | 0.001421 | 0.032097 | up |
| Hpx | 0.176366 | -2.503354 | 0.001427 | 0.032133 | down |
| LOC120102953 | 0.000841 | -10.21639 | 0.001429 | 0.032133 | down |
| Mcpt1 | 0.231042 | -2.113771 | 0.001436 | 0.03222 | down |
| Dpp6 | 0.420485 | -1.249872 | 0.001436 | 0.03222 | down |
| Mcpt9 | 0.444733 | -1.16899 | 0.001466 | 0.032697 | down |
| Pnma8c | 0.123548 | -3.016858 | 0.00148 | 0.032911 | down |
| Arfgef3 | 0.492166 | -1.022784 | 0.001504 | 0.033233 | down |
| Luzp2 | 0.462132 | -1.113622 | 0.00157 | 0.034183 | down |
| Siglech | 0.36697 | -1.446266 | 0.001583 | 0.034301 | down |

| | | | | | |
|---|---|---|---|---|---|
| Lsmem1 | 0.001235 | -9.661755 | 0.001675 | 0.035913 | down |
| Slc12a3 | 0.438141 | -1.190532 | 0.001694 | 0.036196 | down |
| Tnfrsf10b | 0.49388 | -1.017768 | 0.001708 | 0.036332 | down |
| Nr1d1 | 0.35896 | -1.478106 | 0.001724 | 0.036513 | down |
| Apoc3 | 0.10076 | -3.311004 | 0.001745 | 0.036843 | down |
| Pgc | 0.357045 | -1.485822 | 0.001751 | 0.036873 | down |
| Ambp | 0.093083 | -3.425333 | 0.00176 | 0.036959 | down |
| Cpne4 | 0.249131 | -2.005025 | 0.001818 | 0.037777 | down |
| Tmem86b | 0.355062 | -1.493856 | 0.001837 | 0.038095 | down |
| Syt6 | 0.484546 | -1.045294 | 0.001839 | 0.038095 | down |
| LOC120103526 | 2.823569 | 1.4975199 | 0.001891 | 0.038744 | up |
| Glyat | 0.030088 | -5.054691 | 0.001942 | 0.039574 | down |
| Tmem169 | 2.246105 | 1.1674255 | 0.002018 | 0.040731 | up |
| Angptl7 | 0.295903 | -1.756802 | 0.002038 | 0.040931 | down |
| Ptpn5 | 303.798 | 8.2469686 | 0.002144 | 0.042478 | up |
| Itih1 | 0.209394 | -2.255709 | 0.00217 | 0.042922 | down |
| LOC100362949 | 1539.574 | 10.588315 | 0.002229 | 0.043979 | up |
| Scart1 | 2.235353 | 1.1605029 | 0.002259 | 0.044319 | up |
| LOC102555814 | 0.213038 | -2.230816 | 0.002266 | 0.044407 | down |
| Exo1 | 0.453359 | -1.141276 | 0.002274 | 0.044505 | down |
| Hnf4a | 0.133363 | -2.906567 | 0.002319 | 0.045021 | down |
| Tat | 0.317392 | -1.655663 | 0.002353 | 0.045326 | down |
| Sstr1 | 2.053477 | 1.0380688 | 0.002383 | 0.045756 | up |
| Fcrl2 | 0.162228 | -2.623904 | 0.002385 | 0.045759 | down |
| ENSRNOG00000062545 | 2.431526 | 1.2818618 | 0.002388 | 0.045769 | up |
| Rtkn2 | 0.000333 | -11.55136 | 0.002519 | 0.047334 | down |
| Armh4 | 0.480818 | -1.056437 | 0.002529 | 0.047449 | down |
| Cyp2b3 | 0.051906 | -4.267967 | 0.002552 | 0.047763 | down |
| Cntn6 | 0.067245 | -3.894419 | 0.002633 | 0.048759 | down |
| Penk | 0.387469 | -1.367847 | 0.002634 | 0.048759 | down |
| Gja3 | 0.235708 | -2.084925 | 0.002675 | 0.049155 | down |
| Pglyrp2 | 0.373504 | -1.420803 | 0.002703 | 0.04962 | down |
| Stoml3 | 0.399401 | -1.324089 | 0.002724 | 0.049924 | down |

**Table 5. Female BLEO vs Female SHAM microRNA-seq**
Significantly Differentially Expressed microRNAs ($log_2$FC |1|, pvalue < 0.05)

| miR name | fc | $log_2$(fc) | pvalue | regulation |
|---|---|---|---|---|
| rno-miR-449a-3p | 27.4086515 | 4.77655945 | 0.00183352 | up |
| rno-miR-212-3p | 11.9792569 | 3.58246651 | 0.02310434 | up |
| rno-miR-449c-5p_R+4 | 11.7391755 | 3.55325918 | 0.00811993 | up |
| mmu-mir-6240-p3_1ss3TG_1 | 11.2987704 | 3.49809387 | 0.00777637 | up |
| mmu-mir-449b-p3 | 11.2162423 | 3.48751751 | 0.00372985 | up |
| mmu-mir-449b-p5 | 11.2162423 | 3.48751751 | 0.00372985 | up |
| rno-miR-296-3p | 7.54943193 | 2.91636809 | 0.01113314 | up |
| rno-miR-212-5p_R-1 | 6.97831343 | 2.8028784 | 0.03884598 | up |
| mmu-miR-449b_R+3_2ss10-G11-T | 6.93355781 | 2.79359583 | 0.01661055 | up |
| rno-miR-132-5p | 5.66037436 | 2.50089747 | 0.03544841 | up |
| rno-miR-184 | 4.85197091 | 2.2785709 | 0.02027334 | up |
| PC-5p-3418_1419 | 4.84123042 | 2.27537376 | 0.00713778 | up |
| rno-miR-21-3p | 4.72678593 | 2.24085953 | 0.00767766 | up |
| pal-miR-9226-5p_1ss4AG | 4.64385968 | 2.21532438 | 0.00949099 | up |
| mmu-miR-1983 | 4.62170651 | 2.20842565 | 0.04727514 | up |
| hsa-miR-4454_L-2 | 4.39670474 | 2.13642265 | 0.0161356 | up |
| ssc-mir-1285-p3 | 4.24320932 | 2.08515585 | 0.04485264 | up |
| rno-miR-449a-5p | 4.08731977 | 2.03115512 | 0.00972262 | up |
| mmu-miR-26a-2-3p_1ss4GA | 4.03188617 | 2.01145491 | 0.00158665 | up |
| rno-miR-132-3p | 4.00810601 | 2.00292067 | 0.00579044 | up |
| rno-miR-23b-5p | 3.68379305 | 1.88119202 | 0.00143789 | up |
| rno-miR-21-5p_R+1 | 3.65419759 | 1.86955465 | 0.01280314 | up |
| rno-miR-27a-5p | 3.45157923 | 1.7872566 | 0.00848774 | up |
| rno-mir-497-p3 | 3.30018486 | 1.72254684 | 0.00537028 | up |
| rno-miR-511-3p_R+1 | 3.21061599 | 1.68285012 | 0.00019951 | up |
| pal-miR-9993b-3p_1ss15TA | 3.13935487 | 1.65046812 | 0.01801746 | up |
| ppy-mir-27a-p5 | 3.02759484 | 1.59817215 | 0.04414813 | up |
| mmu-miR-335-3p | 2.97752608 | 1.57411415 | 0.01358728 | up |
| rno-miR-16-3p_R+1 | 2.971209 | 1.57105009 | 0.02990691 | up |
| PC-5p-8089_451 | 2.95293126 | 1.56214777 | 0.02022382 | up |
| hsa-miR-4286_R+1 | 2.94427056 | 1.55791025 | 0.02978848 | up |
| rno-let-7c-2-3p | 2.5808382 | 1.3678397 | 0.00063072 | up |
| rno-miR-130b-5p | 2.56234863 | 1.35746678 | 0.01610059 | up |
| rno-miR-511-5p_R+1 | 2.43091001 | 1.28149649 | 0.01971394 | up |
| rno-let-7c-1-3p | 2.38584467 | 1.25450012 | 0.01280495 | up |
| rno-let-7c-2-3p_1ss22CT | 2.30515283 | 1.20486241 | 0.00268888 | up |
| rno-miR-125b-1-3p | 2.22498201 | 1.15379367 | 0.00549226 | up |
| PC-3p-335_45589 | 2.22025883 | 1.15072787 | 0.01938865 | up |

| | | | | |
|---|---|---|---|---|
| rno-miR-3473_1ss22AC | 2.21778122 | 1.14911706 | 0.02223895 | up |
| rno-miR-199a-5p | 2.14623824 | 1.10181023 | 0.01833651 | up |
| pal-miR-9226-5p_L-4 | 2.13476929 | 1.09408016 | 0.01677519 | up |
| pal-miR-9226-5p | 2.1238051 | 1.08665138 | 0.01529116 | up |
| rno-miR-216a-5p | 2.11293069 | 1.07924545 | 0.00528854 | up |
| rno-miR-92b-5p | 2.05608222 | 1.03989796 | 0.010545 | up |
| mmu-miR-16-1-3p_R+1 | 2.01327738 | 1.00954595 | 0.00358775 | up |
| rno-mir-326-p5 | 0.48530472 | 1.04303721 | 0.04277931 | down |
| rno-miR-31a-3p | 0.47908164 | -1.06165657 | 0.02647372 | down |
| rno-miR-133b-3p_R-1 | 0.47232876 | -1.0821367 | 0.02445962 | down |
| pal-mir-9226-p3_1ss18CA | 0.44423415 | -1.1706078 | 0.03786521 | down |
| sha-miR-125a_R+2_2 | 0.43916184 | -1.18717539 | 0.01786443 | down |
| sha-miR-125a_R+2_1 | 0.43916184 | -1.18717539 | 0.01786443 | down |
| mmu-miR-6412_R-2_1ss15AT | 0.41152071 | -1.28096305 | 0.02765388 | down |
| rno-miR-3585-3p_R+3 | 0.41135126 | -1.28155723 | 0.02476466 | down |
| hsa-miR-320b_R+1 | 0.38787994 | -1.36631791 | 0.00485201 | down |
| rno-miR-101a-5p | 0.37928456 | -1.39864747 | 0.01947404 | down |
| mmu-miR-1264-3p | 0.28153017 | -1.82863858 | 0.03759713 | down |
| PC-3p-11628_280 | 0.2534498 | -1.9802281 | 0.01408585 | down |
| rno-miR-3585-5p | 0.24517095 | -2.02814005 | 0.02429082 | down |
| rno-miR-509-3p | 0.21079162 | -2.2461106 | 0.00675018 | down |
| rno-miR-547-5p | 0.20143024 | -2.31164779 | 0.02410984 | down |
| mmu-miR-1298-3p | 0.18699022 | -2.41896528 | 0.00469684 | down |
| rno-miR-208a-5p | 0.14448432 | -2.79101517 | 0.02055638 | down |
| bta-miR-150_1ss8AC | 0.12233562 | -3.03108361 | 0.02878932 | down |
| PC-3p-11106_298 | 0.10007367 | -3.32086568 | 0.00038261 | down |
| hsa-miR-378g | 0.08234654 | -3.60214821 | 0.00796135 | down |
| mdo-miR-497-5p_R+2 | 0.07543527 | -3.7286169 | 0.03283918 | down |

**Table 6. Male BLEO vs Male SHAM microRNA-seq**
Significantly Differentially Expressed microRNAs ($log_2$FC |1|, pvalue < 0.05)

| miR name | fc | $log_2$(fc) | pvalue | regulation |
|---|---|---|---|---|
| rno-miR-212-3p | 43.19925112 | 5.432934398 | 0.014839216 | up |
| rno-miR-212-5p_R-1 | 24.57043705 | 4.618851615 | 0.014398038 | up |
| rno-miR-132-5p | 18.62094804 | 4.218854621 | 0.004994289 | up |
| mmu-mir-6240-p5_1ss13TG | 17.18796105 | 4.103326508 | 0.01568818 | up |
| rno-miR-132-3p | 15.44565375 | 3.949129031 | 0.021548239 | up |
| rno-miR-466c-5p_R+1 | 11.74610983 | 3.554111127 | 0.006441552 | up |
| rno-miR-18a-3p | 11.53038286 | 3.527368513 | 0.004272948 | up |
| cja-miR-9933_L-3_1ss12AT | 10.94685652 | 3.452444742 | 0.038197368 | up |
| PC-5p-5766_706 | 10.13543547 | 3.341336171 | 0.022177221 | up |
| rno-miR-296-3p | 9.295989542 | 3.216608446 | 0.040113854 | up |
| mmu-mir-6240-p3_1ss3TG_1 | 9.181558854 | 3.198739117 | 0.036557516 | up |
| bta-miR-12034_L-1_1ss19TG | 9.077353451 | 3.182271734 | 0.042353457 | up |
| rno-miR-21-5p_R+1 | 8.649549108 | 3.112624929 | 0.001382043 | up |
| mmu-mir-6240-p3_1ss19GT | 7.53356059 | 2.913331888 | 0.023328879 | up |
| rno-miR-511-3p_R+1 | 7.441801039 | 2.89565182 | 0.002080206 | up |
| mmu-mir-6240-p3_1ss3TG_2 | 7.382858295 | 2.884179468 | 0.014067825 | up |
| mmu-mir-6240-p5_1ss3TG | 7.382858295 | 2.884179468 | 0.014067825 | up |
| mmu-miR-1983 | 7.018755531 | 2.811215254 | 1.10911E-06 | up |
| mmu-miR-335-3p | 6.912439701 | 2.78919499 | 0.045859707 | up |
| rno-miR-1247-5p | 6.870185892 | 2.780349136 | 0.007794902 | up |
| mmu-miR-26a-2-3p_1ss4GA | 6.837053022 | 2.773374614 | 0.014514821 | up |
| mmu-mir-6240-p5_1 | 6.514944574 | 2.703752905 | 0.024577049 | up |
| mmu-mir-6240-p5_3 | 6.514944574 | 2.703752905 | 0.024577049 | up |
| mmu-mir-6240-p5_2 | 6.514944574 | 2.703752905 | 0.024577049 | up |
| PC-5p-8089_451 | 6.250361828 | 2.643939708 | 0.002094785 | up |
| rno-miR-298-5p_R-2 | 6.065856493 | 2.600711367 | 0.00378875 | up |
| rno-miR-21-3p | 5.933690917 | 2.568929779 | 0.007964488 | up |
| PC-5p-3418_1419 | 5.73189811 | 2.519012964 | 0.008321993 | up |
| efu-miR-9341_L-2R+1 | 5.722019905 | 2.516524516 | 0.047657389 | up |
| PC-5p-6283_632 | 5.362443665 | 2.422890586 | 0.000473704 | up |
| cgr-miR-1260_L-1R+2 | 4.824452991 | 2.270365375 | 0.017464387 | up |
| ssc-mir-1285-p3 | 4.774882874 | 2.255465345 | 0.012516029 | up |
| rno-miR-196b-5p_R-1 | 4.260974877 | 2.091183545 | 0.026465037 | up |
| rno-miR-155-5p | 4.151752443 | 2.053720422 | 0.002982933 | up |
| rno-miR-214-3p_R+1 | 4.114208793 | 2.040615011 | 0.000778236 | up |
| rno-miR-18a-5p | 4.102191068 | 2.03639469 | 0.003202402 | up |
| efu-miR-9298_R-4_1ss21TC | 4.018395226 | 2.006619466 | 0.002328538 | up |

| | | | | |
|---|---|---|---|---|
| rno-miR-147_R-1 | 3.603418519 | 1.849366223 | 0.008342977 | up |
| rno-miR-219a-5p_R+2 | 3.535971655 | 1.82210671 | 0.000268652 | up |
| PC-3p-8056_454 | 3.373792464 | 1.75437123 | 0.028984729 | up |
| mmu-miR-18b-5p | 3.287461876 | 1.716974164 | 0.028566936 | up |
| rno-miR-511-5p_R+1 | 3.257674783 | 1.703842585 | 0.028713886 | up |
| rno-miR-210-3p | 3.078014503 | 1.622000029 | 0.02959447 | up |
| rno-miR-34a-3p | 3.053822015 | 1.61061598 | 0.008773897 | up |
| rno-miR-1249 | 2.922916705 | 1.547408717 | 0.02607254 | up |
| mmu-miR-223-5p_R-1 | 2.920201297 | 1.546067821 | 0.004366378 | up |
| rno-miR-146b-3p_L+2R-1 | 2.882894146 | 1.527517865 | 0.001338883 | up |
| rno-miR-142-3p_R-1 | 2.881153277 | 1.526646414 | 0.022828949 | up |
| PC-3p-22071_118 | 2.749059108 | 1.458937927 | 0.033921315 | up |
| hsa-miR-1260b_R-1 | 2.688181021 | 1.426630292 | 0.00255065 | up |
| rno-miR-16-3p_R+1 | 2.654435122 | 1.408404881 | 0.001525358 | up |
| rno-miR-877_R+4 | 2.612985582 | 1.385699164 | 0.02314789 | up |
| rno-miR-190b-5p_R+2 | 2.580425082 | 1.367608745 | 0.009846886 | up |
| rno-miR-3473_1ss22AC | 2.521863676 | 1.33449029 | 0.009676355 | up |
| rno-miR-296-5p | 2.520780475 | 1.333870485 | 0.042522173 | up |
| mml-miR-145-5p_R+2_1ss24TA | 2.448128215 | 1.291679118 | 0.007659826 | up |
| rno-miR-501-3p_R-1 | 2.437938245 | 1.285661582 | 0.000221426 | up |
| mmu-miR-147-5p_R-2_1ss9TC | 2.432420278 | 1.282392522 | 0.015519314 | up |
| rno-miR-344a-3p_1ss21TC | 2.407508432 | 1.267540851 | 0.045085802 | up |
| rno-let-7c-2-3p | 2.349647705 | 1.232444462 | 0.008096842 | up |
| mmu-miR-574-5p | 2.300701511 | 1.202073823 | 0.007333168 | up |
| rno-miR-362-5p | 2.292490929 | 1.196916025 | 0.000304949 | up |
| rno-miR-138-5p | 2.290642195 | 1.195752123 | 0.003791169 | up |
| rno-miR-20a-5p | 2.288362341 | 1.194315508 | 0.011573568 | up |
| rno-let-7f-1-3p_R+1 | 2.266116119 | 1.180221789 | 0.041490844 | up |
| rno-miR-130b-5p | 2.244236936 | 1.166224997 | 0.000385771 | up |
| mmu-miR-214-5p | 2.222966498 | 1.152486206 | 0.008423554 | up |
| rno-miR-9a-5p | 2.184377039 | 1.127221897 | 0.018877325 | up |
| rno-miR-324-5p | 2.168371478 | 1.116611935 | 0.042731804 | up |
| eca-miR-450c | 2.144861176 | 1.100884274 | 0.042312633 | up |
| mmu-miR-16-1-3p_R+1 | 2.129258324 | 1.09035099 | 0.00766553 | up |
| rno-miR-378a-5p | 2.116992818 | 1.082016375 | 0.014461334 | up |
| rno-miR-199a-5p | 2.11077403 | 1.077772138 | 0.000212665 | up |
| sha-mir-199a-p5 | 2.077466401 | 1.054825145 | 0.026381221 | up |
| rno-let-7c-2-3p_1ss22CT | 2.077131682 | 1.05459268 | 0.007241516 | up |
| mdo-miR-200b-3p_R+2 | 0.495344344 | -1.013496315 | 0.021686566 | down |
| rno-miR-181c-3p_R-1 | 0.483068135 | -1.049701403 | 0.000573804 | down |
| rno-miR-200a-3p | 0.480556111 | -1.0572232 | 0.000599295 | down |

| | | | | |
|---|---|---|---|---|
| rno-miR-652-5p_L-1R+2 | 0.471901068 | -1.083443659 | 0.01418145 | down |
| rno-miR-26a-5p | 0.467474082 | -1.097041714 | 0.000173302 | down |
| oga-miR-28b_R+3 | 0.465341133 | -1.103639379 | 0.009645616 | down |
| rno-miR-10a-3p_R-1 | 0.464972709 | -1.104782052 | 0.001835606 | down |
| mdo-miR-26-5p_R+1_1ss10TC | 0.458543458 | -1.124869626 | 0.002529974 | down |
| rno-miR-224-5p | 0.451345233 | -1.147696725 | 0.0005857 | down |
| rno-miR-200b-5p | 0.449442436 | -1.153791747 | 0.004417381 | down |
| rno-miR-181c-5p | 0.448637477 | -1.156377952 | 0.000248289 | down |
| rno-miR-322-3p_R+1 | 0.443428898 | -1.173225301 | 5.47338E-05 | down |
| mmu-miR-6412_R-2_1ss15AT | 0.440278669 | -1.183511147 | 0.010660078 | down |
| rno-miR-201-3p | 0.433619566 | -1.205498238 | 0.00914053 | down |
| rno-miR-30c-5p | 0.428848996 | -1.221458353 | 0.000156842 | down |
| rno-miR-200a-5p_R+1 | 0.420423508 | -1.250084754 | 0.004276807 | down |
| rno-miR-96-5p_R-2 | 0.393020132 | -1.347324879 | 0.001127222 | down |
| rno-miR-330-3p_L-1 | 0.358112323 | -1.481515931 | 8.87089E-05 | down |
| mmu-miR-486b-5p_R+2 | 0.354616115 | -1.495669996 | 0.017197945 | down |
| ssc-miR-30a-3p_L-1R+2 | 0.32467308 | -1.622940324 | 0.002024929 | down |
| rno-miR-144-5p | 0.316853945 | -1.658110115 | 0.002003568 | down |
| PC-5p-15155_198 | 0.306048564 | -1.708167497 | 0.007735309 | down |
| PC-3p-11628_280 | 0.303900904 | -1.718327127 | 0.008423544 | down |
| rno-miR-547-3p | 0.303439938 | -1.720517114 | 0.000163537 | down |
| rno-miR-496-3p | 0.302261472 | -1.726130998 | 0.034783392 | down |
| rno-miR-1298 | 0.285424599 | -1.808818416 | 0.005846777 | down |
| rno-miR-3585-3p_R+3 | 0.279899686 | -1.837018226 | 0.000647087 | down |
| tch-miR-26a-2-5p_R-1 | 0.27134044 | -1.881824013 | 0.010987563 | down |
| mmu-miR-1264-3p | 0.267914819 | -1.900153714 | 0.009587792 | down |
| rno-miR-509-3p | 0.262293974 | -1.930743433 | 0.006136724 | down |
| bta-miR-342_R-3 | 0.24858504 | -2.008188617 | 0.013736049 | down |
| rno-miR-3547_R-1_1ss22AT | 0.191942772 | -2.381251864 | 0.011263533 | down |
| rno-miR-3585-5p | 0.185933497 | -2.427141387 | 0.001168127 | down |
| rno-miR-672-3p | 0.182113218 | -2.457092457 | 0.000256459 | down |
| rno-miR-201-5p_R+1 | 0.164645851 | -2.60256194 | 0.000457723 | down |
| mmu-miR-1298-3p | 0.10945582 | -3.191579433 | 0.000492788 | down |
| rno-miR-547-5p | 0.024968574 | -5.323742769 | 0.000203999 | down |

**Table 7. Female BLEO vs Male BLEO microRNA-seq**

Significantly Differentially Expressed microRNAs ($log_2$FC |1|, pvalue < 0.05)

| miR name | fc | $log_2$(fc) | pvalue | regulation |
|---|---|---|---|---|
| rno-miR-672-3p | 4.635953 | 2.212865932 | 0.005896349 | up |
| rno-miR-201-5p_R+1 | 3.291717 | 1.71884038 | 0.009172579 | up |
| rno-miR-144-5p | 2.563168 | 1.357927815 | 0.008431589 | up |
| tch-miR-26a-2-5p_R-1 | 2.491648 | 1.317100153 | 0.03228511 | up |
| rno-miR-1298 | 2.470352 | 1.304716798 | 0.034410408 | up |
| pal-miR-9226-5p_1ss4AG | 2.379203 | 1.250478346 | 0.032505146 | up |
| rno-miR-144-3p | 2.333038 | 1.222210016 | 0.015271201 | up |
| rno-miR-486 | 2.307901 | 1.206581382 | 0.003506299 | up |
| PC-5p-8089_451 | 0.49468 | -1.015432162 | 0.015030397 | down |
| rno-miR-1247-5p | 0.487472 | -1.036608492 | 0.042051179 | down |
| rno-miR-702-3p_R+2 | 0.483367 | -1.048809549 | 0.001398079 | down |
| rno-miR-1306-5p_R-1 | 0.45965 | -1.121392948 | 0.010479871 | down |
| PC-5p-6283_632 | 0.457362 | -1.128590974 | 0.034404265 | down |
| ssc-mir-1285-p3 | 0.45246 | -1.144138275 | 0.047044882 | down |
| rno-miR-18a-5p | 0.446253 | -1.164065213 | 0.009420722 | down |
| rno-miR-21-5p_R+1 | 0.415532 | -1.266969154 | 0.00340876 | down |
| rno-miR-214-3p_R+1 | 0.411489 | -1.28107258 | 0.00092305 | down |
| rno-miR-29a-5p_R-1 | 0.408505 | -1.291573607 | 0.028685297 | down |
| PC-3p-22071_118 | 0.403353 | -1.309886111 | 0.042727435 | down |
| rno-miR-210-3p | 0.394803 | -1.340794823 | 0.041155098 | down |
| mmu-miR-1983 | 0.389076 | -1.361875976 | 0.002994415 | down |
| rno-miR-511-3p_R+1 | 0.385715 | -1.37439312 | 0.006243365 | down |
| rno-miR-147_R-1 | 0.380301 | -1.394784923 | 0.013527595 | down |
| hsa-miR-1260b_R-1 | 0.371776 | -1.427495255 | 0.021142862 | down |
| rno-miR-215 | 0.355853 | -1.490647497 | 0.037233332 | down |
| rno-miR-384-5p | 0.345955 | -1.531344827 | 0.022605062 | down |
| mmu-miR-147-5p_R-2_1ss9TC | 0.342908 | -1.544106219 | 0.011449462 | down |
| rno-miR-138-5p | 0.342884 | -1.544207733 | 0.000601335 | down |
| mmu-miR-26a-2-3p_1ss4GA | 0.323551 | -1.627934623 | 0.029862742 | down |
| rno-miR-1249 | 0.322437 | -1.632912236 | 0.020977354 | down |
| rno-miR-196b-5p_R-1 | 0.320523 | -1.641501216 | 0.049881086 | down |
| mmu-miR-18b-5p | 0.317776 | -1.653916124 | 0.030859927 | down |
| rno-miR-325-3p_R-1 | 0.31679 | -1.658402256 | 0.047119873 | down |
| rno-miR-344a-3p_1ss21TC | 0.305382 | -1.711314247 | 0.020103834 | down |
| rno-miR-298-5p_R-2 | 0.302507 | -1.724959258 | 0.006973795 | down |
| rno-miR-325-5p_R-2 | 0.280536 | -1.833744346 | 0.016120791 | down |
| rno-miR-212-5p_R-1 | 0.264625 | -1.917980607 | 0.030822753 | down |
| rno-miR-132-5p | 0.260811 | -1.938921516 | 0.008801335 | down |

| | | | | |
|---|---|---|---|---|
| rno-miR-335 | 0.258202 | -1.953427756 | 0.029454922 | down |
| PC-5p-5766_706 | 0.237649 | -2.073094806 | 0.036228711 | down |
| rno-miR-466c-5p_R+1 | 0.224123 | -2.157638474 | 0.008207132 | down |
| rno-miR-212-3p | 0.195501 | -2.354753176 | 0.026262325 | down |
| rno-miR-490-5p | 0.189134 | -2.40251671 | 0.033926992 | down |
| hsa-mir-636-p5_1ss18CG | 0.175077 | -2.513939825 | 0.026625838 | down |
| mmu-miR-3473b_R-2 | 0.169748 | -2.558536978 | 0.019902098 | down |
| mmu-miR-3473e_R-3 | 0.169748 | -2.558536978 | 0.019902098 | down |
| mmu-miR-335-3p | 0.169502 | -2.560626556 | 0.049844368 | down |
| PC-3p-41105_46 | 0.164315 | -2.605466236 | 0.037850693 | down |
| cgr-miR-1260_L-1R+2 | 0.143925 | -2.796608625 | 0.002955917 | down |
| PC-3p-29036_79 | 0.12776 | -2.968495891 | 0.032920531 | down |
| PC-5p-36326_56 | 0.096269 | -3.376790375 | 0.000796845 | down |
| rno-miR-466b-5p_L+1R+1 | 0.086564 | -3.530083681 | 0.04055669 | down |
| rno-miR-296-5p | 0.059165 | -4.079120453 | 0.00807475 | down |
| rno-miR-1247-3p | 0.056737 | -4.139554996 | 0.023241937 | down |

**Table 8. Female SHAM vs Male SHAM microRNA-seq**

Significantly Differentially Expressed microRNAs ($log_2FC$ |1|, pvalue < 0.05)

| miR name | fc | log₂(fc) | pvalue | regulation |
|---|---|---|---|---|
| PC-3p-11106_298 | 9.735779193 | 3.283296448 | 0.000570847 | up |
| hsa-miR-320d | 7.777960875 | 2.959391978 | 0.031571062 | up |
| rno-miR-101b-3p_R-2_1ss19GT | 2.828254111 | 1.499911748 | 0.048902334 | up |
| rno-miR-29b-3p | 2.278445887 | 1.188050107 | 0.003379146 | up |
| hsa-miR-320b | 2.185161696 | 1.127740039 | 0.029726952 | up |
| rno-miR-101a-5p | 2.059066624 | 1.041990511 | 0.004987911 | up |
| rno-miR-466c-5p_R+1_1ss18CT | 0.462270843 | -1.113189724 | 0.044418693 | down |
| rno-miR-130b-3p | 0.417080264 | -1.261603048 | 0.021320053 | down |
| hsa-miR-4286_R+1 | 0.413966041 | -1.272415672 | 0.023676267 | down |
| rno-miR-335 | 0.411624906 | -1.280597819 | 0.020918294 | down |
| mmu-miR-335-3p | 0.393505114 | -1.345545711 | 0.019359201 | down |
| mmu-miR-6516-3p_L+1R-1 | 0.373804399 | -1.419644549 | 0.024915007 | down |
| rno-miR-449c-5p_R+4 | 0.364584298 | -1.455675668 | 0.016784247 | down |
| rno-miR-449a-5p | 0.347505269 | -1.524893243 | 0.006821009 | down |
| rno-miR-26a-3p_R+1 | 0.322627463 | -1.632058845 | 0.039107616 | down |
| mmu-miR-449b_R+3_2ss10-G11-T | 0.278402609 | -1.844755362 | 0.00327248 | down |
| rno-mir-3102-p5 | 0.261911857 | -1.932846723 | 0.045549841 | down |
| rno-miR-122-5p_R+1 | 0.216743311 | -2.205940623 | 0.022049491 | down |
| rno-miR-9a-3p_L-1 | 0.153615391 | -2.702605323 | 0.012873902 | down |
| rno-miR-449a-3p | 0.132942465 | -2.911126088 | 0.034123429 | down |
| mmu-mir-449b-p3 | 0.132727701 | -2.913458596 | 0.037177681 | down |
| mmu-mir-449b-p5 | 0.132727701 | -2.913458596 | 0.037177681 | down |
| hsa-mir-4454-p5_1ss19AT | 0.075391937 | -3.729445947 | 0.049998782 | down |

**Table 9. List of geneglobe ID of the individual miRNA PCR primer set from Qiagen miRCURY LNA miRNA PCR assays.**

| microRNA | Accession | Sequence (5') | Geneglobe ID |
|---|---|---|---|
| U6 snRNA (v2) | - | - | YP02119464 |
| rno-miR-672-3p | MIMAT0017312 | ACACACAGTCGCCATCTTCGA | YP02106349 |
